# Efficient secretion of a plastic degrading enzyme from the green algae *Chlamydomonas reinhardtii*

**DOI:** 10.1101/2024.10.23.619606

**Authors:** João Vitor Dutra Molino, Barbara Saucedo, Kalisa Kang, Chloe Walsh, Crisandra Jade Diaz, Marissa Tessman, Stephen Mayfield

**Affiliations:** Division of Biological Sciences, University of California San Diego, La Jolla, California, United States of America; Algenesis Inc., 1238 Sea Village Dr., Cardiff, CA, United States of America

## Abstract

Plastic pollution has become a global crisis, with microplastics contaminating every environment on the planet, including our food, water, and even our bodies. In response, there is a growing interest in developing plastics that biodegrade naturally, thus avoiding the creation of persistent microplastics. As a mechanism to increase the rate of polyester plastic degradation, we examined the potential of using the green microalga *Chlamydomonas reinhardtii* for the expression and secretion of PHL7, an enzyme that breaks down post-consumer polyethylene terephthalate (PET) plastics. We engineered *C. reinhardtii* to secrete active PHL7 enzyme and selected strains showing robust expression, by using agar plates containing a polyester polyurethane (PU) dispersion as an efficient screening tool. This method demonstrated the enzyme’s efficacy in degrading ester bond-containing plastics, such as PET and bio-based polyurethanes, and highlights the potential for microalgae to be implemented in environmental biotechnology. The effectiveness of algal-expressed PHL7 in degrading plastics was shown by incubating PET with the supernatant from engineered strains, resulting in substantial plastic degradation, confirmed by mass spectrometry analysis of terephthalic acid (TPA) formation from PET. Our findings demonstrate the feasibility of polyester plastic recycling using microalgae to produce plastic-degrading enzymes. This eco-friendly approach can support global efforts toward eliminating plastic in our environment, and aligns with the pursuit of low-carbon materials, as these engineered algae can also produce plastic monomer precursors. Finally, this data demonstrates *C. reinhardtii* capabilities for recombinant enzyme production and secretion, offering a “green” alternative to traditional industrial enzyme production methods.

**Graphical Abstract:** 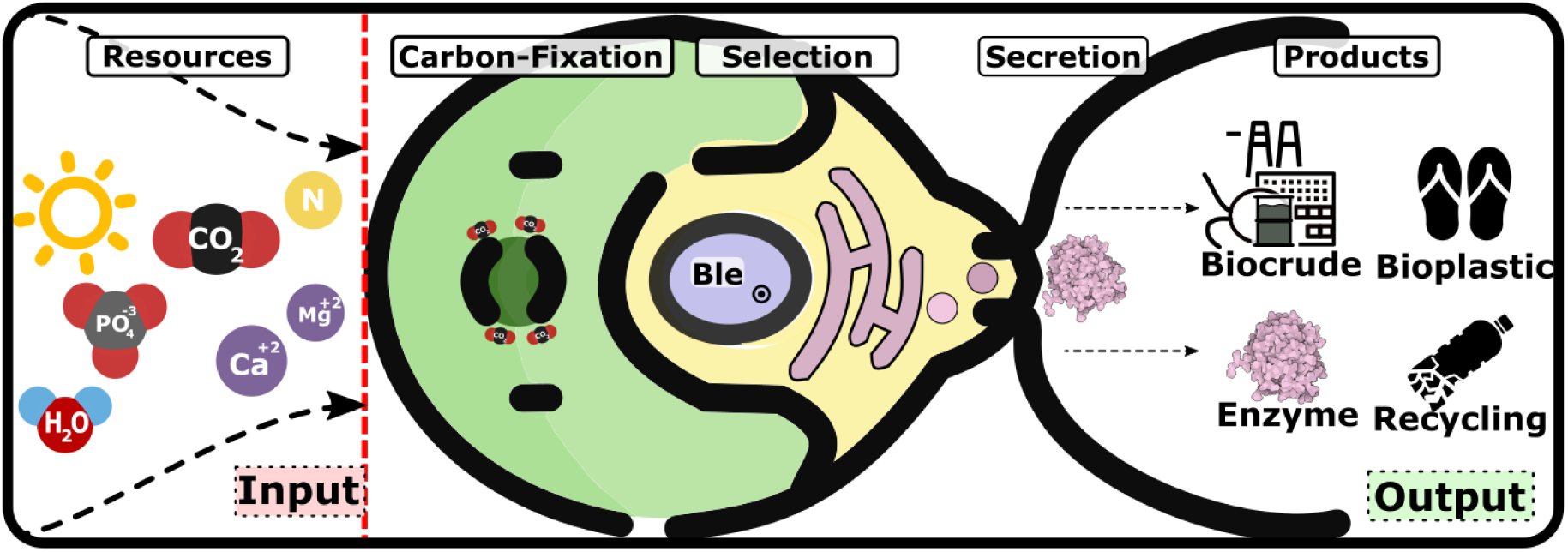

## Introduction

In the 21st century, transitioning to a climate-neutral economy is an urgent and critical challenge, demanding innovative solutions across all facets of society. Biotechnology, especially through the utilization of microalgae, has the potential to play a vital role in this transformation. Microalgae offer substantial environmental benefits, such as replacing fossil-based products with products that offer significantly reduced greenhouse gas emissions (Gupta et al., 2024; Moody et al., 2014). As a sustainable source for bioplastic production, microalgae provide a renewable alternative to petroleum, and contribute to CO_2_ sequestration, addressing two major environmental concerns. However, these microalgal production processes must overcome economic and scaling barriers to realize their full potential and achieve broader adoption. This includes making them cost-competitive with traditional methods and ensuring their cultivation and processing are environmentally sustainable and efficient (Sari et al., 2021). By addressing these challenges, microalgae can become a cornerstone of a more sustainable and climate-resilient future.

Petroleum-based plastics contribute to significant environmental degradation, not only through direct greenhouse gas emissions during production, but also by causing widespread plastic pollution after their useful life (MacLeod et al., 2021). Plastics presently account for about 4.5% of global greenhouse gas emissions, with projections showing a potential increase in emissions if current trends continue (Cabernard et al., 2021). The production of plastic has grown exponentially over the past 70 years, from just two million tons in 1950 to over 450 million tons today, attesting to the widespread use of plastics in many sectors (Ritchie et al., 2023). This increase in production has led to significant environmental challenges, reflected in recycling rates of only 5-6% in 2021, actually down from the 8.7% rate of recycling just three years prior (Greenpeace, 2022). The rest is either landfilled, incinerated, or mismanaged, with a significant portion ending up in natural environments. Unfortunately, between 4.8 and 12.7 million tons of this plastic enter the oceans yearly, contributing to the growing problem of marine pollution (*The Environmental Impacts of Plastics and Micro-Plastics Use, Waste and Pollution*, 2020). This influx of plastics in the marine ecosystem is alarming, as it not only affects marine life but also poses a threat to human health through bioaccumulation in the food chain (Leslie et al., 2022), since it ultimately breaks down into micro- and nano-plastics that can persist for hundreds of years (*The Environmental Impacts of Plastics and Micro-Plastics Use, Waste and Pollution*, 2020). The continued accumulation of plastics is projected to have long-lasting impacts, with some estimates suggesting that by 2050, the mass of plastics in the ocean could surpass that of the mass of fish (*The Environmental Impacts of Plastics and Micro-Plastics Use, Waste and Pollution*, 2020)

Various strategies have been pursued to mitigate the pervasive issue of plastic pollution. These include bans on single-use plastics like plastic bags, straws, and utensils, and promoting reusable alternatives such as metal water bottles and fabric shopping bags. Improving waste collection and recycling systems can also play a crucial role in diverting plastic waste from landfills and oceans (Hasan et al., 2023). Given plastic’s essential role in applications like food spoilage prevention and general consumer goods packaging (Heller et al., 2019; Verghese et al., 2015), eliminating plastics any time soon seems unlikely. However, even today, there are alternatives such as bioplastics, which are plastics derived from bio-based materials. These bio-plastics offer potentially much more environmentally favorable profiles and some of these materials have successfully scaled up to industrial production. (European Commission. Directorate General for Research and Innovation. et al., 2021).

Nonetheless, these options still contribute to greenhouse gas emissions, mainly due to the feedstock production chain and competition with food crops for arable land, water, and energy (Richard Platt, 2021). Some bioplastics are chemically identical to their petroleum-based counterparts, posing similar end-of-life environmental problems (Prieto, 2016). As a result, there is increasing interest in creating bio-based, sustainably sourced plastics that naturally biodegrade, helping to prevent the formation of long-lasting microplastics (Allemann et al., 2024). In this context, microalgae have been explored as a renewable source of bio-crude, capitalizing on features like non-arable land requirements, high productivity per area, and scaling feasibility (Tang et al., 2020). Microalgae capture carbon dioxide during growth and can utilize that carbon to generate biomaterials for plastic production. At scale, it can potentially align with current plastic industry pricing (Beckstrom et al., 2020). The emergence of biodegradable consumer products, such as sneakers made from algae-derived oil, showcases the readiness of this technology for bioplastic production (Beckstrom et al., 2020). To advance the competitiveness and adoption of microalgal bioplastics, it is essential to prioritize cost reduction measures, leverage synergistic byproduct production for increased revenue, and optimize economic efficiency—all while preserving positive environmental impacts.

In this study, we engineered a strain of *Chlamydomonas reinhardtii* to efficiently secrete PHL7, an enzyme capable of degrading post-consumer polyethylene terephthalate (PET) plastics (Sonnendecker et al., 2021). The degradation generates PET monomers that can be recycled into new PET plastics (Tournier et al., 2020). As photosynthetic organisms, microalgae efficiently convert light and carbon dioxide into valuable biomass and bioactive compounds, supporting a closed-loop system from cradle to grave, thereby minimizing environmental impact. Its rapid growth and scalability further underscore its suitability for large-scale cultivation. Extracellular secretion of PHL7 simplifies protein purification and eliminates the need for costly and labor-intensive cell lysis processes. This feature significantly enhances the practicality and economic viability of using microalgae for bioplastic production and other biomass conversion applications.

We envision two promising approaches using our microalgae system. The first involves a single-step process, where algal growth and plastic degradation co-occur within the same medium, providing a streamlined solution for recycling plastic monomers. The second approach is a bifurcated method, separating the algal growth phase from the plastic degradation process, which allows for specialized optimization of each step. These methodologies open new avenues in biotechnology, presenting microalgae as a dual-purpose solution for mitigating plastic pollution while contributing to more sustainable and eco-conscious production of bioplastic feedstock.

## Results

We evaluated the capacity of *C. reinhardtii* CC-1690 for secreting plastic degrading enzymes, using the vector pJP32 (Molino et al., 2018) that employs the *ble* gene as a selection marker and contains a signal peptide to direct an associated recombinant enzyme for secretion. . The recombinant enzyme gene was fused to *ble* gene with a self-cleavage peptide (FMDV-2A) sequence, followed by the signal peptide SP7 sequence, used for targeting the protein to the ER for secretion (Figure 1A). This signal peptide is from a cell wall protein (SAD1p) (Ferris et al., 2010). The plastic degrading enzyme studied is PHL7, an enzyme isolated from a compost metagenome and identified for its potential to break down and recycle polyester plastics, specifically PET (Sonnendecker et al., 2021). To identify colonies capable of secreting PHL7, we employed zeocin selection plates containing Impranil® (Bayer Corporation, Germany), a polyester polyurethane polymer suspension, in two concentrations: 0.5% and 0.75% (v/v). Colonies on these plates indicated the successful incorporation of the vector conferring resistance to zeocin. The capacity to secrete the plastic enzyme was observed by the formation of halos (transparent clearing zones in the opacity generated by Impranil®) around the colonies, indicating both secretion and activity of the enzyme in the area around the colonies. We performed three independent biological transformations and recorded the number of colonies and halos (Table 1).

**Figure 1:**
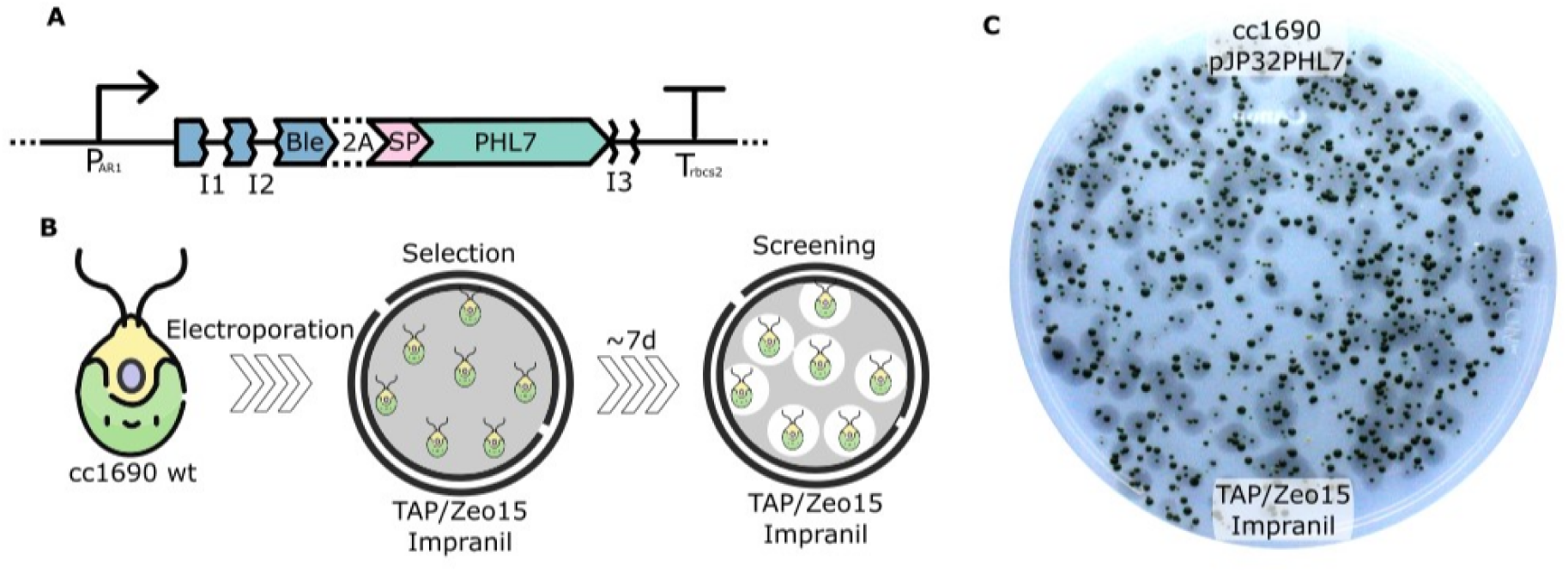
Overview of the vector design, the transformation workflow, and an experimental result. A: Schematic representation of the vector used, including the chimeric Par1 promoter, bleomycin resistance gene, F2A auto-cleavable peptide, SP7 signal peptide, *rbcs2* introns, and the *rbcs2* terminator region. B: Workflow for generating transformants with halos. C: A typical result of the transformed cells with halos, indicating successful expression and secretion of the target protein as designed in the vector. Selection on TAP media plates containing zeocin 15 µg/mL and Impranil® DLN at 0.5% (v/v).

**Table 1:** Summary of the number of colonies obtained and the number of colonies with clearing zones.

| Construct | Impranil® DLN % | #Colonies | #Halos | %Halos |
| --- | --- | --- | --- | --- |
| pJP32PHL7 | 0.5 | 5110 | 1255 | 24.6 |
| pJP32PHL7 | 0.75 | 1904 | 178 | 9.3 |

**\*Note:** Selection plates contain zeocin at 15ug/mL and Impranil® DLN DLN. The colonies came from three independent transformations, and half of each transformation was plated on either 0.5% (v/v) or 0.75% (v/v) Impranil® DLN plates.

On plates containing 0.5% Impranil® DLN, 5110 colonies formed, of which 24.6% (1255) produced halos. On the plates with 0.75% Impranil® DLN only 1904 colonies appeared, with 9.3% (178) forming halos. These results confirm that *C. reinhardtii* can secrete active PHL7, as halo formation indicates (Supplementary Figure 1).

We further screened the colonies with halos in a 96-well plate setup. Strains identified as secreting PHL7 were in TAP media on 96-well plates for five days. The supernatant was recovered and ester bond cleavage activity was determined by measuring fluorescein diacetate (FDA) cleavage activity. To account for native esterase enzymes secreted during the *C. reinhardtii* life cycle (Ves-urai et al., 2021) capable of cleaving the FDA, we measured the parental wild-type strain CC-1690 in the same experiment (Figure 2). A substantial portion of transformants showed increased activity compared to the wild-type. For colonies selected from plates with 0.5% Impranil® DLN, 40.6% (28/69) had higher activity than the averaged results (plus three standard deviations) of wild-type. Plates with 0.75% Impranil® DLN presented 50.9% (27/52) of transformants that exceeded this threshold. An ANOVA followed by a Tukey post-hoc analysis showed significant differences in activity levels between the wild-type and transformed strains, with adjusted p-values of 0.0388 and 0.0441 for 0.5% and 0.75% Impranil® DLN plates, respectively (Figure 2).

**Figure 2:**
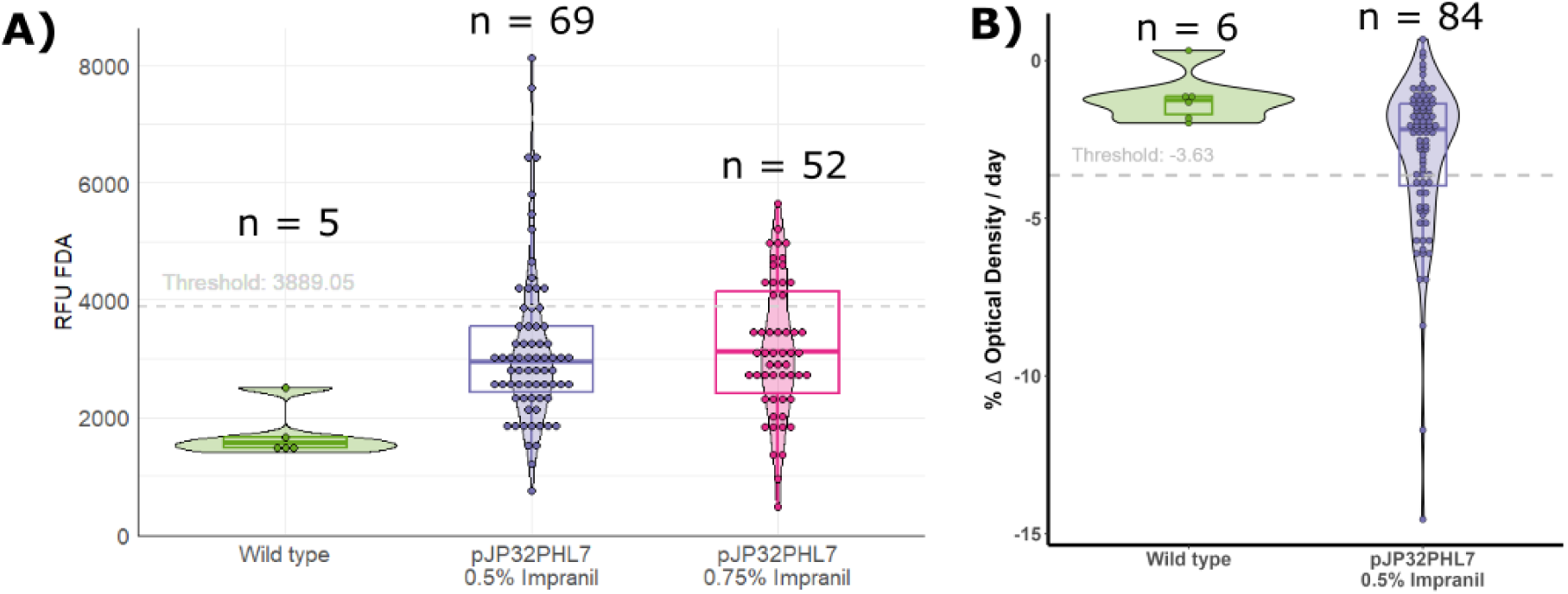
Enzymatic activity of PHL7 produced in *C. reinhardtii*. **A)** Cleavage of ester bond activity in the supernatant by Fluorescein DiAcetate (FDA) assay. B) Relative absorption reduction per day of Impranil® DLN. Wild-type cells are the parental CC1690 strains (green). pJP32PHL7 0.5% Impranil® DLN are the transformants picked from the selection plates containing zeocin 15 ug/mL and 0.5% Impranil® DLN (purple). pJP32PHL7 0.75% Impranil® DLN are the transformants picked from the selection plates containing zeocin 15 ug/mL and 0.75% Impranil® DLN (magenta). A violin plot and a box plot superimpose the bin dot plot to summarize statistics.

In addition to the FDA assay, we explored another strategy to detect enzyme activity using agarose gels supplemented with Impranil® DLN inside 96-well plates (Supplementary Figure 2). Impranil® DLN absorbs strongly in the near UV region (∼350 nm) (Supplementary Figure 3), and we monitored the decrease in absorption as a proxy for substrate degradation (Supplementary Figure 2). Both methods were functional: FDA had a quicker turnaround (∼40 min), while the Impranil® DLN-based assay required several hours (>12 h).

The Impranil® DLN activity assay revealed distinct differences in activity levels between the wild-type and pJP32PHL7 candidate colonies (Figure 2). As expected, the wild-type strain exhibited a narrow distribution of activity values, with all values clustering below the threshold line. The threshold line was calculated as three standard deviations above the mean wild-type baseline activity level. In contrast, the pJP32PHL7 colonies show a broader distribution of activities, with several values exceeding the threshold, indicating the presence of positive strains (Figure 2). This Impranil® DLN activity assay identified 30% positive transformants (25/84).

After the supernatant was used to detect enzyme activity, we blotted the 84 colony cultures onto agar plates containing 0.5% or 0.75% Impranil® DLN as a quality control step to confirm the retention of the halo-forming phenotype (Supp. Figure 4). Halo formation was observed in 81 colonies from the 0.5% plates and 83 colonies from the 0.75% plates, with the absence of halos in three colonies from the 0.5% plates likely due to sampling errors or false positives in densely populated regions on the selection plates. A time-lapse video (<u>Video 1</u>) demonstrates halo formation around cell patches on TAP agar plates containing 0.5% Impranil® DLN and 15 µg/mL zeocin, confirming PHL7 secretion and retention of bleomycin resistance.

The highest-producing strain from the screenings was selected and expanded, and its supernatant was used to run a zymogram containing Impranil® DLN (Figure 3C). We initially observed the presence of two clearing zones on the protein gels, indicating the presence of two isoforms of the enzyme being produced by *C. reinhardtii* in the pJP32PHL7 strain, which we assumed was due to post-translational modifications (PTMs) occurring in the secretory pathway. We confirmed the presence of PHL7 in both cleared regions by mass spectrometry assisted protein sequencing. We identified the presence of 9 and 11 PHL7 peptides from the top and bottom bands, respectively. *In silico* protein sequence analysis using NetN-Glyc-1.0 indicated one possible explanation for the two isoforms (Supp. Figure 5). The sequence holds three possible glycosylation sites, two of them juxtaposed, possibly explaining the two isoforms detected, one with one residual glycosylated and the other with both positions glycosylated. To confirm that glycosylation is the likely PTM being performed on PHL7, we prepared a new PHL7 version (PHL7dg), replacing the asparagine residues on the predicted glycosylation sites with aspartic acid (Supp. Figure 5).

**Figure 3:**
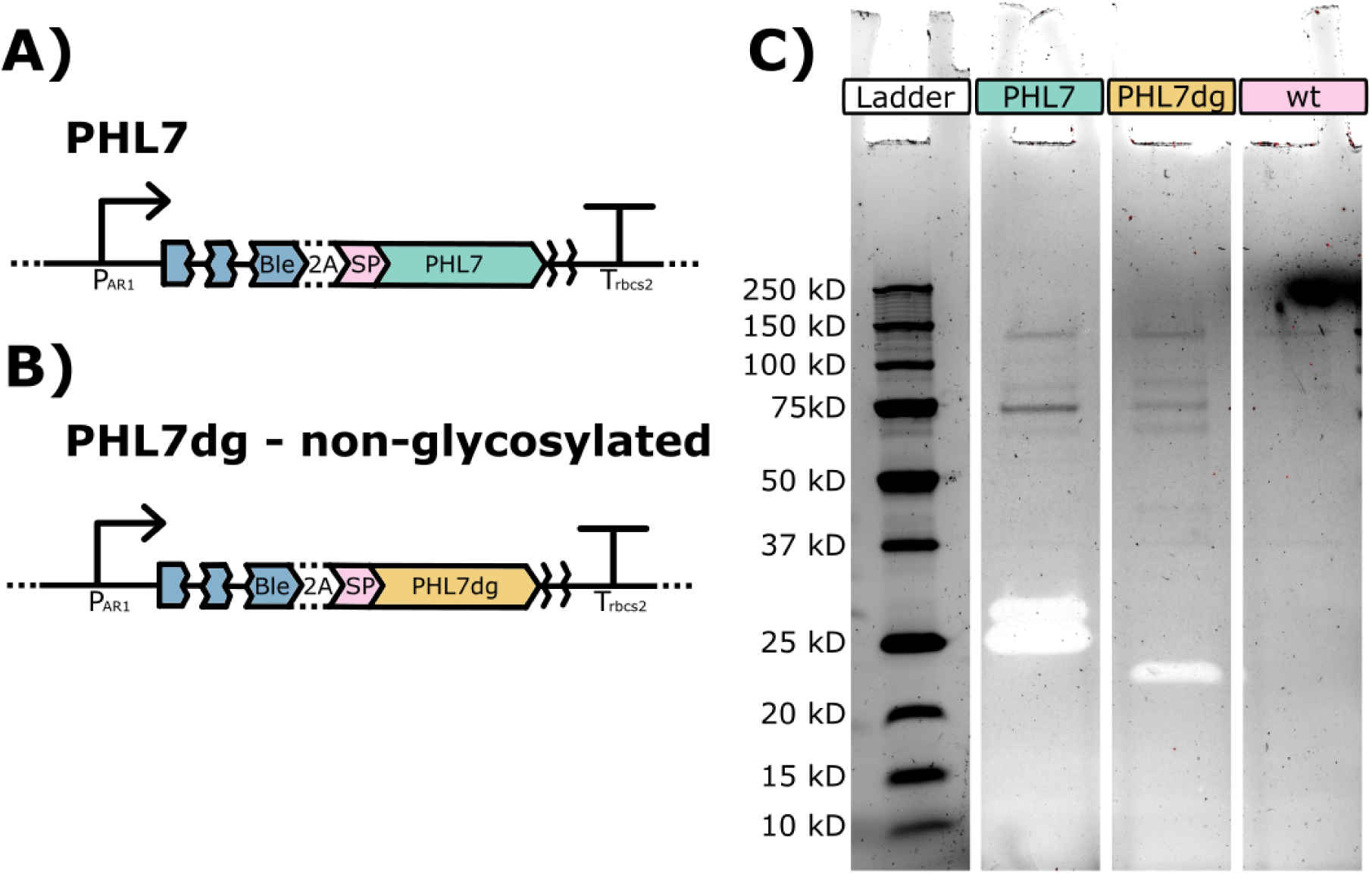
PHL7 glycosilation on the secretory pathway. A) Schematic representation of pJP32PHL7 vector corresponding to sample loaded in lane “PHL7” on zymogram. B) Schematic representation of pJP32PHL7dg (non-glycosylated PHL7) that corresponds to sample loaded in lane “PHL7dg” on zymogram. C) SDS zymogram gel with 1% v/v Impranil® DLN containing Precision Plus Protein™ Unstained Protein Standards, *Strep*-tagged recombinant (Bio-Rad Laboratories #1610363) and 10X concentrated supernatant samples from pJP32PHL7 (PHL7), pJP32PHL7dg (PHL7dg), and wild-type parental cc1690 strain (wt) (from left to right). The gel displayed 2 halos (i.e., transparent bands) in the PHL7 lane and 1 halo in the PHL7dg lane after 1 day of incubation in a 100mM potassium phosphate buffer solution, pH 8.0 at 37°C.

The new vector containing the non-glycosylated PHL7 was transformed and generated several colonies, with a few presenting halos (Supp. Figure 7). We selected one of the colonies from the single transformation event to grow in a flask. We performed a zymogram alongside the original sequence (Figure 3C). We observed a down shift in the band position in the gel, indicating an absence of the PTM observed in the original unmodified enzyme.

We compared the growth performance of the top producer pJP32PHL7 strain to the wild-type parental CC-1690 strain (Figure 4). Both strains grew similarly without any observable difference in growth rate between the wild-type and recombinant strains, with a slightly lower stationary phase density for the PHL7 strain. The secretion of PHL7 enzymes was measured by changes in OD at 350 nm due to enzymatic degradation of Impranil® DLN (Figure 4). The production curve demonstrated that the top producer pJP32PHL7 strain started to secrete detectable levels of enzymes on approximately day 5 of cultivation, coinciding with the onset of the stationary phase (Figure 4). The secretion of PHL7 enzymes exhibited a continual rise throughout the stationary phase. A decrease or stabilization in enzymatic content was not observed within the timeframe studied.

**Figure 4:**
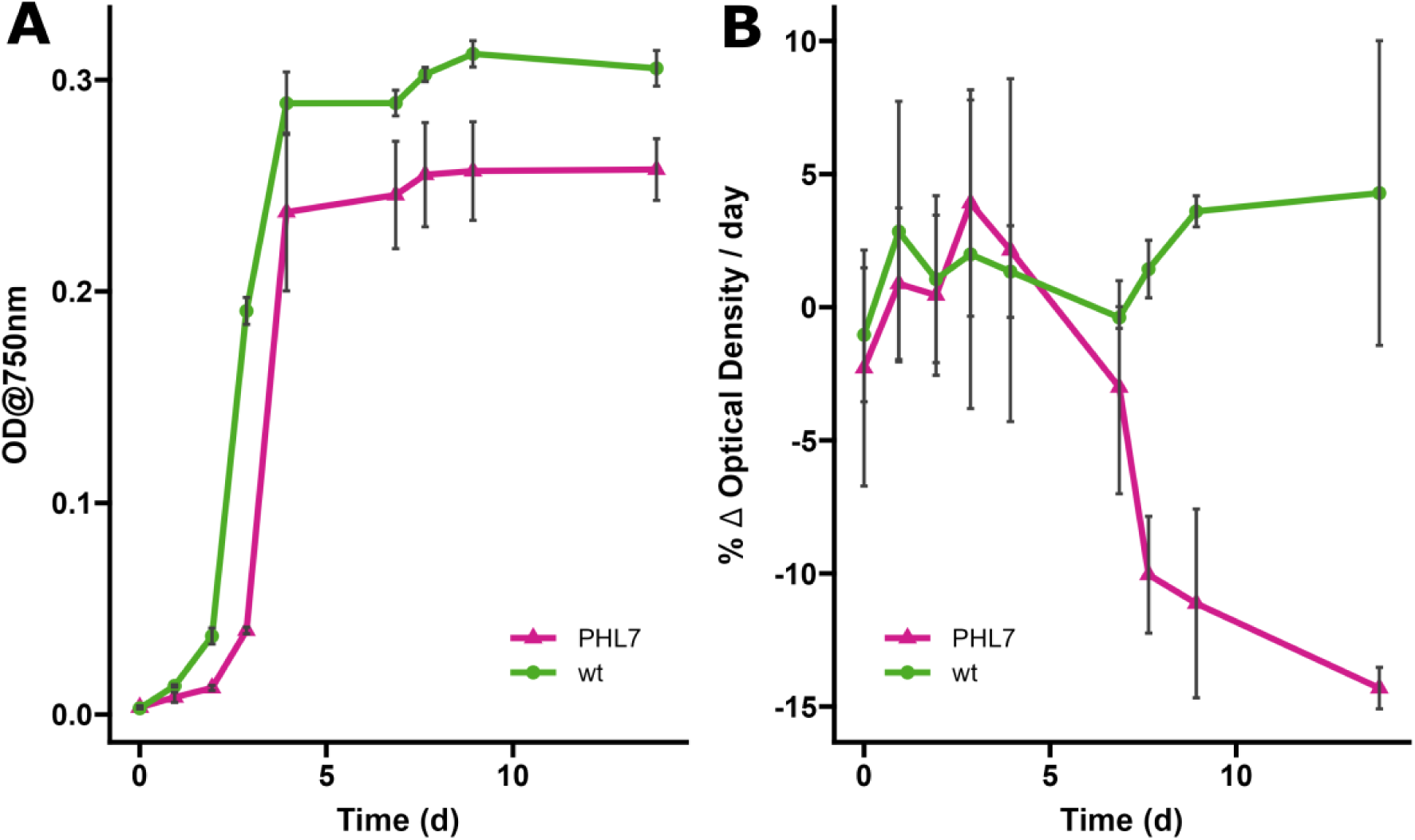
Growth curves and enzyme activity profiles of the parental line (wt) and the recombinant pJP32PHL7 (PHL7) strains over time. Panel A shows the growth curves of the wild-type (`wt`, green circles) and the mutant (`PHL7`, magenta triangles) strains, measured as optical density at 750 nm (OD750) over time. Each point represents the mean OD750 at a given time point, with vertical black error bars indicating the standard deviation across biological replicates (n = 3. Panel B depicts enzyme activity, expressed as the percentage change in optical density per day (% Δ OD/day) of Impranil® DLN, for the same strains over time. Mean enzyme activity is shown with black error bars representing the standard deviation across biological replicates (n = 3). The same symbol patterns were used, the wild-type (`wt`, green circles) and the mutant (`PHL7`, magenta triangles).

To demonstrate that PHL7 produced by *C. reinhardtii* can degrade PET plastic, in addition to degrading the Impranil polyurethane, we incubated a concentrated supernatant sample from the PHL7 strain, and a wild-type control, with approximately 30 mg of PET powder (>50% crystallinity) for seven days at 68°C in 500 mM phosphate buffer (pH 8) (Figure 5). The reaction was monitored by measuring the absorbance at 240 nm, while tracking the formation of TPA, a degradation product of PET (Sonnendecker et al., 2021). A sharp increase in absorbance was observed on the first day, followed by a steady rise over the subsequent seven days. The stark difference in values between the PHL7 strain and the wild-type control (p-value = 6.91e-07) supports the conclusion of PHL7-mediated PET degradation. To confirm the presence of TPA in the reaction media, we submitted the sample for HESI-Orbitrap analysis (LC-MS), which verified TPA’s presence (Figure 5, Supp. Figure 6).

**Figure 5:**
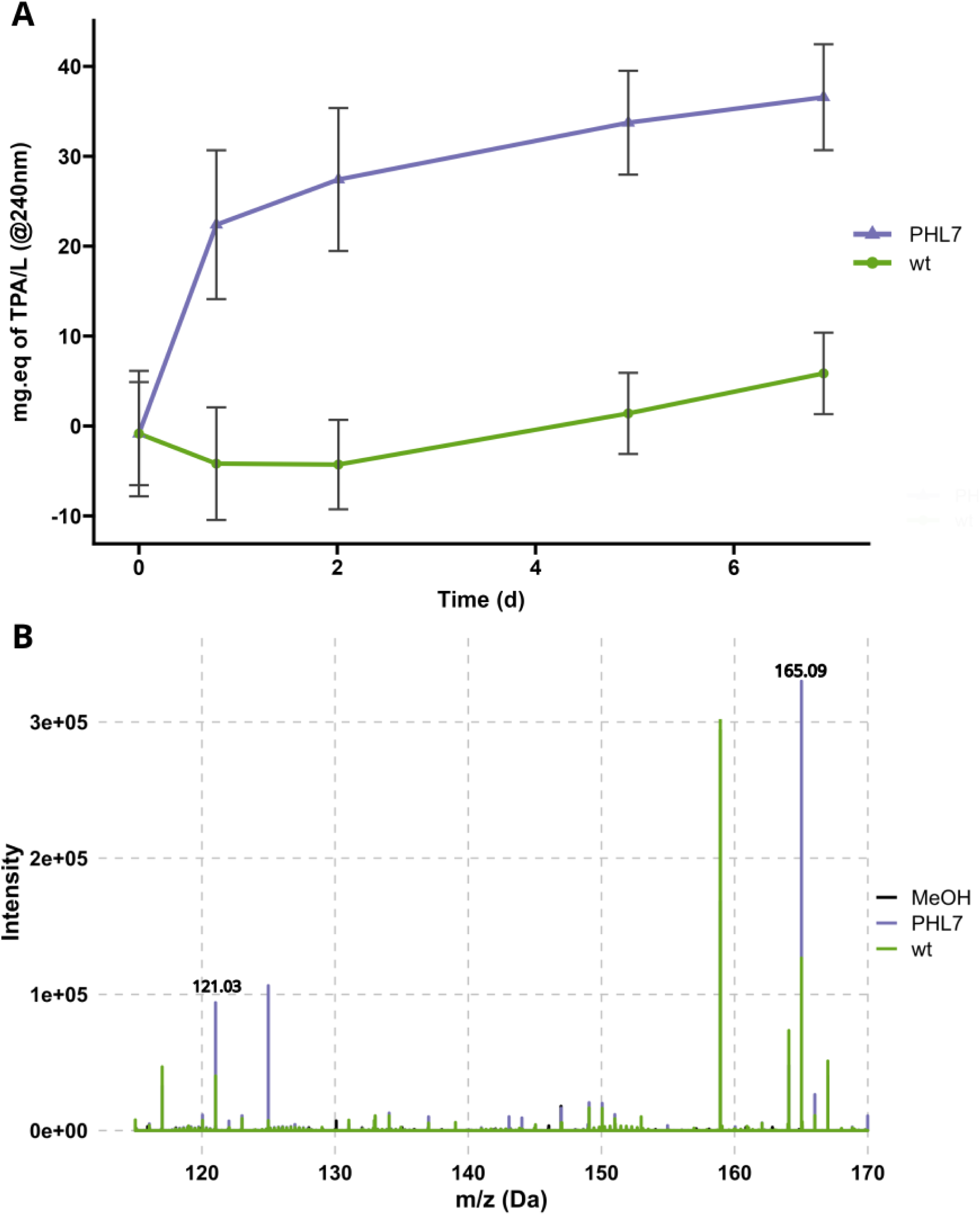
Terephthalic Acid (TPA) release during PET degradation experiment. **A)** The plot shows the TPA concentration (mg equivalent of TPA per liter, calculated by absorbance at 240 nm) over time for wild-type and PHL7 strains, measured during the enzymatic degradation of PET. The absorbance values were normalized to the initial value at time point t0, and the TPA concentration was calculated using the standard curve. Each data point represents biological replicates’ mean TPA concentration (± SD). The TPA concentration trends after day two were statistically analyzed using linear models. Strain-specific differences in TPA production were observed, with the wild type shown in green and PHL7 in purple. n = 2 biological replica, and n = 21 technical replica **B)** Mass spectrometry plot with intensity (y-axis) versus mass-to-charge ratio (m/z) for the range of 110 to 170 m/z, with two peaks highlighted corresponding to **TPA (Terephthalic acid)** in wild-type (wt) and PHL7 samples, with methanol (MeOH) as the blank.

## Discussion

Plastic pollution is a significant challenge on a global scale, with no clear resolution in sight. While plastic recycling processes are in effect today, they are expensive, cumbersome, and generally only work for downcycling, thus limiting their use (Shen & Worrell, 2024). New systems designed around a much more efficient process are required, with lower environmental impact, lower costs, and much easier to scale. One such new system could be built around the biological recycling of polyester plastics, including PET, polyurethanes, and several other polyester-based plastics. Using enzymes to depolymerize polyester plastics, ideally into their monomeric constituents, would allow for actual recycling or even upcycling into new plastic products (Li et al., 2023). This solution would require technical advances and political and social changes to support such an endeavor. Within this framework, polyester plastic recycling using enzymes is a promising strategy with a potential market demand of at least 140000 T/year of recycled PET alone (Tournier et al., 2020). Polyester polyurethanes could also be added to this recycling program at similar levels of material (Rossignolo et al., 2024).

Plastic degrading enzyme production could also be synergic with the production of microalgae bioplastic precursors, potentially displacing petroleum as a raw material supply chain while contributing to CO_2_ capture (Beckstrom et al., 2020). Implementation of this technology would achieve the benefits of providing biodegradable plastics in the market while simultaneously supplying (enzymes) for enzymatically recycling them. Enzymes are pivotal for the bioconversion of plastics, and several enzymes have been studied for plastic degradation (Bahl et al., 2021). Remarkable improvements in available PET degrading enzymes have been made through either bioprospection to identify novel enzymes from the environment (Sonnendecker et al., 2021) or enzyme engineering to improve existing enzymes (Tournier et al., 2020). These enzymes can now be produced in different recombinant systems (https://doi.org/10.5281/zenodo.5811103). Unfortunately, all current industrial-scale growth systems are heterotrophic, which demand fixed carbon feedstocks, partially displacing the benefit of any plastic recycling. A more sustainable alternative is to use phototrophic organisms to produce these enzymes, and microalgae are an ideal candidate due to their innate photosynthetic nature and proven capacity to produce precursors that can be converted to petroleum replacements for fuel and plastic (Gupta et al., 2024; Tang et al., 2020).

Here, we used green microalgae to secrete the polyester depolymerizing enzyme PHL7, and demonstrated its capacity to degrade post-consumer polyester plastics, including both PET and polyester polyurethanes. Our system used the previously developed pJP32 vector (Molino et al., 2018) to drive efficient secretion of this plastic-degrading enzyme from algae cells, and a screening strategy based on halo formation around colonies to detect secreting colonies (Supp. Figure 1). Impranil® is a polyester polyurethane dispersion that decreases the agar plate transparency, and becomes clear when it is enzymatically degraded. While wild-type strains might secrete native enzymes capable of cleaving ester bonds for a specific substrate, no degradation of Impranil® was observed (e.g., colonies without surrounding halo in Supp. Figure 1 and 7). This is expected since *C. reinhardtii* has never been observed to utilize nutrients trapped inside polymeric structures outside the cell for growth (e.g., plastics).

To further characterize the enzyme secreted by *C. reinhardtii*, we screened candidate colonies using the FDA and the Impranil® degradation assays (Figure 2). Both methods successfully identified positive strains, though they differ significantly in sensitivity, specificity, and practicality. The FDA assay offers a faster screening option, with results obtained in approximately 40 minutes. However, this method is not specific to polymer degradation, as native enzymes can also cleave FDA (Liu et al., 2023). This demands the inclusion of proper controls to account for background enzymatic activity unrelated to plastic degradation. While the assay is cost-effective due to its cheap substrate, its utility is limited because a follow-up test is required to confirm the presence of a plastic-degrading enzyme. In contrast, the Impranil® -based assay, though taking longer to produce results, is more specific to polymer degradation. Native enzymes are incapable of cleaving this substrate (see Figure 1 and Suppl. Figure 1), making it a more reliable indicator of plastic degradation potential. Moreover, Impranil® is structurally closer to post-consumer plastics, further aligning the assay with real-world applications. Interestingly, both methods underestimated the actual number of positive strains. Subsequent blotting of strains led to more halo formations than those identified by the FDA or Impranil® assays. This suggests that both enzymatic assays may have relatively high detection limits, potentially missing weaker positive strains. However, they remain useful for comparing candidates, as they provide a more straightforward and objective measurement of enzymatic activity compared to the subjective nature of halo formation. Though more subjective, the halo formation assay appears to have a lower detection limit. This is likely due to the continuous production of the enzyme by live cells near the cell patches, allowing for signal buildup over time. This accumulation likely increases the assay’s sensitivity, enabling it to detect weaker or slower enzyme activity that the FDA and Impranil® assays may miss.

We observed the formation of two clearing zones in a zymogram containing Impranil® as a substrate. The two clearing zones indicate the presence of two active isoforms of the enzyme. Protein sequencing of both regions identified the presence of PHL7 peptides (Supp. Data 1). Analysis of the protein sequence with NetNGlyc 1.0 (Gupta & Brunak, 2002) indicates the presence of three possible glycosylation sites at the 170, 171, and 188 amino acid positions (Supp. Figure 5), which possibly explains the two bands observed, since consecutive glycosylation sites are likely not completely glycosylated (e.g. both positions glycosylated). The peptides in the putative glycosylation region were also not observed in the protein sequence, likely due to unmatched expected peptide mass due to carbohydrate addition. We designed a gene version with substitutions in all three glycosylation sites to further characterize the protein, replacing the asparagine (N) with aspartic acid (D). Aspartic acid residues do not serve as sites for N-linked glycosylation because they lack the necessary amide group that forms the glycosidic bond with the sugar moiety. With the new pJP32PHL7dg vector (Supp. Figure 7) containing the modified version of PHL7 containing aspartic acid residues (PHL7dg), we obtained only a single clearing zone in the zymogram (Figure 3), indicating the absence of glycosylation in PHL7dg, as predicted. Interestingly, the number of colonies displaying halos in our transformation with the pJP32PHL7dg vector was significantly lower, suggesting that removing the glycosylation site in PHL7dg impacted the activity or processing of the protein in algae (Xu & Kieliszewski, 2011). This aligns with findings where adding glycomodules to mVenus resulted in a 12-fold increase in secretion (Ramos-Martinez et al., 2017), and a similar effect may have occurred here. However, the primary goal of the pJP32PHL7dg experiment was to assess the presence of glycosylation in PHL7 via gel shifting, and the observed transformation result is based on a single transformation event. Further exploration is required for a more meaningful conclusion to be drawn.

We conducted an enzymatic degradation assay using PET beads to further validate the activity of PHL7 secreted by *C. reinhardtii*. In this assay, we incubated a concentrated supernatant from the PHL7 strain and a wild-type control in 500 mM phosphate buffer (pH 8) at 68°C. The results showed a rapid increase in absorbance at 240 nm for the PHL7 sample, indicating potential PET degradation. In contrast, the wild-type sample showed a slight decrease in absorbance, likely due to components either precipitating out of the solution or degrading at the elevated temperature.The PET beads used in this assay had high crystallinity (∼50%), while PET bottles have a range of 30-40%, and other PET plastic containers 6-8% (Kawai et al., 2014; Ronkvist et al., 2009). Crystallinity is a key factor in enzyme accessibility, as highly crystalline PET is usually more resistant to enzymatic breakdown (Walter et al., 2022). Despite this challenge, the PHL7 enzyme still demonstrated degradation activity, suggesting its efficacy even on more structurally robust PET substrates. Nevertheless, complete degradation was not observed, either by crystallinity resistance or enzyme losing activity after an extended time.

Interestingly, after day 2, both the wild-type and PHL7 samples displayed similar trends in the increase of absorbance (p-value: 0.5159). This suggests that another process, unrelated to the direct enzymatic activity of PHL7, may be contributing to the signal. However, given that terephthalic acid (TPA), one of the monomers released during PET degradation, was detected by GC-MS in both samples at the final time point, it is possible that some level of PET degradation occurred in both conditions. However, the PHL7 sample exhibited a stronger signal, consistent with the higher 240 nm absorbance, further supporting the role of PHL7 in PET degradation.Overall, the increase in absorbance and the detection of TPA reinforce the conclusion that PHL7 facilitates PET degradation. The additional increase in signal for the wild-type suggests that other environmental or chemical factors may influence PET breakdown, albeit to a lesser extent.

This experiment utilized a concentrate of the supernatant containing the PHL7 enzyme, a potential approach for plastic degradation. Specifically, algal biomass can be separated from the PHL7-containing supernatant and converted into alternative materials, such as animal feed, fuel, or plastic (Gupta et al., 2024). Yet, another strategy involves exposing plastic to growing cultures that secrete active enzymes. To illustrate this, we demonstrated the degradation of a sustainable polyester urethane (sPU) film (Figure 6). Since sPU can be derived from biological sources, our group has previously shown the successful conversion of algae oil and starch to form such materials (Chavarro Gomez et al., 2020; Gupta et al., 2024). After approximately ten days of exposure to the pJP32PHL7 culture, secreting the PHL7 enzyme, the culture breached the sPU film and fell into the receiving flask. The culture was maintained for an additional month, during which no breach occurred with the wild-type strain. This result only illustrates the strategy since no replicas were performed. Still, it is corroborated by the thousands of colonies observed in our Impranil® selection plates, displaying clearing zones (halos) around them after growth (Supp. Figure 1, Supp. Figure 4, Video1).

**Figure 6:**
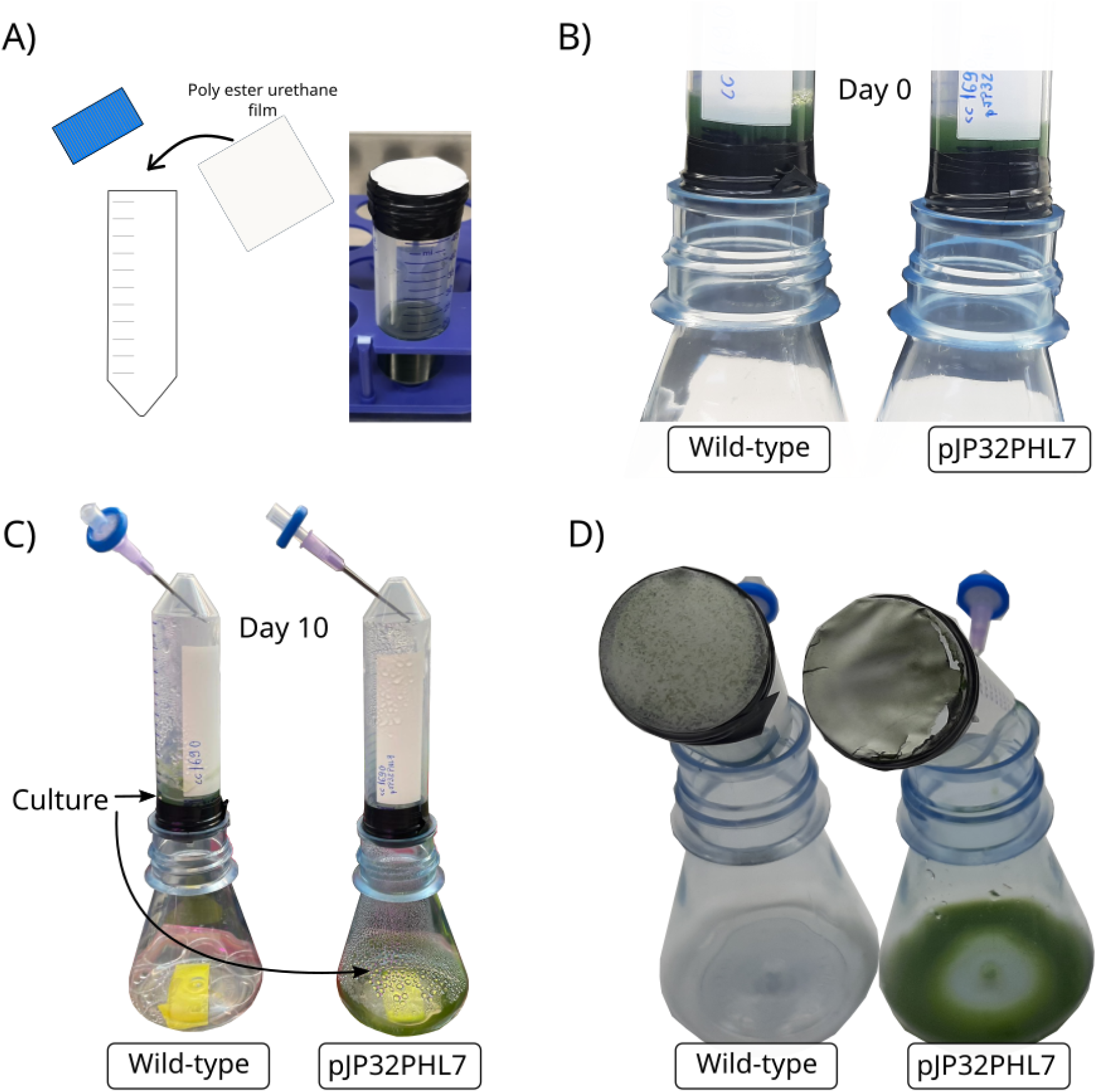
Demonstration of plastic degradation with a culture. (A) Sustainable polyester urethane (sPU) films were taped onto 50 mL centrifuge tubes containing 10 mL of wild-type cc1690 and pJP32PHL7 cell cultures. (B) Centrifuge tubes were inverted and attached to the opening of empty Erlyenmeyer flasks. (C) A syringe with an air filter was inserted into the conical part of each centrifuge tube. After approximately 10 days of growth at 25°C with constant illumination at 80 μmol photons/m²s and agitated at 150 rpm on a rotary shaker, liquid culture was observed in the pJP32PHL flask, and no culture was observed in the wild-type flask, thereby indicating enzymatic degradation of the sPU film by pJP32PHL7. (D) The sPU film for the wild-type displayed no degradation. In contrast, the sPU film for pJP32PHL7 clearly displayed a tear on the perimeter of the film, thus indicating degradation by PHL7 enzymes secreted from the recombinant strain.

Previously, *C. reinhardtii* was used to express IsPETase in the chloroplast, one of the first enzymes shown to degrade PET plastics. PET degradation was confirmed using HPLC and scanning electron microscopy with lysates from the transformed cells, demonstrating the potential of green algae to produce the enzyme (Di Rocco et al., 2023). However, accumulation of enzymes inside the cells poses challenges, as it requires cell disruption to release the enzyme, a resource and energy-intensive process. This also complicates the separation of cell biomass from the enzyme, limiting the biomass from being used to produce fuel or bioplastics. Similarly, other researchers engineered IsPETase to be secreted by the marine diatom *Phaeodactylum tricornutum*, demonstrating PET degradation. Culturing with post-consumer plastic yielded lower enzyme activity, potentially due to suboptimal enzymatic reaction conditions (i.e., 37 ℃) (Brott et al., 2022) since performed at *P. tricornutum* growth condition (i.e., 21 ℃) (Moog et al., 2019).

In the case of *C. reinhardtii* producing PHL7, the optimal culture temperature (∼25°C (Merchant et al., 2007)) differs significantly from the optimal temperature for PHL7 activity on PET (∼70°C,(Sonnendecker et al., 2021)). Therefore, a compartmentalized process may be more economically and environmentally feasible, as matching growth conditions to enzyme requirements for efficient plastic degradation is unlikely. Nonetheless, we have demonstrated that *C. reinhardtii*, while secreting PHL7, can degrade polymers such as Impranil® DLN and sPU plastic film. Further analyses, such as techno-economic assessments (TEA) and life cycle assessments (LCA), are necessary to determine which strategy—compartmentalized or integrated—would be more effective. However, these analyses are beyond the scope of this manuscript.

## Conclusion

Overall, our experiments demonstrate that green algae can efficiently secrete PHL7, an enzyme capable of degrading polyester plastics, and that the enzyme can depolymerize PET plastic and polyurethane plastics. We demonstrated a comprehensive strategy to generate and efficiently screen recombinant strains capable of secreting functional plastic degrading enzymes, employing a polyplastic dispersion (e.g. Impranil® plates). Such a strategy can be applied to further examine secretion of plastic degrading enzymes, or even to help elucidate synthetic biology strategies to increase secretion in different systems. We envisioned two possible strategies to degrade plastic with green algae biologically. A coupled system, in which cultures would not only harbor the required nutrients for cell growth, but also contain plastic material to be degraded and used as a food source for the algae. Ideally, the strain should be further engineered to assimilate the released organic molecules from the plastic in an upcycling process. Such a strategy is challenging since membrane transporters, metabolic engineering to incorporate the required pathway, and the need of matching the cell requirement to growth and the enzyme requirements to degrade plastic are not presently aligned. On the other hand, a compartmentalized strategy appears attainable in the foreseeable future, provided that each process, cell growth and enzyme reaction, can be performed independently and in its optimal setup.

## Material and Methods

### Assembly of transformation vectors

All restriction enzymes used in this study were acquired from New England Biolabs (Ipswich, MA, US). The vectors utilized are derivatives of the pJP32 vector (Molino et al., 2018), available in the Supplementary dataset. These vectors were assembled using the pBlueScript II KS+ (pBSII) backbone. To create the pJP32 PHL7 construct, the required PHL7 codon-optimized sequence was purchased from IDT (Integrated DNA Technologies, San Diego, CA, USA) and integrated into the expression vectors by NEBuilder® HiFi DNA Assembly (NEB - New England Biolabs). The backbone was prepared by PCR using the protocol described in (dx.doi.org/10.17504/protocols.io.bprimm4e), with 20 bp homology arms to the synthesized PHL7 sequence. The deglycosylated version (PHL7dg) was also added in the same fashion. All vectors contain restriction sites flanking the expression cassette for linearization, XbaI on the 5’ side and KpnI on the 3’ end. Final sequences can be found at (https://doi.org/10.5281/zenodo.13959924). All vector maps can be found at Supp. Figure 8.

### Culture conditions and *C. reinhardtii* transformation

Nuclear transformations were performed on the wildtype, cell wall-containing strain *C. reinhardtii* cc1690 (mt+) (<u>Chlamydomonas Resource Center</u> in St. Paul, MN, USA). This strain was propagated in TAP medium at 25 °C, with constant illumination at 80 μmol photons/m²s, and agitated at 150 rpm on a rotary shaker. Growth curves were established using the protocol described in protocols.io (dx.doi.org/10.17504/protocols.io.bpvbmn2n), involving the addition of 160 µL aliquots from 250 mL cultures into a 96-well plate per daily sampling. The absorbance was then measured using an Infinite® M200 PRO plate reader (Tecan, Männedorf, Switzerland), ensuring each strain was represented by three biological replicates. For transformation, *C. reinhardtii* cells were grown to the mid-log phase, achieving a density of 3–6 × 10^6^ cells/mL under the previously mentioned conditions (dx.doi.org/10.17504/protocols.io.bx5cpq2w). Cells were then harvested by centrifugation at 3000 xg for 10 min and resuspended in a MAX Efficiency™ Transformation Reagent for Algae to a 3–6 × 10^8^ cells/mL density. Following a 5–10 min incubation on ice with 500 ng of a double-digested vector plasmid, the cells were electroporated using a Gene Pulser® set to 2000 V/cm and 20 μs. Post-electroporation, the cells were allowed to recover in TAP medium, under gentle agitation in room light, for 18 hours. The recovered cells were then centrifuged, resuspended in 600 μL TAP medium, and spread onto two TAP/agar plates containing either 0.5% or 0.75% (v/v) Impranil® DLN ® with 15 μg/mL of zeocin. Incubation continued under light at 60 μmol photons/m²s and a temperature of 25 °C until colony formation was observed.

### Strain screening

Transformants were screened for enzyme activity by observing areas of Impranil® DLN clearing or “halos” around colonies on TAP agar plates containing 15 µg/mL Zeocin and Impranil® DLN at 0.5% and 0.75% (v/v) (DOI: dx.doi.org/10.17504/protocols.io.rm7vzb695vx1/v1). The total number of colonies was determined using OpenCFU (Geissmann, 2013). These halos indicated the degradation of the Impranil® DLN polymer in the plates. Cultured in 160µL of TAP medium for five days in 96-well plates (Nunc™ Edge™ 96-Well, Nunclon Delta-Treated, Flat-Bottom Microplate, Thermo Scientific™), we selected 84 colonies from these plates following the protocol detailed in dx.doi.org/10.17504/protocols.io.big9kbz6, alongside six wild type colonies and six wells with media as blanks. We used a Thermo plate shaker, Model #4625 (Thermo Scientific, 2555 Kerper Boulevard, Iowa, USA, Thermo Labline 4625 Titer shaker) set to 800 rpm under constant illumination (60 µmol photons/m^2^s) for cultivation. Absorbance and fluorescence measurements were taken using an Infinite® M200 PRO plate reader (Tecan, Männedorf, Switzerland) with complete settings described in the Data Setting file. To establish a baseline for our experiments, the six independent replicates of the parental wild-type strain cc1690 were used as a negative control. Following cultivation, we centrifuged the plates at 3000 xg for 5 minutes to collect the supernatant, which was then used in the enzymatic assay. The remaining cultures were transferred using a microplate replicator onto a rectangular agar plate containing TAP and Impranil® DLN dispersion to confirm the selected colonies’ ability to generate halos.

### Plate reader settings

The Infinite® M200 PRO plate reader (Tecan, Männedorf, Switzerland) plate reader was set to measure cell density and PHL7 activity. Cell density could be followed with chlorophyll fluorescence at Ex. 440 nm Em. 680 nm, and absorbance at 750 nm. A set of protocols followed enzyme activity. Using Fluorescein Diacetate (FDA), a fluorophore that fluoresces at Ex. 490 nm Em. and 520 nm when the ester bonds are cleaved, the detailed protocol was added to protocol.io (dx.doi.org/10.17504/protocols.io.n2bvj3j9blk5/v1). The activity was also followed using a plastic dispersion protocol with Impranil® DLN (Bayer Corporation, Germany), using a gel containing 0.2% (m/v) agarose and 0.25% Impranil® DLN DLN (v/v) to keep Impranil® DLN in suspension and absorbances readings were made in 5 min intervals as detailed described in (dx.doi.org/10.17504/protocols.io.14egn9bxml5d/v1). The enzyme’s ability to degrade post-consumer plastic was followed by a spectroscopy method using UV-transparent 96 well plates and readings at 240 nm. The protocol is fully described in (dx.doi.org/10.17504/protocols.io.bp2l6xp8klqe/v1). All settings are described in Data settings file.

### Zymogram

We utilized the TGX Stain-Free™ FastCast™ Acrylamide Starter Kit (Bio-Rad Laboratories, USA) to prepare upright, SDS zymogram gels. The acrylamide solution was mixed as per the manufacturer’s instructions, with the modification of adding 1% v/v Impranil® DLN to the solution to enable the detection of enzyme activity. This mixture was then poured into a casting frame and allowed to polymerize. Post-polymerization, the gel was placed in the electrophoresis apparatus and run under standard protein gel conditions (120-160V, 1-2h) following the run front with the blue dye. The samples were prepared for electrophoresis by adding 4X Laemmli buffer (#1610747, Bio-Rad Laboratories, USA). Following electrophoresis, the gel was immersed in a 100 mM Potassium Phosphate buffer solution, pH 8.0, and incubated at 37°C until transparent bands (clearing zones) appeared. This incubation step was crucial for developing clearing zones, which indicates enzymatic degradation of the Impranil® DLN within the gel matrix. Clearing zones typically emerged within a couple of days of incubation, allowing for the qualitative assessment of enzyme activity.

### Protein Sequencing

The bands identified in the zymogram of the pJP32 PHL7 supernatant sample were sequenced to confirm the presence of the PHL7 protein. Shortly, the band was cut and reduced into 1 mm cubes, followed by washing steps to remove running buffers and dyes, first with H_2_0, then a 50/50 ACN/H2O, and finally only ACN. The samples were then alkylated, digested with trypsin, and extracted in a 5% formic acid solution for mass spectrometry. The mass spectrometry was performed at the Biomolecular and Proteomics Mass Spectrometry Facility at UC San Diego using a LUMOS Orbi-Trap, and their full protocol can be found under “https://bpmsf.ucsd.edu/training-protocols/protocols.html”.

### PET degradation assay

Polyethylene terephthalate (PET) beads (Goodfellow Cambridge Limited, Huntingdon, UK; Product code ES306000/1) with a maximum particle size of 300 µm and crystalinity of >50% were used for enzymatic degradation studies. The degradation of PET was assessed by quantifying the release of terephthalic acid (TPA) via absorbance at 240 nm using a UV-transparent microplate (UV-Star™ 96-well microplates) (REF). PET plastic beads were washed in 1M potassium phosphate buffer (pH 8.0) and prepared as a slurry with 20-30% solids. In individual PCR tubes, 50 µL of PET slurry, 50 µL of 100 mM potassium phosphate buffer, and 100 µL of plastic-degrading enzyme solution were combined. The reaction was initiated by mixing, followed by centrifugation, and the absorbance at 240 nm was measured to establish a baseline (T0). Tubes were incubated at 68°C for seven days in a thermocycler with a heated lid (105°C) to prevent condensation. After incubation, the tubes were cooled to room temperature, and 100 µL of the supernatant was transferred to UV-Star™ 96-well microplates for absorbance measurement at 240 nm using a TECAN plate reader. A control with the supernatant of a wild-type strain was included to account for non-enzymatic degradation. The extent of PET degradation was calculated by comparing absorbance values from the test samples to those of the control. The absorbance values were converted to milligram equivalents of terephthalic acid (TPA) using a standard curve generated with TPA dissolved in buffer and measured under the same conditions as the samples.

### Monomer Detection - Mass Spectrometry

Enzyme samples were incubated with PET beads plastic in 0.5M potassium phosphate buffer, pH8, at 68°C for seven days. The supernatant was recovered and submitted to a liquid-liquid extraction protocol.

### Liquid-liquid extraction of TPA and LC-MS

The monomers from PET were extracted from a 100 µL enzymatic reaction mixture using a standardized solvent extraction method. An equal volume of ethyl acetate (100 µL) was initially added to the enzymatic reaction mixture in a centrifuge tube, facilitating the monomer’s extraction. 50 uL of HCl ∼10M was added and the mixture was then vortexed vigorously for complete mixing and subsequently centrifuged at 10,000 x g for 5 minutes to enable phase separation. The upper organic layer containing the extracted TPA was carefully transferred to a new tube. To dry ethyl acetate extract was then subjected to solvent evaporation under normal pressure and room temperature overnight to avoid thermal degradation of TPA. The resultant dry TPA extract was resuspended in methanol for mass spectrometry. Monomer detection was performed via direct injection on a HESI-Orbitrap in negative mode. The source temperature was 100C, sheath gas flow was 10, capillary temperature was 350 C, and the spray voltage was 3.5 kV. FTMS scans were taken from 90-500 m/z at 240,000 resolution. The injection flow rate was set at 10uL/min. The EIC at 165.02 m/z, identified as the deprotonated TPA [M-H], was averaged over 30 seconds.

### Plastic Film Degradation

A sustainable polyester urethane (sPU) film derived from algae oil was obtained from Algenesis Materials (PC2). 10 mL of *C. reinhardtii* cc1690 cell cultures and pJP32PHL7 at density 3–6 × 10^7^ cells/mL in TAP medium were added to 50 mL centrifuge tubes (Genesee Scientific 28-108). The centrifuge tubes were sealed with the sPU film, locked into place with black electrical tape, and inverted so that the culture was in direct contact with the sPU film. The inverted tube containing cell culture was fitted tightly into the opening of an Erlenmyer flask. A syringe equipped with a syringe filter (Whatman Uniflo 9916-1302) was inserted into the conical part of the inverted centrifuge tube, establishing an open system that ensures sterility. This configuration allowed for aeration within the system while safeguarding against culture contamination. The cells were cultured in this configuration at a stable temperature of 25°C with constant illumination at 80 μmol photons/m²s and agitated at 150 rpm on a rotary shaker for ten days or until the cell cultures had degraded and penetrated the sPU film.

### Data Analysis

R Statistic version 4.3.2 running in the RStudio 2023.09.1+494 “Desert Sunflower” was used to import and process data, generate the statistical summary, and generate the plots. The codes used are deposited at Zenodo (https://doi.org/10.5281/zenodo.13959987). The data herein was collected from experiments in which, pJP32PHL7 was used to transform the CC1690 strain, and 84 colonies were picked for screening. These colonies were individually assessed through absorbance and fluorescence measurements, providing 84 independent data points per condition in the initial screening phase. For FDA analysis, the presence of cells after centrifugation interferes with the activity measurement due to enzymes inside cells (Chen et al., 2016), and wells with a chlorophyll signal higher than 100 RFU were excluded from the analysis. In flask culture analyses, standard deviation bars represent the variation across three biological replicates of each strain.

## Data Availability

The datasets generated during and/or analysed during the current study are available in the ZENODO repository, https://doi.org/10.5281/zenodo.13981200.

## Author contributions

**JVDM**: Conceptualization, Data curation, Formal analysis, Investigation, Methodology, Visualization, Writing – original draft, Writing – review & editing

**BS:** Investigation, Methodology, Writing – review & editing

**KK:** Investigation, Methodology, Visualization, Writing – review & editing

**CW:** Investigation, Writing – review & editing

**CJD:** Investigation, Writing – review & editing

**MT:** Investigation, Methodology, Writing – review & editing

**SM:** contributed to drafting and revising the original manuscript and secured funding for the research.

## Supporting information

Dataset - https://doi.org/10.5281/zenodo.13981200

## Video

**Video 1**: https://www.youtube.com/watch?v=a2LE5zZe9oo

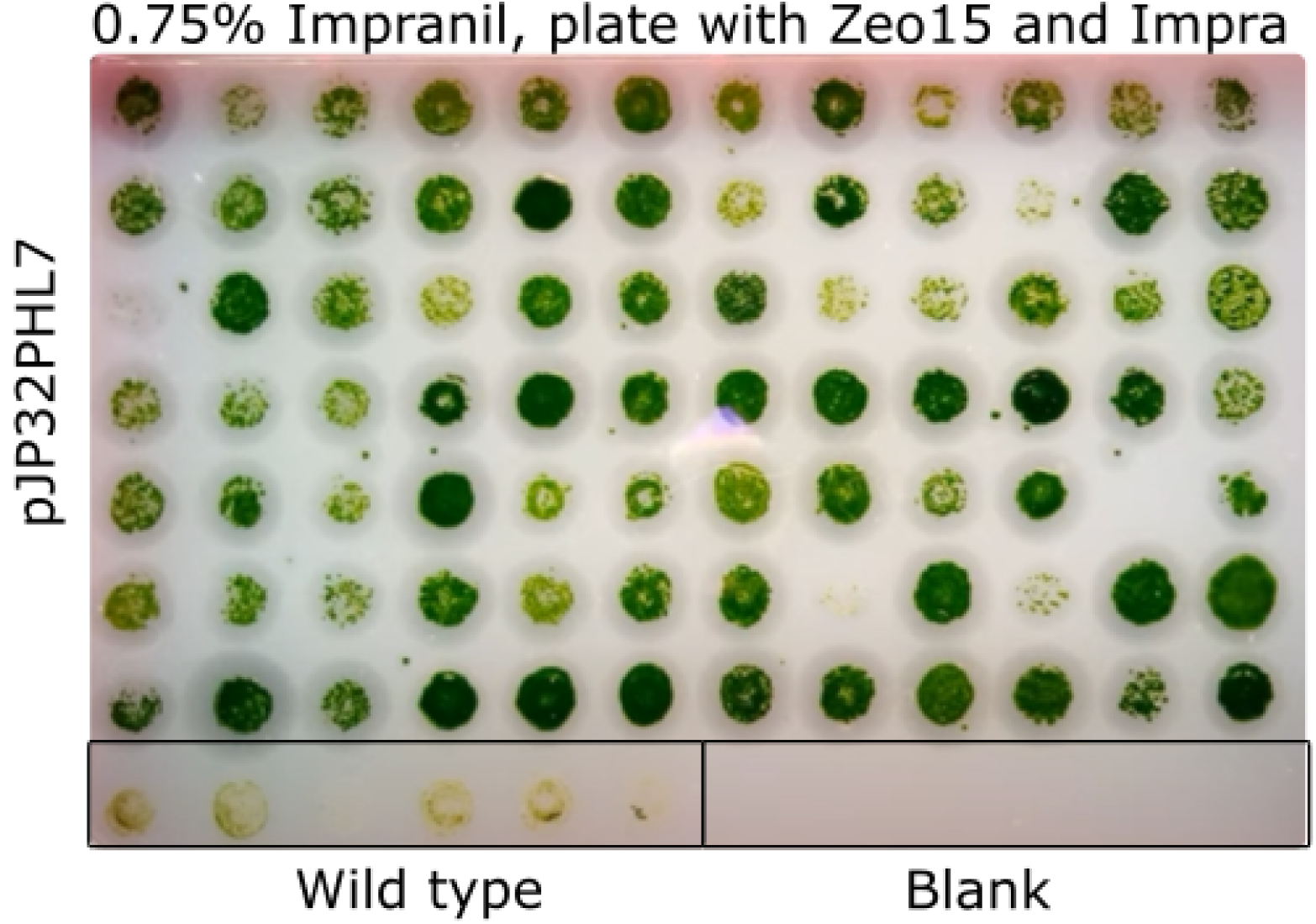

