## Supplementary material for "Efficient secretion of a plastic degrading enzyme from the green algae *Chlamydomonas reinhardtii*": Dataset - https://doi.org/10.5281/zenodo.13981200: peptides.html

peptide list


Peptide Sequence Contains:

Peptide Sample Area >=

Peptide PTM Contains:

Show Peptide From:

With Protein Accession: False

Peptide List

  

| Peptide | -10lgP | Mass | Length | ppm | m/z | RT | Fraction | Scan | Source File | Area Sample 1 | #Feature | #Feature Sample 1 | Accession | PTM | AScore | Found By |
| --- | --- | --- | --- | --- | --- | --- | --- | --- | --- | --- | --- | --- | --- | --- | --- | --- |
| SVGAAGGQEDIVNLVPTAAAATSTGK | 97.51 | 2384.2183 | 26 | 3.2 | 795.7492 | 30.00 | 1 | 6916 | JVDM\_BotE.raw | 4.5271E4 | 1 | 1 | tr\|A0A2K3DP12\|A0A2K3DP12\_CHLRE |  |  | PEAKS DB |
| GPDPTESSIEAVR | 97.26 | 1356.6521 | 13 | 2.4 | 679.3350 | 16.48 | 1 | 3077 | JVDM\_BotE.raw | 2.7415E6 | 1 | 1 | PHL7 |  |  | PEAKS DB |
| VAVLANEQELSVEER | 93.89 | 1684.8632 | 15 | 1.6 | 843.4402 | 23.17 | 1 | 5030 | JVDM\_BotE.raw | 6.8004E5 | 1 | 1 | tr\|Q7X7A7\|Q7X7A7\_CHLRE |  |  | PEAKS DB |
| VIDAGANALVAGSAVFK | 87.38 | 1601.8777 | 17 | 2.3 | 801.9480 | 30.60 | 1 | 7081 | JVDM\_BotE.raw | 1.1643E5 | 1 | 1 | tr\|A8IKW6\|A8IKW6\_CHLRE |  |  | PEAKS DB |
| IASQGFVVITIDTITR | 86.17 | 1732.9723 | 16 | 2.7 | 867.4958 | 38.48 | 1 | 8998 | JVDM\_BotE.raw | 1.4829E6 | 2 | 2 | PHL7 |  |  | PEAKS DB |
| TPTLVVGAQLDTIAPVSSHSEAFYNSLPSDLDK | 82.00 | 3471.7410 | 33 | 2.8 | 868.9449 | 36.25 | 1 | 8484 | JVDM\_BotE.raw | 4.5148E5 | 2 | 2 | PHL7 |  |  | PEAKS DB |
| AAAAASLGQAQDAAAQAQNTAAAAVGNAQTAAGDAAGK | 80.91 | 3323.6089 | 38 | 0.0 | 1108.8770 | 29.70 | 1 | 6835 | JVDM\_BotE.raw | 0 | 0 | 0 | tr\|A0A2K3D8I0\|A0A2K3D8I0\_CHLRE |  |  | PEAKS DB |
| DAAAAAQQQAAAAADAAK | 78.99 | 1612.7804 | 18 | 1.1 | 807.3984 | 17.40 | 1 | 3317 | JVDM\_BotE.raw | 0 | 0 | 0 | tr\|A0A2K3D8I0\|A0A2K3D8I0\_CHLRE |  |  | PEAKS DB |
| AAGDNAQAAAVPSVTEQASEFLAGSWPK | 77.91 | 2772.3354 | 28 | 1.9 | 925.1208 | 38.38 | 1 | 8982 | JVDM\_BotE.raw | 6.3426E4 | 1 | 1 | tr\|A0A2K3D6N5\|A0A2K3D6N5\_CHLRE |  |  | PEAKS DB |
| GQAEAAAAQAQNTAAAAVGNVQTAAADAAAK | 77.81 | 2781.3640 | 31 | 1.2 | 928.1298 | 29.67 | 1 | 6825 | JVDM\_BotE.raw | 1.4042E5 | 1 | 1 | tr\|A0A2K3D8I0\|A0A2K3D8I0\_CHLRE |  |  | PEAKS DB |
| GPFAVAQTTVSR | 77.63 | 1232.6514 | 12 | 2.3 | 617.3344 | 20.39 | 1 | 4192 | JVDM\_BotE.raw | 5.5629E6 | 1 | 1 | PHL7 |  |  | PEAKS DB |
| TAIAVDTILNQK | 77.28 | 1285.7241 | 12 | 2.5 | 643.8710 | 26.92 | 1 | 6082 | JVDM\_BotE.raw | 1.5927E5 | 1 | 1 | tr\|B7U1J0\|B7U1J0\_CHLRE:P26526\|ATPA\_CHLRE |  |  | PEAKS DB |
| MIGATNPLASEPGTIR | 75.42 | 1626.8400 | 16 | 3.8 | 814.4304 | 25.35 | 1 | 5689 | JVDM\_BotE.raw | 6.1475E4 | 1 | 1 | tr\|A8J9H8\|A8J9H8\_CHLRE:tr\|A0A2K3DJF1\|A0A2K3DJF1\_CHLRE |  |  | PEAKS DB |
| GAAASAVGQATAQATAAVDQAK | 75.23 | 1956.9865 | 22 | 1.5 | 653.3371 | 24.46 | 1 | 5439 | JVDM\_BotE.raw | 2.8473E5 | 1 | 1 | tr\|A0A2K3D8I0\|A0A2K3D8I0\_CHLRE |  |  | PEAKS DB |
| SPASDTYIIFGEAK | 75.16 | 1497.7351 | 14 | 1.0 | 749.8756 | 28.74 | 1 | 6556 | JVDM\_BotE.raw | 0 | 0 | 0 | tr\|A0A2K3CWF7\|A0A2K3CWF7\_CHLRE |  |  | PEAKS DB |
| AASAAAEQAQAAAVGAQQAAVEK | 74.68 | 2111.0605 | 23 | 2.3 | 704.6957 | 19.45 | 1 | 3936 | JVDM\_BotE.raw | 8.986E4 | 1 | 1 | tr\|A0A2K3D8I0\|A0A2K3D8I0\_CHLRE |  |  | PEAKS DB |
| EGTFSNLPAGTTIK | 74.37 | 1434.7355 | 14 | 1.8 | 718.3763 | 22.60 | 1 | 4828 | JVDM\_BotE.raw | 1.9599E5 | 1 | 1 | tr\|A8IT01\|A8IT01\_CHLRE |  |  | PEAKS DB |
| AQAVFDQEFQEITLSK | 73.53 | 1852.9207 | 16 | 1.6 | 927.4691 | 31.91 | 1 | 7407 | JVDM\_BotE.raw | 2.0286E5 | 1 | 1 | tr\|Q9FE86\|Q9FE86\_CHLRE |  |  | PEAKS DB |
| AAAAELTLAAYK | 70.65 | 1191.6499 | 12 | 1.9 | 596.8334 | 23.82 | 1 | 5213 | JVDM\_BotE.raw | 8.2982E5 | 1 | 1 | tr\|Q7X7A7\|Q7X7A7\_CHLRE |  |  | PEAKS DB |
| ATVAAGEEALTIR | 69.67 | 1300.6986 | 13 | 2.2 | 651.3580 | 21.62 | 1 | 4543 | JVDM\_BotE.raw | 3.192E6 | 1 | 1 | tr\|A0A2K3CQ54\|A0A2K3CQ54\_CHLRE |  |  | PEAKS DB |
| VAVIVATVTDDVR | 67.94 | 1356.7612 | 13 | 2.1 | 679.3893 | 27.28 | 1 | 6171 | JVDM\_BotE.raw | 7.4732E5 | 2 | 2 | tr\|A8IKZ2\|A8IKZ2\_CHLRE |  |  | PEAKS DB |
| VEYTELQILC(+57.02)PQTIDSVTGYPMDDPR | 67.82 | 3039.4204 | 26 | 1.1 | 1014.1486 | 37.54 | 1 | 8770 | JVDM\_BotE.raw | 1.924E6 | 1 | 1 | tr\|A0A2K3CQ54\|A0A2K3CQ54\_CHLRE | Carbamidomethylation | C10:Carbamidomethylation:1000.00 | PEAKS DB |
| ILQSIEQSEQAK | 63.08 | 1372.7197 | 12 | 2.2 | 687.3687 | 15.42 | 1 | 2819 | JVDM\_BotE.raw | 4.4047E5 | 1 | 1 | tr\|Q7X7A7\|Q7X7A7\_CHLRE |  |  | PEAKS DB |
| VGVAGAVVVLGYNWK | 61.34 | 1530.8558 | 15 | 3.5 | 766.4379 | 35.01 | 1 | 8195 | JVDM\_BotE.raw | 4.7808E4 | 1 | 1 | tr\|A0A2K3D6N5\|A0A2K3D6N5\_CHLRE |  |  | PEAKS DB |
| GPDPTESSIEAVRGPFAVAQTTVSR | 60.11 | 2571.2927 | 25 | 2.4 | 858.1069 | 28.88 | 1 | 6627 | JVDM\_BotE.raw | 3.7583E6 | 1 | 1 | PHL7 |  |  | PEAKS DB |
| AAAAQATAAAAGAQEAASAAAGQATATATDAMTK | 60.03 | 2961.4097 | 34 | 2.8 | 988.1466 | 30.04 | 1 | 6932 | JVDM\_BotE.raw | 7.1196E4 | 1 | 1 | tr\|A0A2K3D8I0\|A0A2K3D8I0\_CHLRE |  |  | PEAKS DB |
| SPNAVNPVVEGK | 59.10 | 1209.6353 | 12 | 1.7 | 605.8259 | 13.31 | 1 | 2185 | JVDM\_BotE.raw | 2.0176E5 | 1 | 1 | tr\|A8IGH1\|A8IGH1\_CHLRE |  |  | PEAKS DB |
| SLQEALASELAAR | 53.97 | 1357.7201 | 13 | 1.7 | 679.8685 | 29.78 | 1 | 6851 | JVDM\_BotE.raw | 8.1197E4 | 1 | 1 | tr\|A8HXL8\|A8HXL8\_CHLRE |  |  | PEAKS DB |
| VAQEKADELPTTNPIR | 53.58 | 1780.9319 | 16 | 2.2 | 594.6525 | 17.75 | 1 | 3412 | JVDM\_BotE.raw | 5.1889E5 | 2 | 2 | tr\|Q7X7A7\|Q7X7A7\_CHLRE |  |  | PEAKS DB |
| EGGLGDLAYPLVADLKK | 52.12 | 1757.9563 | 17 | 1.8 | 586.9938 | 34.62 | 1 | 8124 | JVDM\_BotE.raw | 1.6547E5 | 1 | 1 | tr\|Q9FE86\|Q9FE86\_CHLRE |  |  | PEAKS DB |
| SDIIVSPSILSADFSR | 50.98 | 1705.8887 | 16 | 2.6 | 853.9538 | 35.55 | 1 | 8328 | JVDM\_BotE.raw | 2.2334E4 | 1 | 1 | tr\|A8IKW6\|A8IKW6\_CHLRE |  |  | PEAKS DB |
| IDADLDVFIQK | 50.94 | 1275.6710 | 11 | 2.9 | 638.8446 | 30.91 | 1 | 7190 | JVDM\_BotE.raw | 2.7666E5 | 1 | 1 | tr\|A8JEV1\|A8JEV1\_CHLRE |  |  | PEAKS DB |
| FVDDDLRYEQFLC(+57.02)PAPDDFAISEYR | 50.54 | 3080.3860 | 25 | 4.3 | 1027.8070 | 36.55 | 1 | 8557 | JVDM\_BotE.raw | 1.1317E5 | 1 | 1 | PHL7 | Carbamidomethylation | C13:Carbamidomethylation:1000.00 | PEAKS DB |
| SSQGVPLLAEQASSFFSSK | 50.34 | 1968.9792 | 19 | 2.2 | 985.4991 | 37.02 | 1 | 8653 | JVDM\_BotE.raw | 1.1276E5 | 1 | 1 | tr\|A0A2K3E0C1\|A0A2K3E0C1\_CHLRE:tr\|A0A2K3E094\|A0A2K3E094\_CHLRE:tr\|A0A2K3E0A7\|A0A2K3E0A7\_CHLRE |  |  | PEAKS DB |
| ISGLIYEETR | 49.18 | 1179.6135 | 10 | 2.4 | 590.8154 | 21.89 | 1 | 4629 | JVDM\_BotE.raw | 1.685E5 | 1 | 1 | tr\|A8HVA3\|A8HVA3\_CHLRE:tr\|A8HSB0\|A8HSB0\_CHLRE |  |  | PEAKS DB |
| GLGPGGVELNK | 49.10 | 1039.5662 | 11 | 2.0 | 520.7914 | 18.17 | 1 | 3526 | JVDM\_BotE.raw | 5.6894E5 | 1 | 1 | tr\|A0A2K3CRJ3\|A0A2K3CRJ3\_CHLRE:tr\|Q1ALD7\|Q1ALD7\_CHLRE:tr\|A0A2K3CRI8\|A0A2K3CRI8\_CHLRE |  |  | PEAKS DB |
| TPTLVVGAQLDTIAPVSSHSEAFYN(+.98)SLPSDLDK | 48.69 | 3472.7249 | 33 | 2.4 | 1158.5850 | 35.50 | 1 | 8314 | JVDM\_BotE.raw | 0 | 0 | 0 | PHL7 | Deamidation (NQ) | N25:Deamidation (NQ):0.00 | PEAKS PTM |
| VVIALDSSSHSGADFAAK | 48.60 | 1773.8896 | 18 | 0.0 | 592.3038 | 22.39 | 1 | 4780 | JVDM\_BotE.raw | 1.8993E5 | 1 | 1 | tr\|A0A2K3D6N5\|A0A2K3D6N5\_CHLRE |  |  | PEAKS DB |
| QLVQAGFDLK | 48.43 | 1117.6132 | 10 | 1.9 | 559.8149 | 25.03 | 1 | 5612 | JVDM\_BotE.raw | 1.0514E5 | 1 | 1 | tr\|A0A2K3E0C1\|A0A2K3E0C1\_CHLRE:tr\|A0A2K3E094\|A0A2K3E094\_CHLRE |  |  | PEAKS DB |
| ASSVAQVLNTLK | 48.38 | 1229.6979 | 12 | 1.8 | 615.8573 | 29.25 | 1 | 6724 | JVDM\_BotE.raw | 1.535E5 | 1 | 1 | tr\|B7U1J0\|B7U1J0\_CHLRE:P26526\|ATPA\_CHLRE |  |  | PEAKS DB |
| YEDNFDAVNNLVVIAQDTDKK | 48.13 | 2410.1653 | 21 | -0.7 | 804.3951 | 32.87 | 1 | 7638 | JVDM\_BotE.raw | 1.6315E5 | 1 | 1 | tr\|A0A2K3D661\|A0A2K3D661\_CHLRE |  |  | PEAKS DB |
| AAAAASLGQAQDAAAQ(+.98)AQNTAAAAVGNAQTAAGDAAGK | 47.67 | 3324.5930 | 38 | 6.3 | 1109.2119 | 29.81 | 1 | 6867 | JVDM\_BotE.raw | 0 | 0 | 0 | tr\|A0A2K3D8I0\|A0A2K3D8I0\_CHLRE | Deamidation (NQ) | Q16:Deamidation (NQ):4.64 | PEAKS PTM |
| GDFAIEVGR | 47.52 | 962.4821 | 9 | 0.8 | 482.2487 | 22.17 | 1 | 4705 | JVDM\_BotE.raw | 4.0966E5 | 1 | 1 | tr\|A8J9H8\|A8J9H8\_CHLRE:tr\|A0A2K3DJF1\|A0A2K3DJF1\_CHLRE |  |  | PEAKS DB |
| ADELPTTNPIR | 47.10 | 1225.6302 | 11 | 1.2 | 613.8231 | 19.20 | 1 | 3830 | JVDM\_BotE.raw | 3.0501E5 | 1 | 1 | tr\|Q7X7A7\|Q7X7A7\_CHLRE |  |  | PEAKS DB |
| AAADGAVANAQGAVTSFATK | 46.13 | 1819.9064 | 20 | -1.5 | 910.9591 | 24.16 | 1 | 5324 | JVDM\_BotE.raw | 6.2277E4 | 1 | 1 | tr\|A0A2K3D8I0\|A0A2K3D8I0\_CHLRE |  |  | PEAKS DB |
| QLQAALDHLR | 45.78 | 1163.6411 | 10 | 2.8 | 582.8295 | 19.39 | 1 | 3892 | JVDM\_BotE.raw | 2.7016E6 | 2 | 2 | PHL7 |  |  | PEAKS DB |
| FDLNTLASTK | 45.74 | 1108.5764 | 10 | 2.5 | 555.2969 | 25.51 | 1 | 5738 | JVDM\_BotE.raw | 2.653E5 | 1 | 1 | tr\|A8JEV1\|A8JEV1\_CHLRE |  |  | PEAKS DB |
| TLDAALLPALSADLK | 44.81 | 1510.8606 | 15 | 0.5 | 756.4380 | 37.27 | 1 | 8757 | JVDM\_BotE.raw | 9.8812E3 | 1 | 1 | tr\|A8HP90\|A8HP90\_CHLRE |  |  | PEAKS DB |
| TASTASLQLC(+57.02)R | 44.71 | 1206.6027 | 11 | 1.3 | 604.3094 | 16.77 | 1 | 3136 | JVDM\_BotE.raw | 5.0051E4 | 1 | 1 | tr\|A0A2K3DP12\|A0A2K3DP12\_CHLRE | Carbamidomethylation | C10:Carbamidomethylation:1000.00 | PEAKS DB |
| ELTEADGFVFGFPTR | 44.59 | 1684.8096 | 15 | 3.7 | 843.4152 | 36.17 | 1 | 8463 | JVDM\_BotE.raw | 0 | 0 | 0 | tr\|Q19VH5\|Q19VH5\_CHLRE:tr\|Q19VH4\|Q19VH4\_CHLRE |  |  | PEAKS DB |
| AEDTAAAAQAQAAAAAESAR | 44.31 | 1843.8660 | 20 | -0.1 | 615.6292 | 18.71 | 1 | 3725 | JVDM\_BotE.raw | 1.0125E5 | 1 | 1 | tr\|A0A2K3D8I0\|A0A2K3D8I0\_CHLRE |  |  | PEAKS DB |
| NPLVNNDFENAEGLTK | 44.00 | 1773.8533 | 16 | 2.3 | 887.9359 | 25.16 | 1 | 5663 | JVDM\_BotE.raw | 4.1153E4 | 1 | 1 | Q08365\|RR3\_CHLRE |  |  | PEAKS DB |
| DDYLNAPGETYSVK | 42.36 | 1570.7151 | 14 | 1.7 | 786.3661 | 21.37 | 1 | 4473 | JVDM\_BotE.raw | 5.041E4 | 1 | 1 | tr\|A8JH68\|A8JH68\_CHLRE |  |  | PEAKS DB |
| FVEN(+.98)GGLVVLLGGK | 41.47 | 1401.7867 | 14 | 3.3 | 701.9030 | 35.60 | 1 | 8342 | JVDM\_BotE.raw | 1.2277E4 | 1 | 1 | tr\|A0A2K3E0A7\|A0A2K3E0A7\_CHLRE | Deamidation (NQ) | N4:Deamidation (NQ):1000.00 | PEAKS PTM |
| AAAAQATAAAAGAQ(+.98)EAASAAAGQ(+.98)ATATATDAMTK(+14.02) | 39.38 | 2977.3933 | 34 | 5.1 | 993.4767 | 25.89 | 1 | 5833 | JVDM\_BotE.raw | 0 | 0 | 0 | tr\|A0A2K3D8I0\|A0A2K3D8I0\_CHLRE | Deamidation (NQ); Methylation(KR) | Q14:Deamidation (NQ):50.04;Q23:Deamidation (NQ):30.07;K34:Methylation(KR):1000.00 | PEAKS PTM |
| DYGVLIEDGPDAGVTLR | 39.29 | 1788.8893 | 17 | -1.9 | 895.4503 | 30.65 | 1 | 7101 | JVDM\_BotE.raw | 0 | 0 | 0 | tr\|A8I2V3\|A8I2V3\_CHLRE |  |  | PEAKS DB |
| LAADVMDVPTLIIAR | 39.26 | 1596.8909 | 15 | 2.9 | 799.4550 | 37.79 | 1 | 8862 | JVDM\_BotE.raw | 6.2649E4 | 1 | 1 | tr\|A8J244\|A8J244\_CHLRE |  |  | PEAKS DB |
| LAAEAEAAAAAEAEAAAR | 39.09 | 1655.8114 | 18 | 2.7 | 552.9459 | 24.30 | 1 | 5353 | JVDM\_BotE.raw | 2.3634E4 | 1 | 1 | tr\|A0A2K3D9K7\|A0A2K3D9K7\_CHLRE |  |  | PEAKS DB |
| AGLQFPVGR | 38.66 | 943.5239 | 9 | 1.7 | 472.7700 | 24.35 | 1 | 5377 | JVDM\_BotE.raw | 3.2553E5 | 1 | 1 | tr\|A8HRZ9\|A8HRZ9\_CHLRE:tr\|A8IR81\|A8IR81\_CHLRE:tr\|A8HSB5\|A8HSB5\_CHLRE:tr\|Q42680\|Q42680\_CHLRE:tr\|A8HWM8\|A8HWM8\_CHLRE:tr\|A8HWF6\|A8HWF6\_CHLRE |  |  | PEAKS DB |
| VAIDQTGIATTDR | 37.18 | 1359.6993 | 13 | 2.0 | 680.8583 | 17.21 | 1 | 3262 | JVDM\_BotE.raw | 1.6009E4 | 1 | 1 | tr\|A8J363\|A8J363\_CHLRE |  |  | PEAKS DB |
| WSYPAAGVVSDATLGR(+14.02) | 37.18 | 1662.8365 | 16 | 6.4 | 832.4309 | 27.68 | 1 | 6299 | JVDM\_BotE.raw | 5.8157E4 | 1 | 1 | tr\|A0A2K3D6N5\|A0A2K3D6N5\_CHLRE | Methylation(KR) | R16:Methylation(KR):1000.00 | PEAKS PTM |
| GGAGVGGGGAVGSGTVGR | 36.53 | 1371.6854 | 18 | -5.4 | 686.8463 | 16.75 | 1 | 3129 | JVDM\_BotE.raw | 0 | 0 | 0 |  |  |  | PEAKS DB |
| DLAAASGRGGGGSGSAFKPGK | 36.40 | 1846.9285 | 21 | -1.0 | 616.6495 | 20.63 | 1 | 4281 | JVDM\_BotE.raw | 3.0625E5 | 1 | 1 |  |  |  | PEAKS DB |
| ITSASTVC(+57.02)SHEDSSAVVIPLYTYEDGEDVR | 35.52 | 3299.5139 | 30 | 5.1 | 1100.8508 | 30.14 | 1 | 6947 | JVDM\_BotE.raw | 0 | 0 | 0 | tr\|A0A2K3D6N5\|A0A2K3D6N5\_CHLRE | Carbamidomethylation | C8:Carbamidomethylation:1000.00 | PEAKS DB |
| AAEAQ(+.98)KAASAAAEQAQAAAVGAQQAAVEK(+42.01)AAEAQ(+.98)K | 34.70 | 3351.6541 | 35 | -3.3 | 838.9180 | 24.05 | 1 | 5366 | JVDM\_BotE.raw | 6.1886E6 | 1 | 1 | tr\|A0A2K3D8I0\|A0A2K3D8I0\_CHLRE | Deamidation (NQ); Acetylation (K) | Q5:Deamidation (NQ):19.64;K29:Acetylation (K):13.13;Q34:Deamidation (NQ):3.41 | PEAKS PTM |
| QSPINVPQYQVLDGK | 34.66 | 1684.8784 | 15 | 1.8 | 843.4480 | 26.20 | 1 | 5915 | JVDM\_BotE.raw | 0 | 0 | 0 | tr\|A8IT01\|A8IT01\_CHLRE |  |  | PEAKS DB |
| SGAAAAAGPSAAAIATGEAKGNK | 33.48 | 1940.9915 | 23 | -4.9 | 971.4982 | 34.09 | 1 | 7976 | JVDM\_BotE.raw | 3.2097E4 | 1 | 1 |  |  |  | PEAKS DB |
| SHAEKPLVGSVAPDFK | 33.48 | 1680.8834 | 16 | 1.5 | 561.3026 | 20.12 | 1 | 4130 | JVDM\_BotE.raw | 4.6718E4 | 1 | 1 | tr\|Q9FE86\|Q9FE86\_CHLRE |  |  | PEAKS DB |
| SVGAAGGQEDIVNLVPTAAAATSTGK(+14.02)SNSAK(+28.03) | 33.17 | 2913.5044 | 31 | 5.0 | 972.1802 | 34.56 | 1 | 8101 | JVDM\_BotE.raw | 0 | 0 | 0 | tr\|A0A2K3DP12\|A0A2K3DP12\_CHLRE | Methylation(KR); Dimethylation(KR) | K26:Methylation(KR):0.00;K31:Dimethylation(KR):0.00 | PEAKS PTM |
| AQETAAAAAGQAQAAAADAAAK | 32.93 | 1926.9395 | 22 | 1.7 | 643.3215 | 19.41 | 1 | 3905 | JVDM\_BotE.raw | 1.5324E5 | 1 | 1 | tr\|A0A2K3D8I0\|A0A2K3D8I0\_CHLRE |  |  | PEAKS DB |
| GGGGAGAGGGGGGVLSTR | 32.54 | 1343.6542 | 18 | -6.4 | 672.8301 | 13.30 | 1 | 2186 | JVDM\_BotE.raw | 2.7557E4 | 1 | 1 |  |  |  | PEAKS DB |
| GSGIANTC(+57.02)PVLESGTTNLK | 32.45 | 1917.9465 | 19 | 1.8 | 959.9822 | 23.41 | 1 | 5104 | JVDM\_BotE.raw | 0 | 0 | 0 |  | Carbamidomethylation | C8:Carbamidomethylation:1000.00 | PEAKS DB |
| TIDSVTGYPMDDPR | 31.92 | 1565.7031 | 14 | 1.8 | 783.8602 | 21.41 | 1 | 4497 | JVDM\_BotE.raw | 6.6961E4 | 1 | 1 | tr\|A0A2K3CQ54\|A0A2K3CQ54\_CHLRE |  |  | PEAKS DB |
| KAGASDGNSGAKPVAAAK | 31.48 | 1598.8375 | 18 | -4.6 | 533.9507 | 25.95 | 1 | 5799 | JVDM\_BotE.raw | 7.7739E6 | 2 | 2 |  |  |  | PEAKS DB |
| EGGGEVGGAAGPGPGPALR | 31.23 | 1604.7906 | 19 | 9.9 | 803.4105 | 21.57 | 1 | 4535 | JVDM\_BotE.raw | 9.7735E4 | 1 | 1 |  |  |  | PEAKS DB |
| LINWDAVAQR | 30.75 | 1184.6301 | 10 | 1.2 | 593.3231 | 26.25 | 1 | 5926 | JVDM\_BotE.raw | 0 | 0 | 0 | tr\|A8IGH1\|A8IGH1\_CHLRE |  |  | PEAKS DB |
| AAT(+79.97)VAADAQ(+.98)AAAAKAQDAAAGAAAAAT(+79.97)EKAAEAQK | 30.62 | 3285.4915 | 35 | 0.3 | 1096.1714 | 33.47 | 1 | 7815 | JVDM\_BotE.raw | 5.1671E5 | 1 | 1 | tr\|A0A2K3D8I0\|A0A2K3D8I0\_CHLRE | Phosphorylation (STY); Deamidation (NQ) | T3:Phosphorylation (STY):1000.00;Q9:Deamidation (NQ):0.00;T27:Phosphorylation (STY):1000.00 | PEAKS PTM |
| VALTALTMAEYFR | 30.47 | 1484.7697 | 13 | -1.0 | 743.3914 | 37.91 | 1 | 8879 | JVDM\_BotE.raw | 0 | 0 | 0 |  |  |  | PEAKS DB |
| LLGAGPAGAASGAAGGSLR | 30.32 | 1552.8320 | 19 | -7.8 | 777.4172 | 23.46 | 1 | 5116 | JVDM\_BotE.raw | 0 | 0 | 0 |  |  |  | PEAKS DB |
| EAAAAAAAAGGAVALKSATAGEAGK | 30.24 | 2084.0862 | 25 | -0.3 | 695.7025 | 12.56 | 1 | 1633 | JVDM\_BotE.raw | 0 | 0 | 0 |  |  |  | PEAKS DB |
| STLTFTPEAEGLVK | 29.89 | 1491.7820 | 14 | 2.7 | 746.9003 | 28.40 | 1 | 6455 | JVDM\_BotE.raw | 0 | 0 | 0 | tr\|B7U1J0\|B7U1J0\_CHLRE:P26526\|ATPA\_CHLRE |  |  | PEAKS DB |
| GAAGLGQASPFAAGAPAPDAGAGAGEFGVVAAAGR | 29.47 | 2966.4634 | 35 | 0.7 | 989.8291 | 37.49 | 1 | 8780 | JVDM\_BotE.raw | 1.3084E5 | 1 | 1 |  |  |  | PEAKS DB |
| TAPAFVDLDTR | 29.32 | 1204.6088 | 11 | 3.7 | 603.3139 | 24.84 | 1 | 5561 | JVDM\_BotE.raw | 4.1756E4 | 1 | 1 |  |  |  | PEAKS DB |
| AGAAQAAATAAASQ(+.98)VAEAATGAAASAQSA | 29.21 | 2416.1465 | 29 | -1.0 | 605.0433 | 27.55 | 1 | 6254 | JVDM\_BotE.raw | 0 | 0 | 0 | tr\|A0A2K3D8I0\|A0A2K3D8I0\_CHLRE | Deamidation (NQ) | Q14:Deamidation (NQ):3.58 | PEAKS PTM |
| TGK(+14.02)LSISK(+14.02)SPNAVN(+.98)PVVEGK | 29.17 | 2053.1418 | 20 | -6.5 | 1027.5715 | 34.19 | 1 | 8014 | JVDM\_BotE.raw | 0 | 0 | 0 | tr\|A8IGH1\|A8IGH1\_CHLRE | Methylation(KR); Deamidation (NQ) | K3:Methylation(KR):15.07;K8:Methylation(KR):0.00;N14:Deamidation (NQ):9.42 | PEAKS PTM |
| YGDDFDGQTASLC(+57.02)VR | 29.02 | 1702.7257 | 15 | -0.2 | 852.3700 | 22.60 | 1 | 4837 | JVDM\_BotE.raw | 3.3339E4 | 1 | 1 |  | Carbamidomethylation | C13:Carbamidomethylation:1000.00 | PEAKS DB |
| GGSGGGRKSGAADR | 28.79 | 1231.6017 | 14 | -6.5 | 616.8041 | 13.98 | 1 | 2337 | JVDM\_BotE.raw | 6.5328E5 | 1 | 1 |  |  |  | PEAKS DB |
| RGGGGGGAASSAGGAGAADGEAAAAGGGVGGGGGR | 28.72 | 2584.1721 | 35 | 7.8 | 862.4047 | 21.42 | 1 | 4496 | JVDM\_BotE.raw | 0 | 0 | 0 |  |  |  | PEAKS DB |
| AQDAWPASLEDAK | 28.56 | 1400.6571 | 13 | 1.4 | 701.3368 | 23.56 | 1 | 5155 | JVDM\_BotE.raw | 9.8139E4 | 1 | 1 | tr\|A0A2K3E0C1\|A0A2K3E0C1\_CHLRE:tr\|A0A2K3E094\|A0A2K3E094\_CHLRE:tr\|A0A2K3E0A7\|A0A2K3E0A7\_CHLRE |  |  | PEAKS DB |
| LVAFDNQK | 28.45 | 933.4919 | 8 | 1.7 | 467.7540 | 16.11 | 1 | 2939 | JVDM\_BotE.raw | 3.7404E5 | 1 | 1 |  |  |  | PEAKS DB |
| AGGEC(+57.02)LTFDQLALQRPTGK | 28.26 | 2061.0312 | 19 | 2.0 | 688.0190 | 27.00 | 1 | 6098 | JVDM\_BotE.raw | 1.9028E5 | 1 | 1 | tr\|A8IKZ2\|A8IKZ2\_CHLRE | Carbamidomethylation | C5:Carbamidomethylation:1000.00 | PEAKS DB |
| SVLLVVLTGDR | 28.21 | 1170.6971 | 11 | 1.9 | 586.3569 | 34.70 | 1 | 8147 | JVDM\_BotE.raw | 1.59E5 | 1 | 1 | tr\|A8HXL8\|A8HXL8\_CHLRE |  |  | PEAKS DB |
| LAVNLIPFPR | 27.83 | 1138.6863 | 10 | 2.2 | 570.3517 | 35.79 | 1 | 8388 | JVDM\_BotE.raw | 3.1989E4 | 1 | 1 |  |  |  | PEAKS DB |
| MLAGSQGPASGGVAADGVTPLK | 27.73 | 1983.0095 | 22 | 0.2 | 992.5123 | 28.69 | 1 | 6542 | JVDM\_BotE.raw | 0 | 0 | 0 |  |  |  | PEAKS DB |
| GLFIISPTGVLR | 27.67 | 1271.7601 | 12 | 0.0 | 636.8873 | 37.36 | 1 | 8769 | JVDM\_BotE.raw | 4.5525E4 | 1 | 1 | tr\|A8I2V3\|A8I2V3\_CHLRE |  |  | PEAKS DB |
| LSGWSAPATLT(+79.97)PEK | 27.66 | 1536.7225 | 14 | -7.3 | 769.3629 | 33.61 | 1 | 7885 | JVDM\_BotE.raw | 6.9909E4 | 1 | 1 | tr\|A0A2K3CQ54\|A0A2K3CQ54\_CHLRE | Phosphorylation (STY) | T11:Phosphorylation (STY):0.00 | PEAKS PTM |
| YSIAWLK | 27.33 | 879.4854 | 7 | 1.6 | 440.7507 | 29.05 | 1 | 6640 | JVDM\_BotE.raw | 8.7419E5 | 1 | 1 | PHL7 |  |  | PEAKS DB |
| IEDLSAQTQAAAAEQFK | 27.21 | 1819.8951 | 17 | 0.7 | 910.9554 | 22.93 | 1 | 4936 | JVDM\_BotE.raw | 0 | 0 | 0 | tr\|A0A2K3CWF7\|A0A2K3CWF7\_CHLRE |  |  | PEAKS DB |
| QGGAGGAGGDGSGAGGGGAGSKPR | 27.21 | 1841.8364 | 24 | -2.0 | 921.9236 | 26.49 | 1 | 5983 | JVDM\_BotE.raw | 2.3897E5 | 1 | 1 |  |  |  | PEAKS DB |
| KMIGATNPLASEPGTIR | 26.66 | 1754.9348 | 17 | 2.8 | 585.9872 | 22.49 | 1 | 4796 | JVDM\_BotE.raw | 8.461E4 | 1 | 1 | tr\|A8J9H8\|A8J9H8\_CHLRE:tr\|A0A2K3DJF1\|A0A2K3DJF1\_CHLRE |  |  | PEAKS DB |
| AATVAADAQAAAAK(+14.02)AQDAAAGAAAAATEK | 26.65 | 2540.2830 | 29 | 0.0 | 847.7682 | 26.91 | 1 | 6227 | JVDM\_BotE.raw | 6.2014E5 | 1 | 1 | tr\|A0A2K3D8I0\|A0A2K3D8I0\_CHLRE | Methylation(KR) | K14:Methylation(KR):6.28 | PEAKS PTM |
| AAGAAGGTAGGSSSAGGAK | 26.58 | 1404.6593 | 19 | 5.2 | 703.3406 | 29.49 | 1 | 6775 | JVDM\_BotE.raw | 1.6605E5 | 1 | 1 |  |  |  | PEAKS DB |
| LVLPGELAK | 26.46 | 938.5800 | 9 | 0.6 | 470.2975 | 25.48 | 1 | 5726 | JVDM\_BotE.raw | 1.7562E5 | 1 | 1 |  |  |  | PEAKS DB |
| TSPVLGGGGGGAVAGAAAAGGSGGLGSSGGPVR | 26.36 | 2565.2896 | 33 | -5.8 | 642.3259 | 24.04 | 1 | 5457 | JVDM\_BotE.raw | 1.2706E6 | 1 | 1 |  |  |  | PEAKS DB |
| AGGGGGPPAAKR | 26.34 | 994.5308 | 12 | 4.0 | 498.2747 | 17.55 | 1 | 3442 | JVDM\_BotE.raw | 1.4117E6 | 1 | 1 |  |  |  | PEAKS DB |
| GAGGSSKGAGSSKAGAGGGAPLR | 26.22 | 1856.9452 | 23 | 9.3 | 619.9948 | 16.71 | 1 | 3179 | JVDM\_BotE.raw | 2.9068E5 | 1 | 1 |  |  |  | PEAKS DB |
| AKLDSVLAAVL | 25.94 | 1098.6648 | 11 | 2.0 | 550.3408 | 37.27 | 1 | 8723 | JVDM\_BotE.raw | 2.1268E5 | 1 | 1 | tr\|A8JEV1\|A8JEV1\_CHLRE |  |  | PEAKS DB |
| GGGGGGGGVGGGGDVGAGGGGAGSR | 25.91 | 1742.7679 | 25 | 9.0 | 872.3991 | 28.45 | 1 | 6473 | JVDM\_BotE.raw | 0 | 0 | 0 |  |  |  | PEAKS DB |
| TQIC(+57.02)YGSIGEVR | 25.86 | 1381.6660 | 12 | -1.3 | 691.8394 | 21.07 | 1 | 4388 | JVDM\_BotE.raw | 0 | 0 | 0 |  | Carbamidomethylation | C4:Carbamidomethylation:1000.00 | PEAKS DB |
| QSAGSGGGVGGRGR | 25.85 | 1201.5912 | 14 | 3.3 | 601.8049 | 17.67 | 1 | 3449 | JVDM\_BotE.raw | 3.2967E5 | 1 | 1 |  |  |  | PEAKS DB |
| SPPYALDALEPHMSK | 25.83 | 1654.8025 | 15 | 1.4 | 552.6089 | 27.02 | 1 | 6120 | JVDM\_BotE.raw | 7.1327E4 | 1 | 1 | tr\|A8IGH1\|A8IGH1\_CHLRE |  |  | PEAKS DB |
| TRPYDSSESDYLIR | 25.33 | 1700.8005 | 14 | 1.0 | 567.9413 | 19.77 | 1 | 4023 | JVDM\_BotE.raw | 7.0809E4 | 1 | 1 |  |  |  | PEAKS DB |
| VAGGAAGGAGGSGAATSAAASKARDAQR | 25.16 | 2343.1638 | 28 | -1.4 | 586.7974 | 36.25 | 1 | 8594 | JVDM\_BotE.raw | 9.9506E5 | 1 | 1 |  |  |  | PEAKS DB |
| ASAGAGAGGGGAPPR | 25.09 | 1152.5635 | 15 | 4.6 | 577.2917 | 15.80 | 1 | 2872 | JVDM\_BotE.raw | 1.127E5 | 1 | 1 |  |  |  | PEAKS DB |
| RGMDGSSGSGVALLPK | 25.01 | 1530.7823 | 16 | 6.4 | 766.4034 | 20.79 | 1 | 4308 | JVDM\_BotE.raw | 6.8139E4 | 1 | 1 |  |  |  | PEAKS DB |
| LADLVVALAK | 24.91 | 1011.6328 | 10 | 2.4 | 506.8249 | 32.67 | 1 | 7610 | JVDM\_BotE.raw | 2.1945E4 | 1 | 1 |  |  |  | PEAKS DB |
| MPEDLAAAK | 24.87 | 944.4637 | 9 | -5.9 | 473.2363 | 12.95 | 1 | 2086 | JVDM\_BotE.raw | 5.7766E4 | 1 | 1 |  |  |  | PEAKS DB |
| TIVDDVTIAVEK | 24.77 | 1301.7078 | 12 | 3.2 | 651.8632 | 37.05 | 1 | 8720 | JVDM\_BotE.raw | 6.8999E7 | 2 | 2 |  |  |  | PEAKS DB |
| AAAGRPAAGGGGGGGGGSVR | 24.51 | 1538.7661 | 20 | 6.6 | 770.3954 | 17.86 | 1 | 3454 | JVDM\_BotE.raw | 4.8815E3 | 1 | 1 |  |  |  | PEAKS DB |
| HEDVEAFTIPLYTAAGDEMK | 24.40 | 2236.0356 | 20 | 0.2 | 746.3527 | 34.04 | 1 | 7975 | JVDM\_BotE.raw | 0 | 0 | 0 | tr\|A0A2K3E0C1\|A0A2K3E0C1\_CHLRE:tr\|A0A2K3E0A7\|A0A2K3E0A7\_CHLRE |  |  | PEAKS DB |
| GGGSGGAGAGAAGAGASER | 24.39 | 1416.6342 | 19 | 5.9 | 709.3286 | 14.07 | 1 | 2418 | JVDM\_BotE.raw | 0 | 0 | 0 |  |  |  | PEAKS DB |
| AGPQEQWGALLNK | 24.33 | 1410.7256 | 13 | 2.5 | 706.3718 | 27.17 | 1 | 6163 | JVDM\_BotE.raw | 1.5348E5 | 1 | 1 | tr\|A0A2K3D6N5\|A0A2K3D6N5\_CHLRE |  |  | PEAKS DB |
| MDTPLAASLPGGGGR | 24.14 | 1398.6925 | 15 | 6.2 | 700.3578 | 30.82 | 1 | 7151 | JVDM\_BotE.raw | 0 | 0 | 0 |  |  |  | PEAKS DB |
| AGPAVAAPAGAAAVAAAVAGAVAAGAGGGGDDAR | 23.99 | 2658.3472 | 34 | -9.9 | 887.1142 | 39.52 | 1 | 9235 | JVDM\_BotE.raw | 0 | 0 | 0 |  |  |  | PEAKS DB |
| APAAAGAPAAAAPSAATAAGGVGPSPSGVR | 23.38 | 2428.2458 | 30 | -9.4 | 810.4149 | 30.98 | 1 | 7217 | JVDM\_BotE.raw | 5.3194E4 | 1 | 1 |  |  |  | PEAKS DB |
| LPPSGVAADGAAAAAEAARGAAAAIAAR | 23.21 | 2417.2773 | 28 | -1.1 | 806.7655 | 34.12 | 1 | 7998 | JVDM\_BotE.raw | 0 | 0 | 0 |  |  |  | PEAKS DB |
| IAGGGGGGGGNGSGSAAVK | 23.19 | 1429.6909 | 19 | -6.0 | 715.8484 | 15.18 | 1 | 2711 | JVDM\_BotE.raw | 0 | 0 | 0 |  |  |  | PEAKS DB |
| AVAAEDVAQAAVPEASR | 23.07 | 1653.8322 | 17 | 4.6 | 827.9272 | 25.66 | 1 | 5760 | JVDM\_BotE.raw | 1.7901E5 | 1 | 1 |  |  |  | PEAKS DB |
| EAAASPVAAAVASTQPLSAAPTGILSR | 22.98 | 2506.3391 | 27 | -4.8 | 836.4496 | 35.60 | 1 | 8332 | JVDM\_BotE.raw | 0 | 0 | 0 |  |  |  | PEAKS DB |
| EAAARGALAASHAR | 22.85 | 1350.7115 | 14 | -7.4 | 676.3580 | 12.83 | 1 | 1958 | JVDM\_BotE.raw | 0 | 0 | 0 |  |  |  | PEAKS DB |
| KVPAAGS(+79.97)AANIQ(+.98)PKR | 22.81 | 1587.8134 | 15 | -5.3 | 794.9097 | 37.79 | 1 | 8852 | JVDM\_BotE.raw | 0 | 0 | 0 | tr\|A0A2K3E0C1\|A0A2K3E0C1\_CHLRE | Phosphorylation (STY); Deamidation (NQ) | S7:Phosphorylation (STY):1000.00;Q12:Deamidation (NQ):0.00 | PEAKS PTM |
| LSTDSAGGLGSAAGV | 22.80 | 1261.6150 | 15 | -6.7 | 631.8105 | 20.00 | 1 | 4086 | JVDM\_BotE.raw | 9.3636E4 | 1 | 1 |  |  |  | PEAKS DB |
| RAVGPGGGGGGGGGGLPR | 22.79 | 1434.7439 | 18 | 3.3 | 718.3816 | 24.55 | 1 | 5472 | JVDM\_BotE.raw | 2.3245E4 | 1 | 1 |  |  |  | PEAKS DB |
| GAAAAASAGAAGAGGK | 22.69 | 1157.5789 | 16 | -1.4 | 579.7959 | 18.70 | 1 | 3710 | JVDM\_BotE.raw | 1.0536E7 | 2 | 2 |  |  |  | PEAKS DB |
| QLVQ(+.98)AGFDLKVTADGQ(+.98)PR | 22.69 | 1943.9952 | 18 | 3.4 | 973.0082 | 36.18 | 1 | 8464 | JVDM\_BotE.raw | 0 | 0 | 0 | tr\|A0A2K3E0C1\|A0A2K3E0C1\_CHLRE | Deamidation (NQ) | Q4:Deamidation (NQ):0.00;Q16:Deamidation (NQ):17.01 | PEAKS PTM |
| TAAAGATGAAVAGAAGAGPAAAAAESEAK | 22.66 | 2311.1404 | 29 | 2.3 | 1156.5801 | 33.34 | 1 | 7775 | JVDM\_BotE.raw | 7.3869E4 | 1 | 1 |  |  |  | PEAKS DB |
| APGVQTPVIVR | 22.58 | 1135.6713 | 11 | 4.1 | 568.8452 | 20.88 | 1 | 4337 | JVDM\_BotE.raw | 3.9738E4 | 1 | 1 |  |  |  | PEAKS DB |
| DAGAGTGASPAAGAK | 22.33 | 1200.5735 | 15 | 2.9 | 601.2958 | 19.77 | 1 | 3994 | JVDM\_BotE.raw | 4.0326E5 | 1 | 1 |  |  |  | PEAKS DB |
| EGAPAPGAGGGGKGGGGGK | 22.22 | 1437.6959 | 19 | -0.4 | 719.8550 | 13.55 | 1 | 2405 | JVDM\_BotE.raw | 2.1829E6 | 2 | 2 |  |  |  | PEAKS DB |
| IAGGGGGGGGNGSGSAAV | 22.19 | 1301.5959 | 18 | 5.3 | 651.8087 | 17.59 | 1 | 3333 | JVDM\_BotE.raw | 2.4435E6 | 1 | 1 |  |  |  | PEAKS DB |
| TASGLLSGGAGGGGGGGAVSSPAAA | 22.06 | 1885.9130 | 25 | 0.4 | 629.6452 | 23.10 | 1 | 5093 | JVDM\_BotE.raw | 2.9279E6 | 1 | 1 |  |  |  | PEAKS DB |
| TPSGGSGSGAPGAAAK | 22.05 | 1271.6106 | 16 | 8.1 | 636.8177 | 14.80 | 1 | 2662 | JVDM\_BotE.raw | 3.6588E5 | 1 | 1 |  |  |  | PEAKS DB |
| QPAGGAGGVAAAAAAGSSASGASGGLDYT | 21.83 | 2349.0833 | 29 | -6.5 | 784.0299 | 35.02 | 1 | 8208 | JVDM\_BotE.raw | 0 | 0 | 0 |  |  |  | PEAKS DB |
| DLGPYTSAQDAR | 21.81 | 1292.5996 | 12 | -4.4 | 647.3043 | 20.87 | 1 | 4323 | JVDM\_BotE.raw | 0 | 0 | 0 |  |  |  | PEAKS DB |
| RTVSTADGGAGGTVGR | 21.81 | 1460.7332 | 16 | -7.5 | 731.3683 | 23.27 | 1 | 5045 | JVDM\_BotE.raw | 1.9932E5 | 1 | 1 |  |  |  | PEAKS DB |
| GQGGGGGSSSGAAAAGAVAGAGAGAPGY | 21.75 | 2089.9412 | 28 | -0.9 | 697.6537 | 23.24 | 1 | 5028 | JVDM\_BotE.raw | 1.2692E5 | 1 | 1 |  |  |  | PEAKS DB |
| QPAAAGAAGGGGGTAAVVR | 21.73 | 1537.7960 | 19 | 4.8 | 513.6084 | 20.42 | 1 | 4207 | JVDM\_BotE.raw | 0 | 0 | 0 |  |  |  | PEAKS DB |
| GGGAGGGGGGGGGGGAVSGLR | 21.56 | 1470.6923 | 21 | -3.8 | 736.3506 | 21.92 | 1 | 4642 | JVDM\_BotE.raw | 0 | 0 | 0 |  |  |  | PEAKS DB |
| GVGGGGGGSSVGGGVAVGVK | 21.54 | 1513.7848 | 20 | 5.3 | 757.9037 | 26.49 | 1 | 5982 | JVDM\_BotE.raw | 8.5821E4 | 1 | 1 |  |  |  | PEAKS DB |
| KPTAQAGSASSPKK | 21.52 | 1356.7361 | 14 | -6.7 | 679.3708 | 22.21 | 1 | 4792 | JVDM\_BotE.raw | 2.1171E5 | 1 | 1 |  |  |  | PEAKS DB |
| GAGGGGGGGGSIIASLR | 21.42 | 1342.6953 | 17 | -8.4 | 672.3493 | 16.65 | 1 | 3096 | JVDM\_BotE.raw | 0 | 0 | 0 |  |  |  | PEAKS DB |
| TDALGAYLLTSDADEYDKPFMTGER | 21.38 | 2778.2693 | 25 | 1.7 | 927.0986 | 34.50 | 1 | 8091 | JVDM\_BotE.raw | 0 | 0 | 0 | tr\|A8J244\|A8J244\_CHLRE |  |  | PEAKS DB |
| ELDYLDGAVSNPK | 21.36 | 1419.6881 | 13 | 3.9 | 710.8541 | 23.16 | 1 | 5026 | JVDM\_BotE.raw | 0 | 0 | 0 |  |  |  | PEAKS DB |
| AAAVGGGKAGGATAGGVTAGGATAGGK | 21.32 | 2042.0504 | 27 | 3.1 | 1022.0356 | 10.54 | 1 | 949 | JVDM\_BotE.raw | 0 | 0 | 0 |  |  |  | PEAKS DB |
| TGAAGAAAAAAGGGGGAGGTDPGVAPPL | 21.19 | 2119.0293 | 28 | 4.3 | 707.3534 | 17.79 | 1 | 3429 | JVDM\_BotE.raw | 0 | 0 | 0 |  |  |  | PEAKS DB |
| ASTVADSIVDALEK | 21.16 | 1417.7300 | 14 | 2.7 | 709.8741 | 33.30 | 1 | 7758 | JVDM\_BotE.raw | 7.3643E4 | 1 | 1 | Q08365\|RR3\_CHLRE |  |  | PEAKS DB |
| GGGGRGRGGGAGGSGGIAAAR | 20.93 | 1654.8359 | 21 | -9.7 | 828.4172 | 28.43 | 1 | 6475 | JVDM\_BotE.raw | 1.5356E4 | 1 | 1 |  |  |  | PEAKS DB |
| SLGAGVGAAGEGSGH | 20.87 | 1225.5687 | 15 | 2.0 | 613.7928 | 13.08 | 1 | 2126 | JVDM\_BotE.raw | 3.8487E4 | 1 | 1 |  |  |  | PEAKS DB |
| AAVASAVGAAAAAVALR | 20.87 | 1438.8256 | 17 | -6.3 | 720.4155 | 27.64 | 1 | 6274 | JVDM\_BotE.raw | 3.125E4 | 1 | 1 |  |  |  | PEAKS DB |
| ASC(+57.02)AGGTGEAGAGAGAGAAR | 20.78 | 1618.7117 | 20 | 8.6 | 810.3701 | 29.22 | 1 | 6712 | JVDM\_BotE.raw | 0 | 0 | 0 |  | Carbamidomethylation | C3:Carbamidomethylation:1000.00 | PEAKS DB |
| GAAAAAAVGLAGAGLAGAAAGGAGGGVGGGGGGGGGGR | 20.78 | 2662.3284 | 38 | -9.8 | 888.4414 | 23.14 | 1 | 5015 | JVDM\_BotE.raw | 0 | 0 | 0 |  |  |  | PEAKS DB |
| AGGGGGGGGGGGGGGEGAISPEAVR | 20.75 | 1896.8673 | 25 | 3.9 | 633.2988 | 22.01 | 1 | 4612 | JVDM\_BotE.raw | 5.3575E5 | 1 | 1 |  |  |  | PEAKS DB |
| MAAMLAS(+79.97)KQGAFMGR | 20.73 | 1648.7289 | 15 | -8.9 | 825.3644 | 17.81 | 1 | 3435 | JVDM\_BotE.raw | 0 | 0 | 0 | tr\|A8HXL8\|A8HXL8\_CHLRE | Phosphorylation (STY) | S7:Phosphorylation (STY):1000.00 | PEAKS PTM |
| AGGGGGGGGGAVPR | 20.69 | 1025.5002 | 14 | 1.4 | 513.7581 | 22.22 | 1 | 4727 | JVDM\_BotE.raw | 0 | 0 | 0 |  |  |  | PEAKS DB |
| GSPTASAAGGAAPYK | 20.67 | 1304.6360 | 15 | 6.5 | 653.3295 | 20.84 | 1 | 4329 | JVDM\_BotE.raw | 3.7313E4 | 1 | 1 |  |  |  | PEAKS DB |
| AGPAPGSGSGSGALVR | 20.67 | 1339.6843 | 16 | 2.6 | 670.8512 | 22.94 | 1 | 4910 | JVDM\_BotE.raw | 1.4563E6 | 1 | 1 |  |  |  | PEAKS DB |
| AGGGADGGGGLAAAAITATAR | 20.61 | 1684.8492 | 21 | -2.0 | 562.6226 | 19.49 | 1 | 3956 | JVDM\_BotE.raw | 8.4094E3 | 1 | 1 |  |  |  | PEAKS DB |
| QGRAGAGGDGTVELQAGAVVANGR | 20.54 | 2210.1152 | 24 | -9.7 | 737.7052 | 32.32 | 1 | 7511 | JVDM\_BotE.raw | 0 | 0 | 0 |  |  |  | PEAKS DB |
| TGYGALVVR | 20.43 | 934.5236 | 9 | 0.6 | 468.2693 | 22.56 | 1 | 4824 | JVDM\_BotE.raw | 3.1137E4 | 1 | 1 |  |  |  | PEAKS DB |
| VAGAAGAAAGAGAAAAPAAAS | 20.40 | 1523.7692 | 21 | 7.2 | 762.8973 | 17.12 | 1 | 3242 | JVDM\_BotE.raw | 0 | 0 | 0 |  |  |  | PEAKS DB |
| YEQFLC(+57.02)PAPDDFAISEYR | 20.38 | 2219.9834 | 18 | 3.1 | 1111.0024 | 33.56 | 1 | 7863 | JVDM\_BotE.raw | 7.955E4 | 1 | 1 | PHL7 | Carbamidomethylation | C6:Carbamidomethylation:1000.00 | PEAKS DB |
| GSEGGGGGGDGGLGNGGSR | 20.27 | 1503.6298 | 19 | 4.2 | 752.8253 | 17.64 | 1 | 3381 | JVDM\_BotE.raw | 1.9903E4 | 1 | 1 |  |  |  | PEAKS DB |
| EAAAGASTSAPATTKAMTGK | 20.27 | 1820.8938 | 20 | 4.9 | 911.4586 | 22.97 | 1 | 4950 | JVDM\_BotE.raw | 0 | 0 | 0 |  |  |  | PEAKS DB |
| GSAFTGPGSPLVGAAR | 20.22 | 1443.7469 | 16 | 9.3 | 482.2607 | 21.65 | 1 | 4651 | JVDM\_BotE.raw | 3.5595E5 | 1 | 1 |  |  |  | PEAKS DB |
| AGGGGGAAAAAGLDHGGSTAMRALINWR | 20.22 | 2565.2617 | 28 | 5.0 | 642.3259 | 24.04 | 1 | 5357 | JVDM\_BotE.raw | 1.2706E6 | 1 | 1 |  |  |  | PEAKS DB |
| EAVVAPVTSTQDEERRQSGAAASAR | 20.21 | 2585.2793 | 25 | -4.5 | 862.7631 | 32.54 | 1 | 7565 | JVDM\_BotE.raw | 0 | 0 | 0 |  |  |  | PEAKS DB |
| GALAAAAAGGDVTGGTK | 20.19 | 1386.7102 | 17 | 1.0 | 694.3631 | 22.08 | 1 | 4698 | JVDM\_BotE.raw | 7.4332E4 | 1 | 1 |  |  |  | PEAKS DB |
| AAAAGGGGGGGGSGAATLGR | 20.02 | 1471.7126 | 20 | -3.9 | 736.8607 | 21.97 | 1 | 4659 | JVDM\_BotE.raw | 0 | 0 | 0 |  |  |  | PEAKS DB |
| LTFDEIQGLTYLQVK | 19.88 | 1766.9454 | 15 | 2.2 | 884.4819 | 36.83 | 1 | 8621 | JVDM\_BotE.raw | 1.7816E4 | 1 | 1 |  |  |  | PEAKS DB |
| PGGGAGGGGGAGGGGGGGGGEGGRAVNPLELLGER | 19.87 | 2817.3501 | 35 | -1.6 | 940.1224 | 29.77 | 1 | 6855 | JVDM\_BotE.raw | 0 | 0 | 0 |  |  |  | PEAKS DB |
| GPADGGGGGGGGGKPGAGGKGAGGAAAAGVR | 19.86 | 2305.1270 | 31 | -4.4 | 769.3795 | 27.66 | 1 | 6283 | JVDM\_BotE.raw | 0 | 0 | 0 |  |  |  | PEAKS DB |
| VAGASGAGGGAVRNGGAEAASDGSASAR | 19.77 | 2330.0959 | 28 | 3.8 | 777.7089 | 36.64 | 1 | 8575 | JVDM\_BotE.raw | 0 | 0 | 0 |  |  |  | PEAKS DB |
| ATAAAAGGAGGAGGHR | 19.64 | 1251.6068 | 16 | -1.4 | 626.8098 | 16.73 | 1 | 3108 | JVDM\_BotE.raw | 9.1884E4 | 1 | 1 |  |  |  | PEAKS DB |
| STKAVGGGAGGGGGGGGGGYIGQK | 19.61 | 1905.9292 | 24 | 2.0 | 953.9738 | 28.52 | 1 | 6508 | JVDM\_BotE.raw | 1.6232E4 | 1 | 1 |  |  |  | PEAKS DB |
| QNSAGGAPVSPGAGGPSSSGGAAPAAAAGA | 19.61 | 2349.0945 | 30 | -4.0 | 784.0356 | 14.09 | 1 | 2380 | JVDM\_BotE.raw | 2.1841E5 | 1 | 1 |  |  |  | PEAKS DB |
| DGPAATPLSAAPGAAPPPAPGAGA | 19.58 | 1980.9904 | 24 | 2.3 | 661.3389 | 23.77 | 1 | 5209 | JVDM\_BotE.raw | 0 | 0 | 0 |  |  |  | PEAKS DB |
| AGGAAAGDAAAGGGAAAAAAAGTVR | 19.58 | 1882.9244 | 25 | -7.5 | 628.6440 | 23.80 | 1 | 5224 | JVDM\_BotE.raw | 1.5528E4 | 1 | 1 |  |  |  | PEAKS DB |
| QLTAPAAHAPGGGGGGGNAR | 19.54 | 1715.8451 | 20 | 2.0 | 858.9315 | 28.43 | 1 | 6471 | JVDM\_BotE.raw | 6.4011E4 | 1 | 1 |  |  |  | PEAKS DB |
| APGAGGAAGGLGAPSGL | 19.53 | 1279.6520 | 17 | -2.3 | 640.8318 | 17.20 | 1 | 3259 | JVDM\_BotE.raw | 0 | 0 | 0 |  |  |  | PEAKS DB |
| GAGGGGGPGGGGVAPPAVNR | 19.32 | 1560.7756 | 20 | 0.6 | 781.3956 | 12.74 | 1 | 1860 | JVDM\_BotE.raw | 0 | 0 | 0 |  |  |  | PEAKS DB |
| YLVPSASTTEAAVFYLK | 19.28 | 1858.9716 | 17 | 1.1 | 930.4941 | 34.72 | 1 | 8142 | JVDM\_BotE.raw | 0 | 0 | 0 | tr\|Q7X7A7\|Q7X7A7\_CHLRE |  |  | PEAKS DB |
| GAAAGAGGGAAVGATMGSGAT | 19.24 | 1561.7155 | 21 | -4.1 | 781.8618 | 21.00 | 1 | 4375 | JVDM\_BotE.raw | 1.4621E4 | 1 | 1 |  |  |  | PEAKS DB |
| EAAGGTAAGGGEAAEGWYGGGRR | 19.18 | 2106.9468 | 23 | 8.0 | 703.3285 | 25.78 | 1 | 5804 | JVDM\_BotE.raw | 2.7279E4 | 1 | 1 |  |  |  | PEAKS DB |
| TAPSGGGGVELGGAALC(+57.02)GGGGGGGGGGGGEGGR | 19.18 | 2556.1370 | 33 | -3.5 | 853.0499 | 19.05 | 1 | 3812 | JVDM\_BotE.raw | 0 | 0 | 0 |  | Carbamidomethylation | C17:Carbamidomethylation:1000.00 | PEAKS DB |
| QGAAAVGAGGAVTASPFAAAAVGAGR | 19.13 | 2155.1133 | 26 | 9.9 | 719.3855 | 27.60 | 1 | 6271 | JVDM\_BotE.raw | 0 | 0 | 0 |  |  |  | PEAKS DB |
| SAGAGAGGGAGGGAPLTPGQAAVAAYR | 19.09 | 2212.0984 | 27 | 3.4 | 738.3759 | 29.14 | 1 | 6686 | JVDM\_BotE.raw | 0 | 0 | 0 |  |  |  | PEAKS DB |
| ATAAAGSSAGSGGGGGGGGK | 19.00 | 1433.6494 | 20 | 7.7 | 717.8375 | 15.26 | 1 | 2731 | JVDM\_BotE.raw | 1.263E4 | 1 | 1 |  |  |  | PEAKS DB |
| GGGGGGGGKGAAGGGRGGGGGGGGGR | 18.95 | 1797.8326 | 26 | -5.9 | 899.9183 | 17.53 | 1 | 3346 | JVDM\_BotE.raw | 0 | 0 | 0 |  |  |  | PEAKS DB |
| GTAGGGAVAAATAHPAAAAAMAR | 18.94 | 1920.9587 | 23 | 6.5 | 961.4929 | 38.02 | 1 | 8901 | JVDM\_BotE.raw | 2.3548E4 | 1 | 1 |  |  |  | PEAKS DB |
| GLGGGGGGGGGGGGGGGAGRK | 18.92 | 1455.6926 | 21 | -0.9 | 728.8529 | 21.19 | 1 | 4422 | JVDM\_BotE.raw | 0 | 0 | 0 |  |  |  | PEAKS DB |
| WPAAAGGPGGGGGGALRSGLAGNSR | 18.91 | 2150.0728 | 25 | -6.2 | 1076.0370 | 36.82 | 1 | 8614 | JVDM\_BotE.raw | 0 | 0 | 0 |  |  |  | PEAKS DB |
| KHASGSSPGTAAAAAGAAATT | 18.87 | 1754.8547 | 21 | -4.3 | 585.9564 | 23.55 | 1 | 5146 | JVDM\_BotE.raw | 0 | 0 | 0 |  |  |  | PEAKS DB |
| SNAGVGAGAGAGGGAAK | 18.75 | 1271.6218 | 17 | -0.7 | 636.8177 | 14.80 | 1 | 2584 | JVDM\_BotE.raw | 3.6588E5 | 1 | 1 |  |  |  | PEAKS DB |
| GASSTPSSGSK | 18.69 | 964.4461 | 11 | 9.8 | 483.2351 | 13.03 | 1 | 2120 | JVDM\_BotE.raw | 2.1858E4 | 1 | 1 |  |  |  | PEAKS DB |
| GSGAAAGADGSTQAKK | 18.61 | 1375.6691 | 16 | 8.2 | 459.5674 | 12.71 | 1 | 1812 | JVDM\_BotE.raw | 0 | 0 | 0 |  |  |  | PEAKS DB |
| EALAAAGGGGGPPPPQ | 18.50 | 1345.6626 | 16 | -6.8 | 673.8340 | 19.53 | 1 | 3946 | JVDM\_BotE.raw | 2.156E5 | 1 | 1 |  |  |  | PEAKS DB |
| WLANMAGVNASQAR | 18.49 | 1487.7303 | 14 | -5.6 | 744.8683 | 15.93 | 1 | 2944 | JVDM\_BotE.raw | 2.118E5 | 1 | 1 |  |  |  | PEAKS DB |
| AGAGAGAAGTGGLSDGGR | 18.47 | 1401.6597 | 18 | 3.8 | 701.8398 | 23.65 | 1 | 5174 | JVDM\_BotE.raw | 8.0886E4 | 1 | 1 |  |  |  | PEAKS DB |
| TFNDALADAK | 18.42 | 1064.5138 | 10 | 0.3 | 533.2643 | 17.43 | 1 | 3335 | JVDM\_BotE.raw | 6.6525E4 | 1 | 1 |  |  |  | PEAKS DB |
| GAAAAAAVPHAGAAAGAGP | 18.34 | 1457.7374 | 19 | 5.5 | 729.8800 | 31.85 | 1 | 7397 | JVDM\_BotE.raw | 0 | 0 | 0 |  |  |  | PEAKS DB |
| LGAAAAASAAGGK | 18.27 | 1014.5458 | 13 | -9.4 | 508.2754 | 13.53 | 1 | 2259 | JVDM\_BotE.raw | 0 | 0 | 0 |  |  |  | PEAKS DB |
| GAGGAAGAGAPASPPPAPVK | 18.25 | 1599.8368 | 20 | 6.7 | 800.9310 | 33.15 | 1 | 7715 | JVDM\_BotE.raw | 0 | 0 | 0 |  |  |  | PEAKS DB |
| AGPGGGGGGGRGSPAGGGRR | 18.19 | 1593.7832 | 20 | 4.9 | 797.9028 | 31.83 | 1 | 7388 | JVDM\_BotE.raw | 0 | 0 | 0 |  |  |  | PEAKS DB |
| EAAPAPLQAAAAAAASEEAESVAAEPQEAAAP | 18.17 | 2959.4045 | 32 | 8.3 | 740.8646 | 28.31 | 1 | 6438 | JVDM\_BotE.raw | 0 | 0 | 0 |  |  |  | PEAKS DB |
| GGPAGAGAAPLP | 18.17 | 934.4872 | 12 | 0.1 | 468.2509 | 19.92 | 1 | 4082 | JVDM\_BotE.raw | 1.2249E5 | 1 | 1 |  |  |  | PEAKS DB |
| GADGAKGGPPGKDSAVAAAAAAAAR | 18.07 | 2107.0769 | 25 | -4.3 | 703.3632 | 35.27 | 1 | 8264 | JVDM\_BotE.raw | 0 | 0 | 0 |  |  |  | PEAKS DB |
| AGGGGGPAAGGGGGGGGGGGGGG | 18.03 | 1411.5824 | 23 | -7.8 | 706.7930 | 16.56 | 1 | 3054 | JVDM\_BotE.raw | 2.1463E4 | 1 | 1 |  |  |  | PEAKS DB |
| AGAGPPSAAAAIGGSR | 18.02 | 1309.6738 | 16 | -7.1 | 655.8395 | 12.70 | 1 | 1809 | JVDM\_BotE.raw | 0 | 0 | 0 |  |  |  | PEAKS DB |
| SGGGGGAAGGLGTSAR | 17.99 | 1231.5905 | 16 | 2.7 | 616.8041 | 13.98 | 1 | 2668 | JVDM\_BotE.raw | 6.6079E5 | 2 | 2 |  |  |  | PEAKS DB |
| LGAAGSGGGGGFGSGSPR | 17.97 | 1447.6803 | 18 | 9.0 | 724.8539 | 22.27 | 1 | 4737 | JVDM\_BotE.raw | 0 | 0 | 0 |  |  |  | PEAKS DB |
| MPTGGAAGLGAGAGGGGGAGRGGGR | 17.91 | 1925.9238 | 25 | 8.6 | 642.9874 | 15.33 | 1 | 2744 | JVDM\_BotE.raw | 0 | 0 | 0 |  |  |  | PEAKS DB |
| DAAAGAAAQAQ(+.98)TAAAGLMGSLN(+.98)AAAGK | 17.89 | 2330.1172 | 27 | -4.6 | 777.7094 | 35.27 | 1 | 8197 | JVDM\_BotE.raw | 2.704E6 | 1 | 1 | tr\|A0A2K3D8I0\|A0A2K3D8I0\_CHLRE | Deamidation (NQ) | Q11:Deamidation (NQ):0.00;N22:Deamidation (NQ):3.30 | PEAKS PTM |
| AGGAPAAGGGAGGGQAGGNSRR | 17.88 | 1752.8363 | 22 | -1.0 | 877.4246 | 25.53 | 1 | 5743 | JVDM\_BotE.raw | 0 | 0 | 0 |  |  |  | PEAKS DB |
| LVHDQELSVEER | 17.81 | 1452.7208 | 12 | 1.4 | 485.2482 | 12.96 | 1 | 2057 | JVDM\_BotE.raw | 0 | 0 | 0 |  |  |  | PEAKS DB |
| NILFVIQSPDVFK | 17.73 | 1518.8446 | 13 | 2.6 | 760.4316 | 39.18 | 1 | 9155 | JVDM\_BotE.raw | 3.0111E4 | 1 | 1 | tr\|A0A2K3CWF7\|A0A2K3CWF7\_CHLRE |  |  | PEAKS DB |
| GAAAAVAAAAPQGVK | 17.70 | 1251.6935 | 15 | 4.3 | 626.8567 | 19.09 | 1 | 3824 | JVDM\_BotE.raw | 0 | 0 | 0 |  |  |  | PEAKS DB |
| RAPGSFK | 17.70 | 761.4184 | 7 | 0.2 | 381.7166 | 17.77 | 1 | 3421 | JVDM\_BotE.raw | 0 | 0 | 0 |  |  |  | PEAKS DB |
| TDEAK(+42.01)AAAAELT(+79.97)LAAYK | 17.70 | 1857.8761 | 17 | 2.6 | 929.9478 | 30.69 | 1 | 7100 | JVDM\_BotE.raw | 2.6137E5 | 1 | 1 | tr\|Q7X7A7\|Q7X7A7\_CHLRE | Acetylation (K); Phosphorylation (STY) | K5:Acetylation (K):8.68;T12:Phosphorylation (STY):0.00 | PEAKS PTM |
| GAGGSGAGGSGAGGSRPASR | 17.69 | 1572.7352 | 20 | -8.7 | 787.3680 | 12.58 | 1 | 1661 | JVDM\_BotE.raw | 0 | 0 | 0 |  |  |  | PEAKS DB |
| AGGGAGGAGGGGGGRASSL | 17.64 | 1372.6443 | 19 | -7.3 | 687.3244 | 16.56 | 1 | 3067 | JVDM\_BotE.raw | 0 | 0 | 0 |  |  |  | PEAKS DB |
| GTAAAAGGAPAPGHK | 17.62 | 1232.6261 | 15 | -8.3 | 617.3152 | 16.98 | 1 | 3223 | JVDM\_BotE.raw | 9.0267E4 | 1 | 1 |  |  |  | PEAKS DB |
| ASALADAGAGGSR | 17.60 | 1102.5366 | 13 | -8.0 | 552.2712 | 14.39 | 1 | 2497 | JVDM\_BotE.raw | 1.026E5 | 1 | 1 |  |  |  | PEAKS DB |
| MAAAAGAGGGAGGAGGGGR | 17.59 | 1372.6266 | 19 | 1.5 | 687.3216 | 27.12 | 1 | 6143 | JVDM\_BotE.raw | 4.3407E5 | 1 | 1 |  |  |  | PEAKS DB |
| VRSALPGGGGGGGGGGAGSR | 17.57 | 1582.7924 | 20 | 7.0 | 792.4090 | 21.14 | 1 | 4407 | JVDM\_BotE.raw | 0 | 0 | 0 |  |  |  | PEAKS DB |
| KPGAGARSGGGGGGGGGGGSK | 17.56 | 1584.7716 | 21 | -4.0 | 793.3899 | 17.57 | 1 | 3357 | JVDM\_BotE.raw | 0 | 0 | 0 |  |  |  | PEAKS DB |
| GGGGGIGYGGGGSVSGR | 17.55 | 1350.6276 | 17 | 1.9 | 676.3223 | 14.54 | 1 | 2549 | JVDM\_BotE.raw | 0 | 0 | 0 |  |  |  | PEAKS DB |
| AGAAPAAGGGGGPSC(+57.02)GLGLGFGFSGGGVGGGVFGPQ | 17.47 | 2961.3826 | 36 | 7.3 | 988.1420 | 30.23 | 1 | 6977 | JVDM\_BotE.raw | 0 | 0 | 0 |  | Carbamidomethylation | C15:Carbamidomethylation:1000.00 | PEAKS DB |
| ELALYSAKIAEQSERYQDMVEEMK(+28.03) | 17.46 | 2888.3936 | 24 | -1.9 | 963.8033 | 36.91 | 1 | 8641 | JVDM\_BotE.raw | 1.2065E4 | 1 | 1 | tr\|Q7X7A7\|Q7X7A7\_CHLRE | Dimethylation(KR) | K24:Dimethylation(KR):13.91 | PEAKS PTM |
| TGGSGGGGGGGGGGSSQGTAAGAAVITPK | 17.42 | 2230.0574 | 29 | 2.3 | 744.3615 | 29.10 | 1 | 6678 | JVDM\_BotE.raw | 5.2607E4 | 1 | 1 |  |  |  | PEAKS DB |
| DAAASAAGKAVGGSSTGGNGK | 17.34 | 1732.8340 | 21 | -2.5 | 867.4221 | 21.19 | 1 | 4398 | JVDM\_BotE.raw | 3.6015E5 | 1 | 1 |  |  |  | PEAKS DB |
| PTAATAGGGGSAGGATAGAATATAPPPLAPVR | 17.32 | 2644.3567 | 32 | -1.6 | 882.4581 | 40.43 | 1 | 9456 | JVDM\_BotE.raw | 0 | 0 | 0 |  |  |  | PEAKS DB |
| TGPSLRLGGGLGSGGGR | 17.29 | 1497.8011 | 17 | 3.2 | 500.2759 | 18.84 | 1 | 3750 | JVDM\_BotE.raw | 1.0096E5 | 1 | 1 |  |  |  | PEAKS DB |
| RARNGSAGGTGASASAGK | 17.29 | 1574.7872 | 18 | -8.4 | 788.3943 | 35.27 | 1 | 8279 | JVDM\_BotE.raw | 1.1209E4 | 1 | 1 |  |  |  | PEAKS DB |
| EAALAEVGGGAAAPGTAAAAAGTAAAGS | 17.29 | 2211.0767 | 28 | 8.1 | 738.0388 | 28.06 | 1 | 6481 | JVDM\_BotE.raw | 4.1822E6 | 1 | 1 |  |  |  | PEAKS DB |
| TGVAVAGSGGSASGK | 17.28 | 1204.6047 | 15 | -2.4 | 603.3082 | 20.35 | 1 | 4185 | JVDM\_BotE.raw | 0 | 0 | 0 |  |  |  | PEAKS DB |
| EVRSSGSGSAAVGATA | 17.26 | 1405.6797 | 16 | -1.1 | 703.8464 | 38.27 | 1 | 8970 | JVDM\_BotE.raw | 0 | 0 | 0 |  |  |  | PEAKS DB |
| MNSVALENK | 17.22 | 1004.4961 | 9 | -1.2 | 503.2547 | 17.89 | 1 | 3448 | JVDM\_BotE.raw | 2.8118E5 | 1 | 1 |  |  |  | PEAKS DB |
| DAGAAILGQLPAGAAAGAAAAAAAGDTGAG | 17.13 | 2378.1826 | 30 | 8.4 | 793.7415 | 35.45 | 1 | 8277 | JVDM\_BotE.raw | 2.9047E5 | 1 | 1 |  |  |  | PEAKS DB |
| QNAADALAGMGGISK | 17.13 | 1402.6874 | 15 | 0.5 | 702.3513 | 19.79 | 1 | 4032 | JVDM\_BotE.raw | 5.0873E4 | 1 | 1 |  |  |  | PEAKS DB |
| AGGAAGAAGAGGAAGGAAGGAAGGK | 17.09 | 1682.8083 | 25 | 3.5 | 842.4144 | 17.13 | 1 | 3243 | JVDM\_BotE.raw | 0 | 0 | 0 |  |  |  | PEAKS DB |
| RGSGAGAAPKGLGR | 17.05 | 1253.6952 | 14 | -1.5 | 627.8539 | 35.34 | 1 | 8281 | JVDM\_BotE.raw | 0 | 0 | 0 |  |  |  | PEAKS DB |
| GGGSGSGGGASASASGR | 17.03 | 1278.5548 | 17 | -1.3 | 640.2839 | 17.40 | 1 | 3290 | JVDM\_BotE.raw | 1.5417E5 | 1 | 1 |  |  |  | PEAKS DB |
| ASPGGGGGTAAGGPGAPAPTGVR | 16.95 | 1818.8972 | 23 | 3.1 | 607.3082 | 22.86 | 1 | 4921 | JVDM\_BotE.raw | 0 | 0 | 0 |  |  |  | PEAKS DB |
| TPGSGGGGAIIVR | 16.90 | 1140.6251 | 13 | -6.3 | 571.3162 | 14.80 | 1 | 2677 | JVDM\_BotE.raw | 1.9555E5 | 1 | 1 |  |  |  | PEAKS DB |
| GAGAGAGAGAGVTSSAVTGSGAGGGAGGGGGGGSGPGSK | 16.90 | 2787.2766 | 39 | -0.3 | 930.0992 | 22.87 | 1 | 4924 | JVDM\_BotE.raw | 0 | 0 | 0 |  |  |  | PEAKS DB |
| IGSGAAVGMAAR | 16.90 | 1059.5494 | 12 | 7.1 | 530.7858 | 23.91 | 1 | 5252 | JVDM\_BotE.raw | 0 | 0 | 0 |  |  |  | PEAKS DB |
| EAGVTEAAATDGATSPAAPLPPR | 16.86 | 2149.0649 | 23 | -2.5 | 1075.5371 | 36.65 | 1 | 8580 | JVDM\_BotE.raw | 4.6835E4 | 1 | 1 |  |  |  | PEAKS DB |
| PGGGGGGGGGAGGGGGAGSAQR | 16.86 | 1554.6882 | 22 | 9.0 | 778.3584 | 16.94 | 1 | 3168 | JVDM\_BotE.raw | 1.4809E5 | 1 | 1 |  |  |  | PEAKS DB |
| AAGAGAGSGGGGAAAPR | 16.85 | 1254.6064 | 17 | 2.4 | 628.3120 | 26.91 | 1 | 6088 | JVDM\_BotE.raw | 6.339E4 | 1 | 1 |  |  |  | PEAKS DB |
| DDEDGALGAKGGGVRK | 16.84 | 1543.7590 | 16 | 9.4 | 772.8940 | 26.14 | 1 | 5913 | JVDM\_BotE.raw | 3.0254E4 | 1 | 1 |  |  |  | PEAKS DB |
| LTNSSVSTGLEGVVAVAR | 16.83 | 1758.9475 | 18 | -3.8 | 880.4777 | 28.50 | 1 | 6486 | JVDM\_BotE.raw | 0 | 0 | 0 |  |  |  | PEAKS DB |
| AATGAAASAQS(+79.97)AAAAAVAAAGDK | 16.79 | 1951.9000 | 23 | 3.4 | 976.9606 | 27.34 | 1 | 6164 | JVDM\_BotE.raw | 2.7313E5 | 1 | 1 | tr\|A0A2K3D8I0\|A0A2K3D8I0\_CHLRE | Phosphorylation (STY) | S11:Phosphorylation (STY):5.08 | PEAKS PTM |
| ANGSSTDVAAAVIDSRPAQVSMR | 16.69 | 2302.1335 | 23 | 7.2 | 768.3907 | 27.19 | 1 | 6122 | JVDM\_BotE.raw | 9.7992E4 | 1 | 1 |  |  |  | PEAKS DB |
| RSDGGSLAGNGSTAGSK | 16.69 | 1520.7179 | 17 | 9.5 | 761.3734 | 25.87 | 1 | 5832 | JVDM\_BotE.raw | 0 | 0 | 0 |  |  |  | PEAKS DB |
| ASAVAATGGGSGGDAAAAHWHMARVKIDAAQAMER | 16.69 | 3391.6262 | 35 | -4.1 | 1131.5447 | 30.33 | 1 | 6990 | JVDM\_BotE.raw | 3.4552E5 | 1 | 1 |  |  |  | PEAKS DB |
| EAPAAAAGAGGGGGGGGGGGGR | 16.67 | 1567.7086 | 22 | -8.7 | 784.8547 | 28.79 | 1 | 6586 | JVDM\_BotE.raw | 1.4052E4 | 1 | 1 |  |  |  | PEAKS DB |
| GTAVLPYSSVVYGGGGVVGAGAAGAAGGGSGR | 16.62 | 2678.3411 | 32 | 9.7 | 893.7963 | 35.57 | 1 | 8329 | JVDM\_BotE.raw | 0 | 0 | 0 |  |  |  | PEAKS DB |
| MQELNSLR | 16.55 | 989.4964 | 8 | 1.9 | 495.7564 | 17.85 | 1 | 3432 | JVDM\_BotE.raw | 7.1666E5 | 1 | 1 |  |  |  | PEAKS DB |
| AAAQAQEAAAGAATQ(+.98)AK | 16.54 | 1528.7480 | 17 | -1.2 | 765.3804 | 22.04 | 1 | 4714 | JVDM\_BotE.raw | 9.5974E4 | 1 | 1 | tr\|A0A2K3D8I0\|A0A2K3D8I0\_CHLRE | Deamidation (NQ) | Q15:Deamidation (NQ):6.45 | PEAKS PTM |
| SGGGGGPGGSGGPGGSGGGVR | 16.53 | 1526.6821 | 21 | -2.1 | 764.3467 | 22.71 | 1 | 4867 | JVDM\_BotE.raw | 0 | 0 | 0 |  |  |  | PEAKS DB |
| GGAGGAGGSVGASVHPHIHSR | 16.49 | 1866.9197 | 21 | 2.4 | 623.3153 | 23.13 | 1 | 5013 | JVDM\_BotE.raw | 0 | 0 | 0 |  |  |  | PEAKS DB |
| GSGAGGSSGGGGGGGGGGGGGGGR | 16.49 | 1589.6526 | 24 | 0.1 | 795.8337 | 12.31 | 1 | 1364 | JVDM\_BotE.raw | 0 | 0 | 0 |  |  |  | PEAKS DB |
| INVGPTTPGAGAAAAGGGPGSAAGSPR | 16.44 | 2218.1089 | 27 | -2.7 | 740.3749 | 35.03 | 1 | 8209 | JVDM\_BotE.raw | 0 | 0 | 0 |  |  |  | PEAKS DB |
| RAAAIGGGGGGGGGGARGGR | 16.42 | 1567.8040 | 20 | -1.3 | 784.9083 | 26.99 | 1 | 6102 | JVDM\_BotE.raw | 0 | 0 | 0 |  |  |  | PEAKS DB |
| GAGGGAGPAGGGGGGGGGGGQLLR | 16.40 | 1750.8458 | 24 | -1.7 | 584.6216 | 12.32 | 1 | 1377 | JVDM\_BotE.raw | 0 | 0 | 0 |  |  |  | PEAKS DB |
| AGDAAAAGPVAATTSRPQQPSSGPTGALR | 16.36 | 2662.3423 | 29 | 7.5 | 888.4614 | 38.09 | 1 | 8912 | JVDM\_BotE.raw | 0 | 0 | 0 |  |  |  | PEAKS DB |
| TGLMAGAC(+57.02)LLVGLSNSGPAYAAK | 16.35 | 2221.1235 | 23 | 8.5 | 741.3881 | 24.59 | 1 | 5469 | JVDM\_BotE.raw | 1.8008E5 | 1 | 1 |  | Carbamidomethylation | C8:Carbamidomethylation:1000.00 | PEAKS DB |
| AVAAAAAAGPGSGHGVR | 16.34 | 1418.7378 | 17 | 3.3 | 710.3785 | 18.97 | 1 | 3790 | JVDM\_BotE.raw | 0 | 0 | 0 |  |  |  | PEAKS DB |
| TNIVLEATR | 16.33 | 1015.5662 | 9 | 1.2 | 508.7910 | 19.60 | 1 | 3971 | JVDM\_BotE.raw | 1.0673E5 | 1 | 1 |  |  |  | PEAKS DB |
| GAVPGTASAAGAGRDLTTTTTTSSSRR | 16.32 | 2549.2793 | 27 | 0.3 | 850.7673 | 36.20 | 1 | 8455 | JVDM\_BotE.raw | 2.0574E4 | 1 | 1 |  |  |  | PEAKS DB |
| TLDSAATALGAGGRPGMAAGGGGGG | 16.31 | 2028.9646 | 25 | 3.8 | 677.3314 | 24.97 | 1 | 5594 | JVDM\_BotE.raw | 0 | 0 | 0 |  |  |  | PEAKS DB |
| VSSGAGAGVDMGGAGPSSGSR | 16.30 | 1762.7904 | 21 | 5.7 | 882.4075 | 30.98 | 1 | 7209 | JVDM\_BotE.raw | 2.5337E4 | 1 | 1 |  |  |  | PEAKS DB |
| QPGGGGAAAAPTR | 16.29 | 1109.5577 | 13 | -5.3 | 555.7832 | 30.38 | 1 | 7019 | JVDM\_BotE.raw | 4.6502E4 | 1 | 1 |  |  |  | PEAKS DB |
| GAAGAGGGGGGGGGAAGAGGAAGAG | 16.28 | 1569.6880 | 25 | -6.1 | 785.8465 | 16.23 | 1 | 2973 | JVDM\_BotE.raw | 6.8393E4 | 1 | 1 |  |  |  | PEAKS DB |
| AGGGGAAGASTSVAAAK | 16.28 | 1302.6527 | 17 | -6.6 | 652.3293 | 18.74 | 1 | 3728 | JVDM\_BotE.raw | 4.1776E5 | 1 | 1 |  |  |  | PEAKS DB |
| EPAAAAGSGGGGGGGGGGER | 16.27 | 1527.6661 | 20 | -5.5 | 764.8362 | 28.84 | 1 | 6580 | JVDM\_BotE.raw | 1.573E5 | 1 | 1 |  |  |  | PEAKS DB |
| AVSGGGGGGAGGGGGGGGGGGGGGSGGGGGGLR | 16.25 | 2184.9604 | 33 | 8.2 | 729.3334 | 23.47 | 1 | 5226 | JVDM\_BotE.raw | 2.4747E6 | 1 | 1 |  |  |  | PEAKS DB |
| TLDTAAAAAAAAADTK | 16.24 | 1431.7205 | 16 | 5.0 | 716.8711 | 27.46 | 1 | 6241 | JVDM\_BotE.raw | 1.3228E5 | 1 | 1 |  |  |  | PEAKS DB |
| DGGVAAGAVAIAAATAA | 16.23 | 1355.7045 | 17 | 5.4 | 678.8632 | 20.05 | 1 | 4088 | JVDM\_BotE.raw | 3.2644E5 | 1 | 1 |  |  |  | PEAKS DB |
| AAAGGGGGGGGGGGPGAAGGGGR | 16.22 | 1538.6934 | 23 | 9.3 | 770.3611 | 24.21 | 1 | 5301 | JVDM\_BotE.raw | 4.153E5 | 1 | 1 |  |  |  | PEAKS DB |
| GGGGAGGGGGGGGGGSAHL | 16.21 | 1295.5603 | 19 | -6.7 | 648.7831 | 15.31 | 1 | 2734 | JVDM\_BotE.raw | 0 | 0 | 0 |  |  |  | PEAKS DB |
| NGGAGGGGGGAGGGGAEGGGAAAPR | 16.19 | 1795.7946 | 25 | -8.2 | 898.8972 | 20.72 | 1 | 4275 | JVDM\_BotE.raw | 5.1904E4 | 1 | 1 |  |  |  | PEAKS DB |
| RSAGAGSRGGAAAAAAGGGQ | 16.12 | 1599.7825 | 20 | -3.7 | 800.8956 | 18.16 | 1 | 3493 | JVDM\_BotE.raw | 9.0686E5 | 1 | 1 |  |  |  | PEAKS DB |
| AIADFGSQDK | 16.11 | 1050.4982 | 10 | 2.1 | 526.2574 | 15.68 | 1 | 2836 | JVDM\_BotE.raw | 6.119E4 | 1 | 1 | tr\|A0A2K3D661\|A0A2K3D661\_CHLRE |  |  | PEAKS DB |
| HGKGGGGGGGGGGIGGSGSK | 16.06 | 1496.7080 | 20 | -0.5 | 749.3609 | 12.59 | 1 | 1670 | JVDM\_BotE.raw | 0 | 0 | 0 |  |  |  | PEAKS DB |
| GGGAAGAQGRGLSGSGMAGAGPFPVAGGR | 15.98 | 2427.1824 | 29 | 6.6 | 607.8069 | 24.32 | 1 | 5269 | JVDM\_BotE.raw | 4.9763E5 | 1 | 1 |  |  |  | PEAKS DB |
| RAAAAAAGAANGGGSGGGGDGGGSGQAAR | 15.96 | 2256.0339 | 29 | -6.0 | 753.0141 | 24.35 | 1 | 5456 | JVDM\_BotE.raw | 5.9147E5 | 1 | 1 |  |  |  | PEAKS DB |
| LGGAGGAGGGAGGGGGRR | 15.95 | 1340.6656 | 18 | 3.0 | 671.3421 | 16.63 | 1 | 3104 | JVDM\_BotE.raw | 7.9647E5 | 1 | 1 |  |  |  | PEAKS DB |
| AAGAAAAAAPAGLTGSSVAAR | 15.95 | 1710.9012 | 21 | -5.9 | 571.3043 | 17.99 | 1 | 3502 | JVDM\_BotE.raw | 3.2456E4 | 1 | 1 |  |  |  | PEAKS DB |
| TSLSGSGGGGGGGGAGGGTSGSTMGSGGGGGGGR | 15.93 | 2512.0593 | 34 | -6.0 | 838.3553 | 12.34 | 1 | 1399 | JVDM\_BotE.raw | 0 | 0 | 0 |  |  |  | PEAKS DB |
| AGGGGAAATAAGGGGGGGAAVG | 15.91 | 1470.6810 | 22 | 2.7 | 736.3498 | 24.63 | 1 | 5482 | JVDM\_BotE.raw | 0 | 0 | 0 |  |  |  | PEAKS DB |
| LSRGGPGAGPAGPR | 15.86 | 1248.6687 | 14 | 2.8 | 625.3434 | 33.81 | 1 | 7908 | JVDM\_BotE.raw | 0 | 0 | 0 |  |  |  | PEAKS DB |
| PAGGGVGTGAGGGAGGAGAGGR | 15.84 | 1567.7451 | 22 | -7.7 | 784.8738 | 25.43 | 1 | 5709 | JVDM\_BotE.raw | 0 | 0 | 0 |  |  |  | PEAKS DB |
| AAAGAAGGGGR | 15.81 | 814.4045 | 11 | 9.3 | 408.2133 | 14.98 | 1 | 2666 | JVDM\_BotE.raw | 1.1932E5 | 1 | 1 |  |  |  | PEAKS DB |
| TAAGYGQDGPGGAGGSPLR | 15.75 | 1687.7914 | 19 | 9.4 | 563.6097 | 13.98 | 1 | 2373 | JVDM\_BotE.raw | 5.25E4 | 1 | 1 |  |  |  | PEAKS DB |
| EGGGGGGGGGGGGGGGGFGGGGR | 15.74 | 1590.6519 | 23 | 1.9 | 796.3347 | 12.42 | 1 | 1495 | JVDM\_BotE.raw | 0 | 0 | 0 |  |  |  | PEAKS DB |
| AGPGAGQGGAAAAAAGAGGGGGGGGGGGAVGSAV | 15.73 | 2336.0852 | 34 | 4.6 | 779.7059 | 20.32 | 1 | 3983 | JVDM\_BotE.raw | 1.8921E6 | 1 | 1 |  |  |  | PEAKS DB |
| GGGPGGAGAAQGGGGGGGGK | 15.69 | 1382.6287 | 20 | 6.8 | 692.3263 | 22.10 | 1 | 4712 | JVDM\_BotE.raw | 1.2353E5 | 1 | 1 |  |  |  | PEAKS DB |
| VGGTAAEAQAGLLR | 15.69 | 1312.7098 | 14 | -5.7 | 657.3585 | 23.02 | 1 | 4988 | JVDM\_BotE.raw | 2.4515E4 | 1 | 1 |  |  |  | PEAKS DB |
| PAAGGGAGAGGGGGSAATGQGAAAGGAATGAAAGR | 15.68 | 2538.1919 | 35 | 5.7 | 847.0760 | 23.55 | 1 | 5145 | JVDM\_BotE.raw | 0 | 0 | 0 |  |  |  | PEAKS DB |
| GAGAAAAAAGASGTAAAATGAP | 15.68 | 1612.7804 | 22 | 3.7 | 807.4005 | 17.71 | 1 | 3400 | JVDM\_BotE.raw | 0 | 0 | 0 |  |  |  | PEAKS DB |
| GSGGGVAAAGAVGAVSLQQGAAAAGNVA | 15.68 | 2238.1353 | 28 | -0.6 | 747.0519 | 28.06 | 1 | 6430 | JVDM\_BotE.raw | 1.7673E6 | 1 | 1 |  |  |  | PEAKS DB |
| DAAGATPLAAAAAAR | 15.66 | 1296.6786 | 15 | 5.9 | 649.3504 | 18.82 | 1 | 3745 | JVDM\_BotE.raw | 0 | 0 | 0 |  |  |  | PEAKS DB |
| SVGGSGGGGGGGGSTLR | 15.64 | 1318.6226 | 17 | 2.3 | 660.3201 | 13.93 | 1 | 2362 | JVDM\_BotE.raw | 3.3364E5 | 1 | 1 |  |  |  | PEAKS DB |
| AGGPGGGGGAGGLGAR | 15.64 | 1167.5745 | 16 | -8.4 | 584.7896 | 18.51 | 1 | 3666 | JVDM\_BotE.raw | 0 | 0 | 0 |  |  |  | PEAKS DB |
| WGSGVIVGGGGAADPIGLGGGSSGSGGGGGGGAATR | 15.63 | 2869.3701 | 36 | -9.8 | 718.3428 | 28.79 | 1 | 6598 | JVDM\_BotE.raw | 2.0917E5 | 1 | 1 |  |  |  | PEAKS DB |
| ENAGAHAPGAGK | 15.63 | 1078.5155 | 12 | -6.9 | 360.5100 | 13.62 | 1 | 2263 | JVDM\_BotE.raw | 3.5022E5 | 1 | 1 |  |  |  | PEAKS DB |
| RGGGGAAGGGGGPGGGPR | 15.63 | 1350.6500 | 18 | -8.5 | 676.3265 | 21.62 | 1 | 4541 | JVDM\_BotE.raw | 0 | 0 | 0 |  |  |  | PEAKS DB |
| AGVGPGAGAAAAGVGVSAVSGKKR | 15.62 | 1994.1021 | 24 | 4.8 | 499.5352 | 24.25 | 1 | 5230 | JVDM\_BotE.raw | 2.4793E6 | 1 | 1 |  |  |  | PEAKS DB |
| KVGGGGGGGGAGGGGGHK | 15.62 | 1322.6439 | 18 | -3.0 | 662.3273 | 25.69 | 1 | 5786 | JVDM\_BotE.raw | 1.2107E4 | 1 | 1 |  |  |  | PEAKS DB |
| GRASSAGAGGPGGSGGGGVK | 15.62 | 1542.7498 | 20 | 9.9 | 772.3898 | 26.73 | 1 | 6032 | JVDM\_BotE.raw | 0 | 0 | 0 |  |  |  | PEAKS DB |
| LAGSALLSAGGVSSGAGSGGPNSAEIAAASR | 15.61 | 2642.3259 | 31 | -4.8 | 881.7784 | 23.35 | 1 | 5083 | JVDM\_BotE.raw | 0 | 0 | 0 |  |  |  | PEAKS DB |
| SVGVGISGGGAGGS | 15.58 | 1060.5149 | 14 | -5.9 | 531.2616 | 19.68 | 1 | 3954 | JVDM\_BotE.raw | 2.758E6 | 1 | 1 |  |  |  | PEAKS DB |
| AAGTAAAGTAAAAGGALQ | 15.58 | 1399.7056 | 18 | 0.4 | 700.8604 | 22.25 | 1 | 4725 | JVDM\_BotE.raw | 4.7338E4 | 1 | 1 |  |  |  | PEAKS DB |
| LAGGGAGGAAGGGTPPR | 15.52 | 1322.6691 | 17 | -0.8 | 662.3413 | 14.90 | 1 | 2866 | JVDM\_BotE.raw | 2.454E6 | 1 | 1 |  |  |  | PEAKS DB |
| GGGAGGASSSGGGGGGGK | 15.51 | 1233.5333 | 18 | 4.5 | 617.7767 | 12.64 | 1 | 1735 | JVDM\_BotE.raw | 0 | 0 | 0 |  |  |  | PEAKS DB |
| QQAMAHGNGGAGGSR | 15.51 | 1397.6218 | 15 | -9.5 | 699.8115 | 14.62 | 1 | 2547 | JVDM\_BotE.raw | 4.3637E4 | 1 | 1 |  |  |  | PEAKS DB |
| EGGGGSTLGSAAVAQR | 15.48 | 1416.6957 | 16 | -6.7 | 473.2360 | 16.63 | 1 | 3087 | JVDM\_BotE.raw | 2.7696E5 | 1 | 1 |  |  |  | PEAKS DB |
| GAGAGGPTSGGPGQGVGAGAG | 15.47 | 1538.7073 | 21 | 0.3 | 770.3611 | 24.21 | 1 | 5410 | JVDM\_BotE.raw | 4.153E5 | 1 | 1 |  |  |  | PEAKS DB |
| QLQPAPAPPTAASR | 15.46 | 1403.7521 | 14 | 2.1 | 702.8848 | 25.08 | 1 | 5621 | JVDM\_BotE.raw | 6.8822E3 | 1 | 1 |  |  |  | PEAKS DB |
| GAAGGGGGGGGGADAALVLELSASVDK | 15.46 | 2213.0923 | 27 | -4.6 | 738.7013 | 29.01 | 1 | 6686 | JVDM\_BotE.raw | 2.9843E5 | 1 | 1 |  |  |  | PEAKS DB |
| EAAAAGGGGGRTK | 15.45 | 1101.5526 | 13 | -9.8 | 551.7782 | 12.57 | 1 | 1645 | JVDM\_BotE.raw | 0 | 0 | 0 |  |  |  | PEAKS DB |
| GGGGGGGGAFMAASK | 15.43 | 1180.5294 | 15 | 7.3 | 591.2763 | 18.15 | 1 | 3489 | JVDM\_BotE.raw | 3.162E5 | 1 | 1 |  |  |  | PEAKS DB |
| GGGGGGGGGAGGAGGRK | 15.42 | 1185.5598 | 17 | 5.2 | 593.7903 | 20.98 | 1 | 4371 | JVDM\_BotE.raw | 1.2012E5 | 1 | 1 |  |  |  | PEAKS DB |
| DGGGGGAAPGRVGG | 15.42 | 1083.5057 | 14 | 0.0 | 542.7601 | 24.00 | 1 | 5267 | JVDM\_BotE.raw | 3.006E4 | 1 | 1 |  |  |  | PEAKS DB |
| QAAAAAAEASPPPPPTSGRGR | 15.35 | 1958.9922 | 21 | -7.8 | 653.9996 | 33.00 | 1 | 7674 | JVDM\_BotE.raw | 0 | 0 | 0 |  |  |  | PEAKS DB |
| GAAPVAVAGAVAGAGVAAAGAAAGGEGGAEGG | 15.29 | 2391.1777 | 32 | 8.8 | 798.0735 | 27.98 | 1 | 6364 | JVDM\_BotE.raw | 0 | 0 | 0 |  |  |  | PEAKS DB |
| FITGLAPVNALVK | 15.29 | 1341.8020 | 13 | 5.3 | 448.2770 | 37.06 | 1 | 8718 | JVDM\_BotE.raw | 2.8448E5 | 1 | 1 |  |  |  | PEAKS DB |
| KAAASSGAGAAGGKAK | 15.29 | 1301.7051 | 16 | 4.5 | 651.8627 | 38.07 | 1 | 8910 | JVDM\_BotE.raw | 0 | 0 | 0 |  |  |  | PEAKS DB |
| TAAAGTVVGGAAGTAVGGGR | 15.29 | 1599.8329 | 20 | -9.4 | 800.9162 | 27.00 | 1 | 6103 | JVDM\_BotE.raw | 0 | 0 | 0 |  |  |  | PEAKS DB |
| TGGGGNGGGGGGGVGASGSLLFHTAR | 15.28 | 2157.0310 | 26 | 1.5 | 720.0187 | 28.95 | 1 | 6633 | JVDM\_BotE.raw | 1.7608E5 | 1 | 1 |  |  |  | PEAKS DB |
| EASAGPAAAAAAAAAGAS | 15.26 | 1384.6582 | 18 | -4.0 | 693.3336 | 28.95 | 1 | 6624 | JVDM\_BotE.raw | 0 | 0 | 0 |  |  |  | PEAKS DB |
| TGTGAGGGGGTVAGGSPR | 15.26 | 1415.6753 | 18 | 10.0 | 708.8520 | 19.53 | 1 | 3950 | JVDM\_BotE.raw | 3.2916E4 | 1 | 1 |  |  |  | PEAKS DB |
| DGGASGSAR | 15.25 | 776.3412 | 9 | 6.8 | 389.1805 | 12.63 | 1 | 1723 | JVDM\_BotE.raw | 0 | 0 | 0 |  |  |  | PEAKS DB |
| KGGVASRSPGQR | 15.25 | 1198.6531 | 12 | 4.6 | 600.3365 | 32.76 | 1 | 7657 | JVDM\_BotE.raw | 1.5042E5 | 1 | 1 |  |  |  | PEAKS DB |
| GGPGGGAGGGGGGAAGSTADADR | 15.22 | 1728.7411 | 23 | 6.5 | 865.3834 | 25.43 | 1 | 5721 | JVDM\_BotE.raw | 5.3579E4 | 1 | 1 |  |  |  | PEAKS DB |
| VAGVPLAAGGQGGAGGAGGDGSGAGGGGAGSKPR | 15.21 | 2634.2856 | 34 | 3.3 | 879.1053 | 27.03 | 1 | 6125 | JVDM\_BotE.raw | 2.3968E4 | 1 | 1 |  |  |  | PEAKS DB |
| DVPPGAAGAGGPGGPGGF | 15.15 | 1436.6683 | 18 | 1.9 | 719.3428 | 23.87 | 1 | 5255 | JVDM\_BotE.raw | 3.0062E4 | 1 | 1 |  |  |  | PEAKS DB |
| DAKFAGGGGKAHSAAIAGGGTGGGGGR | 15.13 | 2241.0999 | 27 | -6.4 | 748.0358 | 35.99 | 1 | 8429 | JVDM\_BotE.raw | 0 | 0 | 0 |  |  |  | PEAKS DB |
| WAGAGGGAAAGAAAAVAPAAAPPEQGGPTGPAPLPVP | 15.12 | 3100.5730 | 37 | 4.6 | 776.1541 | 30.77 | 1 | 7136 | JVDM\_BotE.raw | 0 | 0 | 0 |  |  |  | PEAKS DB |
| AAGGGGGGGGSPAAPR | 15.10 | 1195.5693 | 16 | -7.1 | 598.7877 | 14.77 | 1 | 2724 | JVDM\_BotE.raw | 9.6986E5 | 1 | 1 |  |  |  | PEAKS DB |
| AGAAGVHGTALRVGGGPPAPPR | 15.06 | 1965.0656 | 22 | -4.5 | 656.0262 | 29.14 | 1 | 6917 | JVDM\_BotE.raw | 1.7466E7 | 1 | 1 |  |  |  | PEAKS DB |
| AGIATAGGGAAAAAMDAGGETGAASR | 15.06 | 2131.9917 | 26 | -0.3 | 711.6710 | 21.40 | 1 | 4503 | JVDM\_BotE.raw | 4.5813E4 | 1 | 1 |  |  |  | PEAKS DB |
| ASFGGGGVGGGGVGGGGGSR | 15.06 | 1505.6970 | 20 | -2.5 | 753.8539 | 20.75 | 1 | 4273 | JVDM\_BotE.raw | 7.1459E4 | 1 | 1 |  |  |  | PEAKS DB |
| RSGSAGGAAPEAPD | 15.05 | 1241.5636 | 14 | -3.6 | 621.7869 | 16.11 | 1 | 2920 | JVDM\_BotE.raw | 1.5133E5 | 1 | 1 |  |  |  | PEAKS DB |
| MAALGAGGGATGAAAAFVSA | 15.04 | 1620.7930 | 20 | -8.7 | 811.3967 | 24.39 | 1 | 5494 | JVDM\_BotE.raw | 3.2711E5 | 1 | 1 |  |  |  | PEAKS DB |
| AGGGGGGGAGSEVVDGVK | 15.02 | 1429.6797 | 18 | 2.1 | 715.8486 | 14.93 | 1 | 2637 | JVDM\_BotE.raw | 2.4194E5 | 1 | 1 |  |  |  | PEAKS DB |
| QQSLTDSAAAASAAQRAELQRR | 15.01 | 2328.1895 | 22 | -7.5 | 777.0646 | 28.43 | 1 | 6604 | JVDM\_BotE.raw | 2.3723E6 | 1 | 1 |  |  |  | PEAKS DB |
| total 358 peptides |  |  |  |  |  |  |  |  |  |  |  |  |  |  |  |  |
| --- | --- | --- | --- | --- | --- | --- | --- | --- | --- | --- | --- | --- | --- | --- | --- | --- |
