## Supplementary material for "Efficient secretion of a plastic degrading enzyme from the green algae *Chlamydomonas reinhardtii*": Dataset - https://doi.org/10.5281/zenodo.13981200: protein.html

proteins


Summary

  

### 1. Notes

### 2. Result Statistics

**Figure 1.**
False discovery rate (FDR) curve. X axis is the number of peptides being kept. Y axis is the corresponding FDR.

  


  

**Figure 2.**
PSM score distribution.  **(a)**
Distribution of PEAKS peptide score;  **(b)**
Scatterplot of PEAKS peptide score versus precursor mass error.

**(a)**

**(b)**

**Table 1.**
Statistics of data.


|  |  |  |  |  |  |  |  |  |  |  |  |  |
| --- | --- | --- | --- | --- | --- | --- | --- | --- | --- | --- | --- | --- |
|  | #Scans | | | #Features | Identified | | | #Peptides | #Sequences | #Proteins^\*^ | | |
|  | MS1 | MS/MS | #Chimera |  | #PSMs | #Scans | #Features^\*\*^ |  |  | Groups | All | Top |
| Total | 3377 | 8992 | 504 | 17937 | 392 | 390 | 233 | 358 | 355 | 30 | 42 | 42 |
| Sample 1 | 3377 | 8992 | 504 | 17937 | 392 | 390 | 233 | 358 | 355 | 30 | 42 | 42 |

^\* proteins with significant peptides are used in counts.^  
^\*\* features are identified by DB search only.^

**Figure 3.**
Sample overlap for Proteins and Peptides (up to 8 samples).  **(a)**
All Proteins;  **(b)**
Top Proteins;  **(c)**
Peptides;

**(a)**

**Not applicable to only one sample**

**(b)**

**Not applicable to only one sample**

**(c)**

**Not applicable to only one sample**

**Figure 4.**
Distribution of peptide feature detection.  **(a)**
Feature m/z distribution;  **(b)**
Feature RT distribution.

**(a)**

**(b)**

**Figure 5.**
Distribution of identified peptide features.  **(a)**
Feature abundance distribution;  **(b)**
 *De novo*
sequencing validation.

**(a)**

**(b)**

**Table 2.**
Result filtration parameters.

|  |  |
| --- | --- |
| Peptide -10lgP | ≥15 |
| PTM Ascore | ≥0 |
| Protein -10lgP | ≥20 |
| Proteins unique peptides | ≥0 |
| De novo score(%) | ≥50% |


**Table 3.**
Statistics of filtered result.

|  |  |
| --- | --- |
| FDR (Peptide-Spectrum Matches) | 34.2% |
| FDR (Peptide Sequences) | 36.0% |
| FDR (Protein Group) | 0.0% |
| De Novo Only Spectra | 2868 |

**Table 4.**
PTM profile.

| Name | ∆Mass | Position | #PSM | -10lgP | Abundance | AScore || Carbamidomethyl | 57.02 | C | 15 | 67.82 | 1.92E6 | 1000.00 |
| Deamidation | .98 | NQ | 12 | 48.69 |  | 0.00 |
| Phosphorylation | 79.97 | ST | 6 | 30.62 | 5.17E5 | 1000.00 |
| Methylation(KR) | 14.02 | KR | 5 | 39.38 |  | 1000.00 |

### 3. Experiment Control

**Figure 6.**
Precursor mass error of peptide-spectrum matches (PSM) in filtered result.  **(a)**
Distribution of precursor mass error in ppm;  **(b)**
Scatterplot of precursor m/z versus precursor mass error in ppm.

**(a)**

**(b)**

**Table 5.**
Number of identified peptides in each sample by the number of missed cleavages.

|  |  |  |  |  |  |
| --- | --- | --- | --- | --- | --- |
| Missed Cleavages | 0 | 1 | 2 | 3 | 4+ |
| Sample 1 | 270 | 63 | 25 | 0 | 0 |

### 4. Other Information

**Table 6.**
Search parameters.

PEAKS Version: PEAKS Studio 10.6 build 20201221  
Search Engine Name: PEAKS  
Parent Mass Error Tolerance: 10.0 ppm  
Fragment Mass Error Tolerance: 0.5 Da  
Precursor Mass Search Type: monoisotopic  
Enzyme: Trypsin  
Max Missed Cleavages: 2  
Digest Mode: Semispecific  
Fixed Modifications:   
  Carbamidomethylation: 57.02  
Variable Modifications:   
  Acetylation (K): 42.01  
  Deamidation (NQ): 0.98  
  Dimethylation(KR): 28.03  
  Dimethylation(NP): 28.03  
  Dioxidation (M): 31.99  
  Methylation(KR): 14.02  
  Phosphorylation (STY): 79.97  
  Biotinylation: 226.08  
Max Variable PTM Per Peptide: 3  
Database: Chlamydomonas reinhardtii uniprot  
Taxon: All  
Searched Entry: 15000  
FDR Estimation: Enabled  
De novo score(%) threshold: 15  
Peptide hit threshold (-10logP): 30.0  
Peaks run ID: 3  
Merge Options: no merge  
Precursor Options: corrected  
Charge Options: no correction  
Filter Charge: 2 - 8  
Process: true  
Associate chimera: yes

**Table 7.**
Instrument parameters.

Fractions: JVDM\_BotE.raw  
Ion Source: ESI(nano-spray)  
Fragmentation Mode: high energy CID (y and b ions)  
MS Scan Mode: FT-ICR/Orbitrap  
MS/MS Scan Mode: Linear Ion Trap

  

Protein List

  

Protein Accession Contains:

Protein Description Contains:

Protein Sample Area >=

Protein PTM Contains:

| Protein Group | Protein ID | Accession | -10lgP | Coverage (%) | Coverage (%) Sample 1 | Area Sample 1 | #Peptides | #Unique | #Spec Sample 1 | PTM | Avg. Mass | Description |
| --- | --- | --- | --- | --- | --- | --- | --- | --- | --- | --- | --- | --- |
| 1 | 2 | PHL7 | 214.08 | 44 | 44 | 1.7766E7 | 9 | 9 | 20 | Y | 28201 | PHL7 |
| 2 | 1 | tr\|A0A2K3D8I0\|A0A2K3D8I0\_CHLRE | 202.23 | 36 | 36 | 1.1301E7 | 16 | 16 | 21 | Y | 94346 | Uncharacterized protein OS=Chlamydomonas reinhardtii OX=3055 GN=CHLRE\_11g477950v5 PE=4 SV=1 |
| 3 | 3 | tr\|Q7X7A7\|Q7X7A7\_CHLRE | 173.07 | 41 | 41 | 3.0477E6 | 8 | 8 | 11 | Y | 28003 | 14-3-3 protein OS=Chlamydomonas reinhardtii OX=3055 GN=erb14 PE=2 SV=1 |
| 4 | 4 | tr\|A0A2K3D6N5\|A0A2K3D6N5\_CHLRE | 134.09 | 15 | 15 | 5.128E5 | 6 | 6 | 7 | Y | 81251 | Uncharacterized protein OS=Chlamydomonas reinhardtii OX=3055 GN=CHLRE\_12g544450v5 PE=4 SV=1 |
| 7 | 12 | tr\|A0A2K3DP12\|A0A2K3DP12\_CHLRE | 119.86 | 6 | 6 | 9.5323E4 | 3 | 3 | 3 | Y | 79150 | Peptidase\_M11 domain-containing protein OS=Chlamydomonas reinhardtii OX=3055 GN=CHLRE\_06g278165v5 PE=4 SV=1 |
| 13 | 18 | tr\|A8IKW6\|A8IKW6\_CHLRE | 112.87 | 12 | 12 | 1.3877E5 | 2 | 2 | 2 | N | 28401 | Ribulose-phosphate 3-epimerase OS=Chlamydomonas reinhardtii OX=3055 GN=RPE1 PE=3 SV=1 |
| 5 | 16 | tr\|A0A2K3CQ54\|A0A2K3CQ54\_CHLRE | 103.58 | 8 | 8 | 5.2529E6 | 4 | 4 | 6 | Y | 74085 | Peptidase\_M11 domain-containing protein OS=Chlamydomonas reinhardtii OX=3055 GN=CHLRE\_17g718500v5 PE=4 SV=1 |
| 11 | 8 | tr\|B7U1J0\|B7U1J0\_CHLRE | 101.47 | 7 | 7 | 3.1276E5 | 3 | 3 | 3 | N | 54752 | ATP synthase subunit alpha, chloroplastic OS=Chlamydomonas reinhardtii OX=3055 GN=atpA PE=3 SV=1 |
| 11 | 7 | P26526\|ATPA\_CHLRE | 101.47 | 7 | 7 | 3.1276E5 | 3 | 3 | 3 | N | 54752 | ATP synthase subunit alpha, chloroplastic OS=Chlamydomonas reinhardtii OX=3055 GN=atpA PE=1 SV=3 |
| 9 | 5 | tr\|Q9FE86\|Q9FE86\_CHLRE | 99.59 | 21 | 21 | 4.1504E5 | 3 | 3 | 3 | N | 25962 | Peroxiredoxin OS=Chlamydomonas reinhardtii OX=3055 GN=thioredoxin peroxidase PE=1 SV=1 |
| 12 | 19 | tr\|A8J9H8\|A8J9H8\_CHLRE | 99.18 | 17 | 17 | 5.5574E5 | 3 | 3 | 3 | N | 16541 | Nucleoside diphosphate kinase OS=Chlamydomonas reinhardtii OX=3055 GN=FAP103 PE=3 SV=1 |
| 12 | 20 | tr\|A0A2K3DJF1\|A0A2K3DJF1\_CHLRE | 99.18 | 16 | 16 | 5.5574E5 | 3 | 3 | 3 | N | 18054 | Nucleoside diphosphate kinase OS=Chlamydomonas reinhardtii OX=3055 GN=CHLRE\_07g325734v5 PE=3 SV=1 |
| 17 | 14 | tr\|A0A2K3CWF7\|A0A2K3CWF7\_CHLRE | 75.16 | 22 | 22 | 3.0111E4 | 3 | 3 | 3 | N | 21332 | NAC-A/B domain-containing protein OS=Chlamydomonas reinhardtii OX=3055 GN=CHLRE\_15g635600v5 PE=4 SV=1 |
| 6 | 6 | tr\|A0A2K3E0C1\|A0A2K3E0C1\_CHLRE | 74.56 | 10 | 10 | 1.0514E5 | 6 | 3 | 6 | Y | 86252 | Uncharacterized protein OS=Chlamydomonas reinhardtii OX=3055 GN=CHLRE\_02g077800v5 PE=4 SV=1 |
| 6 | 9 | tr\|A0A2K3E094\|A0A2K3E094\_CHLRE | 74.56 | 5 | 5 | 1.0514E5 | 3 | 1 | 3 | N | 81741 | Uncharacterized protein OS=Chlamydomonas reinhardtii OX=3055 GN=CHLRE\_02g077750v5 PE=4 SV=1 |
| 19 | 21 | tr\|A8IT01\|A8IT01\_CHLRE | 74.37 | 8 | 8 | 1.9599E5 | 2 | 2 | 2 | N | 41628 | Carbonic anhydrase OS=Chlamydomonas reinhardtii OX=3055 GN=CAH1 PE=3 SV=1 |
| 10 | 10 | tr\|A8JEV1\|A8JEV1\_CHLRE | 73.81 | 16 | 16 | 7.5464E5 | 3 | 3 | 3 | N | 21824 | Oxygen evolving enhancer protein 3 OS=Chlamydomonas reinhardtii OX=3055 GN=PSBQ PE=3 SV=1 |
| 8 | 13 | tr\|A0A2K3E0A7\|A0A2K3E0A7\_CHLRE | 71.07 | 8 | 8 | 1.2277E4 | 4 | 1 | 4 | Y | 83460 | Uncharacterized protein OS=Chlamydomonas reinhardtii OX=3055 GN=CHLRE\_02g077850v5 PE=4 SV=1 |
| 14 | 15 | tr\|A8IKZ2\|A8IKZ2\_CHLRE | 67.94 | 17 | 17 | 9.3761E5 | 2 | 2 | 3 | Y | 20930 | Ribosomal protein L18 OS=Chlamydomonas reinhardtii OX=3055 GN=RPL18 PE=3 SV=1 |
| 15 | 11 | tr\|A8IGH1\|A8IGH1\_CHLRE | 59.10 | 19 | 19 | 2.7308E5 | 4 | 4 | 4 | Y | 25915 | Superoxide dismutase OS=Chlamydomonas reinhardtii OX=3055 GN=FSD1 PE=2 SV=1 |
| 16 | 22 | tr\|A8HXL8\|A8HXL8\_CHLRE | 53.97 | 11 | 11 | 2.4019E5 | 3 | 3 | 3 | Y | 38761 | Chloroplast ATP synthase gamma chain OS=Chlamydomonas reinhardtii OX=3055 GN=ATPC PE=3 SV=1 |
| 25 | 30 | tr\|A8HVA3\|A8HVA3\_CHLRE | 49.18 | 10 | 10 | 1.685E5 | 1 | 1 | 1 | N | 11457 | Histone H4 OS=Chlamydomonas reinhardtii OX=3055 GN=HFO4 PE=3 SV=1 |
| 25 | 29 | tr\|A8HSB0\|A8HSB0\_CHLRE | 49.18 | 10 | 10 | 1.685E5 | 1 | 1 | 1 | N | 11485 | Histone H4 OS=Chlamydomonas reinhardtii OX=3055 GN=HFO2 PE=3 SV=1 |
| 24 | 27 | tr\|A0A2K3CRJ3\|A0A2K3CRJ3\_CHLRE | 49.10 | 6 | 6 | 5.6894E5 | 1 | 1 | 1 | N | 19601 | Uncharacterized protein OS=Chlamydomonas reinhardtii OX=3055 GN=CHLRE\_17g738000v5 PE=4 SV=1 |
| 24 | 26 | tr\|Q1ALD7\|Q1ALD7\_CHLRE | 49.10 | 6 | 6 | 5.6894E5 | 1 | 1 | 1 | N | 19603 | AGG2 OS=Chlamydomonas reinhardtii OX=3055 GN=CHLRE\_17g738000v5 PE=2 SV=1 |
| 24 | 28 | tr\|A0A2K3CRI8\|A0A2K3CRI8\_CHLRE | 49.10 | 5 | 5 | 5.6894E5 | 1 | 1 | 1 | N | 21866 | Uncharacterized protein OS=Chlamydomonas reinhardtii OX=3055 GN=CHLRE\_17g738000v5 PE=4 SV=1 |
| 20 | 23 | tr\|A0A2K3D661\|A0A2K3D661\_CHLRE | 48.13 | 13 | 13 | 2.2434E5 | 2 | 2 | 2 | N | 25899 | PsbP domain-containing protein OS=Chlamydomonas reinhardtii OX=3055 GN=CHLRE\_12g550850v5 PE=4 SV=1 |
| 26 | 33 | tr\|A8HP90\|A8HP90\_CHLRE | 44.81 | 7 | 7 | 9.8812E3 | 1 | 1 | 1 | N | 24372 | Ribosomal protein L6 OS=Chlamydomonas reinhardtii OX=3055 GN=RPL6 PE=3 SV=1 |
| 27 | 35 | tr\|Q19VH5\|Q19VH5\_CHLRE | 44.59 | 7 | 7 | 0 | 1 | 1 | 1 | N | 21387 | NAD(P)H dehydrogenase (quinone) OS=Chlamydomonas reinhardtii OX=3055 GN=AGG3 PE=2 SV=1 |
| 27 | 34 | tr\|Q19VH4\|Q19VH4\_CHLRE | 44.59 | 7 | 7 | 0 | 1 | 1 | 1 | N | 21415 | NAD(P)H dehydrogenase (quinone) OS=Chlamydomonas reinhardtii OX=3055 GN=CHLRE\_10g456050v5 PE=2 SV=1 |
| 18 | 17 | Q08365\|RR3\_CHLRE | 44.00 | 4 | 4 | 1.148E5 | 2 | 2 | 2 | N | 81855 | 30S ribosomal protein S3, chloroplastic OS=Chlamydomonas reinhardtii OX=3055 GN=rps3 PE=1 SV=2 |
| 28 | 42 | tr\|A8JH68\|A8JH68\_CHLRE | 42.36 | 10 | 10 | 5.041E4 | 1 | 1 | 1 | N | 14875 | Plastocyanin OS=Chlamydomonas reinhardtii OX=3055 GN=PCY1 PE=3 SV=1 |
| 21 | 24 | tr\|A8I2V3\|A8I2V3\_CHLRE | 39.29 | 15 | 15 | 4.5525E4 | 2 | 2 | 2 | N | 21642 | Peroxiredoxin OS=Chlamydomonas reinhardtii OX=3055 GN=PRX2 PE=3 SV=1 |
| 22 | 36 | tr\|A8J244\|A8J244\_CHLRE | 39.26 | 10 | 10 | 6.2649E4 | 2 | 2 | 2 | N | 45749 | Isocitrate lyase OS=Chlamydomonas reinhardtii OX=3055 GN=ICL1 PE=4 SV=1 |
| 29 | 47 | tr\|A0A2K3D9K7\|A0A2K3D9K7\_CHLRE | 39.09 | 2 | 2 | 2.3634E4 | 1 | 1 | 1 | N | 77363 | Uncharacterized protein OS=Chlamydomonas reinhardtii OX=3055 GN=CHLRE\_10g427300v5 PE=3 SV=1 |
| 23 | 44 | tr\|A8HRZ9\|A8HRZ9\_CHLRE | 38.66 | 6 | 6 | 3.2553E5 | 1 | 1 | 1 | N | 15262 | Histone H2A OS=Chlamydomonas reinhardtii OX=3055 GN=CHLRE\_13g567700v5 PE=3 SV=1 |
| 23 | 51 | tr\|A8IR81\|A8IR81\_CHLRE | 38.66 | 7 | 7 | 3.2553E5 | 1 | 1 | 1 | N | 13612 | Histone H2A OS=Chlamydomonas reinhardtii OX=3055 GN=HTA20 PE=3 SV=1 |
| 23 | 50 | tr\|A8HSB5\|A8HSB5\_CHLRE | 38.66 | 7 | 7 | 3.2553E5 | 1 | 1 | 1 | N | 13628 | Histone H2A OS=Chlamydomonas reinhardtii OX=3055 GN=HTA2 PE=3 SV=1 |
| 23 | 52 | tr\|Q42680\|Q42680\_CHLRE | 38.66 | 7 | 7 | 3.2553E5 | 1 | 1 | 1 | N | 13685 | Histone H2A OS=Chlamydomonas reinhardtii OX=3055 GN=ch2a-IV PE=3 SV=1 |
| 23 | 54 | tr\|A8HWM8\|A8HWM8\_CHLRE | 38.66 | 7 | 7 | 3.2553E5 | 1 | 1 | 1 | N | 13854 | Histone H2A OS=Chlamydomonas reinhardtii OX=3055 GN=HTA11 PE=3 SV=1 |
| 23 | 53 | tr\|A8HWF6\|A8HWF6\_CHLRE | 38.66 | 7 | 7 | 3.2553E5 | 1 | 1 | 1 | N | 13841 | Histone H2A OS=Chlamydomonas reinhardtii OX=3055 GN=HTA10 PE=3 SV=1 |
| 700 | 55 | tr\|A8J363\|A8J363\_CHLRE | 37.18 | 2 | 2 | 1.6009E4 | 1 | 1 | 1 | N | 80822 | Matrix metalloproteinase-like protein OS=Chlamydomonas reinhardtii OX=3055 GN=MMP13 PE=4 SV=1 |
| total 42 proteins |  |  |  |  |  |  |  |  |  |  |  |  |
| --- | --- | --- | --- | --- | --- | --- | --- | --- | --- | --- | --- | --- |

  

PHL7

back to list

  

| Protein Coverage
| Supporting Peptides
|

Protein Coverage:

Supporting Peptides:

| Peptide | Uniq | -10lgP | Mass | Length | ppm | m/z | z | RT | Fraction | Scan | Source File | Area Sample 1 | #Feature | #Feature Sample 1 | Start | End | PTM | AScore | Found By |
| --- | --- | --- | --- | --- | --- | --- | --- | --- | --- | --- | --- | --- | --- | --- | --- | --- | --- | --- | --- |
| R.GPDPTESSIEAVR.G | Y | 97.26 | 1356.6521 | 13 | 2.4 | 679.3350 | 2 | 16.48 | 1 | 3077 | JVDM\_BotE.raw | 2.7415E6 | 1 | 1 | 8 | 20 |  |  | PEAKS DB |
| R.IASQGFVVITIDTITR.L | Y | 86.17 | 1732.9723 | 16 | 2.7 | 867.4958 | 2 | 38.48 | 1 | 8998 | JVDM\_BotE.raw | 1.4829E6 | 2 | 2 | 77 | 92 |  |  | PEAKS DB |
| R.TPTLVVGAQLDTIAPVSSHSEAFYNSLPSDLDK.A | Y | 82.00 | 3471.7410 | 33 | 2.8 | 868.9449 | 4 | 36.25 | 1 | 8484 | JVDM\_BotE.raw | 4.5148E5 | 2 | 2 | 167 | 199 |  |  | PEAKS DB |
| R.GPFAVAQTTVSR.L | Y | 77.63 | 1232.6514 | 12 | 2.3 | 617.3344 | 2 | 20.39 | 1 | 4192 | JVDM\_BotE.raw | 5.5629E6 | 1 | 1 | 21 | 32 |  |  | PEAKS DB |
| R.GPDPTESSIEAVRGPFAVAQTTVSR.L | Y | 60.11 | 2571.2927 | 25 | 2.4 | 858.1069 | 3 | 28.88 | 1 | 6627 | JVDM\_BotE.raw | 3.7583E6 | 1 | 1 | 8 | 32 |  |  | PEAKS DB |
| R.FVDDDLRYEQFLC(+57.02)PAPDDFAISEYR.S | Y | 50.54 | 3080.3860 | 25 | 4.3 | 1027.8070 | 3 | 36.55 | 1 | 8557 | JVDM\_BotE.raw | 1.1317E5 | 1 | 1 | 230 | 254 | Carbamidomethylation | C13:Carbamidomethylation:1000.00 | PEAKS DB |
| R.TPTLVVGAQLDTIAPVSSHSEAFYN(+.98)SLPSDLDK.A | Y | 48.69 | 3472.7249 | 33 | 2.4 | 1158.5850 | 3 | 35.50 | 1 | 8314 | JVDM\_BotE.raw | 0 | 0 | 0 | 167 | 199 | Deamidation (NQ) | N25:Deamidation (NQ):0.00 | PEAKS PTM |
| R.QLQAALDHLR.T | Y | 45.78 | 1163.6411 | 10 | 2.8 | 582.8295 | 2 | 19.39 | 1 | 3892 | JVDM\_BotE.raw | 2.7016E6 | 2 | 2 | 102 | 111 |  |  | PEAKS DB |
| K.YSIAWLK.R | Y | 27.33 | 879.4854 | 7 | 1.6 | 440.7507 | 2 | 29.05 | 1 | 6640 | JVDM\_BotE.raw | 8.7419E5 | 1 | 1 | 222 | 228 |  |  | PEAKS DB |
| R.YEQFLC(+57.02)PAPDDFAISEYR.S | Y | 20.38 | 2219.9834 | 18 | 3.1 | 1111.0024 | 2 | 33.56 | 1 | 7863 | JVDM\_BotE.raw | 7.955E4 | 1 | 1 | 237 | 254 | Carbamidomethylation | C6:Carbamidomethylation:1000.00 | PEAKS DB |
| total 10 peptides |  |  |  |  |  |  |  |  |  |  |  |  |  |  |  |  |  |  |  |
| --- | --- | --- | --- | --- | --- | --- | --- | --- | --- | --- | --- | --- | --- | --- | --- | --- | --- | --- | --- |

tr|A0A2K3D8I0|A0A2K3D8I0\_CHLRE

back to list

  

| Protein Coverage
| Supporting Peptides
|

Protein Coverage:

Supporting Peptides:

| Peptide | Uniq | -10lgP | Mass | Length | ppm | m/z | z | RT | Fraction | Scan | Source File | Area Sample 1 | #Feature | #Feature Sample 1 | Start | End | PTM | AScore | Found By |
| --- | --- | --- | --- | --- | --- | --- | --- | --- | --- | --- | --- | --- | --- | --- | --- | --- | --- | --- | --- |
| K.AAAAASLGQAQDAAAQAQNTAAAAVGNAQTAAGDAAGK.A | Y | 80.91 | 3323.6089 | 38 | 0.0 | 1108.8770 | 3 | 29.70 | 1 | 6835 | JVDM\_BotE.raw | 0 | 0 | 0 | 792 | 829 |  |  | PEAKS DB |
| K.DAAAAAQQQAAAAADAAK.G | Y | 78.99 | 1612.7804 | 18 | 1.1 | 807.3984 | 2 | 17.40 | 1 | 3317 | JVDM\_BotE.raw | 0 | 0 | 0 | 865 | 882 |  |  | PEAKS DB |
| R.GQAEAAAAQAQNTAAAAVGNVQTAAADAAAK.A | Y | 77.81 | 2781.3640 | 31 | 1.2 | 928.1298 | 3 | 29.67 | 1 | 6825 | JVDM\_BotE.raw | 1.4042E5 | 1 | 1 | 701 | 731 |  |  | PEAKS DB |
| K.GAAASAVGQATAQATAAVDQAK.A | Y | 75.23 | 1956.9865 | 22 | 1.5 | 653.3371 | 3 | 24.46 | 1 | 5439 | JVDM\_BotE.raw | 2.8473E5 | 1 | 1 | 883 | 904 |  |  | PEAKS DB |
| K.AASAAAEQAQAAAVGAQQAAVEK.A | Y | 74.68 | 2111.0605 | 23 | 2.3 | 704.6957 | 3 | 19.45 | 1 | 3936 | JVDM\_BotE.raw | 8.986E4 | 1 | 1 | 575 | 597 |  |  | PEAKS DB |
| K.AAAAQATAAAAGAQEAASAAAGQATATATDAMTK.A | Y | 60.03 | 2961.4097 | 34 | 2.8 | 988.1466 | 3 | 30.04 | 1 | 6932 | JVDM\_BotE.raw | 7.1196E4 | 1 | 1 | 756 | 789 |  |  | PEAKS DB |
| K.AAAAASLGQAQDAAAQ(+.98)AQNTAAAAVGNAQTAAGDAAGK.A | Y | 47.67 | 3324.5930 | 38 | 6.3 | 1109.2119 | 3 | 29.81 | 1 | 6867 | JVDM\_BotE.raw | 0 | 0 | 0 | 792 | 829 | Deamidation (NQ) | Q16:Deamidation (NQ):4.64 | PEAKS PTM |
| K.AAADGAVANAQGAVTSFATK.A | Y | 46.13 | 1819.9064 | 20 | -1.5 | 910.9591 | 2 | 24.16 | 1 | 5324 | JVDM\_BotE.raw | 6.2277E4 | 1 | 1 | 661 | 680 |  |  | PEAKS DB |
| K.AEDTAAAAQAQAAAAAESAR.G | Y | 44.31 | 1843.8660 | 20 | -0.1 | 615.6292 | 3 | 18.71 | 1 | 3725 | JVDM\_BotE.raw | 1.0125E5 | 1 | 1 | 681 | 700 |  |  | PEAKS DB |
| K.AAAAQATAAAAGAQ(+.98)EAASAAAGQ(+.98)ATATATDAMTK(+14.02).A | Y | 39.38 | 2977.3933 | 34 | 5.1 | 993.4767 | 3 | 25.89 | 1 | 5833 | JVDM\_BotE.raw | 0 | 0 | 0 | 756 | 789 | Deamidation (NQ); Methylation(KR) | Q14:Deamidation (NQ):50.04;Q23:Deamidation (NQ):30.07;K34:Methylation(KR):1000.00 | PEAKS PTM |
| K.AAEAQ(+.98)KAASAAAEQAQAAAVGAQQAAVEK(+42.01)AAEAQ(+.98)K.A | Y | 34.70 | 3351.6541 | 35 | -3.3 | 838.9180 | 4 | 24.05 | 1 | 5366 | JVDM\_BotE.raw | 6.1886E6 | 1 | 1 | 569 | 603 | Deamidation (NQ); Acetylation (K) | Q5:Deamidation (NQ):19.64;K29:Acetylation (K):13.13;Q34:Deamidation (NQ):3.41 | PEAKS PTM |
| K.AQETAAAAAGQAQAAAADAAAK.A | Y | 32.93 | 1926.9395 | 22 | 1.7 | 643.3215 | 3 | 19.41 | 1 | 3905 | JVDM\_BotE.raw | 1.5324E5 | 1 | 1 | 830 | 851 |  |  | PEAKS DB |
| K.AAT(+79.97)VAADAQ(+.98)AAAAKAQDAAAGAAAAAT(+79.97)EKAAEAQK.A | Y | 30.62 | 3285.4915 | 35 | 0.3 | 1096.1714 | 3 | 33.47 | 1 | 7815 | JVDM\_BotE.raw | 5.1671E5 | 1 | 1 | 464 | 498 | Phosphorylation (STY); Deamidation (NQ) | T3:Phosphorylation (STY):1000.00;Q9:Deamidation (NQ):0.00;T27:Phosphorylation (STY):1000.00 | PEAKS PTM |
| K.AGAAQAAATAAASQ(+.98)VAEAATGAAASAQSA.A | Y | 29.21 | 2416.1465 | 29 | -1.0 | 605.0433 | 4 | 27.55 | 1 | 6254 | JVDM\_BotE.raw | 0 | 0 | 0 | 370 | 398 | Deamidation (NQ) | Q14:Deamidation (NQ):3.58 | PEAKS PTM |
| K.AATVAADAQAAAAK(+14.02)AQDAAAGAAAAATEK.A | Y | 26.65 | 2540.2830 | 29 | 0.0 | 847.7682 | 3 | 26.91 | 1 | 6227 | JVDM\_BotE.raw | 6.2014E5 | 1 | 1 | 464 | 492 | Methylation(KR) | K14:Methylation(KR):6.28 | PEAKS PTM |
| R.DAAAGAAAQAQ(+.98)TAAAGLMGSLN(+.98)AAAGK.A | Y | 17.89 | 2330.1172 | 27 | -4.6 | 777.7094 | 3 | 35.27 | 1 | 8197 | JVDM\_BotE.raw | 2.704E6 | 1 | 1 | 923 | 949 | Deamidation (NQ) | Q11:Deamidation (NQ):0.00;N22:Deamidation (NQ):3.30 | PEAKS PTM |
| E.AATGAAASAQS(+79.97)AAAAAVAAAGDK.A | Y | 16.79 | 1951.9000 | 23 | 3.4 | 976.9606 | 2 | 27.34 | 1 | 6164 | JVDM\_BotE.raw | 2.7313E5 | 1 | 1 | 387 | 409 | Phosphorylation (STY) | S11:Phosphorylation (STY):5.08 | PEAKS PTM |
| K.AAAQAQEAAAGAATQ(+.98)AK.A | Y | 16.54 | 1528.7480 | 17 | -1.2 | 765.3804 | 2 | 22.04 | 1 | 4714 | JVDM\_BotE.raw | 9.5974E4 | 1 | 1 | 644 | 660 | Deamidation (NQ) | Q15:Deamidation (NQ):6.45 | PEAKS PTM |
| total 18 peptides |  |  |  |  |  |  |  |  |  |  |  |  |  |  |  |  |  |  |  |
| --- | --- | --- | --- | --- | --- | --- | --- | --- | --- | --- | --- | --- | --- | --- | --- | --- | --- | --- | --- |

tr|Q7X7A7|Q7X7A7\_CHLRE

back to list

  

| Protein Coverage
| Supporting Peptides
|

Protein Coverage:

Supporting Peptides:

| Peptide | Uniq | -10lgP | Mass | Length | ppm | m/z | z | RT | Fraction | Scan | Source File | Area Sample 1 | #Feature | #Feature Sample 1 | Start | End | PTM | AScore | Found By |
| --- | --- | --- | --- | --- | --- | --- | --- | --- | --- | --- | --- | --- | --- | --- | --- | --- | --- | --- | --- |
| K.VAVLANEQELSVEER.N | Y | 93.89 | 1684.8632 | 15 | 1.6 | 843.4402 | 2 | 23.17 | 1 | 5030 | JVDM\_BotE.raw | 6.8004E5 | 1 | 1 | 33 | 47 |  |  | PEAKS DB |
| K.AAAAELTLAAYK.V | Y | 70.65 | 1191.6499 | 12 | 1.9 | 596.8334 | 2 | 23.82 | 1 | 5213 | JVDM\_BotE.raw | 8.2982E5 | 1 | 1 | 147 | 158 |  |  | PEAKS DB |
| R.ILQSIEQSEQAK.G | Y | 63.08 | 1372.7197 | 12 | 2.2 | 687.3687 | 2 | 15.42 | 1 | 2819 | JVDM\_BotE.raw | 4.4047E5 | 1 | 1 | 67 | 78 |  |  | PEAKS DB |
| K.VAQEKADELPTTNPIR.L | Y | 53.58 | 1780.9319 | 16 | 2.2 | 594.6525 | 3 | 17.75 | 1 | 3412 | JVDM\_BotE.raw | 5.1889E5 | 2 | 2 | 159 | 174 |  |  | PEAKS DB |
| K.ADELPTTNPIR.L | Y | 47.10 | 1225.6302 | 11 | 1.2 | 613.8231 | 2 | 19.20 | 1 | 3830 | JVDM\_BotE.raw | 3.0501E5 | 1 | 1 | 164 | 174 |  |  | PEAKS DB |
| K.YLVPSASTTEAAVFYLK.M | Y | 19.28 | 1858.9716 | 17 | 1.1 | 930.4941 | 2 | 34.72 | 1 | 8142 | JVDM\_BotE.raw | 0 | 0 | 0 | 112 | 128 |  |  | PEAKS DB |
| K.TDEAK(+42.01)AAAAELT(+79.97)LAAYK.V | Y | 17.70 | 1857.8761 | 17 | 2.6 | 929.9478 | 2 | 30.69 | 1 | 7100 | JVDM\_BotE.raw | 2.6137E5 | 1 | 1 | 142 | 158 | Acetylation (K); Phosphorylation (STY) | K5:Acetylation (K):8.68;T12:Phosphorylation (STY):0.00 | PEAKS PTM |
| R.ELALYSAKIAEQSERYQDMVEEMK(+28.03).K | Y | 17.46 | 2888.3936 | 24 | -1.9 | 963.8033 | 3 | 36.91 | 1 | 8641 | JVDM\_BotE.raw | 1.2065E4 | 1 | 1 | 8 | 31 | Dimethylation(KR) | K24:Dimethylation(KR):13.91 | PEAKS PTM |
| total 8 peptides |  |  |  |  |  |  |  |  |  |  |  |  |  |  |  |  |  |  |  |
| --- | --- | --- | --- | --- | --- | --- | --- | --- | --- | --- | --- | --- | --- | --- | --- | --- | --- | --- | --- |

tr|A0A2K3D6N5|A0A2K3D6N5\_CHLRE

back to list

  

| Protein Coverage
| Supporting Peptides
|

Protein Coverage:

Supporting Peptides:

| Peptide | Uniq | -10lgP | Mass | Length | ppm | m/z | z | RT | Fraction | Scan | Source File | Area Sample 1 | #Feature | #Feature Sample 1 | Start | End | PTM | AScore | Found By |
| --- | --- | --- | --- | --- | --- | --- | --- | --- | --- | --- | --- | --- | --- | --- | --- | --- | --- | --- | --- |
| K.AAGDNAQAAAVPSVTEQASEFLAGSWPK.T | Y | 77.91 | 2772.3354 | 28 | 1.9 | 925.1208 | 3 | 38.38 | 1 | 8982 | JVDM\_BotE.raw | 6.3426E4 | 1 | 1 | 132 | 159 |  |  | PEAKS DB |
| K.VGVAGAVVVLGYNWK.A | Y | 61.34 | 1530.8558 | 15 | 3.5 | 766.4379 | 2 | 35.01 | 1 | 8195 | JVDM\_BotE.raw | 4.7808E4 | 1 | 1 | 204 | 218 |  |  | PEAKS DB |
| R.VVIALDSSSHSGADFAAK.A | Y | 48.60 | 1773.8896 | 18 | 0.0 | 592.3038 | 3 | 22.39 | 1 | 4780 | JVDM\_BotE.raw | 1.8993E5 | 1 | 1 | 102 | 119 |  |  | PEAKS DB |
| K.WSYPAAGVVSDATLGR(+14.02).V | Y | 37.18 | 1662.8365 | 16 | 6.4 | 832.4309 | 2 | 27.68 | 1 | 6299 | JVDM\_BotE.raw | 5.8157E4 | 1 | 1 | 670 | 685 | Methylation(KR) | R16:Methylation(KR):1000.00 | PEAKS PTM |
| R.ITSASTVC(+57.02)SHEDSSAVVIPLYTYEDGEDVR.V | Y | 35.52 | 3299.5139 | 30 | 5.1 | 1100.8508 | 3 | 30.14 | 1 | 6947 | JVDM\_BotE.raw | 0 | 0 | 0 | 166 | 195 | Carbamidomethylation | C8:Carbamidomethylation:1000.00 | PEAKS DB |
| K.AGPQEQWGALLNK.I | Y | 24.33 | 1410.7256 | 13 | 2.5 | 706.3718 | 2 | 27.17 | 1 | 6163 | JVDM\_BotE.raw | 1.5348E5 | 1 | 1 | 219 | 231 |  |  | PEAKS DB |
| total 6 peptides |  |  |  |  |  |  |  |  |  |  |  |  |  |  |  |  |  |  |  |
| --- | --- | --- | --- | --- | --- | --- | --- | --- | --- | --- | --- | --- | --- | --- | --- | --- | --- | --- | --- |

tr|A0A2K3DP12|A0A2K3DP12\_CHLRE

back to list

  

| Protein Coverage
| Supporting Peptides
|

Protein Coverage:

Supporting Peptides:

| Peptide | Uniq | -10lgP | Mass | Length | ppm | m/z | z | RT | Fraction | Scan | Source File | Area Sample 1 | #Feature | #Feature Sample 1 | Start | End | PTM | AScore | Found By |
| --- | --- | --- | --- | --- | --- | --- | --- | --- | --- | --- | --- | --- | --- | --- | --- | --- | --- | --- | --- |
| R.SVGAAGGQEDIVNLVPTAAAATSTGK.S | Y | 97.51 | 2384.2183 | 26 | 3.2 | 795.7492 | 3 | 30.00 | 1 | 6916 | JVDM\_BotE.raw | 4.5271E4 | 1 | 1 | 200 | 225 |  |  | PEAKS DB |
| K.TASTASLQLC(+57.02)R.A | Y | 44.71 | 1206.6027 | 11 | 1.3 | 604.3094 | 2 | 16.77 | 1 | 3136 | JVDM\_BotE.raw | 5.0051E4 | 1 | 1 | 624 | 634 | Carbamidomethylation | C10:Carbamidomethylation:1000.00 | PEAKS DB |
| R.SVGAAGGQEDIVNLVPTAAAATSTGK(+14.02)SNSAK(+28.03).D | Y | 33.17 | 2913.5044 | 31 | 5.0 | 972.1802 | 3 | 34.56 | 1 | 8101 | JVDM\_BotE.raw | 0 | 0 | 0 | 200 | 230 | Methylation(KR); Dimethylation(KR) | K26:Methylation(KR):0.00;K31:Dimethylation(KR):0.00 | PEAKS PTM |
| total 3 peptides |  |  |  |  |  |  |  |  |  |  |  |  |  |  |  |  |  |  |  |
| --- | --- | --- | --- | --- | --- | --- | --- | --- | --- | --- | --- | --- | --- | --- | --- | --- | --- | --- | --- |

tr|A8IKW6|A8IKW6\_CHLRE

back to list

  

| Protein Coverage
| Supporting Peptides
|

Protein Coverage:

Supporting Peptides:

| Peptide | Uniq | -10lgP | Mass | Length | ppm | m/z | z | RT | Fraction | Scan | Source File | Area Sample 1 | #Feature | #Feature Sample 1 | Start | End | PTM | AScore | Found By |
| --- | --- | --- | --- | --- | --- | --- | --- | --- | --- | --- | --- | --- | --- | --- | --- | --- | --- | --- | --- |
| K.VIDAGANALVAGSAVFK.A | Y | 87.38 | 1601.8777 | 17 | 2.3 | 801.9480 | 2 | 30.60 | 1 | 7081 | JVDM\_BotE.raw | 1.1643E5 | 1 | 1 | 227 | 243 |  |  | PEAKS DB |
| K.SDIIVSPSILSADFSR.L | Y | 50.98 | 1705.8887 | 16 | 2.6 | 853.9538 | 2 | 35.55 | 1 | 8328 | JVDM\_BotE.raw | 2.2334E4 | 1 | 1 | 40 | 55 |  |  | PEAKS DB |
| total 2 peptides |  |  |  |  |  |  |  |  |  |  |  |  |  |  |  |  |  |  |  |
| --- | --- | --- | --- | --- | --- | --- | --- | --- | --- | --- | --- | --- | --- | --- | --- | --- | --- | --- | --- |

tr|A0A2K3CQ54|A0A2K3CQ54\_CHLRE

back to list

  

| Protein Coverage
| Supporting Peptides
|

Protein Coverage:

Supporting Peptides:

| Peptide | Uniq | -10lgP | Mass | Length | ppm | m/z | z | RT | Fraction | Scan | Source File | Area Sample 1 | #Feature | #Feature Sample 1 | Start | End | PTM | AScore | Found By |
| --- | --- | --- | --- | --- | --- | --- | --- | --- | --- | --- | --- | --- | --- | --- | --- | --- | --- | --- | --- |
| R.ATVAAGEEALTIR.N | Y | 69.67 | 1300.6986 | 13 | 2.2 | 651.3580 | 2 | 21.62 | 1 | 4543 | JVDM\_BotE.raw | 3.192E6 | 1 | 1 | 141 | 153 |  |  | PEAKS DB |
| K.VEYTELQILC(+57.02)PQTIDSVTGYPMDDPR.C | Y | 67.82 | 3039.4204 | 26 | 1.1 | 1014.1486 | 3 | 37.54 | 1 | 8770 | JVDM\_BotE.raw | 1.924E6 | 1 | 1 | 110 | 135 | Carbamidomethylation | C10:Carbamidomethylation:1000.00 | PEAKS DB |
| Q.TIDSVTGYPMDDPR.C | Y | 31.92 | 1565.7031 | 14 | 1.8 | 783.8602 | 2 | 21.41 | 1 | 4497 | JVDM\_BotE.raw | 6.6961E4 | 1 | 1 | 122 | 135 |  |  | PEAKS DB |
| K.LSGWSAPATLT(+79.97)PEK.V | Y | 27.66 | 1536.7225 | 14 | -7.3 | 769.3629 | 2 | 33.61 | 1 | 7885 | JVDM\_BotE.raw | 6.9909E4 | 1 | 1 | 246 | 259 | Phosphorylation (STY) | T11:Phosphorylation (STY):0.00 | PEAKS PTM |
| total 4 peptides |  |  |  |  |  |  |  |  |  |  |  |  |  |  |  |  |  |  |  |
| --- | --- | --- | --- | --- | --- | --- | --- | --- | --- | --- | --- | --- | --- | --- | --- | --- | --- | --- | --- |

tr|B7U1J0|B7U1J0\_CHLRE

back to list

  

| Protein Coverage
| Supporting Peptides
|

Protein Coverage:

Supporting Peptides:

| Peptide | Uniq | -10lgP | Mass | Length | ppm | m/z | z | RT | Fraction | Scan | Source File | Area Sample 1 | #Feature | #Feature Sample 1 | Start | End | PTM | AScore | Found By |
| --- | --- | --- | --- | --- | --- | --- | --- | --- | --- | --- | --- | --- | --- | --- | --- | --- | --- | --- | --- |
| K.TAIAVDTILNQK.G | Y | 77.28 | 1285.7241 | 12 | 2.5 | 643.8710 | 2 | 26.92 | 1 | 6082 | JVDM\_BotE.raw | 1.5927E5 | 1 | 1 | 177 | 188 |  |  | PEAKS DB |
| K.ASSVAQVLNTLK.E | Y | 48.38 | 1229.6979 | 12 | 1.8 | 615.8573 | 2 | 29.25 | 1 | 6724 | JVDM\_BotE.raw | 1.535E5 | 1 | 1 | 203 | 214 |  |  | PEAKS DB |
| R.STLTFTPEAEGLVK.Q | Y | 29.89 | 1491.7820 | 14 | 2.7 | 746.9003 | 2 | 28.40 | 1 | 6455 | JVDM\_BotE.raw | 0 | 0 | 0 | 478 | 491 |  |  | PEAKS DB |
| total 3 peptides |  |  |  |  |  |  |  |  |  |  |  |  |  |  |  |  |  |  |  |
| --- | --- | --- | --- | --- | --- | --- | --- | --- | --- | --- | --- | --- | --- | --- | --- | --- | --- | --- | --- |

P26526|ATPA\_CHLRE

back to list

  

| Protein Coverage
| Supporting Peptides
|

Protein Coverage:

Supporting Peptides:

| Peptide | Uniq | -10lgP | Mass | Length | ppm | m/z | z | RT | Fraction | Scan | Source File | Area Sample 1 | #Feature | #Feature Sample 1 | Start | End | PTM | AScore | Found By |
| --- | --- | --- | --- | --- | --- | --- | --- | --- | --- | --- | --- | --- | --- | --- | --- | --- | --- | --- | --- |
| K.TAIAVDTILNQK.G | Y | 77.28 | 1285.7241 | 12 | 2.5 | 643.8710 | 2 | 26.92 | 1 | 6082 | JVDM\_BotE.raw | 1.5927E5 | 1 | 1 | 177 | 188 |  |  | PEAKS DB |
| K.ASSVAQVLNTLK.E | Y | 48.38 | 1229.6979 | 12 | 1.8 | 615.8573 | 2 | 29.25 | 1 | 6724 | JVDM\_BotE.raw | 1.535E5 | 1 | 1 | 203 | 214 |  |  | PEAKS DB |
| R.STLTFTPEAEGLVK.Q | Y | 29.89 | 1491.7820 | 14 | 2.7 | 746.9003 | 2 | 28.40 | 1 | 6455 | JVDM\_BotE.raw | 0 | 0 | 0 | 478 | 491 |  |  | PEAKS DB |
| total 3 peptides |  |  |  |  |  |  |  |  |  |  |  |  |  |  |  |  |  |  |  |
| --- | --- | --- | --- | --- | --- | --- | --- | --- | --- | --- | --- | --- | --- | --- | --- | --- | --- | --- | --- |

tr|Q9FE86|Q9FE86\_CHLRE

back to list

  

| Protein Coverage
| Supporting Peptides
|

Protein Coverage:

Supporting Peptides:

| Peptide | Uniq | -10lgP | Mass | Length | ppm | m/z | z | RT | Fraction | Scan | Source File | Area Sample 1 | #Feature | #Feature Sample 1 | Start | End | PTM | AScore | Found By |
| --- | --- | --- | --- | --- | --- | --- | --- | --- | --- | --- | --- | --- | --- | --- | --- | --- | --- | --- | --- |
| K.AQAVFDQEFQEITLSK.Y | Y | 73.53 | 1852.9207 | 16 | 1.6 | 927.4691 | 2 | 31.91 | 1 | 7407 | JVDM\_BotE.raw | 2.0286E5 | 1 | 1 | 55 | 70 |  |  | PEAKS DB |
| K.EGGLGDLAYPLVADLKK.E | Y | 52.12 | 1757.9563 | 17 | 1.8 | 586.9938 | 3 | 34.62 | 1 | 8124 | JVDM\_BotE.raw | 1.6547E5 | 1 | 1 | 131 | 147 |  |  | PEAKS DB |
| A.SHAEKPLVGSVAPDFK.A | Y | 33.48 | 1680.8834 | 16 | 1.5 | 561.3026 | 3 | 20.12 | 1 | 4130 | JVDM\_BotE.raw | 4.6718E4 | 1 | 1 | 39 | 54 |  |  | PEAKS DB |
| total 3 peptides |  |  |  |  |  |  |  |  |  |  |  |  |  |  |  |  |  |  |  |
| --- | --- | --- | --- | --- | --- | --- | --- | --- | --- | --- | --- | --- | --- | --- | --- | --- | --- | --- | --- |

tr|A8J9H8|A8J9H8\_CHLRE

back to list

  

| Protein Coverage
| Supporting Peptides
|

Protein Coverage:

Supporting Peptides:

| Peptide | Uniq | -10lgP | Mass | Length | ppm | m/z | z | RT | Fraction | Scan | Source File | Area Sample 1 | #Feature | #Feature Sample 1 | Start | End | PTM | AScore | Found By |
| --- | --- | --- | --- | --- | --- | --- | --- | --- | --- | --- | --- | --- | --- | --- | --- | --- | --- | --- | --- |
| K.MIGATNPLASEPGTIR.G | Y | 75.42 | 1626.8400 | 16 | 3.8 | 814.4304 | 2 | 25.35 | 1 | 5689 | JVDM\_BotE.raw | 6.1475E4 | 1 | 1 | 89 | 104 |  |  | PEAKS DB |
| R.GDFAIEVGR.N | Y | 47.52 | 962.4821 | 9 | 0.8 | 482.2487 | 2 | 22.17 | 1 | 4705 | JVDM\_BotE.raw | 4.0966E5 | 1 | 1 | 105 | 113 |  |  | PEAKS DB |
| R.KMIGATNPLASEPGTIR.G | Y | 26.66 | 1754.9348 | 17 | 2.8 | 585.9872 | 3 | 22.49 | 1 | 4796 | JVDM\_BotE.raw | 8.461E4 | 1 | 1 | 88 | 104 |  |  | PEAKS DB |
| total 3 peptides |  |  |  |  |  |  |  |  |  |  |  |  |  |  |  |  |  |  |  |
| --- | --- | --- | --- | --- | --- | --- | --- | --- | --- | --- | --- | --- | --- | --- | --- | --- | --- | --- | --- |

tr|A0A2K3DJF1|A0A2K3DJF1\_CHLRE

back to list

  

| Protein Coverage
| Supporting Peptides
|

Protein Coverage:

Supporting Peptides:

| Peptide | Uniq | -10lgP | Mass | Length | ppm | m/z | z | RT | Fraction | Scan | Source File | Area Sample 1 | #Feature | #Feature Sample 1 | Start | End | PTM | AScore | Found By |
| --- | --- | --- | --- | --- | --- | --- | --- | --- | --- | --- | --- | --- | --- | --- | --- | --- | --- | --- | --- |
| K.MIGATNPLASEPGTIR.G | Y | 75.42 | 1626.8400 | 16 | 3.8 | 814.4304 | 2 | 25.35 | 1 | 5689 | JVDM\_BotE.raw | 6.1475E4 | 1 | 1 | 101 | 116 |  |  | PEAKS DB |
| R.GDFAIEVGR.N | Y | 47.52 | 962.4821 | 9 | 0.8 | 482.2487 | 2 | 22.17 | 1 | 4705 | JVDM\_BotE.raw | 4.0966E5 | 1 | 1 | 117 | 125 |  |  | PEAKS DB |
| R.KMIGATNPLASEPGTIR.G | Y | 26.66 | 1754.9348 | 17 | 2.8 | 585.9872 | 3 | 22.49 | 1 | 4796 | JVDM\_BotE.raw | 8.461E4 | 1 | 1 | 100 | 116 |  |  | PEAKS DB |
| total 3 peptides |  |  |  |  |  |  |  |  |  |  |  |  |  |  |  |  |  |  |  |
| --- | --- | --- | --- | --- | --- | --- | --- | --- | --- | --- | --- | --- | --- | --- | --- | --- | --- | --- | --- |

tr|A0A2K3CWF7|A0A2K3CWF7\_CHLRE

back to list

  

| Protein Coverage
| Supporting Peptides
|

Protein Coverage:

Supporting Peptides:

| Peptide | Uniq | -10lgP | Mass | Length | ppm | m/z | z | RT | Fraction | Scan | Source File | Area Sample 1 | #Feature | #Feature Sample 1 | Start | End | PTM | AScore | Found By |
| --- | --- | --- | --- | --- | --- | --- | --- | --- | --- | --- | --- | --- | --- | --- | --- | --- | --- | --- | --- |
| K.SPASDTYIIFGEAK.I | Y | 75.16 | 1497.7351 | 14 | 1.0 | 749.8756 | 2 | 28.74 | 1 | 6556 | JVDM\_BotE.raw | 0 | 0 | 0 | 91 | 104 |  |  | PEAKS DB |
| K.IEDLSAQTQAAAAEQFK.M | Y | 27.21 | 1819.8951 | 17 | 0.7 | 910.9554 | 2 | 22.93 | 1 | 4936 | JVDM\_BotE.raw | 0 | 0 | 0 | 105 | 121 |  |  | PEAKS DB |
| K.NILFVIQSPDVFK.S | Y | 17.73 | 1518.8446 | 13 | 2.6 | 760.4316 | 2 | 39.18 | 1 | 9155 | JVDM\_BotE.raw | 3.0111E4 | 1 | 1 | 78 | 90 |  |  | PEAKS DB |
| total 3 peptides |  |  |  |  |  |  |  |  |  |  |  |  |  |  |  |  |  |  |  |
| --- | --- | --- | --- | --- | --- | --- | --- | --- | --- | --- | --- | --- | --- | --- | --- | --- | --- | --- | --- |

tr|A0A2K3E0C1|A0A2K3E0C1\_CHLRE

back to list

  

| Protein Coverage
| Supporting Peptides
|

Protein Coverage:

Supporting Peptides:

| Peptide | Uniq | -10lgP | Mass | Length | ppm | m/z | z | RT | Fraction | Scan | Source File | Area Sample 1 | #Feature | #Feature Sample 1 | Start | End | PTM | AScore | Found By |
| --- | --- | --- | --- | --- | --- | --- | --- | --- | --- | --- | --- | --- | --- | --- | --- | --- | --- | --- | --- |
| K.SSQGVPLLAEQASSFFSSK.A | N | 50.34 | 1968.9792 | 19 | 2.2 | 985.4991 | 2 | 37.02 | 1 | 8653 | JVDM\_BotE.raw | 1.1276E5 | 1 | 1 | 144 | 162 |  |  | PEAKS DB |
| K.QLVQAGFDLK.V | Y | 48.43 | 1117.6132 | 10 | 1.9 | 559.8149 | 2 | 25.03 | 1 | 5612 | JVDM\_BotE.raw | 1.0514E5 | 1 | 1 | 54 | 63 |  |  | PEAKS DB |
| K.AQDAWPASLEDAK.T | N | 28.56 | 1400.6571 | 13 | 1.4 | 701.3368 | 2 | 23.56 | 1 | 5155 | JVDM\_BotE.raw | 9.8139E4 | 1 | 1 | 163 | 175 |  |  | PEAKS DB |
| R.HEDVEAFTIPLYTAAGDEMK.V | N | 24.40 | 2236.0356 | 20 | 0.2 | 746.3527 | 3 | 34.04 | 1 | 7975 | JVDM\_BotE.raw | 0 | 0 | 0 | 185 | 204 |  |  | PEAKS DB |
| I.KVPAAGS(+79.97)AANIQ(+.98)PKR.L | Y | 22.81 | 1587.8134 | 15 | -5.3 | 794.9097 | 2 | 37.79 | 1 | 8852 | JVDM\_BotE.raw | 0 | 0 | 0 | 746 | 760 | Phosphorylation (STY); Deamidation (NQ) | S7:Phosphorylation (STY):1000.00;Q12:Deamidation (NQ):0.00 | PEAKS PTM |
| K.QLVQ(+.98)AGFDLKVTADGQ(+.98)PR.L | Y | 22.69 | 1943.9952 | 18 | 3.4 | 973.0082 | 2 | 36.18 | 1 | 8464 | JVDM\_BotE.raw | 0 | 0 | 0 | 54 | 71 | Deamidation (NQ) | Q4:Deamidation (NQ):0.00;Q16:Deamidation (NQ):17.01 | PEAKS PTM |
| total 6 peptides |  |  |  |  |  |  |  |  |  |  |  |  |  |  |  |  |  |  |  |
| --- | --- | --- | --- | --- | --- | --- | --- | --- | --- | --- | --- | --- | --- | --- | --- | --- | --- | --- | --- |

tr|A0A2K3E094|A0A2K3E094\_CHLRE

back to list

  

| Protein Coverage
| Supporting Peptides
|

Protein Coverage:

Supporting Peptides:

| Peptide | Uniq | -10lgP | Mass | Length | ppm | m/z | z | RT | Fraction | Scan | Source File | Area Sample 1 | #Feature | #Feature Sample 1 | Start | End | PTM | AScore | Found By |
| --- | --- | --- | --- | --- | --- | --- | --- | --- | --- | --- | --- | --- | --- | --- | --- | --- | --- | --- | --- |
| K.SSQGVPLLAEQASSFFSSK.A | N | 50.34 | 1968.9792 | 19 | 2.2 | 985.4991 | 2 | 37.02 | 1 | 8653 | JVDM\_BotE.raw | 1.1276E5 | 1 | 1 | 144 | 162 |  |  | PEAKS DB |
| K.QLVQAGFDLK.V | Y | 48.43 | 1117.6132 | 10 | 1.9 | 559.8149 | 2 | 25.03 | 1 | 5612 | JVDM\_BotE.raw | 1.0514E5 | 1 | 1 | 54 | 63 |  |  | PEAKS DB |
| K.AQDAWPASLEDAK.T | N | 28.56 | 1400.6571 | 13 | 1.4 | 701.3368 | 2 | 23.56 | 1 | 5155 | JVDM\_BotE.raw | 9.8139E4 | 1 | 1 | 163 | 175 |  |  | PEAKS DB |
| total 3 peptides |  |  |  |  |  |  |  |  |  |  |  |  |  |  |  |  |  |  |  |
| --- | --- | --- | --- | --- | --- | --- | --- | --- | --- | --- | --- | --- | --- | --- | --- | --- | --- | --- | --- |

tr|A8IT01|A8IT01\_CHLRE

back to list

  

| Protein Coverage
| Supporting Peptides
|

Protein Coverage:

Supporting Peptides:

| Peptide | Uniq | -10lgP | Mass | Length | ppm | m/z | z | RT | Fraction | Scan | Source File | Area Sample 1 | #Feature | #Feature Sample 1 | Start | End | PTM | AScore | Found By |
| --- | --- | --- | --- | --- | --- | --- | --- | --- | --- | --- | --- | --- | --- | --- | --- | --- | --- | --- | --- |
| R.EGTFSNLPAGTTIK.L | Y | 74.37 | 1434.7355 | 14 | 1.8 | 718.3763 | 2 | 22.60 | 1 | 4828 | JVDM\_BotE.raw | 1.9599E5 | 1 | 1 | 228 | 241 |  |  | PEAKS DB |
| K.QSPINVPQYQVLDGK.G | Y | 34.66 | 1684.8784 | 15 | 1.8 | 843.4480 | 2 | 26.20 | 1 | 5915 | JVDM\_BotE.raw | 0 | 0 | 0 | 67 | 81 |  |  | PEAKS DB |
| total 2 peptides |  |  |  |  |  |  |  |  |  |  |  |  |  |  |  |  |  |  |  |
| --- | --- | --- | --- | --- | --- | --- | --- | --- | --- | --- | --- | --- | --- | --- | --- | --- | --- | --- | --- |

tr|A8JEV1|A8JEV1\_CHLRE

back to list

  

| Protein Coverage
| Supporting Peptides
|

Protein Coverage:

Supporting Peptides:

| Peptide | Uniq | -10lgP | Mass | Length | ppm | m/z | z | RT | Fraction | Scan | Source File | Area Sample 1 | #Feature | #Feature Sample 1 | Start | End | PTM | AScore | Found By |
| --- | --- | --- | --- | --- | --- | --- | --- | --- | --- | --- | --- | --- | --- | --- | --- | --- | --- | --- | --- |
| R.IDADLDVFIQK.S | Y | 50.94 | 1275.6710 | 11 | 2.9 | 638.8446 | 2 | 30.91 | 1 | 7190 | JVDM\_BotE.raw | 2.7666E5 | 1 | 1 | 108 | 118 |  |  | PEAKS DB |
| R.FDLNTLASTK.E | Y | 45.74 | 1108.5764 | 10 | 2.5 | 555.2969 | 2 | 25.51 | 1 | 5738 | JVDM\_BotE.raw | 2.653E5 | 1 | 1 | 137 | 146 |  |  | PEAKS DB |
| K.AKLDSVLAAVL | Y | 25.94 | 1098.6648 | 11 | 2.0 | 550.3408 | 2 | 37.27 | 1 | 8723 | JVDM\_BotE.raw | 2.1268E5 | 1 | 1 | 189 | 199 |  |  | PEAKS DB |
| total 3 peptides |  |  |  |  |  |  |  |  |  |  |  |  |  |  |  |  |  |  |  |
| --- | --- | --- | --- | --- | --- | --- | --- | --- | --- | --- | --- | --- | --- | --- | --- | --- | --- | --- | --- |

tr|A0A2K3E0A7|A0A2K3E0A7\_CHLRE

back to list

  

| Protein Coverage
| Supporting Peptides
|

Protein Coverage:

Supporting Peptides:

| Peptide | Uniq | -10lgP | Mass | Length | ppm | m/z | z | RT | Fraction | Scan | Source File | Area Sample 1 | #Feature | #Feature Sample 1 | Start | End | PTM | AScore | Found By |
| --- | --- | --- | --- | --- | --- | --- | --- | --- | --- | --- | --- | --- | --- | --- | --- | --- | --- | --- | --- |
| K.SSQGVPLLAEQASSFFSSK.A | N | 50.34 | 1968.9792 | 19 | 2.2 | 985.4991 | 2 | 37.02 | 1 | 8653 | JVDM\_BotE.raw | 1.1276E5 | 1 | 1 | 130 | 148 |  |  | PEAKS DB |
| K.FVEN(+.98)GGLVVLLGGK.A | Y | 41.47 | 1401.7867 | 14 | 3.3 | 701.9030 | 2 | 35.60 | 1 | 8342 | JVDM\_BotE.raw | 1.2277E4 | 1 | 1 | 87 | 100 | Deamidation (NQ) | N4:Deamidation (NQ):1000.00 | PEAKS PTM |
| K.AQDAWPASLEDAK.T | N | 28.56 | 1400.6571 | 13 | 1.4 | 701.3368 | 2 | 23.56 | 1 | 5155 | JVDM\_BotE.raw | 9.8139E4 | 1 | 1 | 149 | 161 |  |  | PEAKS DB |
| R.HEDVEAFTIPLYTAAGDEMK.V | N | 24.40 | 2236.0356 | 20 | 0.2 | 746.3527 | 3 | 34.04 | 1 | 7975 | JVDM\_BotE.raw | 0 | 0 | 0 | 171 | 190 |  |  | PEAKS DB |
| total 4 peptides |  |  |  |  |  |  |  |  |  |  |  |  |  |  |  |  |  |  |  |
| --- | --- | --- | --- | --- | --- | --- | --- | --- | --- | --- | --- | --- | --- | --- | --- | --- | --- | --- | --- |

tr|A8IKZ2|A8IKZ2\_CHLRE

back to list

  

| Protein Coverage
| Supporting Peptides
|

Protein Coverage:

Supporting Peptides:

| Peptide | Uniq | -10lgP | Mass | Length | ppm | m/z | z | RT | Fraction | Scan | Source File | Area Sample 1 | #Feature | #Feature Sample 1 | Start | End | PTM | AScore | Found By |
| --- | --- | --- | --- | --- | --- | --- | --- | --- | --- | --- | --- | --- | --- | --- | --- | --- | --- | --- | --- |
| K.VAVIVATVTDDVR.L | Y | 67.94 | 1356.7612 | 13 | 2.1 | 679.3893 | 2 | 27.28 | 1 | 6171 | JVDM\_BotE.raw | 7.4732E5 | 2 | 2 | 78 | 90 |  |  | PEAKS DB |
| K.AGGEC(+57.02)LTFDQLALQRPTGK.D | Y | 28.26 | 2061.0312 | 19 | 2.0 | 688.0190 | 3 | 27.00 | 1 | 6098 | JVDM\_BotE.raw | 1.9028E5 | 1 | 1 | 115 | 133 | Carbamidomethylation | C5:Carbamidomethylation:1000.00 | PEAKS DB |
| total 2 peptides |  |  |  |  |  |  |  |  |  |  |  |  |  |  |  |  |  |  |  |
| --- | --- | --- | --- | --- | --- | --- | --- | --- | --- | --- | --- | --- | --- | --- | --- | --- | --- | --- | --- |

tr|A8IGH1|A8IGH1\_CHLRE

back to list

  

| Protein Coverage
| Supporting Peptides
|

Protein Coverage:

Supporting Peptides:

| Peptide | Uniq | -10lgP | Mass | Length | ppm | m/z | z | RT | Fraction | Scan | Source File | Area Sample 1 | #Feature | #Feature Sample 1 | Start | End | PTM | AScore | Found By |
| --- | --- | --- | --- | --- | --- | --- | --- | --- | --- | --- | --- | --- | --- | --- | --- | --- | --- | --- | --- |
| K.SPNAVNPVVEGK.T | Y | 59.10 | 1209.6353 | 12 | 1.7 | 605.8259 | 2 | 13.31 | 1 | 2185 | JVDM\_BotE.raw | 2.0176E5 | 1 | 1 | 176 | 187 |  |  | PEAKS DB |
| K.LINWDAVAQR.Y | Y | 30.75 | 1184.6301 | 10 | 1.2 | 593.3231 | 2 | 26.25 | 1 | 5926 | JVDM\_BotE.raw | 0 | 0 | 0 | 219 | 228 |  |  | PEAKS DB |
| K.TGK(+14.02)LSISK(+14.02)SPNAVN(+.98)PVVEGK.T | Y | 29.17 | 2053.1418 | 20 | -6.5 | 1027.5715 | 2 | 34.19 | 1 | 8014 | JVDM\_BotE.raw | 0 | 0 | 0 | 168 | 187 | Methylation(KR); Deamidation (NQ) | K3:Methylation(KR):15.07;K8:Methylation(KR):0.00;N14:Deamidation (NQ):9.42 | PEAKS PTM |
| K.SPPYALDALEPHMSK.Q | Y | 25.83 | 1654.8025 | 15 | 1.4 | 552.6089 | 3 | 27.02 | 1 | 6120 | JVDM\_BotE.raw | 7.1327E4 | 1 | 1 | 39 | 53 |  |  | PEAKS DB |
| total 4 peptides |  |  |  |  |  |  |  |  |  |  |  |  |  |  |  |  |  |  |  |
| --- | --- | --- | --- | --- | --- | --- | --- | --- | --- | --- | --- | --- | --- | --- | --- | --- | --- | --- | --- |

tr|A8HXL8|A8HXL8\_CHLRE

back to list

  

| Protein Coverage
| Supporting Peptides
|

Protein Coverage:

Supporting Peptides:

| Peptide | Uniq | -10lgP | Mass | Length | ppm | m/z | z | RT | Fraction | Scan | Source File | Area Sample 1 | #Feature | #Feature Sample 1 | Start | End | PTM | AScore | Found By |
| --- | --- | --- | --- | --- | --- | --- | --- | --- | --- | --- | --- | --- | --- | --- | --- | --- | --- | --- | --- |
| R.SLQEALASELAAR.M | Y | 53.97 | 1357.7201 | 13 | 1.7 | 679.8685 | 2 | 29.78 | 1 | 6851 | JVDM\_BotE.raw | 8.1197E4 | 1 | 1 | 300 | 312 |  |  | PEAKS DB |
| K.SVLLVVLTGDR.G | Y | 28.21 | 1170.6971 | 11 | 1.9 | 586.3569 | 2 | 34.70 | 1 | 8147 | JVDM\_BotE.raw | 1.59E5 | 1 | 1 | 110 | 120 |  |  | PEAKS DB |
| MAAMLAS(+79.97)KQGAFMGR.S | Y | 20.73 | 1648.7289 | 15 | -8.9 | 825.3644 | 2 | 17.81 | 1 | 3435 | JVDM\_BotE.raw | 0 | 0 | 0 | 1 | 15 | Phosphorylation (STY) | S7:Phosphorylation (STY):1000.00 | PEAKS PTM |
| total 3 peptides |  |  |  |  |  |  |  |  |  |  |  |  |  |  |  |  |  |  |  |
| --- | --- | --- | --- | --- | --- | --- | --- | --- | --- | --- | --- | --- | --- | --- | --- | --- | --- | --- | --- |

tr|A8HVA3|A8HVA3\_CHLRE

back to list

  

| Protein Coverage
| Supporting Peptides
|

Protein Coverage:

Supporting Peptides:

| Peptide | Uniq | -10lgP | Mass | Length | ppm | m/z | z | RT | Fraction | Scan | Source File | Area Sample 1 | #Feature | #Feature Sample 1 | Start | End | PTM | AScore | Found By |
| --- | --- | --- | --- | --- | --- | --- | --- | --- | --- | --- | --- | --- | --- | --- | --- | --- | --- | --- | --- |
| R.ISGLIYEETR.T | Y | 49.18 | 1179.6135 | 10 | 2.4 | 590.8154 | 2 | 21.89 | 1 | 4629 | JVDM\_BotE.raw | 1.685E5 | 1 | 1 | 47 | 56 |  |  | PEAKS DB |
| total 1 peptides |  |  |  |  |  |  |  |  |  |  |  |  |  |  |  |  |  |  |  |
| --- | --- | --- | --- | --- | --- | --- | --- | --- | --- | --- | --- | --- | --- | --- | --- | --- | --- | --- | --- |

tr|A8HSB0|A8HSB0\_CHLRE

back to list

  

| Protein Coverage
| Supporting Peptides
|

Protein Coverage:

Supporting Peptides:

| Peptide | Uniq | -10lgP | Mass | Length | ppm | m/z | z | RT | Fraction | Scan | Source File | Area Sample 1 | #Feature | #Feature Sample 1 | Start | End | PTM | AScore | Found By |
| --- | --- | --- | --- | --- | --- | --- | --- | --- | --- | --- | --- | --- | --- | --- | --- | --- | --- | --- | --- |
| R.ISGLIYEETR.T | Y | 49.18 | 1179.6135 | 10 | 2.4 | 590.8154 | 2 | 21.89 | 1 | 4629 | JVDM\_BotE.raw | 1.685E5 | 1 | 1 | 47 | 56 |  |  | PEAKS DB |
| total 1 peptides |  |  |  |  |  |  |  |  |  |  |  |  |  |  |  |  |  |  |  |
| --- | --- | --- | --- | --- | --- | --- | --- | --- | --- | --- | --- | --- | --- | --- | --- | --- | --- | --- | --- |

tr|A0A2K3CRJ3|A0A2K3CRJ3\_CHLRE

back to list

  

| Protein Coverage
| Supporting Peptides
|

Protein Coverage:

Supporting Peptides:

| Peptide | Uniq | -10lgP | Mass | Length | ppm | m/z | z | RT | Fraction | Scan | Source File | Area Sample 1 | #Feature | #Feature Sample 1 | Start | End | PTM | AScore | Found By |
| --- | --- | --- | --- | --- | --- | --- | --- | --- | --- | --- | --- | --- | --- | --- | --- | --- | --- | --- | --- |
| R.GLGPGGVELNK.Q | Y | 49.10 | 1039.5662 | 11 | 2.0 | 520.7914 | 2 | 18.17 | 1 | 3526 | JVDM\_BotE.raw | 5.6894E5 | 1 | 1 | 119 | 129 |  |  | PEAKS DB |
| total 1 peptides |  |  |  |  |  |  |  |  |  |  |  |  |  |  |  |  |  |  |  |
| --- | --- | --- | --- | --- | --- | --- | --- | --- | --- | --- | --- | --- | --- | --- | --- | --- | --- | --- | --- |

tr|Q1ALD7|Q1ALD7\_CHLRE

back to list

  

| Protein Coverage
| Supporting Peptides
|

Protein Coverage:

Supporting Peptides:

| Peptide | Uniq | -10lgP | Mass | Length | ppm | m/z | z | RT | Fraction | Scan | Source File | Area Sample 1 | #Feature | #Feature Sample 1 | Start | End | PTM | AScore | Found By |
| --- | --- | --- | --- | --- | --- | --- | --- | --- | --- | --- | --- | --- | --- | --- | --- | --- | --- | --- | --- |
| R.GLGPGGVELNK.Q | Y | 49.10 | 1039.5662 | 11 | 2.0 | 520.7914 | 2 | 18.17 | 1 | 3526 | JVDM\_BotE.raw | 5.6894E5 | 1 | 1 | 119 | 129 |  |  | PEAKS DB |
| total 1 peptides |  |  |  |  |  |  |  |  |  |  |  |  |  |  |  |  |  |  |  |
| --- | --- | --- | --- | --- | --- | --- | --- | --- | --- | --- | --- | --- | --- | --- | --- | --- | --- | --- | --- |

tr|A0A2K3CRI8|A0A2K3CRI8\_CHLRE

back to list

  

| Protein Coverage
| Supporting Peptides
|

Protein Coverage:

Supporting Peptides:

| Peptide | Uniq | -10lgP | Mass | Length | ppm | m/z | z | RT | Fraction | Scan | Source File | Area Sample 1 | #Feature | #Feature Sample 1 | Start | End | PTM | AScore | Found By |
| --- | --- | --- | --- | --- | --- | --- | --- | --- | --- | --- | --- | --- | --- | --- | --- | --- | --- | --- | --- |
| R.GLGPGGVELNK.Q | Y | 49.10 | 1039.5662 | 11 | 2.0 | 520.7914 | 2 | 18.17 | 1 | 3526 | JVDM\_BotE.raw | 5.6894E5 | 1 | 1 | 137 | 147 |  |  | PEAKS DB |
| total 1 peptides |  |  |  |  |  |  |  |  |  |  |  |  |  |  |  |  |  |  |  |
| --- | --- | --- | --- | --- | --- | --- | --- | --- | --- | --- | --- | --- | --- | --- | --- | --- | --- | --- | --- |

tr|A0A2K3D661|A0A2K3D661\_CHLRE

back to list

  

| Protein Coverage
| Supporting Peptides
|

Protein Coverage:

Supporting Peptides:

| Peptide | Uniq | -10lgP | Mass | Length | ppm | m/z | z | RT | Fraction | Scan | Source File | Area Sample 1 | #Feature | #Feature Sample 1 | Start | End | PTM | AScore | Found By |
| --- | --- | --- | --- | --- | --- | --- | --- | --- | --- | --- | --- | --- | --- | --- | --- | --- | --- | --- | --- |
| R.YEDNFDAVNNLVVIAQDTDKK.A | Y | 48.13 | 2410.1653 | 21 | -0.7 | 804.3951 | 3 | 32.87 | 1 | 7638 | JVDM\_BotE.raw | 1.6315E5 | 1 | 1 | 106 | 126 |  |  | PEAKS DB |
| K.AIADFGSQDK.F | Y | 16.11 | 1050.4982 | 10 | 2.1 | 526.2574 | 2 | 15.68 | 1 | 2836 | JVDM\_BotE.raw | 6.119E4 | 1 | 1 | 127 | 136 |  |  | PEAKS DB |
| total 2 peptides |  |  |  |  |  |  |  |  |  |  |  |  |  |  |  |  |  |  |  |
| --- | --- | --- | --- | --- | --- | --- | --- | --- | --- | --- | --- | --- | --- | --- | --- | --- | --- | --- | --- |

tr|A8HP90|A8HP90\_CHLRE

back to list

  

| Protein Coverage
| Supporting Peptides
|

Protein Coverage:

Supporting Peptides:

| Peptide | Uniq | -10lgP | Mass | Length | ppm | m/z | z | RT | Fraction | Scan | Source File | Area Sample 1 | #Feature | #Feature Sample 1 | Start | End | PTM | AScore | Found By |
| --- | --- | --- | --- | --- | --- | --- | --- | --- | --- | --- | --- | --- | --- | --- | --- | --- | --- | --- | --- |
| K.TLDAALLPALSADLK.G | Y | 44.81 | 1510.8606 | 15 | 0.5 | 756.4380 | 2 | 37.27 | 1 | 8757 | JVDM\_BotE.raw | 9.8812E3 | 1 | 1 | 189 | 203 |  |  | PEAKS DB |
| total 1 peptides |  |  |  |  |  |  |  |  |  |  |  |  |  |  |  |  |  |  |  |
| --- | --- | --- | --- | --- | --- | --- | --- | --- | --- | --- | --- | --- | --- | --- | --- | --- | --- | --- | --- |

tr|Q19VH5|Q19VH5\_CHLRE

back to list

  

| Protein Coverage
| Supporting Peptides
|

Protein Coverage:

Supporting Peptides:

| Peptide | Uniq | -10lgP | Mass | Length | ppm | m/z | z | RT | Fraction | Scan | Source File | Area Sample 1 | #Feature | #Feature Sample 1 | Start | End | PTM | AScore | Found By |
| --- | --- | --- | --- | --- | --- | --- | --- | --- | --- | --- | --- | --- | --- | --- | --- | --- | --- | --- | --- |
| K.ELTEADGFVFGFPTR.F | Y | 44.59 | 1684.8096 | 15 | 3.7 | 843.4152 | 2 | 36.17 | 1 | 8463 | JVDM\_BotE.raw | 0 | 0 | 0 | 66 | 80 |  |  | PEAKS DB |
| total 1 peptides |  |  |  |  |  |  |  |  |  |  |  |  |  |  |  |  |  |  |  |
| --- | --- | --- | --- | --- | --- | --- | --- | --- | --- | --- | --- | --- | --- | --- | --- | --- | --- | --- | --- |

tr|Q19VH4|Q19VH4\_CHLRE

back to list

  

| Protein Coverage
| Supporting Peptides
|

Protein Coverage:

Supporting Peptides:

| Peptide | Uniq | -10lgP | Mass | Length | ppm | m/z | z | RT | Fraction | Scan | Source File | Area Sample 1 | #Feature | #Feature Sample 1 | Start | End | PTM | AScore | Found By |
| --- | --- | --- | --- | --- | --- | --- | --- | --- | --- | --- | --- | --- | --- | --- | --- | --- | --- | --- | --- |
| K.ELTEADGFVFGFPTR.F | Y | 44.59 | 1684.8096 | 15 | 3.7 | 843.4152 | 2 | 36.17 | 1 | 8463 | JVDM\_BotE.raw | 0 | 0 | 0 | 66 | 80 |  |  | PEAKS DB |
| total 1 peptides |  |  |  |  |  |  |  |  |  |  |  |  |  |  |  |  |  |  |  |
| --- | --- | --- | --- | --- | --- | --- | --- | --- | --- | --- | --- | --- | --- | --- | --- | --- | --- | --- | --- |

Q08365|RR3\_CHLRE

back to list

  

| Protein Coverage
| Supporting Peptides
|

Protein Coverage:

Supporting Peptides:

| Peptide | Uniq | -10lgP | Mass | Length | ppm | m/z | z | RT | Fraction | Scan | Source File | Area Sample 1 | #Feature | #Feature Sample 1 | Start | End | PTM | AScore | Found By |
| --- | --- | --- | --- | --- | --- | --- | --- | --- | --- | --- | --- | --- | --- | --- | --- | --- | --- | --- | --- |
| K.NPLVNNDFENAEGLTK.L | Y | 44.00 | 1773.8533 | 16 | 2.3 | 887.9359 | 2 | 25.16 | 1 | 5663 | JVDM\_BotE.raw | 4.1153E4 | 1 | 1 | 555 | 570 |  |  | PEAKS DB |
| K.ASTVADSIVDALEK.R | Y | 21.16 | 1417.7300 | 14 | 2.7 | 709.8741 | 2 | 33.30 | 1 | 7758 | JVDM\_BotE.raw | 7.3643E4 | 1 | 1 | 612 | 625 |  |  | PEAKS DB |
| total 2 peptides |  |  |  |  |  |  |  |  |  |  |  |  |  |  |  |  |  |  |  |
| --- | --- | --- | --- | --- | --- | --- | --- | --- | --- | --- | --- | --- | --- | --- | --- | --- | --- | --- | --- |

tr|A8JH68|A8JH68\_CHLRE

back to list

  

| Protein Coverage
| Supporting Peptides
|

Protein Coverage:

Supporting Peptides:

| Peptide | Uniq | -10lgP | Mass | Length | ppm | m/z | z | RT | Fraction | Scan | Source File | Area Sample 1 | #Feature | #Feature Sample 1 | Start | End | PTM | AScore | Found By |
| --- | --- | --- | --- | --- | --- | --- | --- | --- | --- | --- | --- | --- | --- | --- | --- | --- | --- | --- | --- |
| R.DDYLNAPGETYSVK.L | Y | 42.36 | 1570.7151 | 14 | 1.7 | 786.3661 | 2 | 21.37 | 1 | 4473 | JVDM\_BotE.raw | 5.041E4 | 1 | 1 | 106 | 119 |  |  | PEAKS DB |
| total 1 peptides |  |  |  |  |  |  |  |  |  |  |  |  |  |  |  |  |  |  |  |
| --- | --- | --- | --- | --- | --- | --- | --- | --- | --- | --- | --- | --- | --- | --- | --- | --- | --- | --- | --- |

tr|A8I2V3|A8I2V3\_CHLRE

back to list

  

| Protein Coverage
| Supporting Peptides
|

Protein Coverage:

Supporting Peptides:

| Peptide | Uniq | -10lgP | Mass | Length | ppm | m/z | z | RT | Fraction | Scan | Source File | Area Sample 1 | #Feature | #Feature Sample 1 | Start | End | PTM | AScore | Found By |
| --- | --- | --- | --- | --- | --- | --- | --- | --- | --- | --- | --- | --- | --- | --- | --- | --- | --- | --- | --- |
| K.DYGVLIEDGPDAGVTLR.G | Y | 39.29 | 1788.8893 | 17 | -1.9 | 895.4503 | 2 | 30.65 | 1 | 7101 | JVDM\_BotE.raw | 0 | 0 | 0 | 111 | 127 |  |  | PEAKS DB |
| R.GLFIISPTGVLR.Q | Y | 27.67 | 1271.7601 | 12 | 0.0 | 636.8873 | 2 | 37.36 | 1 | 8769 | JVDM\_BotE.raw | 4.5525E4 | 1 | 1 | 128 | 139 |  |  | PEAKS DB |
| total 2 peptides |  |  |  |  |  |  |  |  |  |  |  |  |  |  |  |  |  |  |  |
| --- | --- | --- | --- | --- | --- | --- | --- | --- | --- | --- | --- | --- | --- | --- | --- | --- | --- | --- | --- |

tr|A8J244|A8J244\_CHLRE

back to list

  

| Protein Coverage
| Supporting Peptides
|

Protein Coverage:

Supporting Peptides:

| Peptide | Uniq | -10lgP | Mass | Length | ppm | m/z | z | RT | Fraction | Scan | Source File | Area Sample 1 | #Feature | #Feature Sample 1 | Start | End | PTM | AScore | Found By |
| --- | --- | --- | --- | --- | --- | --- | --- | --- | --- | --- | --- | --- | --- | --- | --- | --- | --- | --- | --- |
| R.LAADVMDVPTLIIAR.T | Y | 39.26 | 1596.8909 | 15 | 2.9 | 799.4550 | 2 | 37.79 | 1 | 8862 | JVDM\_BotE.raw | 6.2649E4 | 1 | 1 | 201 | 215 |  |  | PEAKS DB |
| R.TDALGAYLLTSDADEYDKPFMTGER.T | Y | 21.38 | 2778.2693 | 25 | 1.7 | 927.0986 | 3 | 34.50 | 1 | 8091 | JVDM\_BotE.raw | 0 | 0 | 0 | 216 | 240 |  |  | PEAKS DB |
| total 2 peptides |  |  |  |  |  |  |  |  |  |  |  |  |  |  |  |  |  |  |  |
| --- | --- | --- | --- | --- | --- | --- | --- | --- | --- | --- | --- | --- | --- | --- | --- | --- | --- | --- | --- |

tr|A0A2K3D9K7|A0A2K3D9K7\_CHLRE

back to list

  

| Protein Coverage
| Supporting Peptides
|

Protein Coverage:

Supporting Peptides:

| Peptide | Uniq | -10lgP | Mass | Length | ppm | m/z | z | RT | Fraction | Scan | Source File | Area Sample 1 | #Feature | #Feature Sample 1 | Start | End | PTM | AScore | Found By |
| --- | --- | --- | --- | --- | --- | --- | --- | --- | --- | --- | --- | --- | --- | --- | --- | --- | --- | --- | --- |
| R.LAAEAEAAAAAEAEAAAR.A | Y | 39.09 | 1655.8114 | 18 | 2.7 | 552.9459 | 3 | 24.30 | 1 | 5353 | JVDM\_BotE.raw | 2.3634E4 | 1 | 1 | 627 | 644 |  |  | PEAKS DB |
| total 1 peptides |  |  |  |  |  |  |  |  |  |  |  |  |  |  |  |  |  |  |  |
| --- | --- | --- | --- | --- | --- | --- | --- | --- | --- | --- | --- | --- | --- | --- | --- | --- | --- | --- | --- |

tr|A8HRZ9|A8HRZ9\_CHLRE

back to list

  

| Protein Coverage
| Supporting Peptides
|

Protein Coverage:

Supporting Peptides:

| Peptide | Uniq | -10lgP | Mass | Length | ppm | m/z | z | RT | Fraction | Scan | Source File | Area Sample 1 | #Feature | #Feature Sample 1 | Start | End | PTM | AScore | Found By |
| --- | --- | --- | --- | --- | --- | --- | --- | --- | --- | --- | --- | --- | --- | --- | --- | --- | --- | --- | --- |
| K.AGLQFPVGR.I | Y | 38.66 | 943.5239 | 9 | 1.7 | 472.7700 | 2 | 24.35 | 1 | 5377 | JVDM\_BotE.raw | 3.2553E5 | 1 | 1 | 36 | 44 |  |  | PEAKS DB |
| total 1 peptides |  |  |  |  |  |  |  |  |  |  |  |  |  |  |  |  |  |  |  |
| --- | --- | --- | --- | --- | --- | --- | --- | --- | --- | --- | --- | --- | --- | --- | --- | --- | --- | --- | --- |

tr|A8IR81|A8IR81\_CHLRE

back to list

  

| Protein Coverage
| Supporting Peptides
|

Protein Coverage:

Supporting Peptides:

| Peptide | Uniq | -10lgP | Mass | Length | ppm | m/z | z | RT | Fraction | Scan | Source File | Area Sample 1 | #Feature | #Feature Sample 1 | Start | End | PTM | AScore | Found By |
| --- | --- | --- | --- | --- | --- | --- | --- | --- | --- | --- | --- | --- | --- | --- | --- | --- | --- | --- | --- |
| K.AGLQFPVGR.I | Y | 38.66 | 943.5239 | 9 | 1.7 | 472.7700 | 2 | 24.35 | 1 | 5377 | JVDM\_BotE.raw | 3.2553E5 | 1 | 1 | 21 | 29 |  |  | PEAKS DB |
| total 1 peptides |  |  |  |  |  |  |  |  |  |  |  |  |  |  |  |  |  |  |  |
| --- | --- | --- | --- | --- | --- | --- | --- | --- | --- | --- | --- | --- | --- | --- | --- | --- | --- | --- | --- |

tr|A8HSB5|A8HSB5\_CHLRE

back to list

  

| Protein Coverage
| Supporting Peptides
|

Protein Coverage:

Supporting Peptides:

| Peptide | Uniq | -10lgP | Mass | Length | ppm | m/z | z | RT | Fraction | Scan | Source File | Area Sample 1 | #Feature | #Feature Sample 1 | Start | End | PTM | AScore | Found By |
| --- | --- | --- | --- | --- | --- | --- | --- | --- | --- | --- | --- | --- | --- | --- | --- | --- | --- | --- | --- |
| K.AGLQFPVGR.I | Y | 38.66 | 943.5239 | 9 | 1.7 | 472.7700 | 2 | 24.35 | 1 | 5377 | JVDM\_BotE.raw | 3.2553E5 | 1 | 1 | 21 | 29 |  |  | PEAKS DB |
| total 1 peptides |  |  |  |  |  |  |  |  |  |  |  |  |  |  |  |  |  |  |  |
| --- | --- | --- | --- | --- | --- | --- | --- | --- | --- | --- | --- | --- | --- | --- | --- | --- | --- | --- | --- |

tr|Q42680|Q42680\_CHLRE

back to list

  

| Protein Coverage
| Supporting Peptides
|

Protein Coverage:

Supporting Peptides:

| Peptide | Uniq | -10lgP | Mass | Length | ppm | m/z | z | RT | Fraction | Scan | Source File | Area Sample 1 | #Feature | #Feature Sample 1 | Start | End | PTM | AScore | Found By |
| --- | --- | --- | --- | --- | --- | --- | --- | --- | --- | --- | --- | --- | --- | --- | --- | --- | --- | --- | --- |
| K.AGLQFPVGR.I | Y | 38.66 | 943.5239 | 9 | 1.7 | 472.7700 | 2 | 24.35 | 1 | 5377 | JVDM\_BotE.raw | 3.2553E5 | 1 | 1 | 21 | 29 |  |  | PEAKS DB |
| total 1 peptides |  |  |  |  |  |  |  |  |  |  |  |  |  |  |  |  |  |  |  |
| --- | --- | --- | --- | --- | --- | --- | --- | --- | --- | --- | --- | --- | --- | --- | --- | --- | --- | --- | --- |

tr|A8HWM8|A8HWM8\_CHLRE

back to list

  

| Protein Coverage
| Supporting Peptides
|

Protein Coverage:

Supporting Peptides:

| Peptide | Uniq | -10lgP | Mass | Length | ppm | m/z | z | RT | Fraction | Scan | Source File | Area Sample 1 | #Feature | #Feature Sample 1 | Start | End | PTM | AScore | Found By |
| --- | --- | --- | --- | --- | --- | --- | --- | --- | --- | --- | --- | --- | --- | --- | --- | --- | --- | --- | --- |
| K.AGLQFPVGR.I | Y | 38.66 | 943.5239 | 9 | 1.7 | 472.7700 | 2 | 24.35 | 1 | 5377 | JVDM\_BotE.raw | 3.2553E5 | 1 | 1 | 21 | 29 |  |  | PEAKS DB |
| total 1 peptides |  |  |  |  |  |  |  |  |  |  |  |  |  |  |  |  |  |  |  |
| --- | --- | --- | --- | --- | --- | --- | --- | --- | --- | --- | --- | --- | --- | --- | --- | --- | --- | --- | --- |

tr|A8HWF6|A8HWF6\_CHLRE

back to list

  

| Protein Coverage
| Supporting Peptides
|

Protein Coverage:

Supporting Peptides:

| Peptide | Uniq | -10lgP | Mass | Length | ppm | m/z | z | RT | Fraction | Scan | Source File | Area Sample 1 | #Feature | #Feature Sample 1 | Start | End | PTM | AScore | Found By |
| --- | --- | --- | --- | --- | --- | --- | --- | --- | --- | --- | --- | --- | --- | --- | --- | --- | --- | --- | --- |
| K.AGLQFPVGR.I | Y | 38.66 | 943.5239 | 9 | 1.7 | 472.7700 | 2 | 24.35 | 1 | 5377 | JVDM\_BotE.raw | 3.2553E5 | 1 | 1 | 21 | 29 |  |  | PEAKS DB |
| total 1 peptides |  |  |  |  |  |  |  |  |  |  |  |  |  |  |  |  |  |  |  |
| --- | --- | --- | --- | --- | --- | --- | --- | --- | --- | --- | --- | --- | --- | --- | --- | --- | --- | --- | --- |

tr|A8J363|A8J363\_CHLRE

back to list

  

| Protein Coverage
| Supporting Peptides
|

Protein Coverage:

Supporting Peptides:

| Peptide | Uniq | -10lgP | Mass | Length | ppm | m/z | z | RT | Fraction | Scan | Source File | Area Sample 1 | #Feature | #Feature Sample 1 | Start | End | PTM | AScore | Found By |
| --- | --- | --- | --- | --- | --- | --- | --- | --- | --- | --- | --- | --- | --- | --- | --- | --- | --- | --- | --- |
| R.VAIDQTGIATTDR.L | Y | 37.18 | 1359.6993 | 13 | 2.0 | 680.8583 | 2 | 17.21 | 1 | 3262 | JVDM\_BotE.raw | 1.6009E4 | 1 | 1 | 525 | 537 |  |  | PEAKS DB |
| total 1 peptides |  |  |  |  |  |  |  |  |  |  |  |  |  |  |  |  |  |  |  |
| --- | --- | --- | --- | --- | --- | --- | --- | --- | --- | --- | --- | --- | --- | --- | --- | --- | --- | --- | --- |

Peptide List

  
  

---

Prepared with PEAKS ™ (bioinfor.com)
