## Supplementary material for "Efficient secretion of a plastic degrading enzyme from the green algae *Chlamydomonas reinhardtii*": Dataset - https://doi.org/10.5281/zenodo.13981200: peptides.html

peptide list


Peptide Sequence Contains:

Peptide Sample Area >=

Peptide PTM Contains:

Show Peptide From:

With Protein Accession: False

Peptide List

  

| Peptide | -10lgP | Mass | Length | ppm | m/z | RT | Fraction | Scan | Source File | Area Sample 2 | #Feature | #Feature Sample 2 | Accession | PTM | AScore | Found By |
| --- | --- | --- | --- | --- | --- | --- | --- | --- | --- | --- | --- | --- | --- | --- | --- | --- |
| AAAAASLGQAQDAAAQAQNTAAAAVGNAQTAAGDAAGK | 103.37 | 3323.6089 | 38 | 2.7 | 1108.8799 | 29.67 | 2 | 6748 | JVDM\_TOPE.raw | 2.7853E5 | 2 | 2 | tr\|A0A2K3D8I0\|A0A2K3D8I0\_CHLRE |  |  | PEAKS DB |
| HEDVEAFTIPLYTAAGDETK | 100.81 | 2206.0430 | 20 | 2.6 | 736.3568 | 31.98 | 2 | 7258 | JVDM\_TOPE.raw | 3.5019E5 | 1 | 1 | tr\|A0A2K3E094\|A0A2K3E094\_CHLRE |  |  | PEAKS DB |
| SVGAAGGQEDIVNLVPTAAAATSTGK | 99.14 | 2384.2183 | 26 | 2.2 | 795.7484 | 29.98 | 2 | 6827 | JVDM\_TOPE.raw | 2.6567E5 | 1 | 1 | tr\|A0A2K3DP12\|A0A2K3DP12\_CHLRE |  |  | PEAKS DB |
| GPDPTESSIEAVR | 94.35 | 1356.6521 | 13 | 1.4 | 679.3343 | 16.45 | 2 | 3054 | JVDM\_TOPE.raw | 5.6779E6 | 1 | 1 | PHL7 |  |  | PEAKS DB |
| TVAALYAPSAVLLPTVSNQYR | 93.33 | 2233.2107 | 21 | 1.3 | 745.4118 | 36.17 | 2 | 8323 | JVDM\_TOPE.raw | 2.7241E5 | 2 | 2 | tr\|A0A2K3CTJ7\|A0A2K3CTJ7\_CHLRE |  |  | PEAKS DB |
| IASQGFVVITIDTITR | 92.13 | 1732.9723 | 16 | 2.5 | 867.4956 | 38.43 | 2 | 8913 | JVDM\_TOPE.raw | 2.0567E6 | 2 | 2 | PHL7 |  |  | PEAKS DB |
| LPVDFPAAVASGC(+57.02)AK | 88.47 | 1501.7599 | 15 | 2.7 | 751.8892 | 28.27 | 2 | 6408 | JVDM\_TOPE.raw | 8.9856E4 | 1 | 1 | tr\|A0A2K3DJ22\|A0A2K3DJ22\_CHLRE | Carbamidomethylation | C13:Carbamidomethylation:1000.00 | PEAKS DB |
| AQGAATAATEQAKDAAAAAQQQAAAAADAAK | 88.38 | 2811.3745 | 31 | 1.4 | 938.1334 | 25.47 | 2 | 5780 | JVDM\_TOPE.raw | 1.8533E5 | 1 | 1 | tr\|A0A2K3D8I0\|A0A2K3D8I0\_CHLRE |  |  | PEAKS DB |
| LYDEELQAIAK | 88.34 | 1291.6659 | 11 | 1.5 | 646.8412 | 24.65 | 2 | 5557 | JVDM\_TOPE.raw | 7.0147E6 | 1 | 1 | tr\|A8J6Y8\|A8J6Y8\_CHLRE |  |  | PEAKS DB |
| AAADGAVANAQGAVTSFATK | 88.29 | 1819.9064 | 20 | 3.7 | 910.9639 | 24.13 | 2 | 5430 | JVDM\_TOPE.raw | 1.9643E5 | 2 | 2 | tr\|A0A2K3D8I0\|A0A2K3D8I0\_CHLRE |  |  | PEAKS DB |
| GAAASAVGQATAQATAAVDQAK | 88.08 | 1956.9865 | 22 | 2.7 | 979.5032 | 24.44 | 2 | 5530 | JVDM\_TOPE.raw | 8.8837E5 | 2 | 2 | tr\|A0A2K3D8I0\|A0A2K3D8I0\_CHLRE |  |  | PEAKS DB |
| VIDAGANALVAGSAVFK | 86.20 | 1601.8777 | 17 | 1.9 | 801.9476 | 30.57 | 2 | 6984 | JVDM\_TOPE.raw | 2.9659E4 | 1 | 1 | tr\|A8IKW6\|A8IKW6\_CHLRE |  |  | PEAKS DB |
| TPTLVVGAQLDTIAPVSSHSEAFYNSLPSDLDK | 85.83 | 3471.7410 | 33 | 2.2 | 868.9445 | 36.17 | 2 | 8332 | JVDM\_TOPE.raw | 1.2395E6 | 2 | 2 | PHL7 |  |  | PEAKS DB |
| TLVHNPENMAEAQLLPR | 85.29 | 1931.9886 | 17 | 2.0 | 645.0048 | 25.76 | 2 | 5846 | JVDM\_TOPE.raw | 8.6679E4 | 1 | 1 | tr\|A8IMR9\|A8IMR9\_CHLRE |  |  | PEAKS DB |
| YGDDFDGQTASLC(+57.02)VR | 85.27 | 1702.7257 | 15 | 2.0 | 852.3718 | 22.58 | 2 | 5035 | JVDM\_TOPE.raw | 4.1596E6 | 2 | 2 | tr\|A8J6Y8\|A8J6Y8\_CHLRE | Carbamidomethylation | C13:Carbamidomethylation:1000.00 | PEAKS DB |
| MNSQVSSLSLYDIAGTPGVAADVSHINTK | 85.00 | 2974.4705 | 29 | 2.3 | 992.4997 | 32.93 | 2 | 7484 | JVDM\_TOPE.raw | 0 | 0 | 0 | tr\|A0A2K3D1P1\|A0A2K3D1P1\_CHLRE |  |  | PEAKS DB |
| AQETAAAAAGQAQAAAADAAAK | 83.85 | 1926.9395 | 22 | 1.0 | 964.4780 | 19.45 | 2 | 4039 | JVDM\_TOPE.raw | 3.7236E5 | 1 | 1 | tr\|A0A2K3D8I0\|A0A2K3D8I0\_CHLRE |  |  | PEAKS DB |
| VAVLGAAGGIGQPLSMLMK | 83.13 | 1812.0001 | 19 | 0.4 | 907.0077 | 39.35 | 2 | 9183 | JVDM\_TOPE.raw | 3.8277E4 | 1 | 1 | tr\|A0A2K3D1P1\|A0A2K3D1P1\_CHLRE |  |  | PEAKS DB |
| VEYTELQILC(+57.02)PQTIDSVTGYPMDDPR | 82.55 | 3039.4204 | 26 | 2.3 | 1520.7209 | 37.51 | 2 | 8687 | JVDM\_TOPE.raw | 1.2385E7 | 4 | 4 | tr\|A0A2K3CQ54\|A0A2K3CQ54\_CHLRE | Carbamidomethylation | C10:Carbamidomethylation:1000.00 | PEAKS DB |
| LAAEAEAAAAAEAEAAAR | 82.51 | 1655.8114 | 18 | 1.6 | 552.9453 | 24.27 | 2 | 5458 | JVDM\_TOPE.raw | 3.0578E5 | 2 | 2 | tr\|A0A2K3D9K7\|A0A2K3D9K7\_CHLRE |  |  | PEAKS DB |
| VVIALDSSSHSGADFAAK | 82.15 | 1773.8896 | 18 | 1.8 | 887.9537 | 22.35 | 2 | 4924 | JVDM\_TOPE.raw | 6.4382E5 | 1 | 1 | tr\|A0A2K3D6N5\|A0A2K3D6N5\_CHLRE |  |  | PEAKS DB |
| AGEVQAAAAGAAEQAQAQASAAAAEAQK | 82.07 | 2539.2261 | 28 | 1.6 | 847.4174 | 26.31 | 2 | 5983 | JVDM\_TOPE.raw | 0 | 0 | 0 | tr\|A0A2K3D8I0\|A0A2K3D8I0\_CHLRE |  |  | PEAKS DB |
| LTGTDAYVIPAQHGSAFYSSAEDMSAVSK | 79.55 | 3002.3967 | 29 | 2.5 | 1001.8087 | 27.75 | 2 | 6285 | JVDM\_TOPE.raw | 7.6234E4 | 1 | 1 | tr\|A0A2K3E0A7\|A0A2K3E0A7\_CHLRE |  |  | PEAKS DB |
| VASLYAPTAVLLPTVSDK | 79.46 | 1844.0294 | 18 | 0.9 | 923.0228 | 35.05 | 2 | 8049 | JVDM\_TOPE.raw | 2.4136E5 | 2 | 2 | tr\|A0A2K3CTJ7\|A0A2K3CTJ7\_CHLRE |  |  | PEAKS DB |
| GPFAVAQTTVSR | 79.32 | 1232.6514 | 12 | 2.1 | 617.3342 | 20.39 | 2 | 4297 | JVDM\_TOPE.raw | 1.0429E7 | 1 | 1 | PHL7 |  |  | PEAKS DB |
| ATGETC(+57.02)HIIIETEGK | 79.08 | 1657.7981 | 15 | 2.3 | 829.9082 | 16.76 | 2 | 3147 | JVDM\_TOPE.raw | 2.7798E6 | 2 | 2 | tr\|A8J6Y8\|A8J6Y8\_CHLRE | Carbamidomethylation | C6:Carbamidomethylation:1000.00 | PEAKS DB |
| EPVGVIDNIDGSSPNVR | 77.68 | 1766.8799 | 17 | 2.3 | 884.4492 | 23.49 | 2 | 5258 | JVDM\_TOPE.raw | 2.2935E5 | 2 | 2 | tr\|A0A2K3CTJ7\|A0A2K3CTJ7\_CHLRE |  |  | PEAKS DB |
| GLN(+.98)GAPVVEC(+57.02)TYVESTVTDAPYFASK | 77.00 | 2775.2949 | 26 | 0.0 | 926.1056 | 32.93 | 2 | 7490 | JVDM\_TOPE.raw | 9.5273E4 | 1 | 1 | tr\|A0A2K3D1P1\|A0A2K3D1P1\_CHLRE | Deamidation (NQ); Carbamidomethylation | N3:Deamidation (NQ):1000.00;C10:Carbamidomethylation:1000.00 | PEAKS PTM |
| IDADLDVFIQK | 76.70 | 1275.6710 | 11 | 2.7 | 638.8445 | 30.90 | 2 | 7052 | JVDM\_TOPE.raw | 3.2681E5 | 1 | 1 | tr\|A8JEV1\|A8JEV1\_CHLRE |  |  | PEAKS DB |
| MIGATNPLASEPGTIR | 76.51 | 1626.8400 | 16 | 2.4 | 814.4292 | 25.34 | 2 | 5746 | JVDM\_TOPE.raw | 3.0907E5 | 1 | 1 | tr\|A8J9H8\|A8J9H8\_CHLRE:tr\|A0A2K3DJF1\|A0A2K3DJF1\_CHLRE |  |  | PEAKS DB |
| SSQGVPLLAEQASSFFSSK | 76.47 | 1968.9792 | 19 | 0.6 | 657.3341 | 36.98 | 2 | 8537 | JVDM\_TOPE.raw | 1.1703E6 | 2 | 2 | tr\|A0A2K3E094\|A0A2K3E094\_CHLRE:tr\|A0A2K3E0A7\|A0A2K3E0A7\_CHLRE |  |  | PEAKS DB |
| GSGIANTC(+57.02)PVLESGTTNLK | 76.35 | 1917.9465 | 19 | 4.5 | 959.9849 | 23.38 | 2 | 5237 | JVDM\_TOPE.raw | 3.3615E4 | 1 | 1 | tr\|A8J0E4\|A8J0E4\_CHLRE | Carbamidomethylation | C8:Carbamidomethylation:1000.00 | PEAKS DB |
| ATVAAGEEALTIR | 75.53 | 1300.6986 | 13 | 1.8 | 651.3577 | 21.61 | 2 | 4773 | JVDM\_TOPE.raw | 2.6005E7 | 1 | 1 | tr\|A0A2K3CQ54\|A0A2K3CQ54\_CHLRE |  |  | PEAKS DB |
| TVPTKLEEGEMPLNTYSNK | 75.40 | 2150.0564 | 19 | 2.4 | 1076.0381 | 22.50 | 2 | 4981 | JVDM\_TOPE.raw | 3.1033E6 | 2 | 2 | tr\|A8J6Y8\|A8J6Y8\_CHLRE |  |  | PEAKS DB |
| AAEAQAAAAGAAEQAQAAAANAQQAAADAAGK | 75.24 | 2808.3386 | 32 | 1.6 | 937.1216 | 28.56 | 2 | 6471 | JVDM\_TOPE.raw | 7.6782E4 | 1 | 1 | tr\|A0A2K3D8I0\|A0A2K3D8I0\_CHLRE |  |  | PEAKS DB |
| ASSVAQVLNTLK | 73.85 | 1229.6979 | 12 | 2.1 | 615.8575 | 29.22 | 2 | 6641 | JVDM\_TOPE.raw | 3.0606E5 | 1 | 1 | tr\|B7U1J0\|B7U1J0\_CHLRE:P26526\|ATPA\_CHLRE |  |  | PEAKS DB |
| FVQAGAEVSALLGR | 73.17 | 1416.7725 | 14 | 2.0 | 709.3949 | 31.52 | 2 | 7158 | JVDM\_TOPE.raw | 1.0473E5 | 1 | 1 | P06541\|ATPB\_CHLRE |  |  | PEAKS DB |
| GLC(+57.02)SNFLC(+57.02)DATPGTEISMTGPTGK | 72.12 | 2513.1235 | 24 | 4.8 | 1257.5751 | 31.38 | 2 | 7134 | JVDM\_TOPE.raw | 5.3155E5 | 1 | 1 | tr\|A8J6Y8\|A8J6Y8\_CHLRE | Carbamidomethylation | C3:Carbamidomethylation:1000.00;C8:Carbamidomethylation:1000.00 | PEAKS DB |
| GC(+57.02)PALTVNAVDSYNK | 70.75 | 1607.7614 | 15 | 2.0 | 804.8895 | 21.59 | 2 | 4689 | JVDM\_TOPE.raw | 1.6598E5 | 1 | 1 | tr\|A0A2K3CTJ7\|A0A2K3CTJ7\_CHLRE | Carbamidomethylation | C2:Carbamidomethylation:1000.00 | PEAKS DB |
| LVHDQELSVEER | 70.52 | 1452.7208 | 12 | 0.5 | 485.2478 | 12.86 | 2 | 1892 | JVDM\_TOPE.raw | 1.9723E6 | 2 | 2 | tr\|A8JGV6\|A8JGV6\_CHLRE |  |  | PEAKS DB |
| AAGLSSITVTSDINAVK | 70.42 | 1645.8887 | 17 | 3.0 | 823.9540 | 27.14 | 2 | 6155 | JVDM\_TOPE.raw | 1.0669E5 | 1 | 1 | tr\|A8ILJ9\|A8ILJ9\_CHLRE |  |  | PEAKS DB |
| SLQEALASELAAR | 70.42 | 1357.7201 | 13 | 2.1 | 679.8688 | 29.71 | 2 | 6756 | JVDM\_TOPE.raw | 2.0197E5 | 1 | 1 | tr\|A8HXL8\|A8HXL8\_CHLRE |  |  | PEAKS DB |
| VAPEEHPVLLTEAPLNPK | 70.14 | 1953.0570 | 18 | 4.1 | 977.5398 | 26.13 | 2 | 5946 | JVDM\_TOPE.raw | 1.3114E6 | 1 | 1 | tr\|A8JAV1\|A8JAV1\_CHLRE |  |  | PEAKS DB |
| VGVAGAVVVLGYNWK | 70.13 | 1530.8558 | 15 | 2.8 | 766.4373 | 34.89 | 2 | 8017 | JVDM\_TOPE.raw | 8.0497E4 | 1 | 1 | tr\|A0A2K3D6N5\|A0A2K3D6N5\_CHLRE |  |  | PEAKS DB |
| FVDDDLRYEQFLC(+57.02)PAPDDFAISEYR | 69.77 | 3080.3860 | 25 | 2.3 | 1027.8049 | 36.44 | 2 | 8410 | JVDM\_TOPE.raw | 8.4096E4 | 1 | 1 | PHL7 | Carbamidomethylation | C13:Carbamidomethylation:1000.00 | PEAKS DB |
| AASAAAEQAQAAAVGAQQAAVEK | 69.44 | 2111.0605 | 23 | 2.6 | 704.6959 | 19.48 | 2 | 4036 | JVDM\_TOPE.raw | 2.4879E5 | 1 | 1 | tr\|A0A2K3D8I0\|A0A2K3D8I0\_CHLRE |  |  | PEAKS DB |
| EAAEHTLLAYK | 68.49 | 1244.6400 | 11 | 1.6 | 623.3283 | 17.45 | 2 | 3370 | JVDM\_TOPE.raw | 3.9859E5 | 1 | 1 | tr\|A8JGV6\|A8JGV6\_CHLRE |  |  | PEAKS DB |
| ELDYLDGAVSNPK | 68.47 | 1419.6881 | 13 | 1.0 | 710.8521 | 23.11 | 2 | 5159 | JVDM\_TOPE.raw | 8.4813E4 | 1 | 1 | tr\|A8JC04\|A8JC04\_CHLRE |  |  | PEAKS DB |
| AYGVLTEDGISLR | 68.37 | 1392.7249 | 13 | 3.0 | 697.3718 | 28.21 | 2 | 6390 | JVDM\_TOPE.raw | 4.1242E5 | 1 | 1 | tr\|Q9FE86\|Q9FE86\_CHLRE |  |  | PEAKS DB |
| AGGEC(+57.02)LTFDQLALQRPTGK | 67.65 | 2061.0312 | 19 | 1.8 | 688.0189 | 26.93 | 2 | 6111 | JVDM\_TOPE.raw | 1.6893E5 | 1 | 1 | tr\|A8IKZ2\|A8IKZ2\_CHLRE | Carbamidomethylation | C5:Carbamidomethylation:1000.00 | PEAKS DB |
| LAADVMDVPTLIIAR | 66.34 | 1596.8909 | 15 | 3.3 | 799.4553 | 37.77 | 2 | 8762 | JVDM\_TOPE.raw | 5.2588E4 | 1 | 1 | tr\|A8J244\|A8J244\_CHLRE |  |  | PEAKS DB |
| VAVIVATVTDDVR | 66.22 | 1356.7612 | 13 | 2.1 | 679.3893 | 27.26 | 2 | 6185 | JVDM\_TOPE.raw | 4.0074E5 | 1 | 1 | tr\|A8IKZ2\|A8IKZ2\_CHLRE |  |  | PEAKS DB |
| IAEEAGAVAVMALER | 66.22 | 1528.7919 | 15 | 1.7 | 765.4045 | 29.76 | 2 | 6770 | JVDM\_TOPE.raw | 8.0948E4 | 1 | 1 | tr\|A0A2K3CWZ6\|A0A2K3CWZ6\_CHLRE |  |  | PEAKS DB |
| IEDLSAQTQAAAAEQFK | 66.12 | 1819.8951 | 17 | 3.3 | 910.9578 | 22.90 | 2 | 5102 | JVDM\_TOPE.raw | 3.2087E4 | 1 | 1 | tr\|A0A2K3CWF7\|A0A2K3CWF7\_CHLRE |  |  | PEAKS DB |
| ADDVAVQVGATAAGAAGAKPPAGVPK | 66.09 | 2288.2124 | 26 | 1.3 | 763.7457 | 22.72 | 2 | 5054 | JVDM\_TOPE.raw | 5.1235E4 | 1 | 1 | tr\|A0A2K3D8I0\|A0A2K3D8I0\_CHLRE |  |  | PEAKS DB |
| AQAVFDQEFQEITLSK | 65.92 | 1852.9207 | 16 | 1.7 | 927.4692 | 31.90 | 2 | 7243 | JVDM\_TOPE.raw | 2.9274E5 | 1 | 1 | tr\|Q9FE86\|Q9FE86\_CHLRE |  |  | PEAKS DB |
| YEQFLC(+57.02)PAPDDFAISEYR | 65.79 | 2219.9834 | 18 | 1.1 | 1111.0002 | 33.50 | 2 | 7663 | JVDM\_TOPE.raw | 4.5602E5 | 2 | 2 | PHL7 | Carbamidomethylation | C6:Carbamidomethylation:1000.00 | PEAKS DB |
| DAAAAAQQQAAAAADAAK | 65.14 | 1612.7804 | 18 | 5.6 | 807.4020 | 17.20 | 2 | 3292 | JVDM\_TOPE.raw | 1.1098E5 | 1 | 1 | tr\|A0A2K3D8I0\|A0A2K3D8I0\_CHLRE |  |  | PEAKS DB |
| KVIGVTSLDIVR | 64.27 | 1298.7921 | 12 | 1.5 | 433.9386 | 26.02 | 2 | 5915 | JVDM\_TOPE.raw | 3.1815E5 | 1 | 1 | tr\|P93106\|P93106\_CHLRE |  |  | PEAKS DB |
| VGSDGSAELKEDDGIDYAATTVQLPGGER | 63.94 | 2949.3838 | 29 | 0.9 | 984.1361 | 26.62 | 2 | 6042 | JVDM\_TOPE.raw | 0 | 0 | 0 | tr\|A8J0E4\|A8J0E4\_CHLRE |  |  | PEAKS DB |
| SPNAVNPVVEGK | 63.87 | 1209.6353 | 12 | 2.1 | 605.8262 | 13.19 | 2 | 2057 | JVDM\_TOPE.raw | 3.9422E5 | 1 | 1 | tr\|A8IGH1\|A8IGH1\_CHLRE |  |  | PEAKS DB |
| LALALGDEVGQPLPLAAASNAQYIAAR | 62.51 | 2692.4548 | 27 | 3.0 | 898.4949 | 37.69 | 2 | 8745 | JVDM\_TOPE.raw | 5.061E4 | 1 | 1 | tr\|A0A2K3DP15\|A0A2K3DP15\_CHLRE |  |  | PEAKS DB |
| GPDPTESSIEAVRGPFAVAQTTVSR | 62.50 | 2571.2927 | 25 | 2.4 | 858.1069 | 28.89 | 2 | 6569 | JVDM\_TOPE.raw | 6.2711E6 | 1 | 1 | PHL7 |  |  | PEAKS DB |
| KMIGATNPLASEPGTIR | 62.15 | 1754.9348 | 17 | 2.6 | 585.9871 | 22.37 | 2 | 4932 | JVDM\_TOPE.raw | 3.1472E5 | 1 | 1 | tr\|A8J9H8\|A8J9H8\_CHLRE:tr\|A0A2K3DJF1\|A0A2K3DJF1\_CHLRE |  |  | PEAKS DB |
| LVTDFADGAYTAHAAGR | 61.52 | 1734.8325 | 17 | 1.4 | 579.2856 | 21.86 | 2 | 4774 | JVDM\_TOPE.raw | 2.5121E5 | 1 | 1 | tr\|A0A2K3DZ46\|A0A2K3DZ46\_CHLRE |  |  | PEAKS DB |
| YEDNFDAVNNLVVIAQDTDKK | 61.17 | 2410.1653 | 21 | 2.2 | 804.3975 | 32.79 | 2 | 7436 | JVDM\_TOPE.raw | 2.4859E5 | 1 | 1 | tr\|A0A2K3D661\|A0A2K3D661\_CHLRE |  |  | PEAKS DB |
| VASLYAPTAVLLPTVSDKVR | 61.16 | 2099.1990 | 20 | 2.4 | 700.7419 | 33.57 | 2 | 7687 | JVDM\_TOPE.raw | 2.5472E5 | 1 | 1 | tr\|A0A2K3CTJ7\|A0A2K3CTJ7\_CHLRE |  |  | PEAKS DB |
| AAGDNAQAAAVPSVTEQASEFLAGSWPK | 61.06 | 2772.3354 | 28 | 2.0 | 925.1210 | 38.36 | 2 | 8908 | JVDM\_TOPE.raw | 2.2897E5 | 1 | 1 | tr\|A0A2K3D6N5\|A0A2K3D6N5\_CHLRE |  |  | PEAKS DB |
| AAQDIALVDLPPTHPIR | 60.56 | 1826.0050 | 17 | 1.9 | 609.6768 | 28.26 | 2 | 6412 | JVDM\_TOPE.raw | 2.1308E6 | 2 | 2 | tr\|A8JGV6\|A8JGV6\_CHLRE |  |  | PEAKS DB |
| VIGVTSLDIVR | 59.77 | 1170.6971 | 11 | 1.4 | 586.3567 | 29.76 | 2 | 6771 | JVDM\_TOPE.raw | 3.3453E5 | 1 | 1 | tr\|P93106\|P93106\_CHLRE |  |  | PEAKS DB |
| GVALIAPDTSPR | 59.73 | 1195.6560 | 12 | 0.8 | 598.8358 | 21.70 | 2 | 4725 | JVDM\_TOPE.raw | 2.0243E5 | 1 | 1 | tr\|A8J7F8\|A8J7F8\_CHLRE:tr\|A0A2K3DW85\|A0A2K3DW85\_CHLRE:tr\|A0A2K3DW83\|A0A2K3DW83\_CHLRE |  |  | PEAKS DB |
| AEDTAAAAQAQAAAAAESAR | 59.62 | 1843.8660 | 20 | 3.0 | 615.6311 | 18.73 | 2 | 3783 | JVDM\_TOPE.raw | 1.5539E5 | 1 | 1 | tr\|A0A2K3D8I0\|A0A2K3D8I0\_CHLRE |  |  | PEAKS DB |
| ADETVAVEEVAAAEPASGK | 59.09 | 1842.8846 | 19 | 2.1 | 922.4515 | 21.98 | 2 | 4833 | JVDM\_TOPE.raw | 2.0515E4 | 1 | 1 | tr\|A8HRZ0\|A8HRZ0\_CHLRE |  |  | PEAKS DB |
| ASEAATGALSEIAAVDGEGGAAK | 58.95 | 2044.9912 | 23 | -0.2 | 1023.5027 | 27.55 | 2 | 6258 | JVDM\_TOPE.raw | 1.635E4 | 1 | 1 | tr\|A0A2K3D8I0\|A0A2K3D8I0\_CHLRE |  |  | PEAKS DB |
| SHAEKPLVGSVAPDFK | 58.89 | 1680.8834 | 16 | 2.4 | 561.3031 | 20.09 | 2 | 4241 | JVDM\_TOPE.raw | 2.5658E5 | 1 | 1 | tr\|Q9FE86\|Q9FE86\_CHLRE |  |  | PEAKS DB |
| SPPYALDALEPHMSK | 58.86 | 1654.8025 | 15 | 2.5 | 552.6095 | 26.96 | 2 | 6116 | JVDM\_TOPE.raw | 2.211E5 | 1 | 1 | tr\|A8IGH1\|A8IGH1\_CHLRE |  |  | PEAKS DB |
| EGGLGDLAYPLVADLKK | 58.75 | 1757.9563 | 17 | 2.1 | 586.9940 | 34.51 | 2 | 7919 | JVDM\_TOPE.raw | 2.7066E5 | 1 | 1 | tr\|Q9FE86\|Q9FE86\_CHLRE |  |  | PEAKS DB |
| AAAAQATAAAAGAQEAASAAAGQATATATDAMTK | 58.65 | 2961.4097 | 34 | 1.9 | 988.1458 | 30.05 | 2 | 6837 | JVDM\_TOPE.raw | 1.6568E5 | 1 | 1 | tr\|A0A2K3D8I0\|A0A2K3D8I0\_CHLRE |  |  | PEAKS DB |
| AAAASAQGALAAATGQAR | 58.56 | 1555.8066 | 18 | 1.2 | 778.9115 | 19.92 | 2 | 4157 | JVDM\_TOPE.raw | 1.5906E5 | 1 | 1 | tr\|A0A2K3D8I0\|A0A2K3D8I0\_CHLRE |  |  | PEAKS DB |
| ELEDAAIVQAFTR | 57.92 | 1461.7463 | 13 | 0.6 | 731.8809 | 32.81 | 2 | 7441 | JVDM\_TOPE.raw | 4.1742E4 | 1 | 1 | tr\|A0A2K3DZ46\|A0A2K3DZ46\_CHLRE |  |  | PEAKS DB |
| GQAEAAAAQAQNTAAAAVGNVQTAAADAAAK | 57.63 | 2781.3640 | 31 | 3.1 | 928.1315 | 29.63 | 2 | 6727 | JVDM\_TOPE.raw | 2.4063E5 | 1 | 1 | tr\|A0A2K3D8I0\|A0A2K3D8I0\_CHLRE |  |  | PEAKS DB |
| IGSLLDQSITR | 57.14 | 1201.6666 | 11 | 0.2 | 601.8407 | 24.69 | 2 | 5594 | JVDM\_TOPE.raw | 4.4689E4 | 1 | 1 | tr\|A0A2K3E350\|A0A2K3E350\_CHLRE |  |  | PEAKS DB |
| VAIDQTGIATTDR | 57.10 | 1359.6993 | 13 | 1.7 | 680.8581 | 17.14 | 2 | 3285 | JVDM\_TOPE.raw | 9.5905E4 | 1 | 1 | tr\|A8J363\|A8J363\_CHLRE |  |  | PEAKS DB |
| ATMSAEVLDALTK | 56.78 | 1348.6908 | 13 | 2.7 | 675.3545 | 32.06 | 2 | 7282 | JVDM\_TOPE.raw | 1.097E5 | 1 | 1 | tr\|A0A2K3D1P1\|A0A2K3D1P1\_CHLRE |  |  | PEAKS DB |
| SIISTQLGTELEK | 56.52 | 1417.7664 | 13 | 2.1 | 709.8920 | 26.83 | 2 | 6082 | JVDM\_TOPE.raw | 1.8672E5 | 1 | 1 | tr\|Q6UKY5\|Q6UKY5\_CHLRE |  |  | PEAKS DB |
| GGVIMDVTTAEEAR | 55.65 | 1447.6976 | 14 | 2.4 | 724.8578 | 23.51 | 2 | 5296 | JVDM\_TOPE.raw | 2.0724E5 | 1 | 1 | tr\|A0A2K3CWZ6\|A0A2K3CWZ6\_CHLRE |  |  | PEAKS DB |
| QLVQAGFDLK | 54.48 | 1117.6132 | 10 | 1.0 | 559.8144 | 25.03 | 2 | 5676 | JVDM\_TOPE.raw | 6.3693E5 | 1 | 1 | tr\|A0A2K3E094\|A0A2K3E094\_CHLRE |  |  | PEAKS DB |
| LEEGEMPLNTYSNK | 54.26 | 1623.7450 | 14 | 3.6 | 812.8827 | 20.86 | 2 | 4479 | JVDM\_TOPE.raw | 4.0525E4 | 1 | 1 | tr\|A8J6Y8\|A8J6Y8\_CHLRE |  |  | PEAKS DB |
| GIVDSEDLPLNISR | 53.87 | 1526.7939 | 14 | 2.5 | 764.4061 | 29.52 | 2 | 6715 | JVDM\_TOPE.raw | 0 | 0 | 0 | tr\|A8J1U1\|A8J1U1\_CHLRE |  |  | PEAKS DB |
| KAGEVQAAAAGAAEQAQAQASAAAAEAQK | 53.69 | 2667.3210 | 29 | 2.4 | 890.1164 | 24.59 | 2 | 5570 | JVDM\_TOPE.raw | 0 | 0 | 0 | tr\|A0A2K3D8I0\|A0A2K3D8I0\_CHLRE |  |  | PEAKS DB |
| SFSLGAAPSTK | 53.49 | 1064.5502 | 11 | 1.1 | 533.2830 | 19.13 | 2 | 3962 | JVDM\_TOPE.raw | 2.2551E5 | 1 | 1 | tr\|A8HXL8\|A8HXL8\_CHLRE |  |  | PEAKS DB |
| GTFLDSDC(+57.02)FDQVGTK | 53.49 | 1688.7352 | 15 | 2.4 | 845.3770 | 26.93 | 2 | 6118 | JVDM\_TOPE.raw | 7.5896E4 | 1 | 1 | tr\|A0A2K3CTJ7\|A0A2K3CTJ7\_CHLRE | Carbamidomethylation | C8:Carbamidomethylation:1000.00 | PEAKS DB |
| SESVIAGGADPC(+57.02)TGAVSESANLLSALAR | 53.48 | 2702.3181 | 28 | 3.2 | 901.7828 | 37.51 | 2 | 8698 | JVDM\_TOPE.raw | 4.6999E4 | 1 | 1 | tr\|A0A2K3DJ22\|A0A2K3DJ22\_CHLRE | Carbamidomethylation | C12:Carbamidomethylation:1000.00 | PEAKS DB |
| STLTFTPEAEGLVK | 53.33 | 1491.7820 | 14 | 2.6 | 746.9002 | 28.36 | 2 | 6435 | JVDM\_TOPE.raw | 1.1766E5 | 1 | 1 | tr\|B7U1J0\|B7U1J0\_CHLRE:P26526\|ATPA\_CHLRE |  |  | PEAKS DB |
| TISAFSPISNPINAPWGVK | 53.14 | 1998.0574 | 19 | 3.2 | 1000.0392 | 36.17 | 2 | 8336 | JVDM\_TOPE.raw | 1.373E4 | 1 | 1 | tr\|A8J7F8\|A8J7F8\_CHLRE:tr\|A0A2K3DW85\|A0A2K3DW85\_CHLRE:tr\|A0A2K3DW83\|A0A2K3DW83\_CHLRE |  |  | PEAKS DB |
| LSTLNIQAYDAAR | 53.09 | 1434.7467 | 13 | 2.2 | 718.3822 | 24.56 | 2 | 5549 | JVDM\_TOPE.raw | 6.5093E4 | 1 | 1 | tr\|Q9LD42\|Q9LD42\_CHLRE |  |  | PEAKS DB |
| NAIVQVDATPFK | 53.07 | 1301.6979 | 12 | 2.4 | 651.8578 | 25.75 | 2 | 5842 | JVDM\_TOPE.raw | 1.0858E5 | 1 | 1 | tr\|A8HVQ1\|A8HVQ1\_CHLRE |  |  | PEAKS DB |
| VALTALTMAEYFR | 52.19 | 1484.7697 | 13 | 3.7 | 743.3948 | 37.84 | 2 | 8787 | JVDM\_TOPE.raw | 4.4348E4 | 1 | 1 | P06541\|ATPB\_CHLRE |  |  | PEAKS DB |
| GC(+57.02)DLVIIPAGVPR | 52.13 | 1365.7438 | 13 | 3.0 | 683.8812 | 30.51 | 2 | 6978 | JVDM\_TOPE.raw | 1.5891E5 | 1 | 1 | tr\|A0A2K3D1P1\|A0A2K3D1P1\_CHLRE | Carbamidomethylation | C2:Carbamidomethylation:1000.00 | PEAKS DB |
| AGPQEQWGALLNK | 51.84 | 1410.7256 | 13 | 1.6 | 706.3712 | 27.17 | 2 | 6147 | JVDM\_TOPE.raw | 3.855E5 | 1 | 1 | tr\|A0A2K3D6N5\|A0A2K3D6N5\_CHLRE |  |  | PEAKS DB |
| AGAAASAVQSTAASAVEK | 51.60 | 1588.8057 | 18 | 1.2 | 795.4111 | 16.39 | 2 | 3021 | JVDM\_TOPE.raw | 2.7426E3 | 1 | 1 | tr\|A0A2K3E7M7\|A0A2K3E7M7\_CHLRE |  |  | PEAKS DB |
| DYGVLIEDGPDAGVTLR | 51.51 | 1788.8893 | 17 | 1.1 | 895.4529 | 30.59 | 2 | 6987 | JVDM\_TOPE.raw | 2.5548E4 | 1 | 1 | tr\|A8I2V3\|A8I2V3\_CHLRE |  |  | PEAKS DB |
| FVEN(+.98)GGLVVLLGGK | 51.27 | 1401.7867 | 14 | 1.9 | 701.9020 | 35.53 | 2 | 8156 | JVDM\_TOPE.raw | 1.1712E5 | 1 | 1 | tr\|A0A2K3E0A7\|A0A2K3E0A7\_CHLRE | Deamidation (NQ) | N4:Deamidation (NQ):1000.00 | PEAKS PTM |
| LINWDAVAQR | 51.20 | 1184.6301 | 10 | 2.3 | 593.3237 | 26.21 | 2 | 5959 | JVDM\_TOPE.raw | 3.6451E5 | 1 | 1 | tr\|A8IGH1\|A8IGH1\_CHLRE |  |  | PEAKS DB |
| SVLLVVLTGDR | 51.19 | 1170.6971 | 11 | 1.9 | 586.3569 | 34.63 | 2 | 7958 | JVDM\_TOPE.raw | 2.6572E5 | 1 | 1 | tr\|A8HXL8\|A8HXL8\_CHLRE |  |  | PEAKS DB |
| GLGPGGVELNK | 50.86 | 1039.5662 | 11 | 0.8 | 520.7908 | 18.12 | 2 | 3574 | JVDM\_TOPE.raw | 7.1637E5 | 1 | 1 | tr\|A0A2K3CRJ3\|A0A2K3CRJ3\_CHLRE:tr\|Q1ALD7\|Q1ALD7\_CHLRE:tr\|A0A2K3CRI8\|A0A2K3CRI8\_CHLRE |  |  | PEAKS DB |
| AQDAWPASLEDAK | 50.77 | 1400.6571 | 13 | 1.5 | 701.3369 | 23.58 | 2 | 5316 | JVDM\_TOPE.raw | 8.6981E5 | 1 | 1 | tr\|A0A2K3E094\|A0A2K3E094\_CHLRE:tr\|A0A2K3E0A7\|A0A2K3E0A7\_CHLRE |  |  | PEAKS DB |
| AAATGAAAAAGSQLASAQEAGK | 50.67 | 1871.9336 | 22 | 0.9 | 624.9857 | 16.37 | 2 | 3018 | JVDM\_TOPE.raw | 0 | 0 | 0 | tr\|A0A2K3DMT2\|A0A2K3DMT2\_CHLRE |  |  | PEAKS DB |
| DDLFNTNAGIVK | 50.57 | 1305.6565 | 12 | 2.5 | 653.8372 | 25.91 | 2 | 5876 | JVDM\_TOPE.raw | 3.6104E5 | 1 | 1 | tr\|P93106\|P93106\_CHLRE |  |  | PEAKS DB |
| FAN(+.98)GTVSIANTLAGLTPK | 50.37 | 1774.9465 | 18 | 2.5 | 888.4828 | 36.72 | 2 | 8488 | JVDM\_TOPE.raw | 4.1581E4 | 1 | 1 | tr\|A8JC04\|A8JC04\_CHLRE | Deamidation (NQ) | N3:Deamidation (NQ):61.82 | PEAKS PTM |
| GDFAIEVGR | 49.99 | 962.4821 | 9 | 0.2 | 482.2484 | 22.17 | 2 | 4857 | JVDM\_TOPE.raw | 1.1754E6 | 1 | 1 | tr\|A8J9H8\|A8J9H8\_CHLRE:tr\|A0A2K3DJF1\|A0A2K3DJF1\_CHLRE |  |  | PEAKS DB |
| SIVGTSLEVIQK | 49.93 | 1272.7289 | 12 | 2.5 | 637.3733 | 26.87 | 2 | 6099 | JVDM\_TOPE.raw | 2.2313E5 | 1 | 1 | tr\|A8J1A3\|A8J1A3\_CHLRE:tr\|A0A2K3DE61\|A0A2K3DE61\_CHLRE |  |  | PEAKS DB |
| GADLIVIPAGVPR | 49.77 | 1276.7502 | 13 | 3.3 | 639.3845 | 31.38 | 2 | 7137 | JVDM\_TOPE.raw | 5.9717E5 | 1 | 1 | tr\|P93106\|P93106\_CHLRE |  |  | PEAKS DB |
| TAIAVDTILNQK | 49.20 | 1285.7241 | 12 | 2.2 | 643.8707 | 26.91 | 2 | 6101 | JVDM\_TOPE.raw | 3.2313E5 | 1 | 1 | tr\|B7U1J0\|B7U1J0\_CHLRE:P26526\|ATPA\_CHLRE |  |  | PEAKS DB |
| GFDKDGLAEALR | 49.19 | 1290.6567 | 12 | 1.7 | 646.3367 | 23.06 | 2 | 5149 | JVDM\_TOPE.raw | 3.9778E4 | 1 | 1 | tr\|A0A2K3D1P1\|A0A2K3D1P1\_CHLRE |  |  | PEAKS DB |
| FDLNTLASTK | 48.55 | 1108.5764 | 10 | 2.1 | 555.2966 | 25.53 | 2 | 5786 | JVDM\_TOPE.raw | 4.9507E5 | 1 | 1 | tr\|A8JEV1\|A8JEV1\_CHLRE |  |  | PEAKS DB |
| VALVYGQMNEPPGAR | 47.79 | 1600.8031 | 15 | 7.0 | 801.4144 | 22.97 | 2 | 5073 | JVDM\_TOPE.raw | 6.9289E4 | 1 | 1 | P06541\|ATPB\_CHLRE |  |  | PEAKS DB |
| MPSAVGYQPTLATEMGGLQER | 47.69 | 2235.0664 | 21 | 2.1 | 746.0310 | 29.53 | 2 | 6716 | JVDM\_TOPE.raw | 0 | 0 | 0 | P06541\|ATPB\_CHLRE |  |  | PEAKS DB |
| VLPGVAALDEK | 46.73 | 1110.6284 | 11 | 0.7 | 556.3219 | 24.52 | 2 | 5547 | JVDM\_TOPE.raw | 2.0309E5 | 1 | 1 | tr\|A8JC04\|A8JC04\_CHLRE |  |  | PEAKS DB |
| AMMPELLASIEK | 46.67 | 1331.6829 | 12 | 1.9 | 666.8500 | 35.31 | 2 | 8120 | JVDM\_TOPE.raw | 1.3983E5 | 1 | 1 | tr\|A0A2K3D1P1\|A0A2K3D1P1\_CHLRE |  |  | PEAKS DB |
| GMMPGIQDMLER | 46.49 | 1376.6250 | 12 | 2.2 | 689.3213 | 32.39 | 2 | 7322 | JVDM\_TOPE.raw | 1.8587E6 | 1 | 1 | tr\|A8J6Y8\|A8J6Y8\_CHLRE |  |  | PEAKS DB |
| ITSASTVC(+57.02)SHEDSSAVVIPLYTYEDGEDVR | 46.21 | 3299.5139 | 30 | 2.6 | 1100.8481 | 30.00 | 2 | 6845 | JVDM\_TOPE.raw | 6.1684E4 | 1 | 1 | tr\|A0A2K3D6N5\|A0A2K3D6N5\_CHLRE | Carbamidomethylation | C8:Carbamidomethylation:1000.00 | PEAKS DB |
| ASTVADSIVDALEK | 45.96 | 1417.7300 | 14 | 3.3 | 709.8746 | 33.28 | 2 | 7574 | JVDM\_TOPE.raw | 4.1721E4 | 1 | 1 | Q08365\|RR3\_CHLRE |  |  | PEAKS DB |
| HAPNAVLEIITNPVN(+.98)STVPIAVETLK | 45.71 | 2740.5010 | 26 | 0.2 | 914.5078 | 38.74 | 2 | 9023 | JVDM\_TOPE.raw | 0 | 0 | 0 | tr\|P93106\|P93106\_CHLRE | Deamidation (NQ) | N15:Deamidation (NQ):16.03 | PEAKS PTM |
| GPVGAIDKYEIR | 45.64 | 1316.7087 | 12 | 2.3 | 439.9112 | 19.99 | 2 | 4239 | JVDM\_TOPE.raw | 3.6154E5 | 1 | 1 | tr\|A0A2K3CTJ7\|A0A2K3CTJ7\_CHLRE |  |  | PEAKS DB |
| AAAAELTLAAYK | 45.17 | 1191.6499 | 12 | 1.5 | 596.8331 | 23.81 | 2 | 5355 | JVDM\_TOPE.raw | 2.1097E5 | 1 | 1 | tr\|Q7X7A7\|Q7X7A7\_CHLRE |  |  | PEAKS DB |
| HAPNAVLEIITNPVNSTVPIAVETLK | 44.37 | 2739.5171 | 26 | 2.2 | 914.1816 | 40.38 | 2 | 9416 | JVDM\_TOPE.raw | 4.828E4 | 1 | 1 | tr\|P93106\|P93106\_CHLRE |  |  | PEAKS DB |
| LFASLGNPVSLEK | 43.07 | 1373.7554 | 13 | 1.7 | 687.8861 | 29.92 | 2 | 6812 | JVDM\_TOPE.raw | 0 | 0 | 0 | tr\|A0A2K3DZB3\|A0A2K3DZB3\_CHLRE |  |  | PEAKS DB |
| TPTLVVGAQ(+.98)LDTIAPVSSHSEAFYN(+.98)SLPSDLDK | 42.94 | 3473.7090 | 33 | 8.8 | 1158.9204 | 35.45 | 2 | 8142 | JVDM\_TOPE.raw | 2.0264E4 | 1 | 1 | PHL7 | Deamidation (NQ) | Q9:Deamidation (NQ):1000.00;N25:Deamidation (NQ):1000.00 | PEAKS PTM |
| TTLVNLFSNLEK | 42.92 | 1377.7504 | 12 | 1.3 | 689.8834 | 37.21 | 2 | 8627 | JVDM\_TOPE.raw | 3.2444E4 | 1 | 1 | Q08365\|RR3\_CHLRE |  |  | PEAKS DB |
| NPLVNNDFENAEGLTK | 42.35 | 1773.8533 | 16 | 0.9 | 887.9348 | 25.22 | 2 | 5733 | JVDM\_TOPE.raw | 2.0028E4 | 1 | 1 | Q08365\|RR3\_CHLRE |  |  | PEAKS DB |
| AAEEAGLDLPYSC(+57.02)R | 42.06 | 1550.7035 | 14 | 2.6 | 776.3610 | 23.53 | 2 | 5283 | JVDM\_TOPE.raw | 0 | 0 | 0 | tr\|A8IV40\|A8IV40\_CHLRE | Carbamidomethylation | C13:Carbamidomethylation:1000.00 | PEAKS DB |
| VLSGYNDVIMAR | 41.74 | 1336.6809 | 12 | 1.7 | 669.3489 | 24.49 | 2 | 5542 | JVDM\_TOPE.raw | 7.1279E4 | 1 | 1 | tr\|A8ILJ9\|A8ILJ9\_CHLRE |  |  | PEAKS DB |
| ATVAAGEEALTIRN | 41.65 | 1414.7416 | 14 | 1.9 | 708.3794 | 19.90 | 2 | 4202 | JVDM\_TOPE.raw | 6.8017E4 | 1 | 1 | tr\|A0A2K3CQ54\|A0A2K3CQ54\_CHLRE |  |  | PEAKS DB |
| LLAPSADVVAAGEAAQPHC(+57.02)SR | 41.48 | 2119.0481 | 21 | 3.5 | 707.3591 | 22.84 | 2 | 5090 | JVDM\_TOPE.raw | 1.4618E4 | 1 | 1 | tr\|A0A2K3D7P8\|A0A2K3D7P8\_CHLRE | Carbamidomethylation | C19:Carbamidomethylation:1000.00 | PEAKS DB |
| RPFVAIVGGSK | 41.27 | 1129.6608 | 11 | 1.7 | 565.8386 | 19.92 | 2 | 4206 | JVDM\_TOPE.raw | 5.5773E4 | 1 | 1 | tr\|A8JC04\|A8JC04\_CHLRE |  |  | PEAKS DB |
| VAQPLGGYDSGLPK | 41.15 | 1400.7300 | 14 | -1.3 | 701.3713 | 22.16 | 2 | 4865 | JVDM\_TOPE.raw | 8.6358E4 | 1 | 1 | tr\|A8J363\|A8J363\_CHLRE |  |  | PEAKS DB |
| MNSQ(+.98)VSSLSLYDIAGTPGVAADVSHINTK | 40.55 | 2975.4546 | 29 | 6.9 | 992.8323 | 32.87 | 2 | 7484 | JVDM\_TOPE.raw | 9.3073E4 | 1 | 1 | tr\|A0A2K3D1P1\|A0A2K3D1P1\_CHLRE | Deamidation (NQ) | Q4:Deamidation (NQ):26.31 | PEAKS PTM |
| LYSIASSR | 40.32 | 895.4763 | 8 | 1.3 | 448.7460 | 15.18 | 2 | 2710 | JVDM\_TOPE.raw | 5.6978E6 | 1 | 1 | tr\|A8J6Y8\|A8J6Y8\_CHLRE |  |  | PEAKS DB |
| ILQSIEQSEQAK | 39.89 | 1372.7197 | 12 | 1.3 | 687.3680 | 15.30 | 2 | 2698 | JVDM\_TOPE.raw | 6.568E4 | 1 | 1 | tr\|Q7X7A7\|Q7X7A7\_CHLRE |  |  | PEAKS DB |
| VLILDADNAGSAVR | 39.81 | 1412.7623 | 14 | 3.3 | 707.3907 | 24.90 | 2 | 5659 | JVDM\_TOPE.raw | 8.3093E3 | 1 | 1 | tr\|A0A2K3CPA4\|A0A2K3CPA4\_CHLRE |  |  | PEAKS DB |
| GPDPTESSIEAVRGPF | 39.19 | 1657.7947 | 16 | 3.1 | 829.9072 | 27.62 | 2 | 6270 | JVDM\_TOPE.raw | 1.4792E4 | 1 | 1 | PHL7 |  |  | PEAKS DB |
| LLAAGQGEAVVK | 38.56 | 1154.6659 | 12 | 2.3 | 578.3416 | 17.28 | 2 | 3314 | JVDM\_TOPE.raw | 2.0075E5 | 1 | 1 | tr\|A0A2K3CQL0\|A0A2K3CQL0\_CHLRE:tr\|A0A2K3CQP7\|A0A2K3CQP7\_CHLRE |  |  | PEAKS DB |
| AAGLDAVDTVSVVK | 38.48 | 1343.7296 | 14 | 4.5 | 672.8751 | 24.68 | 2 | 5596 | JVDM\_TOPE.raw | 9.6926E4 | 1 | 1 | tr\|O48949\|O48949\_CHLRE |  |  | PEAKS DB |
| TVPTKLEEGEMPLN(+.98)TYSNK | 38.40 | 2151.0405 | 19 | 2.7 | 718.0227 | 22.37 | 2 | 4934 | JVDM\_TOPE.raw | 1.5243E5 | 1 | 1 | tr\|A8J6Y8\|A8J6Y8\_CHLRE | Deamidation (NQ) | N14:Deamidation (NQ):34.30 | PEAKS PTM |
| AALAAVPAVC(+57.02)R | 38.37 | 1097.6016 | 11 | 3.0 | 549.8097 | 20.00 | 2 | 4223 | JVDM\_TOPE.raw | 1.0497E5 | 1 | 1 | tr\|A0A2K3CQL0\|A0A2K3CQL0\_CHLRE:tr\|A0A2K3CQP7\|A0A2K3CQP7\_CHLRE | Carbamidomethylation | C10:Carbamidomethylation:1000.00 | PEAKS DB |
| VTGYTGPEELGAC(+57.02)LK | 38.28 | 1593.7709 | 15 | 2.7 | 797.8948 | 24.27 | 2 | 5445 | JVDM\_TOPE.raw | 4.4207E5 | 1 | 1 | tr\|P93106\|P93106\_CHLRE | Carbamidomethylation | C13:Carbamidomethylation:1000.00 | PEAKS DB |
| QILC(+57.02)PQTIDSVTGYPMDDPR | 38.02 | 2305.0718 | 20 | 1.9 | 769.3660 | 30.14 | 2 | 6859 | JVDM\_TOPE.raw | 2.2885E5 | 1 | 1 | tr\|A0A2K3CQ54\|A0A2K3CQ54\_CHLRE | Carbamidomethylation | C4:Carbamidomethylation:1000.00 | PEAKS DB |
| QLQAALDHLR | 38.01 | 1163.6411 | 10 | 1.2 | 388.8881 | 19.33 | 2 | 3981 | JVDM\_TOPE.raw | 5.9083E6 | 2 | 2 | PHL7 |  |  | PEAKS DB |
| LAEQAERYDEMVEEMKK | 37.42 | 2097.9709 | 17 | 1.4 | 700.3319 | 21.95 | 2 | 4799 | JVDM\_TOPE.raw | 8.8522E5 | 2 | 2 | tr\|A8JGV6\|A8JGV6\_CHLRE |  |  | PEAKS DB |
| TNIVLEATR | 36.79 | 1015.5662 | 9 | 1.1 | 508.7909 | 19.58 | 2 | 4062 | JVDM\_TOPE.raw | 2.2058E5 | 1 | 1 | tr\|A8HMY6\|A8HMY6\_CHLRE |  |  | PEAKS DB |
| LDYALSR | 36.43 | 836.4392 | 7 | 1.4 | 419.2274 | 16.44 | 2 | 2991 | JVDM\_TOPE.raw | 5.1405E6 | 1 | 1 | tr\|A8J6Y8\|A8J6Y8\_CHLRE |  |  | PEAKS DB |
| ALTSGVDAATAGR | 36.18 | 1188.6099 | 13 | 1.1 | 595.3129 | 13.37 | 2 | 2116 | JVDM\_TOPE.raw | 8.6069E4 | 1 | 1 | tr\|A0A2K3E3X9\|A0A2K3E3X9\_CHLRE |  |  | PEAKS DB |
| AAVTIPVMAK | 36.17 | 999.5787 | 10 | 1.9 | 500.7976 | 24.23 | 2 | 5467 | JVDM\_TOPE.raw | 1.0159E5 | 1 | 1 | tr\|A0A2K3CWZ6\|A0A2K3CWZ6\_CHLRE |  |  | PEAKS DB |
| ALNYDGNWISC(+57.02)K | 36.12 | 1439.6504 | 12 | 2.7 | 720.8344 | 24.27 | 2 | 5475 | JVDM\_TOPE.raw | 1.1377E5 | 1 | 1 | tr\|A0A2K3D6N5\|A0A2K3D6N5\_CHLRE | Carbamidomethylation | C11:Carbamidomethylation:1000.00 | PEAKS DB |
| VSAASLLDVSTTTDKK | 35.66 | 1634.8727 | 16 | 1.6 | 545.9657 | 24.14 | 2 | 5434 | JVDM\_TOPE.raw | 2.2817E5 | 1 | 1 | tr\|A0A2K3D661\|A0A2K3D661\_CHLRE |  |  | PEAKS DB |
| TIDSVTGYPMDDPR | 35.43 | 1565.7031 | 14 | 2.4 | 783.8607 | 21.42 | 2 | 4705 | JVDM\_TOPE.raw | 3.8917E5 | 1 | 1 | tr\|A0A2K3CQ54\|A0A2K3CQ54\_CHLRE |  |  | PEAKS DB |
| VLLLPADANAPLIC(+57.02)VATGTGIAPFR | 35.35 | 2549.4038 | 25 | 1.7 | 850.8100 | 40.57 | 2 | 9488 | JVDM\_TOPE.raw | 0 | 0 | 0 | tr\|A8J6Y8\|A8J6Y8\_CHLRE | Carbamidomethylation | C14:Carbamidomethylation:1000.00 | PEAKS DB |
| LAVNLIPFPR | 35.27 | 1138.6863 | 10 | 2.1 | 570.3516 | 35.74 | 2 | 8215 | JVDM\_TOPE.raw | 9.359E4 | 1 | 1 | tr\|A8IXZ0\|A8IXZ0\_CHLRE |  |  | PEAKS DB |
| VVAQAFGK | 33.53 | 818.4650 | 8 | 0.3 | 410.2399 | 13.31 | 2 | 2086 | JVDM\_TOPE.raw | 9.4349E5 | 1 | 1 | tr\|A0A2K3D6N5\|A0A2K3D6N5\_CHLRE |  |  | PEAKS DB |
| VEYTELQILCPQT(+79.97)IDSVTGYPMDDPR | 33.45 | 3062.3655 | 26 | 5.8 | 766.6031 | 37.44 | 2 | 8686 | JVDM\_TOPE.raw | 0 | 0 | 0 | tr\|A0A2K3CQ54\|A0A2K3CQ54\_CHLRE | Phosphorylation (STY) | T13:Phosphorylation (STY):5.77 | PEAKS PTM |
| TISFNTWC(+57.02)R | 33.40 | 1183.5444 | 9 | 2.4 | 592.7809 | 25.61 | 2 | 5813 | JVDM\_TOPE.raw | 2.8165E5 | 1 | 1 | tr\|A0A2K3E094\|A0A2K3E094\_CHLRE:tr\|A0A2K3E0A7\|A0A2K3E0A7\_CHLRE | Carbamidomethylation | C8:Carbamidomethylation:1000.00 | PEAKS DB |
| NQWHVEVY | 33.37 | 1073.4930 | 8 | 1.0 | 537.7543 | 24.02 | 2 | 5392 | JVDM\_TOPE.raw | 6.8438E5 | 1 | 1 | tr\|A8J6Y8\|A8J6Y8\_CHLRE |  |  | PEAKS DB |
| AAPAANGGGASGAAAAVK | 32.97 | 1410.7214 | 18 | 2.0 | 706.3694 | 20.31 | 2 | 4318 | JVDM\_TOPE.raw | 0 | 0 | 0 | tr\|A0A2K3DFK4\|A0A2K3DFK4\_CHLRE |  |  | PEAKS DB |
| RGSAAGGGAAAPPSLPGATAAATSSSGTGGGGGGGGR | 32.62 | 2910.3928 | 37 | -4.0 | 728.6025 | 20.60 | 2 | 4404 | JVDM\_TOPE.raw | 8.5138E4 | 1 | 1 | tr\|A0A2K3D7S0\|A0A2K3D7S0\_CHLRE |  |  | PEAKS DB |
| HDVGIGGGGNGGGGALSSGGPR | 32.58 | 1834.8669 | 22 | 5.4 | 918.4457 | 20.83 | 2 | 4469 | JVDM\_TOPE.raw | 0 | 0 | 0 | tr\|A0A2K3E270\|A0A2K3E270\_CHLRE:tr\|A0A2K3E262\|A0A2K3E262\_CHLRE |  |  | PEAKS DB |
| DGPQPGWGDLLAK | 32.46 | 1352.6724 | 13 | 0.1 | 677.3435 | 29.89 | 2 | 6810 | JVDM\_TOPE.raw | 7.162E4 | 1 | 1 | tr\|A0A2K3DZ46\|A0A2K3DZ46\_CHLRE |  |  | PEAKS DB |
| FVDDDLRYEQ(+.98)FLC(+57.02)PAPDDFAISEYR | 32.19 | 3081.3701 | 25 | 9.0 | 1028.1399 | 36.48 | 2 | 8419 | JVDM\_TOPE.raw | 0 | 0 | 0 | PHL7 | Deamidation (NQ); Carbamidomethylation | Q10:Deamidation (NQ):1000.00;C13:Carbamidomethylation:1000.00 | PEAKS PTM |
| LTGTDAYVIPAQ(+.98)HGSAFYSSAEDMSAVSK | 32.18 | 3003.3806 | 29 | 6.6 | 1002.1407 | 27.78 | 2 | 6313 | JVDM\_TOPE.raw | 1.3806E5 | 1 | 1 | tr\|A0A2K3E0A7\|A0A2K3E0A7\_CHLRE | Deamidation (NQ) | Q12:Deamidation (NQ):1000.00 | PEAKS PTM |
| SDIIVSPSILSADFSR | 32.02 | 1705.8887 | 16 | 2.8 | 853.9540 | 35.53 | 2 | 8153 | JVDM\_TOPE.raw | 0 | 0 | 0 | tr\|A8IKW6\|A8IKW6\_CHLRE |  |  | PEAKS DB |
| LESARGGGPAATAAAR | 31.86 | 1454.7589 | 16 | 8.0 | 728.3926 | 17.13 | 2 | 3277 | JVDM\_TOPE.raw | 0 | 0 | 0 | tr\|A0A2K3CWQ4\|A0A2K3CWQ4\_CHLRE |  |  | PEAKS DB |
| GLETTDGVMESPQ(+.98)SVVFQEAENR | 31.65 | 2523.1436 | 23 | 7.2 | 842.0612 | 29.85 | 2 | 6801 | JVDM\_TOPE.raw | 4.5335E4 | 1 | 1 | tr\|A8ILJ9\|A8ILJ9\_CHLRE | Deamidation (NQ) | Q13:Deamidation (NQ):61.30 | PEAKS PTM |
| GLFIISPTGVLR | 31.65 | 1271.7601 | 12 | 2.1 | 636.8887 | 37.34 | 2 | 8653 | JVDM\_TOPE.raw | 1.0668E5 | 1 | 1 | tr\|A8I2V3\|A8I2V3\_CHLRE |  |  | PEAKS DB |
| LTFDEIQGLTYLQVK | 31.35 | 1766.9454 | 15 | -3.8 | 884.4766 | 36.84 | 2 | 8507 | JVDM\_TOPE.raw | 0 | 0 | 0 | tr\|A8J0E4\|A8J0E4\_CHLRE |  |  | PEAKS DB |
| AAESAAASAAQAQATAASAVSAVQDK | 31.28 | 2345.1458 | 26 | 3.1 | 782.7250 | 25.94 | 2 | 5890 | JVDM\_TOPE.raw | 0 | 0 | 0 | tr\|A0A2K3D8I0\|A0A2K3D8I0\_CHLRE |  |  | PEAKS DB |
| TGSSVPAIK | 31.17 | 858.4811 | 9 | 3.4 | 430.2493 | 12.91 | 2 | 1936 | JVDM\_TOPE.raw | 1.1617E5 | 1 | 1 | tr\|A8HRZ0\|A8HRZ0\_CHLRE |  |  | PEAKS DB |
| AGLQFPVGR | 31.10 | 943.5239 | 9 | 2.3 | 472.7703 | 24.38 | 2 | 5490 | JVDM\_TOPE.raw | 3.1308E5 | 1 | 1 |  |  |  | PEAKS DB |
| AITGVYVQPK | 31.05 | 1074.6073 | 10 | 1.1 | 538.3115 | 17.66 | 2 | 3445 | JVDM\_TOPE.raw | 9.5715E4 | 1 | 1 |  |  |  | PEAKS DB |
| GSGGSGGGRGGGKAAAGK | 31.01 | 1387.6915 | 18 | 2.3 | 694.8546 | 12.56 | 2 | 1567 | JVDM\_TOPE.raw | 0 | 0 | 0 |  |  |  | PEAKS DB |
| VAAGGGGGGGGGTAAAASGGGK | 30.84 | 1543.7339 | 22 | 9.0 | 772.8812 | 30.95 | 2 | 7058 | JVDM\_TOPE.raw | 8.3681E4 | 1 | 1 |  |  |  | PEAKS DB |
| LTYTLDAMSGSFK | 30.79 | 1432.6908 | 13 | 3.4 | 717.3551 | 28.93 | 2 | 6572 | JVDM\_TOPE.raw | 1.5469E4 | 1 | 1 | tr\|A8J0E4\|A8J0E4\_CHLRE |  |  | PEAKS DB |
| HEDVEAFTIPLYTAAGDEMK | 30.63 | 2236.0356 | 20 | 3.5 | 746.3551 | 33.87 | 2 | 7785 | JVDM\_TOPE.raw | 1.0031E5 | 1 | 1 | tr\|A0A2K3E0A7\|A0A2K3E0A7\_CHLRE |  |  | PEAKS DB |
| GQVPNIYNALTIR | 30.58 | 1457.7991 | 13 | 1.6 | 729.9080 | 31.77 | 2 | 7232 | JVDM\_TOPE.raw | 3.9054E4 | 1 | 1 | P06541\|ATPB\_CHLRE |  |  | PEAKS DB |
| TAPAFVDLDTR | 30.31 | 1204.6088 | 11 | 2.2 | 603.3130 | 24.85 | 2 | 5627 | JVDM\_TOPE.raw | 1.6021E5 | 1 | 1 | P06541\|ATPB\_CHLRE |  |  | PEAKS DB |
| AKLDSVLAAVL | 30.17 | 1098.6648 | 11 | 1.2 | 550.3403 | 37.15 | 2 | 8608 | JVDM\_TOPE.raw | 3.5223E5 | 1 | 1 | tr\|A8JEV1\|A8JEV1\_CHLRE |  |  | PEAKS DB |
| GTSLDSDC(+57.02)FDQVGTK | 30.08 | 1628.6989 | 15 | 1.6 | 815.3580 | 19.45 | 2 | 4019 | JVDM\_TOPE.raw | 1.0192E5 | 1 | 1 |  | Carbamidomethylation | C8:Carbamidomethylation:1000.00 | PEAKS DB |
| AKAAAEGGAGAALGAAAASSTAAPGK | 29.66 | 2097.0813 | 26 | -4.9 | 700.0309 | 28.32 | 2 | 6417 | JVDM\_TOPE.raw | 0 | 0 | 0 |  |  |  | PEAKS DB |
| SGAAYISTLEGR | 29.57 | 1223.6146 | 12 | 2.0 | 612.8158 | 21.88 | 2 | 4794 | JVDM\_TOPE.raw | 0 | 0 | 0 |  |  |  | PEAKS DB |
| VYGPIEIPC(+57.02)TGSQPTK | 29.56 | 1745.8658 | 16 | 2.2 | 873.9421 | 25.09 | 2 | 5691 | JVDM\_TOPE.raw | 9.6984E4 | 1 | 1 | tr\|A0A2K3DP12\|A0A2K3DP12\_CHLRE | Carbamidomethylation | C9:Carbamidomethylation:1000.00 | PEAKS DB |
| VVSGLPEDTPEQPVLPTVAGVTLR | 28.95 | 2473.3428 | 24 | 5.2 | 825.4592 | 34.61 | 2 | 7946 | JVDM\_TOPE.raw | 0 | 0 | 0 |  |  |  | PEAKS DB |
| VAVLANEQELSVEER | 28.91 | 1684.8632 | 15 | -0.3 | 843.4386 | 23.18 | 2 | 5194 | JVDM\_TOPE.raw | 6.7261E4 | 1 | 1 | tr\|Q7X7A7\|Q7X7A7\_CHLRE |  |  | PEAKS DB |
| LYQGGAGAGGMPGGAPGAGAAPSGGSGAGPK | 28.48 | 2482.1658 | 31 | 1.8 | 828.3973 | 16.67 | 2 | 3138 | JVDM\_TOPE.raw | 0 | 0 | 0 |  |  |  | PEAKS DB |
| EVSALLGGAASSSGGAGGKK | 28.30 | 1702.8849 | 20 | -8.1 | 852.4428 | 29.46 | 2 | 6700 | JVDM\_TOPE.raw | 9.8831E4 | 1 | 1 |  |  |  | PEAKS DB |
| LTLSAYDPVTR | 28.16 | 1234.6558 | 11 | 1.9 | 618.3363 | 24.90 | 2 | 5647 | JVDM\_TOPE.raw | 6.0102E4 | 1 | 1 |  |  |  | PEAKS DB |
| LTPVVAR | 28.09 | 754.4701 | 7 | 1.6 | 378.2429 | 13.58 | 2 | 2184 | JVDM\_TOPE.raw | 1.3424E5 | 1 | 1 | tr\|A8JC04\|A8JC04\_CHLRE |  |  | PEAKS DB |
| QGGAGGAGGDGSGAGGGGAGSKPR | 27.98 | 1841.8364 | 24 | -2.7 | 921.9230 | 26.51 | 2 | 6025 | JVDM\_TOPE.raw | 6.1331E4 | 1 | 1 |  |  |  | PEAKS DB |
| IAEGAAMIR | 27.92 | 930.4957 | 9 | -0.2 | 466.2550 | 16.17 | 2 | 2962 | JVDM\_TOPE.raw | 1.2289E5 | 1 | 1 | tr\|A0A2K3CWZ6\|A0A2K3CWZ6\_CHLRE |  |  | PEAKS DB |
| YSIAWLK | 27.85 | 879.4854 | 7 | 2.3 | 440.7510 | 29.00 | 2 | 6577 | JVDM\_TOPE.raw | 1.4417E6 | 1 | 1 | PHL7 |  |  | PEAKS DB |
| KEPVGVIDNIDGSSPNVR | 27.66 | 1894.9749 | 18 | 1.9 | 632.6667 | 21.03 | 2 | 4493 | JVDM\_TOPE.raw | 1.972E5 | 1 | 1 | tr\|A0A2K3CTJ7\|A0A2K3CTJ7\_CHLRE |  |  | PEAKS DB |
| FGSVSC(+57.02)LYYDNR | 27.51 | 1479.6453 | 12 | -0.1 | 740.8298 | 23.42 | 2 | 5251 | JVDM\_TOPE.raw | 0 | 0 | 0 |  | Carbamidomethylation | C6:Carbamidomethylation:1000.00 | PEAKS DB |
| DAGAGTGASPAAGAK | 27.30 | 1200.5735 | 15 | 2.9 | 601.2958 | 19.70 | 2 | 4120 | JVDM\_TOPE.raw | 4.3175E5 | 1 | 1 |  |  |  | PEAKS DB |
| AGGVAGAGGAAGAGGARAGSGGGVR | 27.09 | 1867.9360 | 25 | 4.6 | 934.9796 | 27.58 | 2 | 6251 | JVDM\_TOPE.raw | 0 | 0 | 0 |  |  |  | PEAKS DB |
| TGPPAVTVEDADKLK | 27.07 | 1539.8143 | 15 | 1.2 | 514.2794 | 18.99 | 2 | 3898 | JVDM\_TOPE.raw | 9.9186E4 | 1 | 1 | tr\|O48949\|O48949\_CHLRE |  |  | PEAKS DB |
| LGAAAAASAAGGK | 26.80 | 1014.5458 | 13 | 1.5 | 508.2809 | 35.37 | 2 | 8114 | JVDM\_TOPE.raw | 2.6829E5 | 2 | 2 |  |  |  | PEAKS DB |
| ELIIGDR | 26.68 | 814.4548 | 7 | 1.9 | 408.2355 | 18.74 | 2 | 3800 | JVDM\_TOPE.raw | 3.0707E5 | 1 | 1 | tr\|B7U1J0\|B7U1J0\_CHLRE:P26526\|ATPA\_CHLRE |  |  | PEAKS DB |
| AFGGAGTGPGSVASGGAPGSGGAK | 26.64 | 1873.8917 | 24 | 4.1 | 625.6404 | 21.43 | 2 | 4637 | JVDM\_TOPE.raw | 1.3298E5 | 1 | 1 |  |  |  | PEAKS DB |
| GLETTDGVMESPQSVVFQEAENR | 26.61 | 2522.1594 | 23 | 1.6 | 841.7284 | 29.97 | 2 | 6821 | JVDM\_TOPE.raw | 0 | 0 | 0 | tr\|A8ILJ9\|A8ILJ9\_CHLRE |  |  | PEAKS DB |
| WGELLEK | 26.48 | 873.4596 | 7 | 0.8 | 437.7374 | 23.13 | 2 | 5157 | JVDM\_TOPE.raw | 5.748E5 | 1 | 1 | tr\|A0A2K3E094\|A0A2K3E094\_CHLRE:tr\|A0A2K3E0A7\|A0A2K3E0A7\_CHLRE |  |  | PEAKS DB |
| FAAAIHAQYPGK | 26.30 | 1272.6615 | 12 | 1.9 | 425.2286 | 16.51 | 2 | 3050 | JVDM\_TOPE.raw | 1.6491E5 | 1 | 1 | tr\|A8J244\|A8J244\_CHLRE |  |  | PEAKS DB |
| HLAAAAAAGGGGGGR | 26.13 | 1192.6061 | 15 | 0.7 | 597.3107 | 21.33 | 2 | 4627 | JVDM\_TOPE.raw | 0 | 0 | 0 |  |  |  | PEAKS DB |
| AAGAGAAGASAGAAAPVLTG | 26.08 | 1510.7739 | 20 | 3.1 | 756.3966 | 31.93 | 2 | 7253 | JVDM\_TOPE.raw | 0 | 0 | 0 |  |  |  | PEAKS DB |
| LAGGAGAPAAPAVPAAGGGA | 25.82 | 1502.7841 | 20 | -6.0 | 752.3948 | 21.16 | 2 | 4559 | JVDM\_TOPE.raw | 1.1837E5 | 1 | 1 |  |  |  | PEAKS DB |
| MTAGGAAGGGGGAVSVSAR | 25.62 | 1532.7365 | 19 | 4.9 | 767.3793 | 32.42 | 2 | 7350 | JVDM\_TOPE.raw | 0 | 0 | 0 |  |  |  | PEAKS DB |
| ISGLIYEETR | 25.60 | 1179.6135 | 10 | 0.9 | 590.8146 | 21.88 | 2 | 4808 | JVDM\_TOPE.raw | 1.5526E5 | 1 | 1 |  |  |  | PEAKS DB |
| GAAGLGQASPFAAGAPAPDAGAGAGEFGVVAAAGR | 25.56 | 2966.4634 | 35 | -3.4 | 989.8251 | 37.47 | 2 | 8694 | JVDM\_TOPE.raw | 3.9856E4 | 1 | 1 |  |  |  | PEAKS DB |
| AVGAGAGDGGAGAGAGLAER | 25.27 | 1583.7651 | 20 | -2.8 | 792.8876 | 22.35 | 2 | 4918 | JVDM\_TOPE.raw | 3.1927E4 | 1 | 1 |  |  |  | PEAKS DB |
| PGGGGAAAVGSLTAAASAGGGGK | 25.09 | 1740.8754 | 23 | 6.1 | 581.3026 | 19.78 | 2 | 4170 | JVDM\_TOPE.raw | 2.0443E5 | 1 | 1 |  |  |  | PEAKS DB |
| GPSGGGGVAAAGGAAGGDPPR | 25.02 | 1634.7760 | 21 | 9.1 | 818.4027 | 17.43 | 2 | 3378 | JVDM\_TOPE.raw | 0 | 0 | 0 |  |  |  | PEAKS DB |
| GSVAVGDEAEAVR | 25.01 | 1258.6152 | 13 | 2.8 | 630.3167 | 13.32 | 2 | 2073 | JVDM\_TOPE.raw | 4.6791E5 | 1 | 1 |  |  |  | PEAKS DB |
| VINKVYGPIEIPC(+57.02)TGSQPTK | 24.99 | 2200.1562 | 20 | 3.2 | 734.3950 | 25.04 | 2 | 5688 | JVDM\_TOPE.raw | 3.0425E4 | 1 | 1 | tr\|A0A2K3DP12\|A0A2K3DP12\_CHLRE | Carbamidomethylation | C13:Carbamidomethylation:1000.00 | PEAKS DB |
| APTDGAAGAHTAK | 24.97 | 1166.5680 | 13 | -8.4 | 584.2864 | 25.15 | 2 | 5711 | JVDM\_TOPE.raw | 0 | 0 | 0 |  |  |  | PEAKS DB |
| FTAGVGVVSAR | 24.96 | 1062.5822 | 11 | -2.4 | 532.2971 | 17.59 | 2 | 3434 | JVDM\_TOPE.raw | 0 | 0 | 0 |  |  |  | PEAKS DB |
| AAGAAGGTAGGSSSAGGAK | 24.96 | 1404.6593 | 19 | 4.8 | 703.3403 | 29.46 | 2 | 6696 | JVDM\_TOPE.raw | 1.5772E5 | 1 | 1 |  |  |  | PEAKS DB |
| ASTAVTTDMSKR | 24.67 | 1266.6238 | 12 | 1.3 | 423.2158 | 12.41 | 2 | 1486 | JVDM\_TOPE.raw | 1.9971E6 | 1 | 1 | tr\|A8J6Y8\|A8J6Y8\_CHLRE |  |  | PEAKS DB |
| LVAFDNQK | 24.65 | 933.4919 | 8 | 1.3 | 467.7538 | 16.01 | 2 | 2892 | JVDM\_TOPE.raw | 7.6448E5 | 1 | 1 |  |  |  | PEAKS DB |
| AGAGSGGGGGGGGVNGSAAEPLLQPQLR | 24.52 | 2391.1890 | 28 | 6.0 | 798.0751 | 27.65 | 2 | 6265 | JVDM\_TOPE.raw | 0 | 0 | 0 |  |  |  | PEAKS DB |
| LVLPGELAK | 24.37 | 938.5800 | 9 | 1.3 | 470.2979 | 25.47 | 2 | 5785 | JVDM\_TOPE.raw | 1.814E5 | 1 | 1 |  |  |  | PEAKS DB |
| RGSGDGGGEAGGGVDLAGQASLQSR | 24.09 | 2258.0635 | 25 | 6.7 | 753.7001 | 23.85 | 2 | 5363 | JVDM\_TOPE.raw | 6.9096E4 | 1 | 1 |  |  |  | PEAKS DB |
| TLAALAAGGGAASPAGGGGAADGFTPK | 24.08 | 2213.1077 | 27 | -9.7 | 1107.5504 | 23.71 | 2 | 5330 | JVDM\_TOPE.raw | 0 | 0 | 0 |  |  |  | PEAKS DB |
| LDNIVFR | 24.01 | 875.4865 | 7 | 1.6 | 438.7512 | 23.29 | 2 | 5203 | JVDM\_TOPE.raw | 1.8522E5 | 1 | 1 |  |  |  | PEAKS DB |
| VAGAAGAAAGAGAAAAPAAAS | 24.00 | 1523.7692 | 21 | 9.3 | 762.8989 | 16.96 | 2 | 3235 | JVDM\_TOPE.raw | 2.6555E4 | 1 | 1 |  |  |  | PEAKS DB |
| GIVYGKPVNQGITQLKPAR | 23.64 | 2038.1687 | 19 | 1.8 | 680.3981 | 19.79 | 2 | 4162 | JVDM\_TOPE.raw | 1.7965E5 | 2 | 2 |  |  |  | PEAKS DB |
| VNAAYVIGTSTK | 23.52 | 1222.6558 | 12 | 5.2 | 612.3383 | 18.75 | 2 | 3815 | JVDM\_TOPE.raw | 2.8123E4 | 1 | 1 |  |  |  | PEAKS DB |
| AGGGGAAATAAGGGGGGGAAVGGGGGGKGG | 23.50 | 1997.9263 | 30 | 5.2 | 999.9756 | 25.04 | 2 | 5683 | JVDM\_TOPE.raw | 1.1414E5 | 1 | 1 |  |  |  | PEAKS DB |
| GGAAVGGADPAAAP | 23.48 | 1080.5199 | 14 | 1.9 | 541.2682 | 12.79 | 2 | 1836 | JVDM\_TOPE.raw | 0 | 0 | 0 |  |  |  | PEAKS DB |
| MSQEVANFFVSEAR | 23.40 | 1613.7507 | 14 | 2.0 | 807.8843 | 30.00 | 2 | 6832 | JVDM\_TOPE.raw | 6.781E4 | 1 | 1 |  |  |  | PEAKS DB |
| AAGGSKAAAGKGGAGK | 23.40 | 1257.6788 | 16 | 3.5 | 629.8489 | 29.75 | 2 | 6763 | JVDM\_TOPE.raw | 6.8767E4 | 1 | 1 |  |  |  | PEAKS DB |
| ATGLVAVDAMIPVGR | 23.16 | 1468.8071 | 15 | -0.3 | 735.4106 | 33.56 | 2 | 7680 | JVDM\_TOPE.raw | 0 | 0 | 0 | tr\|B7U1J0\|B7U1J0\_CHLRE:P26526\|ATPA\_CHLRE |  |  | PEAKS DB |
| LAGVYDPK | 23.16 | 861.4596 | 8 | 1.2 | 431.7376 | 14.37 | 2 | 2395 | JVDM\_TOPE.raw | 4.0843E5 | 1 | 1 | tr\|P93106\|P93106\_CHLRE |  |  | PEAKS DB |
| GDGGLGPGAAAGSAAAMGTPQSKQR | 23.15 | 2212.0654 | 25 | 6.3 | 738.3671 | 23.14 | 2 | 5195 | JVDM\_TOPE.raw | 8.9286E4 | 1 | 1 |  |  |  | PEAKS DB |
| ADELPTTNPIR | 22.92 | 1225.6302 | 11 | 3.3 | 613.8244 | 19.14 | 2 | 3951 | JVDM\_TOPE.raw | 0 | 0 | 0 | tr\|Q7X7A7\|Q7X7A7\_CHLRE |  |  | PEAKS DB |
| GALAAAPLSGGLQTLVIR | 22.91 | 1707.0043 | 18 | -0.9 | 570.0082 | 39.20 | 2 | 9113 | JVDM\_TOPE.raw | 1.2688E5 | 1 | 1 |  |  |  | PEAKS DB |
| VALEAC(+57.02)TQAR | 22.87 | 1117.5549 | 10 | 2.1 | 559.7859 | 12.79 | 2 | 1865 | JVDM\_TOPE.raw | 4.0142E4 | 1 | 1 |  | Carbamidomethylation | C6:Carbamidomethylation:1000.00 | PEAKS DB |
| TIVDDVTIAVEK | 22.82 | 1301.7078 | 12 | 2.0 | 651.8625 | 37.00 | 2 | 8528 | JVDM\_TOPE.raw | 5.5043E7 | 2 | 2 |  |  |  | PEAKS DB |
| ADETVAVEEVAAAEPASGKK | 22.75 | 1970.9796 | 20 | 2.1 | 658.0018 | 19.97 | 2 | 4172 | JVDM\_TOPE.raw | 7.7876E5 | 1 | 1 | tr\|A8HRZ0\|A8HRZ0\_CHLRE |  |  | PEAKS DB |
| LAATLAARGAAAASAAR | 22.60 | 1511.8531 | 17 | 3.3 | 756.9363 | 33.37 | 2 | 7620 | JVDM\_TOPE.raw | 2.323E4 | 1 | 1 |  |  |  | PEAKS DB |
| RTLDGMAAGSGGGGGGK | 22.55 | 1447.6837 | 17 | 5.6 | 724.8532 | 23.30 | 2 | 5217 | JVDM\_TOPE.raw | 0 | 0 | 0 |  |  |  | PEAKS DB |
| FLKPSVAGFLLQK | 22.52 | 1446.8599 | 13 | 2.2 | 483.2950 | 31.85 | 2 | 7252 | JVDM\_TOPE.raw | 4.5437E4 | 1 | 1 | tr\|A8JC04\|A8JC04\_CHLRE |  |  | PEAKS DB |
| PGGAASAAAAAGPAPAPAPR | 22.52 | 1627.8430 | 20 | -2.9 | 814.9264 | 31.63 | 2 | 7190 | JVDM\_TOPE.raw | 3.4957E5 | 1 | 1 |  |  |  | PEAKS DB |
| ALGYQGSWSAC(+57.02)K | 22.51 | 1326.6027 | 12 | 1.8 | 664.3098 | 19.45 | 2 | 4018 | JVDM\_TOPE.raw | 1.6304E5 | 1 | 1 | tr\|A0A2K3E094\|A0A2K3E094\_CHLRE | Carbamidomethylation | C11:Carbamidomethylation:1000.00 | PEAKS DB |
| VLLGPN(+.98)GVAK | 22.48 | 967.5702 | 10 | 2.2 | 484.7935 | 19.70 | 2 | 4216 | JVDM\_TOPE.raw | 8.4824E5 | 1 | 1 | tr\|P93106\|P93106\_CHLRE | Deamidation (NQ) | N6:Deamidation (NQ):1000.00 | PEAKS PTM |
| AGSAGVGAAEAAAAK | 22.38 | 1200.6099 | 15 | 2.7 | 601.3138 | 20.66 | 2 | 4398 | JVDM\_TOPE.raw | 3.4627E5 | 1 | 1 |  |  |  | PEAKS DB |
| AGGGGGGSVRAR | 22.32 | 1000.5162 | 12 | 3.7 | 501.2672 | 20.96 | 2 | 4497 | JVDM\_TOPE.raw | 1.3472E5 | 1 | 1 |  |  |  | PEAKS DB |
| AAGGGGGAGGGSVREGGGGGPGGLTHIK | 22.18 | 2247.1104 | 28 | -9.3 | 750.0371 | 27.47 | 2 | 6232 | JVDM\_TOPE.raw | 0 | 0 | 0 |  |  |  | PEAKS DB |
| QSAAAASAAPPPPPPR | 22.14 | 1484.7734 | 16 | 2.7 | 743.3960 | 25.69 | 2 | 5830 | JVDM\_TOPE.raw | 0 | 0 | 0 |  |  |  | PEAKS DB |
| GYSFTTTAER | 22.09 | 1131.5197 | 10 | 1.1 | 566.7677 | 16.01 | 2 | 2898 | JVDM\_TOPE.raw | 2.405E5 | 1 | 1 | tr\|A8JAV1\|A8JAV1\_CHLRE |  |  | PEAKS DB |
| AGGGGGGGGGGGGGGGAGANAVASTGQRR | 22.08 | 2184.0127 | 29 | 4.3 | 729.0146 | 20.43 | 2 | 4326 | JVDM\_TOPE.raw | 4.0097E5 | 1 | 1 |  |  |  | PEAKS DB |
| GSGGGGATGAGGASSSAGSAGGK | 22.01 | 1664.7350 | 23 | 8.9 | 833.3822 | 18.49 | 2 | 3742 | JVDM\_TOPE.raw | 6.326E3 | 1 | 1 |  |  |  | PEAKS DB |
| VEYTELQILC(+57.02)PQTIDSVTGYPMDD | 21.84 | 2786.2666 | 24 | 2.0 | 929.7647 | 39.70 | 2 | 9259 | JVDM\_TOPE.raw | 6.4405E4 | 1 | 1 | tr\|A0A2K3CQ54\|A0A2K3CQ54\_CHLRE | Carbamidomethylation | C10:Carbamidomethylation:1000.00 | PEAKS DB |
| DVAGHAPSRR | 21.83 | 1064.5475 | 10 | 4.1 | 533.2832 | 15.78 | 2 | 2850 | JVDM\_TOPE.raw | 3.8556E4 | 1 | 1 |  |  |  | PEAKS DB |
| TLSLTLK | 21.69 | 774.4851 | 7 | 2.3 | 388.2507 | 23.33 | 2 | 5221 | JVDM\_TOPE.raw | 2.692E6 | 1 | 1 |  |  |  | PEAKS DB |
| GTAAAAAAAAVAAGGSGAGGSAGASGGAGGGGGGGGFASR | 21.55 | 2904.3457 | 40 | 5.0 | 969.1274 | 32.93 | 2 | 7485 | JVDM\_TOPE.raw | 0 | 0 | 0 |  |  |  | PEAKS DB |
| EAPAPAPPGAKADGK | 21.49 | 1375.7095 | 15 | -6.0 | 688.8579 | 21.15 | 2 | 4514 | JVDM\_TOPE.raw | 3.2217E6 | 1 | 1 |  |  |  | PEAKS DB |
| LAASVLNC(+57.02)GLR | 21.49 | 1172.6335 | 11 | 5.2 | 587.3271 | 23.06 | 2 | 5147 | JVDM\_TOPE.raw | 0 | 0 | 0 |  | Carbamidomethylation | C8:Carbamidomethylation:1000.00 | PEAKS DB |
| AGVGAGGSGDVAAAGLPGR | 21.47 | 1538.7800 | 19 | -1.4 | 513.9332 | 22.34 | 2 | 4867 | JVDM\_TOPE.raw | 1.169E5 | 1 | 1 |  |  |  | PEAKS DB |
| DGGGGGGGVPIAVPPGR | 21.45 | 1418.7266 | 17 | 8.2 | 710.3764 | 18.81 | 2 | 3840 | JVDM\_TOPE.raw | 0 | 0 | 0 |  |  |  | PEAKS DB |
| FVDDDLR | 21.41 | 878.4133 | 7 | 0.5 | 440.2142 | 15.82 | 2 | 2905 | JVDM\_TOPE.raw | 1.1021E6 | 1 | 1 | PHL7 |  |  | PEAKS DB |
| AGGGAGGGGGAGGGGGAGPR | 21.35 | 1353.6133 | 20 | 1.5 | 677.8149 | 21.61 | 2 | 4711 | JVDM\_TOPE.raw | 8.3907E4 | 1 | 1 |  |  |  | PEAKS DB |
| KPAAAAPAPPK | 21.31 | 1017.5970 | 11 | -1.5 | 509.8051 | 28.66 | 2 | 6482 | JVDM\_TOPE.raw | 1.4915E5 | 1 | 1 |  |  |  | PEAKS DB |
| TAPAPGGGVGKGAVITAGGGGGGDGGK | 21.13 | 2122.0767 | 27 | 5.8 | 708.3702 | 29.25 | 2 | 6643 | JVDM\_TOPE.raw | 2.7808E5 | 1 | 1 |  |  |  | PEAKS DB |
| AASGAAAAGAAAGPVGTR | 21.09 | 1425.7324 | 18 | -1.4 | 713.8725 | 30.09 | 2 | 6850 | JVDM\_TOPE.raw | 0 | 0 | 0 |  |  |  | PEAKS DB |
| GTGGAAAPAAGAAAAPVAAAAP | 21.03 | 1660.8531 | 22 | -7.7 | 831.4275 | 28.65 | 2 | 6486 | JVDM\_TOPE.raw | 0 | 0 | 0 |  |  |  | PEAKS DB |
| IAGGGGGGGGNGSGSAAVK | 20.88 | 1429.6909 | 19 | -6.2 | 715.8483 | 14.87 | 2 | 2534 | JVDM\_TOPE.raw | 7.508E4 | 1 | 1 |  |  |  | PEAKS DB |
| AGGAGVTGGTSALGGAR | 20.88 | 1358.6902 | 17 | 2.2 | 680.3539 | 12.40 | 2 | 1442 | JVDM\_TOPE.raw | 0 | 0 | 0 |  |  |  | PEAKS DB |
| GDTTQPFTLTSLVDVGR | 20.88 | 1805.9159 | 17 | 5.8 | 903.9705 | 36.23 | 2 | 8353 | JVDM\_TOPE.raw | 1.1321E4 | 1 | 1 |  |  |  | PEAKS DB |
| AYPGQFR | 20.80 | 837.4133 | 7 | 0.4 | 419.7141 | 13.37 | 2 | 2237 | JVDM\_TOPE.raw | 4.4914E6 | 1 | 1 | tr\|A8J6Y8\|A8J6Y8\_CHLRE |  |  | PEAKS DB |
| MYEYVTEELPALLR | 20.73 | 1725.8647 | 14 | 6.9 | 863.9456 | 37.74 | 2 | 8753 | JVDM\_TOPE.raw | 0 | 0 | 0 | tr\|A8J7F8\|A8J7F8\_CHLRE:tr\|A0A2K3DW85\|A0A2K3DW85\_CHLRE:tr\|A0A2K3DW83\|A0A2K3DW83\_CHLRE |  |  | PEAKS DB |
| ALSGGAGGAVLLPGNGS | 20.72 | 1396.7310 | 17 | -2.0 | 699.3713 | 25.33 | 2 | 5759 | JVDM\_TOPE.raw | 4.0137E4 | 1 | 1 |  |  |  | PEAKS DB |
| GGGGDGKGPGGGGGGGGGRVGGVR | 20.58 | 1837.8890 | 24 | -7.0 | 919.9454 | 28.31 | 2 | 6416 | JVDM\_TOPE.raw | 0 | 0 | 0 |  |  |  | PEAKS DB |
| RGGGGLVGGGSGGAVGSSL | 20.57 | 1500.7644 | 19 | 2.0 | 751.3910 | 31.94 | 2 | 7260 | JVDM\_TOPE.raw | 3.4867E4 | 1 | 1 |  |  |  | PEAKS DB |
| GAAAAASGGGADELK | 20.43 | 1244.5996 | 15 | 2.3 | 623.3085 | 5.40 | 2 | 185 | JVDM\_TOPE.raw | 1.051E5 | 1 | 1 |  |  |  | PEAKS DB |
| DKGGGGAAAGGAAAGAAAGAAAGA | 20.15 | 1697.8080 | 24 | -0.7 | 849.9106 | 23.08 | 2 | 5154 | JVDM\_TOPE.raw | 0 | 0 | 0 |  |  |  | PEAKS DB |
| ELAAAGGGSSLQTK | 20.14 | 1288.6622 | 14 | 3.0 | 645.3403 | 18.16 | 2 | 3609 | JVDM\_TOPE.raw | 0 | 0 | 0 |  |  |  | PEAKS DB |
| GC(+57.02)EVLYIAQR | 20.10 | 1207.6019 | 10 | 5.1 | 604.8113 | 22.37 | 2 | 4922 | JVDM\_TOPE.raw | 3.2257E4 | 1 | 1 |  | Carbamidomethylation | C2:Carbamidomethylation:1000.00 | PEAKS DB |
| TTAAAGGGGGGGGGGGAGGAGGRK | 20.08 | 1714.8094 | 24 | -5.8 | 858.4070 | 30.14 | 2 | 6853 | JVDM\_TOPE.raw | 2.3544E5 | 1 | 1 |  |  |  | PEAKS DB |
| GAAPPPSLAPGGK | 20.06 | 1118.6084 | 13 | -4.0 | 560.3093 | 35.85 | 2 | 8231 | JVDM\_TOPE.raw | 4.6624E5 | 1 | 1 |  |  |  | PEAKS DB |
| GAPAVAAAAAAAAAGGGAGGGAGGAP | 20.06 | 1818.8972 | 26 | -0.1 | 607.3063 | 23.31 | 2 | 5187 | JVDM\_TOPE.raw | 5.7435E6 | 1 | 1 | tr\|A0A2K3DFK4\|A0A2K3DFK4\_CHLRE |  |  | PEAKS DB |
| LNMAPTIPAAR | 20.05 | 1153.6277 | 11 | 0.7 | 577.8215 | 22.54 | 2 | 4986 | JVDM\_TOPE.raw | 4.8937E4 | 1 | 1 |  |  |  | PEAKS DB |
| AVATGGAGGGAGGGAGGGVAGAR | 19.99 | 1654.8135 | 23 | 1.5 | 828.4153 | 28.39 | 2 | 6429 | JVDM\_TOPE.raw | 1.4381E5 | 1 | 1 |  |  |  | PEAKS DB |
| GGAASPGGTPAPPAAADGGGEAAAAA | 19.99 | 2018.9292 | 26 | -4.4 | 673.9807 | 18.82 | 2 | 3843 | JVDM\_TOPE.raw | 0 | 0 | 0 |  |  |  | PEAKS DB |
| VGAAAGGAGAGAGVAHHGAQ | 19.98 | 1585.7709 | 20 | -5.7 | 793.8882 | 17.02 | 2 | 3281 | JVDM\_TOPE.raw | 2.2574E6 | 1 | 1 |  |  |  | PEAKS DB |
| GAGAGVGIAGGAGGGGGGGGAGARVA | 19.95 | 1837.9142 | 26 | -2.4 | 919.9622 | 29.06 | 2 | 6813 | JVDM\_TOPE.raw | 1.239E6 | 1 | 1 |  |  |  | PEAKS DB |
| SAGGSGGGGGSGAAAGGGGSR | 19.90 | 1490.6458 | 21 | 2.1 | 746.3317 | 16.48 | 2 | 3063 | JVDM\_TOPE.raw | 0 | 0 | 0 |  |  |  | PEAKS DB |
| AALTGGGVTMTSGGNHSRGGFSDDTAR | 19.90 | 2579.1782 | 27 | -5.8 | 860.7283 | 21.31 | 2 | 4620 | JVDM\_TOPE.raw | 0 | 0 | 0 |  |  |  | PEAKS DB |
| SSGTFAVK | 19.90 | 795.4127 | 8 | 1.3 | 398.7141 | 12.91 | 2 | 1946 | JVDM\_TOPE.raw | 9.2055E4 | 1 | 1 | tr\|A0A2K3CWZ6\|A0A2K3CWZ6\_CHLRE |  |  | PEAKS DB |
| AGPGGGGGGAGAVPARPPPHSGAKRR | 19.89 | 2291.2107 | 26 | 9.1 | 1146.6230 | 35.75 | 2 | 8220 | JVDM\_TOPE.raw | 5.0709E4 | 1 | 1 |  |  |  | PEAKS DB |
| LGVGGGGAAGPAAKRR | 19.76 | 1393.7902 | 16 | -7.0 | 697.8975 | 39.40 | 2 | 9208 | JVDM\_TOPE.raw | 4.7855E4 | 1 | 1 |  |  |  | PEAKS DB |
| GAIGLGPPAAGGGGTGVGGK | 19.75 | 1549.8212 | 20 | -1.3 | 775.9169 | 26.35 | 2 | 5989 | JVDM\_TOPE.raw | 2.0121E4 | 1 | 1 |  |  |  | PEAKS DB |
| ASAAGAGAVPAK | 19.72 | 969.5243 | 12 | 1.2 | 485.7700 | 13.26 | 2 | 2097 | JVDM\_TOPE.raw | 1.136E5 | 1 | 1 |  |  |  | PEAKS DB |
| RKVGGGGGGGGAGGGGGHK | 19.72 | 1478.7450 | 19 | -8.5 | 740.3735 | 24.82 | 2 | 5641 | JVDM\_TOPE.raw | 2.7297E4 | 1 | 1 |  |  |  | PEAKS DB |
| ARGLAAPASGGGSGGGGR | 19.71 | 1454.7338 | 18 | -7.4 | 728.3688 | 15.14 | 2 | 2656 | JVDM\_TOPE.raw | 0 | 0 | 0 |  |  |  | PEAKS DB |
| AAAAGSSAGGGAAAC(+57.02)GGGGGGK | 19.67 | 1618.7117 | 22 | 9.3 | 810.3707 | 20.57 | 2 | 4369 | JVDM\_TOPE.raw | 1.058E5 | 1 | 1 |  | Carbamidomethylation | C15:Carbamidomethylation:1000.00 | PEAKS DB |
| SPPAGANGAAVSPSGGGAAPAAVGTR | 19.65 | 2147.0718 | 26 | 7.7 | 1074.5514 | 11.38 | 2 | 1131 | JVDM\_TOPE.raw | 0 | 0 | 0 |  |  |  | PEAKS DB |
| SGAGGGGGPGITSAALEAGLGAR | 19.60 | 1882.9496 | 23 | -5.7 | 628.6536 | 27.21 | 2 | 6174 | JVDM\_TOPE.raw | 0 | 0 | 0 |  |  |  | PEAKS DB |
| TEDVVFVQTTSADVAK | 19.58 | 1708.8519 | 16 | 0.1 | 855.4333 | 23.91 | 2 | 5371 | JVDM\_TOPE.raw | 0 | 0 | 0 | tr\|O48949\|O48949\_CHLRE |  |  | PEAKS DB |
| GPNDGGFGGGIKAAAGGVAAGAGGR | 19.57 | 2041.0089 | 25 | -2.4 | 681.3419 | 25.09 | 2 | 5706 | JVDM\_TOPE.raw | 1.7027E5 | 1 | 1 |  |  |  | PEAKS DB |
| GAGAGFSAAASAAAAS | 19.57 | 1236.5734 | 16 | 5.9 | 619.2976 | 18.66 | 2 | 3752 | JVDM\_TOPE.raw | 1.6301E5 | 1 | 1 |  |  |  | PEAKS DB |
| QYTPTQALVAR | 19.54 | 1246.6670 | 11 | 2.0 | 624.3420 | 19.04 | 2 | 3922 | JVDM\_TOPE.raw | 0 | 0 | 0 |  |  |  | PEAKS DB |
| AGTAAAAAGGGAAAK | 19.36 | 1114.5730 | 15 | -5.7 | 558.2906 | 16.36 | 2 | 3005 | JVDM\_TOPE.raw | 3.1712E4 | 1 | 1 |  |  |  | PEAKS DB |
| RAGGGSKAAVSR | 19.33 | 1115.6160 | 12 | 1.5 | 558.8161 | 19.92 | 2 | 4211 | JVDM\_TOPE.raw | 0 | 0 | 0 |  |  |  | PEAKS DB |
| GADSITPGGSAGGAAAAAAAAAAAAGTAT | 19.31 | 2228.0669 | 29 | 8.1 | 743.7023 | 26.19 | 2 | 5962 | JVDM\_TOPE.raw | 0 | 0 | 0 |  |  |  | PEAKS DB |
| EAAAMAAAAASR | 19.29 | 1089.5237 | 12 | 1.4 | 545.7699 | 13.26 | 2 | 2082 | JVDM\_TOPE.raw | 0 | 0 | 0 |  |  |  | PEAKS DB |
| RAVAAAAAGSGAGGGGGGK | 19.23 | 1441.7385 | 19 | 7.5 | 721.8820 | 37.64 | 2 | 8730 | JVDM\_TOPE.raw | 1.665E5 | 1 | 1 |  |  |  | PEAKS DB |
| QATSQAHAAAAATGTSGSA | 19.20 | 1657.7656 | 19 | 5.0 | 829.8942 | 22.54 | 2 | 5017 | JVDM\_TOPE.raw | 1.8368E5 | 1 | 1 |  |  |  | PEAKS DB |
| AVVAAAAAAAAALSPAGGGGGGGGGGGGGGGDSP | 19.19 | 2449.1582 | 34 | 7.0 | 817.3990 | 23.92 | 2 | 5380 | JVDM\_TOPE.raw | 5.9744E4 | 1 | 1 |  |  |  | PEAKS DB |
| LLVAGGPGEVGR | 19.18 | 1123.6349 | 12 | 2.2 | 562.8259 | 19.82 | 2 | 4137 | JVDM\_TOPE.raw | 1.0179E6 | 1 | 1 |  |  |  | PEAKS DB |
| GMEVVDTGKPLSVPVGK | 19.17 | 1711.9178 | 17 | 2.4 | 571.6479 | 23.60 | 2 | 5311 | JVDM\_TOPE.raw | 3.5529E4 | 1 | 1 | P06541\|ATPB\_CHLRE |  |  | PEAKS DB |
| LSLGAAGAAAGAGG | 19.07 | 1042.5406 | 14 | 2.1 | 522.2787 | 19.71 | 2 | 4091 | JVDM\_TOPE.raw | 4.6384E5 | 1 | 1 |  |  |  | PEAKS DB |
| QLLGGGSDAGGEVAGEEEEDTELVAAVAVEVGRR | 19.02 | 3368.6331 | 34 | 3.8 | 843.1687 | 27.38 | 2 | 6210 | JVDM\_TOPE.raw | 0 | 0 | 0 |  |  |  | PEAKS DB |
| VNLVC(+57.02)VGR | 19.00 | 915.4960 | 8 | -0.6 | 458.7550 | 18.77 | 2 | 3813 | JVDM\_TOPE.raw | 1.6723E5 | 1 | 1 | tr\|A8HXL8\|A8HXL8\_CHLRE | Carbamidomethylation | C5:Carbamidomethylation:1000.00 | PEAKS DB |
| YPNAAAGGAPGAAAAAAGHHPLQDGPDLPPR | 18.93 | 2887.4114 | 31 | -8.7 | 963.4694 | 35.80 | 2 | 8234 | JVDM\_TOPE.raw | 4.6847E5 | 1 | 1 |  |  |  | PEAKS DB |
| GGSGSSGSSSTAGK | 18.84 | 1125.4897 | 14 | 5.9 | 563.7555 | 15.89 | 2 | 2862 | JVDM\_TOPE.raw | 1.0696E5 | 1 | 1 |  |  |  | PEAKS DB |
| GTGASDAAAGAAANGASSR | 18.83 | 1561.7080 | 19 | 8.4 | 781.8679 | 19.85 | 2 | 4186 | JVDM\_TOPE.raw | 0 | 0 | 0 |  |  |  | PEAKS DB |
| AGGGAGGPGGGGGGGAAGAAAVGAGGV | 18.81 | 1793.8405 | 27 | 3.7 | 897.9308 | 14.03 | 2 | 2256 | JVDM\_TOPE.raw | 2.3574E5 | 1 | 1 |  |  |  | PEAKS DB |
| SLGGGGGGGGGGSRSL | 18.72 | 1231.5905 | 16 | 2.8 | 616.8042 | 12.67 | 2 | 1749 | JVDM\_TOPE.raw | 6.4365E5 | 1 | 1 |  |  |  | PEAKS DB |
| RGGGGGGGGGGGLGLIR | 18.70 | 1353.7225 | 17 | 6.6 | 677.8730 | 5.25 | 2 | 52 | JVDM\_TOPE.raw | 0 | 0 | 0 |  |  |  | PEAKS DB |
| FPTAALGSVI | 18.69 | 974.5436 | 10 | -2.5 | 488.2779 | 12.71 | 2 | 1759 | JVDM\_TOPE.raw | 0 | 0 | 0 |  |  |  | PEAKS DB |
| TGALIDAAVSALPGR | 18.65 | 1410.7831 | 15 | -5.9 | 706.3947 | 18.99 | 2 | 3902 | JVDM\_TOPE.raw | 0 | 0 | 0 |  |  |  | PEAKS DB |
| IQDKEGIPPDQQR | 18.64 | 1522.7739 | 13 | 2.4 | 508.5998 | 12.45 | 2 | 1393 | JVDM\_TOPE.raw | 1.2856E6 | 1 | 1 |  |  |  | PEAKS DB |
| LSVAAAASGSK | 18.61 | 960.5240 | 11 | 1.1 | 481.2698 | 18.08 | 2 | 3577 | JVDM\_TOPE.raw | 0 | 0 | 0 |  |  |  | PEAKS DB |
| SVAAPGSAGGGGDR | 18.54 | 1157.5425 | 14 | 6.4 | 579.7822 | 12.86 | 2 | 1914 | JVDM\_TOPE.raw | 2.6731E5 | 1 | 1 |  |  |  | PEAKS DB |
| GSVAVGDEAEAVRAE | 18.47 | 1458.6949 | 15 | 4.2 | 730.3578 | 15.86 | 2 | 2865 | JVDM\_TOPE.raw | 0 | 0 | 0 |  |  |  | PEAKS DB |
| TLDTAAAAAAAAADTK | 18.45 | 1431.7205 | 16 | 5.2 | 716.8712 | 27.45 | 2 | 6216 | JVDM\_TOPE.raw | 2.2578E5 | 1 | 1 |  |  |  | PEAKS DB |
| PGGSGAAGPAVGADTSAHR | 18.45 | 1634.7760 | 19 | -1.7 | 818.3939 | 21.90 | 2 | 4801 | JVDM\_TOPE.raw | 0 | 0 | 0 |  |  |  | PEAKS DB |
| RGGGGGGGAAGGGVVAV | 18.38 | 1254.6428 | 17 | -7.3 | 628.3241 | 20.31 | 2 | 4354 | JVDM\_TOPE.raw | 7.3301E4 | 1 | 1 |  |  |  | PEAKS DB |
| IPDVAGGGGGGAGGGGGAGGGGGR | 18.33 | 1723.7986 | 24 | -4.8 | 862.9025 | 33.50 | 2 | 7653 | JVDM\_TOPE.raw | 0 | 0 | 0 |  |  |  | PEAKS DB |
| GAASVLVGGGGAGGGGGGGGGGGGGGAGGGGGGGGGGAR | 18.33 | 2581.1726 | 39 | -2.6 | 861.3959 | 20.20 | 2 | 4284 | JVDM\_TOPE.raw | 0 | 0 | 0 |  |  |  | PEAKS DB |
| MAGGRSGGGAAGAPGR | 18.28 | 1328.6367 | 16 | -1.2 | 665.3248 | 18.44 | 2 | 3677 | JVDM\_TOPE.raw | 5.0808E5 | 1 | 1 |  |  |  | PEAKS DB |
| GAGAGGGGGGGGGAAGGGRGPGGG | 18.28 | 1581.6992 | 24 | 0.7 | 791.8574 | 16.93 | 2 | 3197 | JVDM\_TOPE.raw | 1.4393E4 | 1 | 1 |  |  |  | PEAKS DB |
| VAEAAAAGAAHAAAAGGNAVK | 18.27 | 1747.8965 | 21 | 7.1 | 874.9617 | 22.34 | 2 | 4923 | JVDM\_TOPE.raw | 0 | 0 | 0 |  |  |  | PEAKS DB |
| AVAGGGGGGGGPLAGHELL | 18.26 | 1545.7899 | 19 | -5.4 | 773.8981 | 22.43 | 2 | 4940 | JVDM\_TOPE.raw | 1.3333E5 | 1 | 1 |  |  |  | PEAKS DB |
| LLSGGSAAAAAAAAAFGGSK | 18.20 | 1647.8579 | 20 | 5.5 | 550.2963 | 25.17 | 2 | 5727 | JVDM\_TOPE.raw | 4.0697E4 | 1 | 1 |  |  |  | PEAKS DB |
| GAGVGIAGGAGGGGGGGGAGAR | 18.19 | 1539.7501 | 22 | 2.6 | 770.8843 | 19.19 | 2 | 3970 | JVDM\_TOPE.raw | 0 | 0 | 0 |  |  |  | PEAKS DB |
| GAAAASLAPMGSDVSGLGGPTPSSLGR | 18.14 | 2383.1802 | 27 | -7.1 | 795.3950 | 40.31 | 2 | 9394 | JVDM\_TOPE.raw | 3.1375E3 | 1 | 1 |  |  |  | PEAKS DB |
| LAGDGGAGSGSGGGGGGGR | 18.14 | 1402.6185 | 19 | 3.8 | 702.3192 | 16.19 | 2 | 3000 | JVDM\_TOPE.raw | 1.2654E5 | 1 | 1 |  |  |  | PEAKS DB |
| GAGYDAASPSALQAAESR | 18.09 | 1720.8015 | 18 | 3.9 | 861.4114 | 23.64 | 2 | 5307 | JVDM\_TOPE.raw | 0 | 0 | 0 |  |  |  | PEAKS DB |
| KGGGGSGGAASASERAR | 18.02 | 1474.7236 | 17 | 2.8 | 738.3712 | 12.56 | 2 | 1565 | JVDM\_TOPE.raw | 0 | 0 | 0 |  |  |  | PEAKS DB |
| GAKGSASAAGAGGR | 18.01 | 1116.5635 | 14 | 2.8 | 559.2906 | 14.80 | 2 | 2537 | JVDM\_TOPE.raw | 0 | 0 | 0 |  |  |  | PEAKS DB |
| AGGGGGPPAAKR | 17.89 | 994.5308 | 12 | 4.8 | 498.2750 | 17.48 | 2 | 3453 | JVDM\_TOPE.raw | 1.1617E6 | 1 | 1 |  |  |  | PEAKS DB |
| LTYYTPDYVVR | 17.79 | 1388.6976 | 11 | 2.7 | 695.3580 | 25.19 | 2 | 5716 | JVDM\_TOPE.raw | 8.9301E4 | 1 | 1 |  |  |  | PEAKS DB |
| GAATASSASGRPGGARSGQQSRHQK | 17.75 | 2409.1970 | 25 | -4.9 | 804.0690 | 27.11 | 2 | 6162 | JVDM\_TOPE.raw | 1.3512E4 | 1 | 1 |  |  |  | PEAKS DB |
| AVGAAAEAAPAAVPAGPEAGTTGR | 17.74 | 2062.0442 | 24 | -2.7 | 688.3535 | 26.87 | 2 | 6104 | JVDM\_TOPE.raw | 0 | 0 | 0 |  |  |  | PEAKS DB |
| LRHSDGGGSGGGAGGPPAAR | 17.70 | 1732.8353 | 20 | 6.0 | 867.4302 | 26.48 | 2 | 6022 | JVDM\_TOPE.raw | 3.2055E4 | 1 | 1 |  |  |  | PEAKS DB |
| GLNYEEWVEGLK | 17.66 | 1435.6982 | 12 | 2.1 | 718.8579 | 32.93 | 2 | 7460 | JVDM\_TOPE.raw | 7.2447E5 | 1 | 1 | tr\|A8J6Y8\|A8J6Y8\_CHLRE |  |  | PEAKS DB |
| KGTKGVEPK | 17.66 | 942.5498 | 9 | 1.3 | 472.2828 | 12.87 | 2 | 1904 | JVDM\_TOPE.raw | 0 | 0 | 0 |  |  |  | PEAKS DB |
| LAEEEDKLNNK | 17.63 | 1301.6462 | 11 | -4.9 | 651.8272 | 18.06 | 2 | 3565 | JVDM\_TOPE.raw | 0 | 0 | 0 |  |  |  | PEAKS DB |
| RAPGAAGAAGGGGGGGGVGGGGGGGGGASGQDPGQLSR | 17.63 | 2905.3523 | 38 | 5.0 | 969.4629 | 34.09 | 2 | 7821 | JVDM\_TOPE.raw | 2.3318E5 | 1 | 1 |  |  |  | PEAKS DB |
| EPAAAAGSGGGGGGGGGGER | 17.55 | 1527.6661 | 20 | 3.8 | 764.8433 | 13.95 | 2 | 2355 | JVDM\_TOPE.raw | 1.6813E5 | 1 | 1 |  |  |  | PEAKS DB |
| SAGAGAGGGAGGGAPLTPGQAAVAAYR | 17.55 | 2212.0984 | 27 | -0.8 | 1107.0555 | 27.86 | 2 | 6311 | JVDM\_TOPE.raw | 9.5433E4 | 1 | 1 |  |  |  | PEAKS DB |
| AAGGGGGAAAALAGGPPQ | 17.46 | 1349.6687 | 18 | 9.7 | 675.8481 | 12.67 | 2 | 1682 | JVDM\_TOPE.raw | 0 | 0 | 0 |  |  |  | PEAKS DB |
| RGGAAGGDSR | 17.44 | 902.4318 | 10 | -7.1 | 452.2200 | 13.16 | 2 | 2059 | JVDM\_TOPE.raw | 8.0046E4 | 1 | 1 |  |  |  | PEAKS DB |
| GVAAGYAASLQLDASGDR | 17.37 | 1720.8380 | 18 | -7.3 | 861.4200 | 23.48 | 2 | 5256 | JVDM\_TOPE.raw | 7.8373E4 | 1 | 1 |  |  |  | PEAKS DB |
| MGGGGGGGGGGGGGGTLRKGGALYSKPAPVQR | 17.35 | 2770.4045 | 32 | 9.4 | 924.4841 | 35.97 | 2 | 8276 | JVDM\_TOPE.raw | 0 | 0 | 0 |  |  |  | PEAKS DB |
| GGAGAAGAAAGGVVSPKGGAG | 17.33 | 1538.7800 | 21 | -1.4 | 513.9332 | 22.34 | 2 | 4967 | JVDM\_TOPE.raw | 1.169E5 | 1 | 1 |  |  |  | PEAKS DB |
| KAAAAGATADGATADGGAT | 17.33 | 1546.7223 | 19 | 2.4 | 774.3702 | 29.63 | 2 | 6734 | JVDM\_TOPE.raw | 4.9206E4 | 1 | 1 |  |  |  | PEAKS DB |
| EAAGLSTAAAGAAAGGAGGGR | 17.31 | 1642.8022 | 21 | 5.4 | 548.6110 | 18.09 | 2 | 3582 | JVDM\_TOPE.raw | 0 | 0 | 0 |  |  |  | PEAKS DB |
| TQIC(+57.02)YGSIGEVR | 17.25 | 1381.6660 | 12 | 3.3 | 691.8426 | 21.04 | 2 | 4533 | JVDM\_TOPE.raw | 6.2336E4 | 1 | 1 |  | Carbamidomethylation | C4:Carbamidomethylation:1000.00 | PEAKS DB |
| HGAGAGAAAGGGWAAGLAALQGGGG | 17.22 | 1961.9456 | 25 | 3.4 | 981.9834 | 27.00 | 2 | 6138 | JVDM\_TOPE.raw | 5.6067E4 | 1 | 1 |  |  |  | PEAKS DB |
| EGGGGGGGGGGSGSGKGKAKPVR | 17.22 | 1869.9404 | 23 | 1.7 | 935.9791 | 15.12 | 2 | 2652 | JVDM\_TOPE.raw | 0 | 0 | 0 |  |  |  | PEAKS DB |
| KAADAAAAAAAAKPAATPEPSKPAAK | 17.18 | 2345.2703 | 26 | 6.1 | 587.3284 | 24.73 | 2 | 5610 | JVDM\_TOPE.raw | 0 | 0 | 0 |  |  |  | PEAKS DB |
| TGLVGPTVGHLADK | 17.18 | 1363.7460 | 14 | 1.7 | 682.8814 | 24.44 | 2 | 5508 | JVDM\_TOPE.raw | 3.5098E5 | 1 | 1 |  |  |  | PEAKS DB |
| GTYGISEHDPLSPDASAYGAGSR | 17.17 | 2307.0403 | 23 | 7.5 | 770.0265 | 18.51 | 2 | 3736 | JVDM\_TOPE.raw | 0 | 0 | 0 |  |  |  | PEAKS DB |
| SGIGGGAGGGGAEGGGGGDAAASGGSSGSR | 17.16 | 2233.9543 | 30 | 4.7 | 745.6622 | 17.49 | 2 | 3403 | JVDM\_TOPE.raw | 4.3364E4 | 1 | 1 |  |  |  | PEAKS DB |
| REEAAAAAAALLSKSR | 17.14 | 1613.8849 | 16 | 5.8 | 807.9544 | 23.96 | 2 | 5381 | JVDM\_TOPE.raw | 0 | 0 | 0 |  |  |  | PEAKS DB |
| GVFTNVTSPTTK | 17.14 | 1250.6506 | 12 | 1.4 | 626.3335 | 19.58 | 2 | 4065 | JVDM\_TOPE.raw | 5.7418E4 | 1 | 1 |  |  |  | PEAKS DB |
| ATAVLATDIDTDSLTAFVK | 17.12 | 1951.0149 | 19 | 2.1 | 976.5167 | 35.63 | 2 | 8183 | JVDM\_TOPE.raw | 1.5207E4 | 1 | 1 | tr\|O48949\|O48949\_CHLRE |  |  | PEAKS DB |
| LSSGAAAGAAAATGVR | 17.08 | 1329.7001 | 16 | -7.2 | 665.8525 | 21.88 | 2 | 4800 | JVDM\_TOPE.raw | 2.8708E4 | 1 | 1 |  |  |  | PEAKS DB |
| VMVIC(+57.02)SNAK | 17.08 | 1020.5096 | 9 | 3.4 | 511.2639 | 14.95 | 2 | 2592 | JVDM\_TOPE.raw | 2.5016E4 | 1 | 1 |  | Carbamidomethylation | C5:Carbamidomethylation:1000.00 | PEAKS DB |
| GGVGGGGGGGGVGSTGGAAVGPGAGAGPGGGGGGGAGK | 17.03 | 2606.2180 | 38 | -5.5 | 869.7418 | 27.70 | 2 | 6276 | JVDM\_TOPE.raw | 0 | 0 | 0 |  |  |  | PEAKS DB |
| YLNIISK | 17.03 | 849.4960 | 7 | 4.7 | 425.7573 | 21.40 | 2 | 4622 | JVDM\_TOPE.raw | 2.8893E5 | 1 | 1 | Q08365\|RR3\_CHLRE |  |  | PEAKS DB |
| ALEGEIYDTFK | 17.03 | 1284.6238 | 11 | 2.4 | 643.3207 | 27.26 | 2 | 6186 | JVDM\_TOPE.raw | 5.6701E4 | 1 | 1 | tr\|O48949\|O48949\_CHLRE |  |  | PEAKS DB |
| ATLAITDDTR | 17.01 | 1075.5509 | 10 | 0.7 | 538.7831 | 16.92 | 2 | 3208 | JVDM\_TOPE.raw | 5.2825E4 | 1 | 1 | tr\|A8JC04\|A8JC04\_CHLRE |  |  | PEAKS DB |
| ELTEADGFVFGFPTR | 16.91 | 1684.8096 | 15 | 3.7 | 843.4152 | 36.06 | 2 | 8299 | JVDM\_TOPE.raw | 3.1435E4 | 1 | 1 |  |  |  | PEAKS DB |
| LLAGGGGSGSR | 16.90 | 930.4883 | 11 | 4.9 | 466.2537 | 23.77 | 2 | 5337 | JVDM\_TOPE.raw | 4.1031E4 | 1 | 1 |  |  |  | PEAKS DB |
| LGGGGGGAAAAGPGGVQQS | 16.85 | 1467.7065 | 19 | -8.9 | 734.8540 | 30.03 | 2 | 6849 | JVDM\_TOPE.raw | 3.523E4 | 1 | 1 |  |  |  | PEAKS DB |
| AVASGAGSAAGGTPK | 16.83 | 1200.6099 | 15 | 2.7 | 601.3138 | 20.66 | 2 | 4474 | JVDM\_TOPE.raw | 3.4627E5 | 1 | 1 |  |  |  | PEAKS DB |
| RSDGGSLAGNGSTAGSK | 16.82 | 1520.7179 | 17 | -0.4 | 761.3659 | 28.49 | 2 | 6452 | JVDM\_TOPE.raw | 0 | 0 | 0 |  |  |  | PEAKS DB |
| GAPGGAGPDGDK | 16.75 | 997.4464 | 12 | 0.3 | 499.7307 | 14.82 | 2 | 2557 | JVDM\_TOPE.raw | 9.3454E3 | 1 | 1 |  |  |  | PEAKS DB |
| GAGGAVAAAPGGASAAATAAV | 16.75 | 1567.7954 | 21 | 0.8 | 784.9056 | 26.99 | 2 | 6124 | JVDM\_TOPE.raw | 0 | 0 | 0 |  |  |  | PEAKS DB |
| ARAATGGGGGGGNGGVR | 16.73 | 1370.6763 | 17 | 5.1 | 686.3489 | 21.83 | 2 | 4768 | JVDM\_TOPE.raw | 9.3266E4 | 1 | 1 |  |  |  | PEAKS DB |
| WLAGEDGATGGRR | 16.62 | 1344.6534 | 13 | 9.1 | 673.3401 | 21.61 | 2 | 4701 | JVDM\_TOPE.raw | 5.3961E4 | 1 | 1 |  |  |  | PEAKS DB |
| AGATGGRGAGVGPAGR | 16.60 | 1310.6803 | 16 | -3.5 | 437.8992 | 38.23 | 2 | 8866 | JVDM\_TOPE.raw | 2.2293E5 | 1 | 1 |  |  |  | PEAKS DB |
| MGAVATASGGAAT | 16.57 | 1063.4968 | 13 | -3.6 | 532.7538 | 13.91 | 2 | 2282 | JVDM\_TOPE.raw | 0 | 0 | 0 |  |  |  | PEAKS DB |
| LTAAGEYGYYC(+57.02)EPHQGAGMVGK | 16.52 | 2358.0408 | 22 | 2.8 | 787.0231 | 20.00 | 2 | 4221 | JVDM\_TOPE.raw | 1.8245E4 | 1 | 1 |  | Carbamidomethylation | C11:Carbamidomethylation:1000.00 | PEAKS DB |
| DAPGC(+57.02)SARGGHYPTK | 16.51 | 1572.7103 | 15 | -3.7 | 787.3595 | 19.18 | 2 | 3963 | JVDM\_TOPE.raw | 3.4847E4 | 1 | 1 |  | Carbamidomethylation | C5:Carbamidomethylation:1000.00 | PEAKS DB |
| EAVGGVIVALNA | 16.50 | 1111.6237 | 12 | 3.1 | 556.8208 | 19.01 | 2 | 3906 | JVDM\_TOPE.raw | 2.2824E4 | 1 | 1 |  |  |  | PEAKS DB |
| AGGAAGGGGGGDGSAGGGGGGSAR | 16.50 | 1673.7101 | 24 | 7.4 | 837.8685 | 20.28 | 2 | 4355 | JVDM\_TOPE.raw | 6.1613E5 | 1 | 1 |  |  |  | PEAKS DB |
| AQAPGGPGGGGGPGPGGGGGGGGGGGGGGGAGR | 16.50 | 2272.0076 | 33 | 7.7 | 758.3489 | 17.99 | 2 | 3542 | JVDM\_TOPE.raw | 0 | 0 | 0 |  |  |  | PEAKS DB |
| ASGGAGGAAAAADGVAVPDGGERR | 16.49 | 2038.9779 | 24 | -5.9 | 680.6625 | 23.96 | 2 | 5400 | JVDM\_TOPE.raw | 4.4935E4 | 1 | 1 |  |  |  | PEAKS DB |
| ATGAAGAAESGLAGK | 16.49 | 1230.6204 | 15 | 2.8 | 616.3192 | 18.29 | 2 | 3683 | JVDM\_TOPE.raw | 3.1619E5 | 1 | 1 |  |  |  | PEAKS DB |
| QINTSARAAR | 16.41 | 1086.5894 | 10 | 5.1 | 544.3047 | 25.12 | 2 | 5703 | JVDM\_TOPE.raw | 0 | 0 | 0 |  |  |  | PEAKS DB |
| GDAAGPGTGASAAMDAEK | 16.41 | 1575.6835 | 18 | 6.1 | 526.2383 | 22.90 | 2 | 5098 | JVDM\_TOPE.raw | 1.4087E5 | 1 | 1 |  |  |  | PEAKS DB |
| DGGGGGGGTSGAK | 16.39 | 976.4210 | 13 | -5.0 | 489.2153 | 12.81 | 2 | 1850 | JVDM\_TOPE.raw | 0 | 0 | 0 |  |  |  | PEAKS DB |
| GEANWYGGAAQGGGGAGVGPRGTGAR | 16.36 | 2330.0901 | 26 | 5.5 | 777.7083 | 35.02 | 2 | 8023 | JVDM\_TOPE.raw | 9.2289E5 | 1 | 1 |  |  |  | PEAKS DB |
| AGASAAAAGAVFGEAASGGAK | 16.36 | 1690.8274 | 21 | -3.8 | 564.6143 | 17.43 | 2 | 3367 | JVDM\_TOPE.raw | 9.7866E3 | 1 | 1 |  |  |  | PEAKS DB |
| ESVLLPSASGSLSASA | 16.36 | 1474.7515 | 16 | -5.5 | 738.3790 | 30.05 | 2 | 6862 | JVDM\_TOPE.raw | 3.8501E4 | 1 | 1 |  |  |  | PEAKS DB |
| AGAAASAAQSSAAAAVEKASG | 16.34 | 1745.8544 | 21 | 9.5 | 873.9427 | 26.96 | 2 | 6120 | JVDM\_TOPE.raw | 0 | 0 | 0 | tr\|A0A2K3E7M7\|A0A2K3E7M7\_CHLRE |  |  | PEAKS DB |
| EANGSGGVVGK | 16.30 | 973.4828 | 11 | 2.4 | 487.7499 | 14.37 | 2 | 2407 | JVDM\_TOPE.raw | 1.0087E5 | 1 | 1 |  |  |  | PEAKS DB |
| GPGPAGPPSALGSAVQAQLR | 16.30 | 1829.9747 | 20 | -0.4 | 915.9943 | 25.83 | 2 | 5859 | JVDM\_TOPE.raw | 0 | 0 | 0 |  |  |  | PEAKS DB |
| GGPAAAGAYVK | 16.30 | 960.5028 | 11 | 5.0 | 481.2611 | 24.50 | 2 | 5543 | JVDM\_TOPE.raw | 0 | 0 | 0 |  |  |  | PEAKS DB |
| EGARVPGGGGGGVR | 16.29 | 1224.6323 | 14 | 7.9 | 409.2213 | 12.29 | 2 | 1355 | JVDM\_TOPE.raw | 0 | 0 | 0 |  |  |  | PEAKS DB |
| GSGGGSGGGGSGGAAF | 16.28 | 1138.4639 | 16 | 4.4 | 570.2417 | 12.45 | 2 | 1480 | JVDM\_TOPE.raw | 0 | 0 | 0 |  |  |  | PEAKS DB |
| AAAAAASKAAELAAGDEHASVDGDAAGGGGAAAPK | 16.27 | 2976.4172 | 35 | -3.4 | 993.1430 | 25.79 | 2 | 5849 | JVDM\_TOPE.raw | 0 | 0 | 0 |  |  |  | PEAKS DB |
| AGK(+14.02)GSATLSMAYAAAR | 16.27 | 1538.7875 | 16 | -6.3 | 513.9332 | 16.59 | 2 | 3033 | JVDM\_TOPE.raw | 4.0015E6 | 1 | 1 | tr\|P93106\|P93106\_CHLRE | Methylation(KR) | K3:Methylation(KR):16.42 | PEAKS PTM |
| ASAAAGGGGPLLPAASAH | 16.25 | 1474.7528 | 18 | -6.5 | 738.3789 | 29.50 | 2 | 6712 | JVDM\_TOPE.raw | 0 | 0 | 0 |  |  |  | PEAKS DB |
| AAAGGGGGAGGGSSAVTGGAGPHGHR | 16.25 | 2029.9426 | 26 | 9.8 | 677.6614 | 24.92 | 2 | 5660 | JVDM\_TOPE.raw | 7.1374E3 | 1 | 1 |  |  |  | PEAKS DB |
| TKGEAGTGNVVEAVR(+14.02) | 16.24 | 1500.7896 | 15 | 5.2 | 751.4059 | 30.48 | 2 | 6951 | JVDM\_TOPE.raw | 1.6497E5 | 1 | 1 | tr\|A0A2K3CWZ6\|A0A2K3CWZ6\_CHLRE | Methylation(KR) | R15:Methylation(KR):57.37 | PEAKS PTM |
| EAAAKAEAALATAR | 16.20 | 1342.7205 | 14 | 6.3 | 672.3718 | 28.42 | 2 | 6438 | JVDM\_TOPE.raw | 0 | 0 | 0 |  |  |  | PEAKS DB |
| QSGGGGGGGGGGGSGGGGAGGPKGGDSGGGK | 16.19 | 2200.9441 | 31 | 4.1 | 1101.4839 | 12.56 | 2 | 1562 | JVDM\_TOPE.raw | 0 | 0 | 0 |  |  |  | PEAKS DB |
| GAGGGGGGSAAAARPLLSAALAAR | 16.10 | 1979.0659 | 24 | 5.5 | 660.6995 | 30.47 | 2 | 6966 | JVDM\_TOPE.raw | 9.0272E4 | 1 | 1 |  |  |  | PEAKS DB |
| RHDGGGGGGAAAAGAGVEAAA | 16.09 | 1678.7771 | 21 | 6.0 | 840.4009 | 30.39 | 2 | 6923 | JVDM\_TOPE.raw | 9.8819E4 | 1 | 1 |  |  |  | PEAKS DB |
| EATAAPGPKSPGKSE | 16.02 | 1425.7100 | 15 | 6.6 | 713.8669 | 17.82 | 2 | 3492 | JVDM\_TOPE.raw | 1.2768E5 | 1 | 1 |  |  |  | PEAKS DB |
| GGGGGHGAGGGSGAGGSK | 16.01 | 1283.5603 | 18 | 5.0 | 642.7906 | 16.05 | 2 | 2888 | JVDM\_TOPE.raw | 3.0101E5 | 1 | 1 |  |  |  | PEAKS DB |
| GGAGGSTAGGGGGSGGNGASPR | 15.98 | 1644.7200 | 22 | -1.4 | 823.3661 | 19.36 | 2 | 4020 | JVDM\_TOPE.raw | 0 | 0 | 0 |  |  |  | PEAKS DB |
| GGGGGGGGGGGGGGGGGGGGPGR | 15.96 | 1468.6151 | 23 | 5.6 | 735.3189 | 12.36 | 2 | 1415 | JVDM\_TOPE.raw | 0 | 0 | 0 |  |  |  | PEAKS DB |
| AVAMASSSSGGAGGSGR | 15.93 | 1408.6365 | 17 | 0.6 | 705.3259 | 23.18 | 2 | 5181 | JVDM\_TOPE.raw | 0 | 0 | 0 |  |  |  | PEAKS DB |
| KASGGAGGAAAAADGVAV | 15.92 | 1399.7054 | 18 | -2.5 | 700.8583 | 24.22 | 2 | 5454 | JVDM\_TOPE.raw | 0 | 0 | 0 |  |  |  | PEAKS DB |
| GGGGGGGGADAALVLELSASVDKAASGGDK | 15.92 | 2543.2463 | 30 | 5.0 | 848.7603 | 27.52 | 2 | 6242 | JVDM\_TOPE.raw | 0 | 0 | 0 |  |  |  | PEAKS DB |
| MGGGGPGGGPPGK | 15.91 | 1024.4760 | 13 | -9.3 | 513.2405 | 13.45 | 2 | 2264 | JVDM\_TOPE.raw | 1.3732E6 | 1 | 1 |  |  |  | PEAKS DB |
| AGGGAGGAGGGGGGR | 15.89 | 1014.4590 | 15 | -1.6 | 508.2360 | 12.24 | 2 | 1302 | JVDM\_TOPE.raw | 0 | 0 | 0 |  |  |  | PEAKS DB |
| EGGGVAGGVDGPLHQR | 15.87 | 1504.7382 | 16 | -6.5 | 753.3715 | 19.92 | 2 | 4201 | JVDM\_TOPE.raw | 7.7191E4 | 1 | 1 |  |  |  | PEAKS DB |
| AGAAAS(+79.97)AAQSTAASAVEKAGSLTQR | 15.85 | 2354.1226 | 25 | 3.6 | 785.7176 | 34.77 | 2 | 7989 | JVDM\_TOPE.raw | 0 | 0 | 0 | tr\|A0A2K3E7M7\|A0A2K3E7M7\_CHLRE | Phosphorylation (STY) | S6:Phosphorylation (STY):0.00 | PEAKS PTM |
| GEGGGGGRAGPAVAAGAAEAAAR | 15.83 | 1879.9248 | 23 | -2.6 | 627.6473 | 24.64 | 2 | 5583 | JVDM\_TOPE.raw | 2.5087E4 | 1 | 1 |  |  |  | PEAKS DB |
| YLPTPVTK | 15.83 | 917.5222 | 8 | 0.6 | 459.7687 | 17.11 | 2 | 3269 | JVDM\_TOPE.raw | 1.5597E5 | 1 | 1 |  |  |  | PEAKS DB |
| AGAAASAAQ(+.98)STAASAVEKASGAATAAQ(+.98)AAAGSAAEK | 15.82 | 3047.4641 | 36 | 8.0 | 1016.8368 | 33.66 | 2 | 7789 | JVDM\_TOPE.raw | 3.8661E5 | 1 | 1 | tr\|A0A2K3E7M7\|A0A2K3E7M7\_CHLRE | Deamidation (NQ) | Q9:Deamidation (NQ):1000.00;Q27:Deamidation (NQ):1000.00 | PEAKS PTM |
| EAGASGASLVAALK | 15.80 | 1243.6771 | 14 | 1.4 | 622.8467 | 24.80 | 2 | 5632 | JVDM\_TOPE.raw | 3.3209E4 | 1 | 1 |  |  |  | PEAKS DB |
| EGGAAGAALQR | 15.79 | 999.5097 | 11 | -6.1 | 500.7591 | 20.80 | 2 | 4464 | JVDM\_TOPE.raw | 0 | 0 | 0 |  |  |  | PEAKS DB |
| AAAAGAGAGGGAGAGAGAGAGAGAGAGAGAGAGAGAGAGAGAR | 15.78 | 2877.3574 | 43 | 6.4 | 960.1325 | 25.39 | 2 | 5768 | JVDM\_TOPE.raw | 0 | 0 | 0 |  |  |  | PEAKS DB |
| AGLETLYGAALR | 15.77 | 1233.6716 | 12 | 1.7 | 617.8441 | 16.35 | 2 | 3058 | JVDM\_TOPE.raw | 2.8337E6 | 1 | 1 |  |  |  | PEAKS DB |
| EGGGGGGGGGGGGGGGGFGG | 15.74 | 1320.5078 | 20 | 1.7 | 441.1773 | 12.69 | 2 | 1721 | JVDM\_TOPE.raw | 0 | 0 | 0 |  |  |  | PEAKS DB |
| TAPAAPGGAGGGGSFGR | 15.70 | 1386.6639 | 17 | -4.9 | 694.3359 | 38.05 | 2 | 8825 | JVDM\_TOPE.raw | 0 | 0 | 0 |  |  |  | PEAKS DB |
| RGGGTSAAATAAAAGHV | 15.69 | 1424.7120 | 17 | 4.3 | 713.3663 | 24.39 | 2 | 5464 | JVDM\_TOPE.raw | 6.7342E5 | 1 | 1 |  |  |  | PEAKS DB |
| GQEIVISTVGAAALSEQPK | 15.68 | 1897.0156 | 19 | 0.9 | 949.5159 | 28.69 | 2 | 6501 | JVDM\_TOPE.raw | 0 | 0 | 0 |  |  |  | PEAKS DB |
| EASSGASAHADNASGQVAMTPSRGSSGSSGK | 15.66 | 2848.2642 | 31 | -4.0 | 713.0705 | 16.59 | 2 | 3104 | JVDM\_TOPE.raw | 0 | 0 | 0 |  |  |  | PEAKS DB |
| LAGGVGAPGYR | 15.66 | 1016.5403 | 11 | -7.2 | 509.2737 | 6.04 | 2 | 189 | JVDM\_TOPE.raw | 1.0113E5 | 1 | 1 |  |  |  | PEAKS DB |
| ASSLLGLNSGAGSSR | 15.66 | 1375.7056 | 15 | -7.5 | 688.8549 | 17.96 | 2 | 3566 | JVDM\_TOPE.raw | 3.8081E4 | 1 | 1 |  |  |  | PEAKS DB |
| KADPAPAATPAAK | 15.63 | 1207.6560 | 13 | -7.4 | 604.8308 | 26.86 | 2 | 6081 | JVDM\_TOPE.raw | 1.621E6 | 1 | 1 |  |  |  | PEAKS DB |
| AGGSGGGAAAGGAAAVGR | 15.60 | 1313.6436 | 18 | 4.5 | 657.8320 | 24.64 | 2 | 5580 | JVDM\_TOPE.raw | 3.7387E4 | 1 | 1 |  |  |  | PEAKS DB |
| KAEAAAAAAAAGGGAAPG | 15.54 | 1381.6949 | 18 | 4.1 | 691.8576 | 29.98 | 2 | 6826 | JVDM\_TOPE.raw | 1.2402E5 | 1 | 1 |  |  |  | PEAKS DB |
| GSSAEPLAKLLLGK | 15.52 | 1382.8132 | 14 | 9.0 | 692.4201 | 39.96 | 2 | 9346 | JVDM\_TOPE.raw | 2.1048E6 | 1 | 1 |  |  |  | PEAKS DB |
| LPRPAGGGGSSGGPAGTK | 15.50 | 1522.7852 | 18 | -1.6 | 508.6015 | 23.49 | 2 | 5268 | JVDM\_TOPE.raw | 0 | 0 | 0 |  |  |  | PEAKS DB |
| GSDGGGGGGGGGGSSGSSGGGGGGGGLQGPAESAR | 15.49 | 2618.0938 | 35 | -3.6 | 873.7021 | 19.00 | 2 | 3905 | JVDM\_TOPE.raw | 0 | 0 | 0 |  |  |  | PEAKS DB |
| LSGSGSGAGGGRGGR | 15.49 | 1231.6017 | 15 | 7.0 | 616.8124 | 22.02 | 2 | 4853 | JVDM\_TOPE.raw | 1.622E5 | 1 | 1 |  |  |  | PEAKS DB |
| AGKGAAPAGAATSTRDAAGGEEGK | 15.45 | 2100.0195 | 24 | -3.4 | 1051.0134 | 28.16 | 2 | 6393 | JVDM\_TOPE.raw | 3.4154E4 | 1 | 1 |  |  |  | PEAKS DB |
| GAGPGSGGGGGGGGGGLTWTGSALELAK | 15.43 | 2285.1035 | 28 | -9.2 | 762.7014 | 30.17 | 2 | 6881 | JVDM\_TOPE.raw | 3.1283E5 | 1 | 1 |  |  |  | PEAKS DB |
| EEGGVKTSIIIKK | 15.39 | 1400.8239 | 13 | 4.7 | 701.4225 | 24.58 | 2 | 5572 | JVDM\_TOPE.raw | 4.7459E3 | 1 | 1 |  |  |  | PEAKS DB |
| EAGATGSSAGGAR | 15.38 | 1090.5002 | 13 | 9.4 | 546.2625 | 13.89 | 2 | 2297 | JVDM\_TOPE.raw | 8.1192E5 | 1 | 1 |  |  |  | PEAKS DB |
| RAALSAVPAVC(+57.02)R | 15.35 | 1269.6975 | 12 | 9.8 | 635.8622 | 24.02 | 2 | 5401 | JVDM\_TOPE.raw | 0 | 0 | 0 |  | Carbamidomethylation | C11:Carbamidomethylation:1000.00 | PEAKS DB |
| ATGGKAAVSTDTLQR | 15.33 | 1474.7739 | 15 | 5.3 | 738.3981 | 18.64 | 2 | 3774 | JVDM\_TOPE.raw | 0 | 0 | 0 |  |  |  | PEAKS DB |
| VSAGLSGAEAGYGSAVER | 15.33 | 1679.8114 | 18 | -4.9 | 560.9417 | 12.63 | 2 | 1647 | JVDM\_TOPE.raw | 0 | 0 | 0 |  |  |  | PEAKS DB |
| AGGGAGGGSLHRDAAAAR | 15.33 | 1550.7661 | 18 | -1.2 | 776.3894 | 24.02 | 2 | 5396 | JVDM\_TOPE.raw | 3.8705E4 | 1 | 1 |  |  |  | PEAKS DB |
| EGGGSNGGSGGGGVGSR | 15.32 | 1347.5763 | 17 | -3.1 | 674.7933 | 15.94 | 2 | 2912 | JVDM\_TOPE.raw | 1.0634E5 | 1 | 1 |  |  |  | PEAKS DB |
| GAASLDGSAFVA | 15.31 | 1064.5138 | 12 | -0.9 | 533.2637 | 13.37 | 2 | 2109 | JVDM\_TOPE.raw | 3.8034E4 | 1 | 1 |  |  |  | PEAKS DB |
| SSAGGGGAGAVVTT | 15.29 | 1090.5254 | 14 | -1.7 | 546.2690 | 13.25 | 2 | 2082 | JVDM\_TOPE.raw | 1.0449E5 | 1 | 1 |  |  |  | PEAKS DB |
| GGGGAGGAAGGGGGGGGWVVTGAHRR | 15.28 | 2092.0059 | 26 | -0.5 | 698.3422 | 32.99 | 2 | 7501 | JVDM\_TOPE.raw | 0 | 0 | 0 |  |  |  | PEAKS DB |
| ASGGGGGAGAAAAPAGR | 15.25 | 1254.6064 | 17 | -4.1 | 419.2077 | 12.52 | 2 | 1532 | JVDM\_TOPE.raw | 0 | 0 | 0 |  |  |  | PEAKS DB |
| IPFWEGQSYGVIPPGTK(+14.02)IN(+.98)SK(+28.03) | 15.25 | 2360.2415 | 21 | 5.1 | 787.7584 | 32.06 | 2 | 7266 | JVDM\_TOPE.raw | 2.2242E5 | 1 | 1 | tr\|A8J6Y8\|A8J6Y8\_CHLRE | Methylation(KR); Deamidation (NQ); Dimethylation(KR) | K17:Methylation(KR):2.76;N19:Deamidation (NQ):47.33;K21:Dimethylation(KR):2.76 | PEAKS PTM |
| VGSGAAGPAHAQHSTSPSPPSR | 15.24 | 2054.9883 | 22 | 0.1 | 686.0034 | 12.60 | 2 | 1594 | JVDM\_TOPE.raw | 0 | 0 | 0 |  |  |  | PEAKS DB |
| RTVPTKLEEGEMPLNTYSNK | 15.19 | 2306.1577 | 20 | 3.2 | 769.7290 | 20.00 | 2 | 4235 | JVDM\_TOPE.raw | 1.1472E4 | 1 | 1 | tr\|A8J6Y8\|A8J6Y8\_CHLRE |  |  | PEAKS DB |
| VEYTELQILC(+57.02)PQ | 15.18 | 1491.7279 | 12 | 1.1 | 746.8721 | 32.68 | 2 | 7400 | JVDM\_TOPE.raw | 2.4394E5 | 1 | 1 | tr\|A0A2K3CQ54\|A0A2K3CQ54\_CHLRE | Carbamidomethylation | C10:Carbamidomethylation:1000.00 | PEAKS DB |
| QGPSSLAVEGGR | 15.14 | 1156.5836 | 12 | 1.7 | 579.3000 | 18.85 | 2 | 3786 | JVDM\_TOPE.raw | 9.1305E6 | 1 | 1 |  |  |  | PEAKS DB |
| ASGGGAGGGGGGGGGGAAGRPRSAATR | 15.13 | 2083.0015 | 27 | -3.8 | 1042.5040 | 30.19 | 2 | 6876 | JVDM\_TOPE.raw | 0 | 0 | 0 |  |  |  | PEAKS DB |
| QVSSGGDLLR | 15.13 | 1030.5406 | 10 | 3.3 | 516.2793 | 16.96 | 2 | 3225 | JVDM\_TOPE.raw | 1.1739E5 | 1 | 1 |  |  |  | PEAKS DB |
| ARGGGGGGGGGGAVSDAVVLYGGPGAG | 15.12 | 2129.0249 | 27 | -6.0 | 710.6780 | 24.30 | 2 | 5484 | JVDM\_TOPE.raw | 2.0105E4 | 1 | 1 |  |  |  | PEAKS DB |
| AAATMAGGGVAR | 15.12 | 1031.5182 | 12 | 7.5 | 516.7703 | 19.41 | 2 | 4035 | JVDM\_TOPE.raw | 7.2777E4 | 1 | 1 |  |  |  | PEAKS DB |
| KVAASSSGSSSR | 15.11 | 1122.5629 | 12 | -8.7 | 562.2838 | 13.05 | 2 | 2010 | JVDM\_TOPE.raw | 0 | 0 | 0 |  |  |  | PEAKS DB |
| EGGGGPGFSFGAPAGTSGIGGFGA | 15.08 | 2010.9071 | 24 | -2.4 | 671.3080 | 19.55 | 2 | 4072 | JVDM\_TOPE.raw | 0 | 0 | 0 |  |  |  | PEAKS DB |
| EGGGGGGGGGGGGGGGGFGGGG | 15.08 | 1434.5508 | 22 | -6.3 | 718.2781 | 12.81 | 2 | 2014 | JVDM\_TOPE.raw | 1.0993E6 | 1 | 1 |  |  |  | PEAKS DB |
| EAPGGGGGGGGAAPAPAYR | 15.07 | 1568.7330 | 19 | -2.0 | 785.3722 | 26.13 | 2 | 5947 | JVDM\_TOPE.raw | 0 | 0 | 0 |  |  |  | PEAKS DB |
| EDGKAAEAATAAPSTSR | 15.05 | 1631.7750 | 17 | -9.2 | 816.8873 | 23.48 | 2 | 5259 | JVDM\_TOPE.raw | 0 | 0 | 0 |  |  |  | PEAKS DB |
| GSGDGGGTAGGGGGVFGVTGR | 15.04 | 1678.7659 | 21 | 1.3 | 840.3913 | 21.93 | 2 | 4823 | JVDM\_TOPE.raw | 1.941E4 | 1 | 1 |  |  |  | PEAKS DB |
| EAGDVSSAAGAGSAAR | 15.04 | 1375.6327 | 16 | 9.4 | 688.8301 | 21.66 | 2 | 4726 | JVDM\_TOPE.raw | 0 | 0 | 0 |  |  |  | PEAKS DB |
| AAGAGVASSALRRR | 15.02 | 1341.7589 | 14 | 4.6 | 671.8898 | 24.43 | 2 | 5517 | JVDM\_TOPE.raw | 8.9245E4 | 1 | 1 |  |  |  | PEAKS DB |
| GGGGGGGGGGGGGGGGGGGGGGGGGYSV | 15.02 | 1792.7108 | 28 | 10.1 | 897.3717 | 21.16 | 2 | 4580 | JVDM\_TOPE.raw | 1.1259E5 | 1 | 1 |  |  |  | PEAKS DB |
| total 484 peptides |  |  |  |  |  |  |  |  |  |  |  |  |  |  |  |  |
| --- | --- | --- | --- | --- | --- | --- | --- | --- | --- | --- | --- | --- | --- | --- | --- | --- |
