## Supplementary material for "Efficient secretion of a plastic degrading enzyme from the green algae *Chlamydomonas reinhardtii*": Dataset - https://doi.org/10.5281/zenodo.13981200: protein.html

proteins


Summary

  

### 1. Notes

### 2. Result Statistics

**Figure 1.**
False discovery rate (FDR) curve. X axis is the number of peptides being kept. Y axis is the corresponding FDR.

  


  

**Figure 2.**
PSM score distribution.  **(a)**
Distribution of PEAKS peptide score;  **(b)**
Scatterplot of PEAKS peptide score versus precursor mass error.

**(a)**

**(b)**

**Table 1.**
Statistics of data.


|  |  |  |  |  |  |  |  |  |  |  |  |  |
| --- | --- | --- | --- | --- | --- | --- | --- | --- | --- | --- | --- | --- |
|  | #Scans | | | #Features | Identified | | | #Peptides | #Sequences | #Proteins^\*^ | | |
|  | MS1 | MS/MS | #Chimera |  | #PSMs | #Scans | #Features^\*\*^ |  |  | Groups | All | Top |
| Total | 3085 | 8625 | 532 | 19758 | 559 | 557 | 376 | 484 | 476 | 62 | 77 | 71 |
| Sample 2 | 3085 | 8625 | 532 | 19758 | 559 | 557 | 376 | 484 | 476 | 62 | 77 | 71 |

^\* proteins with significant peptides are used in counts.^  
^\*\* features are identified by DB search only.^

**Figure 3.**
Sample overlap for Proteins and Peptides (up to 8 samples).  **(a)**
All Proteins;  **(b)**
Top Proteins;  **(c)**
Peptides;

**(a)**

**Not applicable to only one sample**

**(b)**

**Not applicable to only one sample**

**(c)**

**Not applicable to only one sample**

**Figure 4.**
Distribution of peptide feature detection.  **(a)**
Feature m/z distribution;  **(b)**
Feature RT distribution.

**(a)**

**(b)**

**Figure 5.**
Distribution of identified peptide features.  **(a)**
Feature abundance distribution;  **(b)**
 *De novo*
sequencing validation.

**(a)**

**(b)**

**Table 2.**
Result filtration parameters.

|  |  |
| --- | --- |
| Peptide -10lgP | ≥15 |
| PTM Ascore | ≥0 |
| Protein -10lgP | ≥20 |
| Proteins unique peptides | ≥0 |
| De novo score(%) | ≥50% |


**Table 3.**
Statistics of filtered result.

|  |  |
| --- | --- |
| FDR (Peptide-Spectrum Matches) | 18.8% |
| FDR (Peptide Sequences) | 21.1% |
| FDR (Protein Group) | 0.0% |
| De Novo Only Spectra | 2620 |

**Table 4.**
PTM profile.

| Name | ∆Mass | Position | #PSM | -10lgP | Abundance | AScore || Carbamidomethyl | 57.02 | C | 52 | 88.47 | 8.99E4 | 1000.00 |
| Deamidation | .98 | NQ | 14 | 77.00 | 9.53E4 | 1000.00 |

### 3. Experiment Control

**Figure 6.**
Precursor mass error of peptide-spectrum matches (PSM) in filtered result.  **(a)**
Distribution of precursor mass error in ppm;  **(b)**
Scatterplot of precursor m/z versus precursor mass error in ppm.

**(a)**

**(b)**

**Table 5.**
Number of identified peptides in each sample by the number of missed cleavages.

|  |  |  |  |  |  |
| --- | --- | --- | --- | --- | --- |
| Missed Cleavages | 0 | 1 | 2 | 3 | 4+ |
| Sample 2 | 364 | 101 | 19 | 0 | 0 |

### 4. Other Information

**Table 6.**
Search parameters.

PEAKS Version: PEAKS Studio 10.6 build 20201221  
Search Engine Name: PEAKS  
Parent Mass Error Tolerance: 10.0 ppm  
Fragment Mass Error Tolerance: 0.5 Da  
Precursor Mass Search Type: monoisotopic  
Enzyme: Trypsin  
Max Missed Cleavages: 2  
Digest Mode: Semispecific  
Fixed Modifications:   
  Carbamidomethylation: 57.02  
Variable Modifications:   
  Acetylation (K): 42.01  
  Deamidation (NQ): 0.98  
  Dimethylation(KR): 28.03  
  Dimethylation(NP): 28.03  
  Dioxidation (M): 31.99  
  Methylation(KR): 14.02  
  Phosphorylation (STY): 79.97  
  Biotinylation: 226.08  
Max Variable PTM Per Peptide: 3  
Database: Chlamydomonas reinhardtii uniprot  
Taxon: All  
Searched Entry: 15000  
FDR Estimation: Enabled  
De novo score(%) threshold: 15  
Peptide hit threshold (-10logP): 30.0  
Peaks run ID: 7  
Merge Options: no merge  
Precursor Options: corrected  
Charge Options: no correction  
Filter Charge: 2 - 8  
Process: true  
Associate chimera: yes

**Table 7.**
Instrument parameters.

Fractions: JVDM\_TOPE.raw  
Ion Source: ESI(nano-spray)  
Fragmentation Mode: high energy CID (y and b ions)  
MS Scan Mode: FT-ICR/Orbitrap  
MS/MS Scan Mode: Linear Ion Trap

  

Protein List

  

Protein Accession Contains:

Protein Description Contains:

Protein Sample Area >=

Protein PTM Contains:

| Protein Group | Protein ID | Accession | -10lgP | Coverage (%) | Coverage (%) Sample 2 | Area Sample 2 | #Peptides | #Unique | #Spec Sample 2 | PTM | Avg. Mass | Description |
| --- | --- | --- | --- | --- | --- | --- | --- | --- | --- | --- | --- | --- |
| 2 | 1 | tr\|A0A2K3D8I0\|A0A2K3D8I0\_CHLRE | 288.98 | 39 | 39 | 3.1459E6 | 17 | 17 | 25 | N | 94346 | Uncharacterized protein OS=Chlamydomonas reinhardtii OX=3055 GN=CHLRE\_11g477950v5 PE=4 SV=1 |
| 1 | 2 | PHL7 | 231.84 | 44 | 44 | 3.4701E7 | 11 | 11 | 35 | Y | 28201 | PHL7 |
| 3 | 212 | tr\|A8J6Y8\|A8J6Y8\_CHLRE | 221.96 | 57 | 57 | 3.861E7 | 16 | 16 | 25 | Y | 38267 | Ferredoxin--NADP reductase, chloroplastic OS=Chlamydomonas reinhardtii OX=3055 GN=FNR1 PE=3 SV=1 |
| 5 | 5722 | tr\|A0A2K3CTJ7\|A0A2K3CTJ7\_CHLRE | 204.32 | 29 | 29 | 1.7985E6 | 8 | 8 | 11 | Y | 38468 | Uncharacterized protein OS=Chlamydomonas reinhardtii OX=3055 GN=CHLRE\_16g661750v5 PE=4 SV=1 |
| 8 | 5723 | tr\|A0A2K3D1P1\|A0A2K3D1P1\_CHLRE | 191.73 | 36 | 36 | 6.7485E5 | 7 | 7 | 8 | Y | 35139 | Malate dehydrogenase OS=Chlamydomonas reinhardtii OX=3055 GN=CHLRE\_12g483950v5 PE=3 SV=1 |
| 9 | 9 | tr\|A0A2K3E094\|A0A2K3E094\_CHLRE | 176.58 | 12 | 12 | 1.1502E6 | 7 | 3 | 10 | Y | 81741 | Uncharacterized protein OS=Chlamydomonas reinhardtii OX=3055 GN=CHLRE\_02g077750v5 PE=4 SV=1 |
| 7 | 4 | tr\|A0A2K3D6N5\|A0A2K3D6N5\_CHLRE | 170.59 | 16 | 16 | 2.4577E6 | 7 | 7 | 9 | Y | 81251 | Uncharacterized protein OS=Chlamydomonas reinhardtii OX=3055 GN=CHLRE\_12g544450v5 PE=4 SV=1 |
| 6 | 13 | tr\|A0A2K3E0A7\|A0A2K3E0A7\_CHLRE | 154.25 | 14 | 14 | 3.3142E5 | 7 | 2 | 11 | Y | 83460 | Uncharacterized protein OS=Chlamydomonas reinhardtii OX=3055 GN=CHLRE\_02g077850v5 PE=4 SV=1 |
| 4 | 16 | tr\|A0A2K3CQ54\|A0A2K3CQ54\_CHLRE | 150.80 | 6 | 6 | 3.9384E7 | 7 | 7 | 15 | Y | 74085 | Peptidase\_M11 domain-containing protein OS=Chlamydomonas reinhardtii OX=3055 GN=CHLRE\_17g718500v5 PE=4 SV=1 |
| 10 | 5725 | tr\|P93106\|P93106\_CHLRE | 138.98 | 32 | 32 | 7.3594E6 | 9 | 9 | 11 | Y | 36602 | Malate dehydrogenase OS=Chlamydomonas reinhardtii OX=3055 GN=MDH2 PE=2 SV=1 |
| 15 | 5 | tr\|Q9FE86\|Q9FE86\_CHLRE | 135.66 | 26 | 26 | 1.2324E6 | 4 | 4 | 4 | N | 25962 | Peroxiredoxin OS=Chlamydomonas reinhardtii OX=3055 GN=thioredoxin peroxidase PE=1 SV=1 |
| 11 | 693 | tr\|A8JGV6\|A8JGV6\_CHLRE | 134.32 | 22 | 22 | 5.3869E6 | 4 | 4 | 7 | N | 29514 | 14-3-3 protein OS=Chlamydomonas reinhardtii OX=3055 GN=FTT1 PE=3 SV=1 |
| 13 | 43 | P06541\|ATPB\_CHLRE | 127.12 | 22 | 22 | 4.5316E5 | 7 | 7 | 8 | N | 52040 | ATP synthase subunit beta, chloroplastic OS=Chlamydomonas reinhardtii OX=3055 GN=atpB PE=1 SV=3 |
| 20 | 19 | tr\|A8J9H8\|A8J9H8\_CHLRE | 124.25 | 17 | 17 | 1.7992E6 | 3 | 3 | 3 | N | 16541 | Nucleoside diphosphate kinase OS=Chlamydomonas reinhardtii OX=3055 GN=FAP103 PE=3 SV=1 |
| 20 | 20 | tr\|A0A2K3DJF1\|A0A2K3DJF1\_CHLRE | 124.25 | 16 | 16 | 1.7992E6 | 3 | 3 | 3 | N | 18054 | Nucleoside diphosphate kinase OS=Chlamydomonas reinhardtii OX=3055 GN=CHLRE\_07g325734v5 PE=3 SV=1 |
| 12 | 294 | tr\|A8JC04\|A8JC04\_CHLRE | 119.56 | 18 | 18 | 6.1776E5 | 7 | 7 | 7 | Y | 49032 | Phosphoglycerate kinase OS=Chlamydomonas reinhardtii OX=3055 GN=PGK1 PE=3 SV=1 |
| 17 | 8 | tr\|B7U1J0\|B7U1J0\_CHLRE | 116.91 | 12 | 12 | 1.0539E6 | 5 | 5 | 5 | N | 54752 | ATP synthase subunit alpha, chloroplastic OS=Chlamydomonas reinhardtii OX=3055 GN=atpA PE=3 SV=1 |
| 17 | 7 | P26526\|ATPA\_CHLRE | 116.91 | 12 | 12 | 1.0539E6 | 5 | 5 | 5 | N | 54752 | ATP synthase subunit alpha, chloroplastic OS=Chlamydomonas reinhardtii OX=3055 GN=atpA PE=1 SV=3 |
| 31 | 5735 | tr\|A0A2K3DJ22\|A0A2K3DJ22\_CHLRE | 115.21 | 2 | 2 | 1.3686E5 | 2 | 2 | 2 | Y | 199243 | Uncharacterized protein OS=Chlamydomonas reinhardtii OX=3055 GN=CHLRE\_07g321400v5 PE=4 SV=1 |
| 18 | 22 | tr\|A8HXL8\|A8HXL8\_CHLRE | 114.23 | 12 | 12 | 8.6044E5 | 4 | 4 | 5 | Y | 38761 | Chloroplast ATP synthase gamma chain OS=Chlamydomonas reinhardtii OX=3055 GN=ATPC PE=3 SV=1 |
| 21 | 11 | tr\|A8IGH1\|A8IGH1\_CHLRE | 110.37 | 16 | 16 | 9.7983E5 | 3 | 3 | 3 | N | 25915 | Superoxide dismutase OS=Chlamydomonas reinhardtii OX=3055 GN=FSD1 PE=2 SV=1 |
| 22 | 37 | tr\|A8J0E4\|A8J0E4\_CHLRE | 108.32 | 26 | 26 | 4.9084E4 | 4 | 4 | 4 | Y | 30580 | Oxygen-evolving enhancer protein 1 of photosystem II OS=Chlamydomonas reinhardtii OX=3055 GN=PSBO PE=1 SV=1 |
| 16 | 5726 | tr\|A0A2K3CWZ6\|A0A2K3CWZ6\_CHLRE | 106.10 | 24 | 24 | 7.6969E5 | 6 | 6 | 6 | Y | 31543 | SOR\_SNZ domain-containing protein OS=Chlamydomonas reinhardtii OX=3055 GN=CHLRE\_15g643550v5 PE=3 SV=1 |
| 34 | 18 | tr\|A8IKW6\|A8IKW6\_CHLRE | 102.21 | 12 | 12 | 2.9659E4 | 2 | 2 | 2 | N | 28401 | Ribulose-phosphate 3-epimerase OS=Chlamydomonas reinhardtii OX=3055 GN=RPE1 PE=3 SV=1 |
| 27 | 5728 | tr\|A8ILJ9\|A8ILJ9\_CHLRE | 101.84 | 14 | 14 | 2.233E5 | 3 | 3 | 4 | Y | 40386 | Ornithine carbamoyltransferase OS=Chlamydomonas reinhardtii OX=3055 GN=OTC1 PE=3 SV=1 |
| 19 | 5727 | tr\|A0A2K3DZ46\|A0A2K3DZ46\_CHLRE | 101.31 | 5 | 5 | 3.6457E5 | 3 | 3 | 3 | N | 90598 | 4Fe-4S ferredoxin-type domain-containing protein OS=Chlamydomonas reinhardtii OX=3055 GN=CHLRE\_03g204450v5 PE=4 SV=1 |
| 28 | 10 | tr\|A8JEV1\|A8JEV1\_CHLRE | 100.98 | 16 | 16 | 1.1741E6 | 3 | 3 | 3 | N | 21824 | Oxygen evolving enhancer protein 3 OS=Chlamydomonas reinhardtii OX=3055 GN=PSBQ PE=3 SV=1 |
| 32 | 15 | tr\|A8IKZ2\|A8IKZ2\_CHLRE | 100.76 | 17 | 17 | 5.6967E5 | 2 | 2 | 2 | Y | 20930 | Ribosomal protein L18 OS=Chlamydomonas reinhardtii OX=3055 GN=RPL18 PE=3 SV=1 |
| 24 | 12 | tr\|A0A2K3DP12\|A0A2K3DP12\_CHLRE | 99.14 | 6 | 6 | 3.9308E5 | 3 | 3 | 4 | Y | 79150 | Peptidase\_M11 domain-containing protein OS=Chlamydomonas reinhardtii OX=3055 GN=CHLRE\_06g278165v5 PE=4 SV=1 |
| 29 | 5729 | tr\|A8J7F8\|A8J7F8\_CHLRE | 86.31 | 16 | 16 | 2.1616E5 | 3 | 3 | 3 | N | 31316 | S-formylglutathione hydrolase OS=Chlamydomonas reinhardtii OX=3055 GN=CHLRE\_03g158800v5 PE=3 SV=1 |
| 29 | 5730 | tr\|A0A2K3DW85\|A0A2K3DW85\_CHLRE | 86.31 | 16 | 16 | 2.1616E5 | 3 | 3 | 3 | N | 31545 | S-formylglutathione hydrolase OS=Chlamydomonas reinhardtii OX=3055 GN=CHLRE\_03g158800v5 PE=3 SV=1 |
| 29 | 5731 | tr\|A0A2K3DW83\|A0A2K3DW83\_CHLRE | 86.31 | 15 | 15 | 2.1616E5 | 3 | 3 | 3 | N | 32148 | S-formylglutathione hydrolase OS=Chlamydomonas reinhardtii OX=3055 GN=CHLRE\_03g158800v5 PE=3 SV=1 |
| 51 | 5743 | tr\|A8IMR9\|A8IMR9\_CHLRE | 85.29 | 5 | 5 | 8.6679E4 | 1 | 1 | 1 | N | 36007 | Predicted protein OS=Chlamydomonas reinhardtii OX=3055 GN=CHLRE\_06g307750v5 PE=4 SV=1 |
| 30 | 47 | tr\|A0A2K3D9K7\|A0A2K3D9K7\_CHLRE | 82.51 | 2 | 2 | 3.0578E5 | 1 | 1 | 2 | N | 77363 | Uncharacterized protein OS=Chlamydomonas reinhardtii OX=3055 GN=CHLRE\_10g427300v5 PE=3 SV=1 |
| 14 | 17 | Q08365\|RR3\_CHLRE | 81.55 | 7 | 7 | 3.8312E5 | 4 | 4 | 5 | N | 81855 | 30S ribosomal protein S3, chloroplastic OS=Chlamydomonas reinhardtii OX=3055 GN=rps3 PE=1 SV=2 |
| 33 | 23 | tr\|A0A2K3D661\|A0A2K3D661\_CHLRE | 79.00 | 15 | 15 | 4.7676E5 | 2 | 2 | 2 | N | 25899 | PsbP domain-containing protein OS=Chlamydomonas reinhardtii OX=3055 GN=CHLRE\_12g550850v5 PE=4 SV=1 |
| 26 | 55 | tr\|A8J363\|A8J363\_CHLRE | 77.68 | 4 | 4 | 1.8226E5 | 2 | 2 | 2 | N | 80822 | Matrix metalloproteinase-like protein OS=Chlamydomonas reinhardtii OX=3055 GN=MMP13 PE=4 SV=1 |
| 41 | 5738 | tr\|A8JAV1\|A8JAV1\_CHLRE | 70.14 | 7 | 7 | 1.5519E6 | 2 | 2 | 3 | N | 41836 | Actin OS=Chlamydomonas reinhardtii OX=3055 GN=IDA5 PE=3 SV=1 |
| 42 | 24 | tr\|A8I2V3\|A8I2V3\_CHLRE | 67.34 | 15 | 15 | 1.3223E5 | 2 | 2 | 2 | N | 21642 | Peroxiredoxin OS=Chlamydomonas reinhardtii OX=3055 GN=PRX2 PE=3 SV=1 |
| 43 | 36 | tr\|A8J244\|A8J244\_CHLRE | 66.34 | 6 | 6 | 2.1749E5 | 2 | 2 | 2 | N | 45749 | Isocitrate lyase OS=Chlamydomonas reinhardtii OX=3055 GN=ICL1 PE=4 SV=1 |
| 46 | 14 | tr\|A0A2K3CWF7\|A0A2K3CWF7\_CHLRE | 66.12 | 8 | 8 | 3.2087E4 | 1 | 1 | 2 | N | 21332 | NAC-A/B domain-containing protein OS=Chlamydomonas reinhardtii OX=3055 GN=CHLRE\_15g635600v5 PE=4 SV=1 |
| 23 | 3 | tr\|Q7X7A7\|Q7X7A7\_CHLRE | 65.12 | 20 | 20 | 3.4392E5 | 4 | 4 | 4 | N | 28003 | 14-3-3 protein OS=Chlamydomonas reinhardtii OX=3055 GN=erb14 PE=2 SV=1 |
| 44 | 5739 | tr\|A0A2K3DP15\|A0A2K3DP15\_CHLRE | 62.51 | 7 | 7 | 5.061E4 | 1 | 1 | 1 | N | 39384 | Uncharacterized protein OS=Chlamydomonas reinhardtii OX=3055 GN=CHLRE\_06g278148v5 PE=4 SV=1 |
| 40 | 5737 | tr\|A8HRZ0\|A8HRZ0\_CHLRE | 59.09 | 11 | 11 | 9.1544E5 | 3 | 3 | 3 | N | 27356 | Histone H1 OS=Chlamydomonas reinhardtii OX=3055 GN=HON1 PE=3 SV=1 |
| 25 | 5733 | tr\|A0A2K3CQL0\|A0A2K3CQL0\_CHLRE | 57.75 | 3 | 3 | 3.0571E5 | 2 | 2 | 2 | Y | 77435 | TROVE domain-containing protein OS=Chlamydomonas reinhardtii OX=3055 GN=CHLRE\_17g725750v5 PE=3 SV=1 |
| 25 | 5734 | tr\|A0A2K3CQP7\|A0A2K3CQP7\_CHLRE | 57.75 | 3 | 3 | 3.0571E5 | 2 | 2 | 2 | Y | 83399 | TROVE domain-containing protein OS=Chlamydomonas reinhardtii OX=3055 GN=CHLRE\_17g726850v5 PE=3 SV=1 |
| 38 | 5740 | tr\|A0A2K3E350\|A0A2K3E350\_CHLRE | 57.14 | 2 | 2 | 4.4689E4 | 1 | 1 | 1 | N | 54500 | Glutamine synthetase OS=Chlamydomonas reinhardtii OX=3055 GN=CHLRE\_02g113200v5 PE=3 SV=1 |
| 52 | 5744 | tr\|Q6UKY5\|Q6UKY5\_CHLRE | 56.52 | 11 | 11 | 1.8672E5 | 1 | 1 | 1 | N | 12424 | Acyl carrier protein OS=Chlamydomonas reinhardtii OX=3055 GN=ACP2 PE=2 SV=1 |
| 53 | 5745 | tr\|A8J1U1\|A8J1U1\_CHLRE | 53.87 | 2 | 2 | 0 | 1 | 1 | 1 | N | 80682 | Heat shock protein 90A OS=Chlamydomonas reinhardtii OX=3055 GN=HSP90A PE=3 SV=1 |
| 54 | 5746 | tr\|Q9LD42\|Q9LD42\_CHLRE | 53.09 | 4 | 4 | 6.5093E4 | 1 | 1 | 1 | N | 37941 | Fe-assimilating protein 1 OS=Chlamydomonas reinhardtii OX=3055 GN=h43 PE=2 SV=1 |
| 55 | 5747 | tr\|A8HVQ1\|A8HVQ1\_CHLRE | 53.07 | 6 | 6 | 1.0858E5 | 1 | 1 | 1 | N | 23896 | 40S ribosomal protein S8 OS=Chlamydomonas reinhardtii OX=3055 GN=RPS8 PE=3 SV=1 |
| 36 | 5736 | tr\|A0A2K3E7M7\|A0A2K3E7M7\_CHLRE | 51.60 | 7 | 7 | 3.8935E5 | 4 | 4 | 4 | Y | 143415 | SMP-LTD domain-containing protein OS=Chlamydomonas reinhardtii OX=3055 GN=CHLRE\_01g044600v5 PE=3 SV=1 |
| 56 | 27 | tr\|A0A2K3CRJ3\|A0A2K3CRJ3\_CHLRE | 50.86 | 6 | 6 | 7.1637E5 | 1 | 1 | 1 | N | 19601 | Uncharacterized protein OS=Chlamydomonas reinhardtii OX=3055 GN=CHLRE\_17g738000v5 PE=4 SV=1 |
| 56 | 26 | tr\|Q1ALD7\|Q1ALD7\_CHLRE | 50.86 | 6 | 6 | 7.1637E5 | 1 | 1 | 1 | N | 19603 | AGG2 OS=Chlamydomonas reinhardtii OX=3055 GN=CHLRE\_17g738000v5 PE=2 SV=1 |
| 56 | 28 | tr\|A0A2K3CRI8\|A0A2K3CRI8\_CHLRE | 50.86 | 5 | 5 | 7.1637E5 | 1 | 1 | 1 | N | 21866 | Uncharacterized protein OS=Chlamydomonas reinhardtii OX=3055 GN=CHLRE\_17g738000v5 PE=4 SV=1 |
| 35 | 511 | tr\|A0A2K3DMT2\|A0A2K3DMT2\_CHLRE | 50.67 | 1 | 1 | 0 | 1 | 1 | 1 | N | 322939 | Uncharacterized protein OS=Chlamydomonas reinhardtii OX=3055 GN=CHLRE\_06g263250v5 PE=4 SV=1 |
| 57 | 5748 | tr\|A8J1A3\|A8J1A3\_CHLRE | 49.93 | 8 | 8 | 2.2313E5 | 1 | 1 | 1 | N | 17036 | Ribosomal protein L24 OS=Chlamydomonas reinhardtii OX=3055 GN=RPL24 PE=3 SV=1 |
| 57 | 5749 | tr\|A0A2K3DE61\|A0A2K3DE61\_CHLRE | 49.93 | 7 | 7 | 2.2313E5 | 1 | 1 | 1 | N | 19704 | Uncharacterized protein OS=Chlamydomonas reinhardtii OX=3055 GN=CHLRE\_09g391097v5 PE=3 SV=1 |
| 58 | 5750 | tr\|A0A2K3DZB3\|A0A2K3DZB3\_CHLRE | 43.07 | 4 | 4 | 0 | 1 | 1 | 1 | N | 39841 | Uncharacterized protein OS=Chlamydomonas reinhardtii OX=3055 GN=CHLRE\_03g202950v5 PE=4 SV=1 |
| 59 | 5751 | tr\|A8IV40\|A8IV40\_CHLRE | 42.06 | 11 | 11 | 0 | 1 | 1 | 1 | Y | 13232 | Ferredoxin OS=Chlamydomonas reinhardtii OX=3055 GN=PETF PE=3 SV=1 |
| 45 | 5741 | tr\|A0A2K3D7P8\|A0A2K3D7P8\_CHLRE | 41.48 | 1 | 1 | 1.4618E4 | 1 | 1 | 1 | Y | 158774 | Uncharacterized protein OS=Chlamydomonas reinhardtii OX=3055 GN=CHLRE\_11g467691v5 PE=3 SV=1 |
| 48 | 5753 | tr\|A0A2K3CPA4\|A0A2K3CPA4\_CHLRE | 39.81 | 8 | 8 | 8.3093E3 | 1 | 1 | 2 | N | 18371 | Uncharacterized protein OS=Chlamydomonas reinhardtii OX=3055 GN=CHLRE\_17g706600v5 PE=4 SV=1 |
| 37 | 5732 | tr\|O48949\|O48949\_CHLRE | 38.48 | 14 | 14 | 2.6802E5 | 5 | 5 | 5 | N | 58237 | Protein disulfide-isomerase OS=Chlamydomonas reinhardtii OX=3055 GN=PDI PE=2 SV=1 |
| 60 | 735 | tr\|A8HMY6\|A8HMY6\_CHLRE | 36.79 | 5 | 5 | 2.2058E5 | 1 | 1 | 1 | N | 18986 | Predicted protein OS=Chlamydomonas reinhardtii OX=3055 GN=CHLRE\_01g045200v5 PE=4 SV=1 |
| 49 | 5754 | tr\|A0A2K3E3X9\|A0A2K3E3X9\_CHLRE | 36.18 | 4 | 4 | 8.6069E4 | 1 | 1 | 1 | N | 35776 | Uncharacterized protein OS=Chlamydomonas reinhardtii OX=3055 GN=CHLRE\_02g144005v5 PE=3 SV=1 |
| 61 | 223 | tr\|A8IXZ0\|A8IXZ0\_CHLRE | 35.27 | 2 | 2 | 9.359E4 | 1 | 1 | 1 | N | 49619 | Tubulin beta chain OS=Chlamydomonas reinhardtii OX=3055 GN=TUB1 PE=3 SV=1 |
| 47 | 564 | tr\|A0A2K3DFK4\|A0A2K3DFK4\_CHLRE | 32.97 | 2 | 2 | 5.7435E6 | 2 | 2 | 2 | N | 211099 | SWIRM domain-containing protein OS=Chlamydomonas reinhardtii OX=3055 GN=CHLRE\_09g410300v5 PE=3 SV=1 |
| 39 | 1031 | tr\|A0A2K3D7S0\|A0A2K3D7S0\_CHLRE | 32.62 | 5 | 5 | 8.5138E4 | 1 | 1 | 2 | N | 81200 | Histone deacetylase OS=Chlamydomonas reinhardtii OX=3055 GN=CHLRE\_11g467706v5 PE=4 SV=1 |
| 50 | 989 | tr\|A0A2K3E270\|A0A2K3E270\_CHLRE | 32.58 | 2 | 2 | 0 | 1 | 1 | 1 | N | 144274 | Protein kinase domain-containing protein OS=Chlamydomonas reinhardtii OX=3055 GN=CHLRE\_02g098600v5 PE=4 SV=1 |
| 50 | 990 | tr\|A0A2K3E262\|A0A2K3E262\_CHLRE | 32.58 | 2 | 2 | 0 | 1 | 1 | 1 | N | 144513 | Protein kinase domain-containing protein OS=Chlamydomonas reinhardtii OX=3055 GN=CHLRE\_02g098600v5 PE=4 SV=1 |
| 62 | 5757 | tr\|A0A2K3CWQ4\|A0A2K3CWQ4\_CHLRE | 31.86 | 1 | 1 | 0 | 1 | 1 | 1 | N | 116027 | RAP domain-containing protein OS=Chlamydomonas reinhardtii OX=3055 GN=CHLRE\_15g639308v5 PE=4 SV=1 |
| total 71 proteins |  |  |  |  |  |  |  |  |  |  |  |  |
| --- | --- | --- | --- | --- | --- | --- | --- | --- | --- | --- | --- | --- |

  

tr|A0A2K3D8I0|A0A2K3D8I0\_CHLRE

back to list

  

| Protein Coverage
| Supporting Peptides
|

Protein Coverage:

Supporting Peptides:

| Peptide | Uniq | -10lgP | Mass | Length | ppm | m/z | z | RT | Fraction | Scan | Source File | Area Sample 2 | #Feature | #Feature Sample 2 | Start | End | PTM | AScore | Found By |
| --- | --- | --- | --- | --- | --- | --- | --- | --- | --- | --- | --- | --- | --- | --- | --- | --- | --- | --- | --- |
| K.AAAAASLGQAQDAAAQAQNTAAAAVGNAQTAAGDAAGK.A | Y | 103.37 | 3323.6089 | 38 | 2.7 | 1108.8799 | 3 | 29.67 | 2 | 6748 | JVDM\_TOPE.raw | 2.7853E5 | 2 | 2 | 792 | 829 |  |  | PEAKS DB |
| K.AQGAATAATEQAKDAAAAAQQQAAAAADAAK.G | Y | 88.38 | 2811.3745 | 31 | 1.4 | 938.1334 | 3 | 25.47 | 2 | 5780 | JVDM\_TOPE.raw | 1.8533E5 | 1 | 1 | 852 | 882 |  |  | PEAKS DB |
| K.AAADGAVANAQGAVTSFATK.A | Y | 88.29 | 1819.9064 | 20 | 3.7 | 910.9639 | 2 | 24.13 | 2 | 5430 | JVDM\_TOPE.raw | 1.9643E5 | 2 | 2 | 661 | 680 |  |  | PEAKS DB |
| K.GAAASAVGQATAQATAAVDQAK.A | Y | 88.08 | 1956.9865 | 22 | 2.7 | 979.5032 | 2 | 24.44 | 2 | 5530 | JVDM\_TOPE.raw | 8.8837E5 | 2 | 2 | 883 | 904 |  |  | PEAKS DB |
| K.AQETAAAAAGQAQAAAADAAAK.A | Y | 83.85 | 1926.9395 | 22 | 1.0 | 964.4780 | 2 | 19.45 | 2 | 4039 | JVDM\_TOPE.raw | 3.7236E5 | 1 | 1 | 830 | 851 |  |  | PEAKS DB |
| K.AGEVQAAAAGAAEQAQAQASAAAAEAQK.A | Y | 82.07 | 2539.2261 | 28 | 1.6 | 847.4174 | 3 | 26.31 | 2 | 5983 | JVDM\_TOPE.raw | 0 | 0 | 0 | 616 | 643 |  |  | PEAKS DB |
| K.AAEAQAAAAGAAEQAQAAAANAQQAAADAAGK.A | Y | 75.24 | 2808.3386 | 32 | 1.6 | 937.1216 | 3 | 28.56 | 2 | 6471 | JVDM\_TOPE.raw | 7.6782E4 | 1 | 1 | 511 | 542 |  |  | PEAKS DB |
| K.AASAAAEQAQAAAVGAQQAAVEK.A | Y | 69.44 | 2111.0605 | 23 | 2.6 | 704.6959 | 3 | 19.48 | 2 | 4036 | JVDM\_TOPE.raw | 2.4879E5 | 1 | 1 | 575 | 597 |  |  | PEAKS DB |
| K.ADDVAVQVGATAAGAAGAKPPAGVPK.K | Y | 66.09 | 2288.2124 | 26 | 1.3 | 763.7457 | 3 | 22.72 | 2 | 5054 | JVDM\_TOPE.raw | 5.1235E4 | 1 | 1 | 973 | 998 |  |  | PEAKS DB |
| K.DAAAAAQQQAAAAADAAK.G | Y | 65.14 | 1612.7804 | 18 | 5.6 | 807.4020 | 2 | 17.20 | 2 | 3292 | JVDM\_TOPE.raw | 1.1098E5 | 1 | 1 | 865 | 882 |  |  | PEAKS DB |
| K.AEDTAAAAQAQAAAAAESAR.G | Y | 59.62 | 1843.8660 | 20 | 3.0 | 615.6311 | 3 | 18.73 | 2 | 3783 | JVDM\_TOPE.raw | 1.5539E5 | 1 | 1 | 681 | 700 |  |  | PEAKS DB |
| K.ASEAATGALSEIAAVDGEGGAAK.A | Y | 58.95 | 2044.9912 | 23 | -0.2 | 1023.5027 | 2 | 27.55 | 2 | 6258 | JVDM\_TOPE.raw | 1.635E4 | 1 | 1 | 950 | 972 |  |  | PEAKS DB |
| K.AAAAQATAAAAGAQEAASAAAGQATATATDAMTK.A | Y | 58.65 | 2961.4097 | 34 | 1.9 | 988.1458 | 3 | 30.05 | 2 | 6837 | JVDM\_TOPE.raw | 1.6568E5 | 1 | 1 | 756 | 789 |  |  | PEAKS DB |
| K.AAAASAQGALAAATGQAR.D | Y | 58.56 | 1555.8066 | 18 | 1.2 | 778.9115 | 2 | 19.92 | 2 | 4157 | JVDM\_TOPE.raw | 1.5906E5 | 1 | 1 | 905 | 922 |  |  | PEAKS DB |
| R.GQAEAAAAQAQNTAAAAVGNVQTAAADAAAK.A | Y | 57.63 | 2781.3640 | 31 | 3.1 | 928.1315 | 3 | 29.63 | 2 | 6727 | JVDM\_TOPE.raw | 2.4063E5 | 1 | 1 | 701 | 731 |  |  | PEAKS DB |
| A.KAGEVQAAAAGAAEQAQAQASAAAAEAQK.A | Y | 53.69 | 2667.3210 | 29 | 2.4 | 890.1164 | 3 | 24.59 | 2 | 5570 | JVDM\_TOPE.raw | 0 | 0 | 0 | 615 | 643 |  |  | PEAKS DB |
| K.AAESAAASAAQAQATAASAVSAVQDK.A | Y | 31.28 | 2345.1458 | 26 | 3.1 | 782.7250 | 3 | 25.94 | 2 | 5890 | JVDM\_TOPE.raw | 0 | 0 | 0 | 438 | 463 |  |  | PEAKS DB |
| total 17 peptides |  |  |  |  |  |  |  |  |  |  |  |  |  |  |  |  |  |  |  |
| --- | --- | --- | --- | --- | --- | --- | --- | --- | --- | --- | --- | --- | --- | --- | --- | --- | --- | --- | --- |

PHL7

back to list

  

| Protein Coverage
| Supporting Peptides
|

Protein Coverage:

Supporting Peptides:

| Peptide | Uniq | -10lgP | Mass | Length | ppm | m/z | z | RT | Fraction | Scan | Source File | Area Sample 2 | #Feature | #Feature Sample 2 | Start | End | PTM | AScore | Found By |
| --- | --- | --- | --- | --- | --- | --- | --- | --- | --- | --- | --- | --- | --- | --- | --- | --- | --- | --- | --- |
| R.GPDPTESSIEAVR.G | Y | 94.35 | 1356.6521 | 13 | 1.4 | 679.3343 | 2 | 16.45 | 2 | 3054 | JVDM\_TOPE.raw | 5.6779E6 | 1 | 1 | 8 | 20 |  |  | PEAKS DB |
| R.IASQGFVVITIDTITR.L | Y | 92.13 | 1732.9723 | 16 | 2.5 | 867.4956 | 2 | 38.43 | 2 | 8913 | JVDM\_TOPE.raw | 2.0567E6 | 2 | 2 | 77 | 92 |  |  | PEAKS DB |
| R.TPTLVVGAQLDTIAPVSSHSEAFYNSLPSDLDK.A | Y | 85.83 | 3471.7410 | 33 | 2.2 | 868.9445 | 4 | 36.17 | 2 | 8332 | JVDM\_TOPE.raw | 1.2395E6 | 2 | 2 | 167 | 199 |  |  | PEAKS DB |
| R.GPFAVAQTTVSR.L | Y | 79.32 | 1232.6514 | 12 | 2.1 | 617.3342 | 2 | 20.39 | 2 | 4297 | JVDM\_TOPE.raw | 1.0429E7 | 1 | 1 | 21 | 32 |  |  | PEAKS DB |
| R.FVDDDLRYEQFLC(+57.02)PAPDDFAISEYR.S | Y | 69.77 | 3080.3860 | 25 | 2.3 | 1027.8049 | 3 | 36.44 | 2 | 8410 | JVDM\_TOPE.raw | 8.4096E4 | 1 | 1 | 230 | 254 | Carbamidomethylation | C13:Carbamidomethylation:1000.00 | PEAKS DB |
| R.YEQFLC(+57.02)PAPDDFAISEYR.S | Y | 65.79 | 2219.9834 | 18 | 1.1 | 1111.0002 | 2 | 33.50 | 2 | 7663 | JVDM\_TOPE.raw | 4.5602E5 | 2 | 2 | 237 | 254 | Carbamidomethylation | C6:Carbamidomethylation:1000.00 | PEAKS DB |
| R.GPDPTESSIEAVRGPFAVAQTTVSR.L | Y | 62.50 | 2571.2927 | 25 | 2.4 | 858.1069 | 3 | 28.89 | 2 | 6569 | JVDM\_TOPE.raw | 6.2711E6 | 1 | 1 | 8 | 32 |  |  | PEAKS DB |
| R.TPTLVVGAQ(+.98)LDTIAPVSSHSEAFYN(+.98)SLPSDLDK.A | Y | 42.94 | 3473.7090 | 33 | 8.8 | 1158.9204 | 3 | 35.45 | 2 | 8142 | JVDM\_TOPE.raw | 2.0264E4 | 1 | 1 | 167 | 199 | Deamidation (NQ) | Q9:Deamidation (NQ):1000.00;N25:Deamidation (NQ):1000.00 | PEAKS PTM |
| R.GPDPTESSIEAVRGPF.A | Y | 39.19 | 1657.7947 | 16 | 3.1 | 829.9072 | 2 | 27.62 | 2 | 6270 | JVDM\_TOPE.raw | 1.4792E4 | 1 | 1 | 8 | 23 |  |  | PEAKS DB |
| R.QLQAALDHLR.T | Y | 38.01 | 1163.6411 | 10 | 1.2 | 388.8881 | 3 | 19.33 | 2 | 3981 | JVDM\_TOPE.raw | 5.9083E6 | 2 | 2 | 102 | 111 |  |  | PEAKS DB |
| R.FVDDDLRYEQ(+.98)FLC(+57.02)PAPDDFAISEYR.S | Y | 32.19 | 3081.3701 | 25 | 9.0 | 1028.1399 | 3 | 36.48 | 2 | 8419 | JVDM\_TOPE.raw | 0 | 0 | 0 | 230 | 254 | Deamidation (NQ); Carbamidomethylation | Q10:Deamidation (NQ):1000.00;C13:Carbamidomethylation:1000.00 | PEAKS PTM |
| K.YSIAWLK.R | Y | 27.85 | 879.4854 | 7 | 2.3 | 440.7510 | 2 | 29.00 | 2 | 6577 | JVDM\_TOPE.raw | 1.4417E6 | 1 | 1 | 222 | 228 |  |  | PEAKS DB |
| R.FVDDDLR.Y | Y | 21.41 | 878.4133 | 7 | 0.5 | 440.2142 | 2 | 15.82 | 2 | 2905 | JVDM\_TOPE.raw | 1.1021E6 | 1 | 1 | 230 | 236 |  |  | PEAKS DB |
| total 13 peptides |  |  |  |  |  |  |  |  |  |  |  |  |  |  |  |  |  |  |  |
| --- | --- | --- | --- | --- | --- | --- | --- | --- | --- | --- | --- | --- | --- | --- | --- | --- | --- | --- | --- |

tr|A8J6Y8|A8J6Y8\_CHLRE

back to list

  

| Protein Coverage
| Supporting Peptides
|

Protein Coverage:

Supporting Peptides:

| Peptide | Uniq | -10lgP | Mass | Length | ppm | m/z | z | RT | Fraction | Scan | Source File | Area Sample 2 | #Feature | #Feature Sample 2 | Start | End | PTM | AScore | Found By |
| --- | --- | --- | --- | --- | --- | --- | --- | --- | --- | --- | --- | --- | --- | --- | --- | --- | --- | --- | --- |
| K.LYDEELQAIAK.A | Y | 88.34 | 1291.6659 | 11 | 1.5 | 646.8412 | 2 | 24.65 | 2 | 5557 | JVDM\_TOPE.raw | 7.0147E6 | 1 | 1 | 244 | 254 |  |  | PEAKS DB |
| R.YGDDFDGQTASLC(+57.02)VR.R | Y | 85.27 | 1702.7257 | 15 | 2.0 | 852.3718 | 2 | 22.58 | 2 | 5035 | JVDM\_TOPE.raw | 4.1596E6 | 2 | 2 | 131 | 145 | Carbamidomethylation | C13:Carbamidomethylation:1000.00 | PEAKS DB |
| K.ATGETC(+57.02)HIIIETEGK.I | Y | 79.08 | 1657.7981 | 15 | 2.3 | 829.9082 | 2 | 16.76 | 2 | 3147 | JVDM\_TOPE.raw | 2.7798E6 | 2 | 2 | 78 | 92 | Carbamidomethylation | C6:Carbamidomethylation:1000.00 | PEAKS DB |
| R.TVPTKLEEGEMPLNTYSNK.A | Y | 75.40 | 2150.0564 | 19 | 2.4 | 1076.0381 | 2 | 22.50 | 2 | 4981 | JVDM\_TOPE.raw | 3.1033E6 | 2 | 2 | 42 | 60 |  |  | PEAKS DB |
| K.GLC(+57.02)SNFLC(+57.02)DATPGTEISMTGPTGK.V | Y | 72.12 | 2513.1235 | 24 | 4.8 | 1257.5751 | 2 | 31.38 | 2 | 7134 | JVDM\_TOPE.raw | 5.3155E5 | 1 | 1 | 163 | 186 | Carbamidomethylation | C3:Carbamidomethylation:1000.00;C8:Carbamidomethylation:1000.00 | PEAKS DB |
| K.LEEGEMPLNTYSNK.A | Y | 54.26 | 1623.7450 | 14 | 3.6 | 812.8827 | 2 | 20.86 | 2 | 4479 | JVDM\_TOPE.raw | 4.0525E4 | 1 | 1 | 47 | 60 |  |  | PEAKS DB |
| K.GMMPGIQDMLER.V | Y | 46.49 | 1376.6250 | 12 | 2.2 | 689.3213 | 2 | 32.39 | 2 | 7322 | JVDM\_TOPE.raw | 1.8587E6 | 1 | 1 | 308 | 319 |  |  | PEAKS DB |
| R.LYSIASSR.Y | Y | 40.32 | 895.4763 | 8 | 1.3 | 448.7460 | 2 | 15.18 | 2 | 2710 | JVDM\_TOPE.raw | 5.6978E6 | 1 | 1 | 123 | 130 |  |  | PEAKS DB |
| R.TVPTKLEEGEMPLN(+.98)TYSNK.A | Y | 38.40 | 2151.0405 | 19 | 2.7 | 718.0227 | 3 | 22.37 | 2 | 4934 | JVDM\_TOPE.raw | 1.5243E5 | 1 | 1 | 42 | 60 | Deamidation (NQ) | N14:Deamidation (NQ):34.30 | PEAKS PTM |
| R.LDYALSR.E | Y | 36.43 | 836.4392 | 7 | 1.4 | 419.2274 | 2 | 16.44 | 2 | 2991 | JVDM\_TOPE.raw | 5.1405E6 | 1 | 1 | 262 | 268 |  |  | PEAKS DB |
| K.VLLLPADANAPLIC(+57.02)VATGTGIAPFR.S | Y | 35.35 | 2549.4038 | 25 | 1.7 | 850.8100 | 3 | 40.57 | 2 | 9488 | JVDM\_TOPE.raw | 0 | 0 | 0 | 187 | 211 | Carbamidomethylation | C14:Carbamidomethylation:1000.00 | PEAKS DB |
| K.NQWHVEVY | Y | 33.37 | 1073.4930 | 8 | 1.0 | 537.7543 | 2 | 24.02 | 2 | 5392 | JVDM\_TOPE.raw | 6.8438E5 | 1 | 1 | 339 | 346 |  |  | PEAKS DB |
| K.ASTAVTTDMSKR.T | Y | 24.67 | 1266.6238 | 12 | 1.3 | 423.2158 | 3 | 12.41 | 2 | 1486 | JVDM\_TOPE.raw | 1.9971E6 | 1 | 1 | 30 | 41 |  |  | PEAKS DB |
| K.AYPGQFR.L | Y | 20.80 | 837.4133 | 7 | 0.4 | 419.7141 | 2 | 13.37 | 2 | 2237 | JVDM\_TOPE.raw | 4.4914E6 | 1 | 1 | 255 | 261 |  |  | PEAKS DB |
| K.GLNYEEWVEGLK.H | Y | 17.66 | 1435.6982 | 12 | 2.1 | 718.8579 | 2 | 32.93 | 2 | 7460 | JVDM\_TOPE.raw | 7.2447E5 | 1 | 1 | 325 | 336 |  |  | PEAKS DB |
| K.IPFWEGQSYGVIPPGTK(+14.02)IN(+.98)SK(+28.03).G | Y | 15.25 | 2360.2415 | 21 | 5.1 | 787.7584 | 3 | 32.06 | 2 | 7266 | JVDM\_TOPE.raw | 2.2242E5 | 1 | 1 | 93 | 113 | Methylation(KR); Deamidation (NQ); Dimethylation(KR) | K17:Methylation(KR):2.76;N19:Deamidation (NQ):47.33;K21:Dimethylation(KR):2.76 | PEAKS PTM |
| K.RTVPTKLEEGEMPLNTYSNK.A | Y | 15.19 | 2306.1577 | 20 | 3.2 | 769.7290 | 3 | 20.00 | 2 | 4235 | JVDM\_TOPE.raw | 1.1472E4 | 1 | 1 | 41 | 60 |  |  | PEAKS DB |
| total 17 peptides |  |  |  |  |  |  |  |  |  |  |  |  |  |  |  |  |  |  |  |
| --- | --- | --- | --- | --- | --- | --- | --- | --- | --- | --- | --- | --- | --- | --- | --- | --- | --- | --- | --- |

tr|A0A2K3CTJ7|A0A2K3CTJ7\_CHLRE

back to list

  

| Protein Coverage
| Supporting Peptides
|

Protein Coverage:

Supporting Peptides:

| Peptide | Uniq | -10lgP | Mass | Length | ppm | m/z | z | RT | Fraction | Scan | Source File | Area Sample 2 | #Feature | #Feature Sample 2 | Start | End | PTM | AScore | Found By |
| --- | --- | --- | --- | --- | --- | --- | --- | --- | --- | --- | --- | --- | --- | --- | --- | --- | --- | --- | --- |
| K.TVAALYAPSAVLLPTVSNQYR.T | Y | 93.33 | 2233.2107 | 21 | 1.3 | 745.4118 | 3 | 36.17 | 2 | 8323 | JVDM\_TOPE.raw | 2.7241E5 | 2 | 2 | 115 | 135 |  |  | PEAKS DB |
| K.VASLYAPTAVLLPTVSDK.V | Y | 79.46 | 1844.0294 | 18 | 0.9 | 923.0228 | 2 | 35.05 | 2 | 8049 | JVDM\_TOPE.raw | 2.4136E5 | 2 | 2 | 241 | 258 |  |  | PEAKS DB |
| K.EPVGVIDNIDGSSPNVR.F | Y | 77.68 | 1766.8799 | 17 | 2.3 | 884.4492 | 2 | 23.49 | 2 | 5258 | JVDM\_TOPE.raw | 2.2935E5 | 2 | 2 | 277 | 293 |  |  | PEAKS DB |
| K.GC(+57.02)PALTVNAVDSYNK.A | Y | 70.75 | 1607.7614 | 15 | 2.0 | 804.8895 | 2 | 21.59 | 2 | 4689 | JVDM\_TOPE.raw | 1.6598E5 | 1 | 1 | 33 | 47 | Carbamidomethylation | C2:Carbamidomethylation:1000.00 | PEAKS DB |
| K.VASLYAPTAVLLPTVSDKVR.N | Y | 61.16 | 2099.1990 | 20 | 2.4 | 700.7419 | 3 | 33.57 | 2 | 7687 | JVDM\_TOPE.raw | 2.5472E5 | 1 | 1 | 241 | 260 |  |  | PEAKS DB |
| K.GTFLDSDC(+57.02)FDQVGTK.G | Y | 53.49 | 1688.7352 | 15 | 2.4 | 845.3770 | 2 | 26.93 | 2 | 6118 | JVDM\_TOPE.raw | 7.5896E4 | 1 | 1 | 50 | 64 | Carbamidomethylation | C8:Carbamidomethylation:1000.00 | PEAKS DB |
| K.GPVGAIDKYEIR.H | Y | 45.64 | 1316.7087 | 12 | 2.3 | 439.9112 | 3 | 19.99 | 2 | 4239 | JVDM\_TOPE.raw | 3.6154E5 | 1 | 1 | 152 | 163 |  |  | PEAKS DB |
| K.KEPVGVIDNIDGSSPNVR.F | Y | 27.66 | 1894.9749 | 18 | 1.9 | 632.6667 | 3 | 21.03 | 2 | 4493 | JVDM\_TOPE.raw | 1.972E5 | 1 | 1 | 276 | 293 |  |  | PEAKS DB |
| total 8 peptides |  |  |  |  |  |  |  |  |  |  |  |  |  |  |  |  |  |  |  |
| --- | --- | --- | --- | --- | --- | --- | --- | --- | --- | --- | --- | --- | --- | --- | --- | --- | --- | --- | --- |

tr|A0A2K3D1P1|A0A2K3D1P1\_CHLRE

back to list

  

| Protein Coverage
| Supporting Peptides
|

Protein Coverage:

Supporting Peptides:

| Peptide | Uniq | -10lgP | Mass | Length | ppm | m/z | z | RT | Fraction | Scan | Source File | Area Sample 2 | #Feature | #Feature Sample 2 | Start | End | PTM | AScore | Found By |
| --- | --- | --- | --- | --- | --- | --- | --- | --- | --- | --- | --- | --- | --- | --- | --- | --- | --- | --- | --- |
| K.MNSQVSSLSLYDIAGTPGVAADVSHINTK.A | Y | 85.00 | 2974.4705 | 29 | 2.3 | 992.4997 | 3 | 32.93 | 2 | 7484 | JVDM\_TOPE.raw | 0 | 0 | 0 | 52 | 80 |  |  | PEAKS DB |
| K.VAVLGAAGGIGQPLSMLMK.M | Y | 83.13 | 1812.0001 | 19 | 0.4 | 907.0077 | 2 | 39.35 | 2 | 9183 | JVDM\_TOPE.raw | 3.8277E4 | 1 | 1 | 33 | 51 |  |  | PEAKS DB |
| R.GLN(+.98)GAPVVEC(+57.02)TYVESTVTDAPYFASK.V | Y | 77.00 | 2775.2949 | 26 | 0.0 | 926.1056 | 3 | 32.93 | 2 | 7490 | JVDM\_TOPE.raw | 9.5273E4 | 1 | 1 | 270 | 295 | Deamidation (NQ); Carbamidomethylation | N3:Deamidation (NQ):1000.00;C10:Carbamidomethylation:1000.00 | PEAKS PTM |
| K.ATMSAEVLDALTK.R | Y | 56.78 | 1348.6908 | 13 | 2.7 | 675.3545 | 2 | 32.06 | 2 | 7282 | JVDM\_TOPE.raw | 1.097E5 | 1 | 1 | 221 | 233 |  |  | PEAKS DB |
| R.GC(+57.02)DLVIIPAGVPR.K | Y | 52.13 | 1365.7438 | 13 | 3.0 | 683.8812 | 2 | 30.51 | 2 | 6978 | JVDM\_TOPE.raw | 1.5891E5 | 1 | 1 | 97 | 109 | Carbamidomethylation | C2:Carbamidomethylation:1000.00 | PEAKS DB |
| K.GFDKDGLAEALR.G | Y | 49.19 | 1290.6567 | 12 | 1.7 | 646.3367 | 2 | 23.06 | 2 | 5149 | JVDM\_TOPE.raw | 3.9778E4 | 1 | 1 | 85 | 96 |  |  | PEAKS DB |
| K.AMMPELLASIEK.G | Y | 46.67 | 1331.6829 | 12 | 1.9 | 666.8500 | 2 | 35.31 | 2 | 8120 | JVDM\_TOPE.raw | 1.3983E5 | 1 | 1 | 322 | 333 |  |  | PEAKS DB |
| K.MNSQ(+.98)VSSLSLYDIAGTPGVAADVSHINTK.A | Y | 40.55 | 2975.4546 | 29 | 6.9 | 992.8323 | 3 | 32.87 | 2 | 7484 | JVDM\_TOPE.raw | 9.3073E4 | 1 | 1 | 52 | 80 | Deamidation (NQ) | Q4:Deamidation (NQ):26.31 | PEAKS PTM |
| total 8 peptides |  |  |  |  |  |  |  |  |  |  |  |  |  |  |  |  |  |  |  |
| --- | --- | --- | --- | --- | --- | --- | --- | --- | --- | --- | --- | --- | --- | --- | --- | --- | --- | --- | --- |

tr|A0A2K3E094|A0A2K3E094\_CHLRE

back to list

  

| Protein Coverage
| Supporting Peptides
|

Protein Coverage:

Supporting Peptides:

| Peptide | Uniq | -10lgP | Mass | Length | ppm | m/z | z | RT | Fraction | Scan | Source File | Area Sample 2 | #Feature | #Feature Sample 2 | Start | End | PTM | AScore | Found By |
| --- | --- | --- | --- | --- | --- | --- | --- | --- | --- | --- | --- | --- | --- | --- | --- | --- | --- | --- | --- |
| R.HEDVEAFTIPLYTAAGDETK.V | Y | 100.81 | 2206.0430 | 20 | 2.6 | 736.3568 | 3 | 31.98 | 2 | 7258 | JVDM\_TOPE.raw | 3.5019E5 | 1 | 1 | 185 | 204 |  |  | PEAKS DB |
| K.SSQGVPLLAEQASSFFSSK.A | N | 76.47 | 1968.9792 | 19 | 0.6 | 657.3341 | 3 | 36.98 | 2 | 8537 | JVDM\_TOPE.raw | 1.1703E6 | 2 | 2 | 144 | 162 |  |  | PEAKS DB |
| K.QLVQAGFDLK.V | Y | 54.48 | 1117.6132 | 10 | 1.0 | 559.8144 | 2 | 25.03 | 2 | 5676 | JVDM\_TOPE.raw | 6.3693E5 | 1 | 1 | 54 | 63 |  |  | PEAKS DB |
| K.AQDAWPASLEDAK.T | N | 50.77 | 1400.6571 | 13 | 1.5 | 701.3369 | 2 | 23.58 | 2 | 5316 | JVDM\_TOPE.raw | 8.6981E5 | 1 | 1 | 163 | 175 |  |  | PEAKS DB |
| K.TISFNTWC(+57.02)R.H | N | 33.40 | 1183.5444 | 9 | 2.4 | 592.7809 | 2 | 25.61 | 2 | 5813 | JVDM\_TOPE.raw | 2.8165E5 | 1 | 1 | 176 | 184 | Carbamidomethylation | C8:Carbamidomethylation:1000.00 | PEAKS DB |
| R.WGELLEK.L | N | 26.48 | 873.4596 | 7 | 0.8 | 437.7374 | 2 | 23.13 | 2 | 5157 | JVDM\_TOPE.raw | 5.748E5 | 1 | 1 | 234 | 240 |  |  | PEAKS DB |
| K.ALGYQGSWSAC(+57.02)K.H | Y | 22.51 | 1326.6027 | 12 | 1.8 | 664.3098 | 2 | 19.45 | 2 | 4018 | JVDM\_TOPE.raw | 1.6304E5 | 1 | 1 | 125 | 136 | Carbamidomethylation | C11:Carbamidomethylation:1000.00 | PEAKS DB |
| total 7 peptides |  |  |  |  |  |  |  |  |  |  |  |  |  |  |  |  |  |  |  |
| --- | --- | --- | --- | --- | --- | --- | --- | --- | --- | --- | --- | --- | --- | --- | --- | --- | --- | --- | --- |

tr|A0A2K3D6N5|A0A2K3D6N5\_CHLRE

back to list

  

| Protein Coverage
| Supporting Peptides
|

Protein Coverage:

Supporting Peptides:

| Peptide | Uniq | -10lgP | Mass | Length | ppm | m/z | z | RT | Fraction | Scan | Source File | Area Sample 2 | #Feature | #Feature Sample 2 | Start | End | PTM | AScore | Found By |
| --- | --- | --- | --- | --- | --- | --- | --- | --- | --- | --- | --- | --- | --- | --- | --- | --- | --- | --- | --- |
| R.VVIALDSSSHSGADFAAK.A | Y | 82.15 | 1773.8896 | 18 | 1.8 | 887.9537 | 2 | 22.35 | 2 | 4924 | JVDM\_TOPE.raw | 6.4382E5 | 1 | 1 | 102 | 119 |  |  | PEAKS DB |
| K.VGVAGAVVVLGYNWK.A | Y | 70.13 | 1530.8558 | 15 | 2.8 | 766.4373 | 2 | 34.89 | 2 | 8017 | JVDM\_TOPE.raw | 8.0497E4 | 1 | 1 | 204 | 218 |  |  | PEAKS DB |
| K.AAGDNAQAAAVPSVTEQASEFLAGSWPK.T | Y | 61.06 | 2772.3354 | 28 | 2.0 | 925.1210 | 3 | 38.36 | 2 | 8908 | JVDM\_TOPE.raw | 2.2897E5 | 1 | 1 | 132 | 159 |  |  | PEAKS DB |
| K.AGPQEQWGALLNK.I | Y | 51.84 | 1410.7256 | 13 | 1.6 | 706.3712 | 2 | 27.17 | 2 | 6147 | JVDM\_TOPE.raw | 3.855E5 | 1 | 1 | 219 | 231 |  |  | PEAKS DB |
| R.ITSASTVC(+57.02)SHEDSSAVVIPLYTYEDGEDVR.V | Y | 46.21 | 3299.5139 | 30 | 2.6 | 1100.8481 | 3 | 30.00 | 2 | 6845 | JVDM\_TOPE.raw | 6.1684E4 | 1 | 1 | 166 | 195 | Carbamidomethylation | C8:Carbamidomethylation:1000.00 | PEAKS DB |
| K.ALNYDGNWISC(+57.02)K.A | Y | 36.12 | 1439.6504 | 12 | 2.7 | 720.8344 | 2 | 24.27 | 2 | 5475 | JVDM\_TOPE.raw | 1.1377E5 | 1 | 1 | 120 | 131 | Carbamidomethylation | C11:Carbamidomethylation:1000.00 | PEAKS DB |
| R.VVAQAFGK.V | Y | 33.53 | 818.4650 | 8 | 0.3 | 410.2399 | 2 | 13.31 | 2 | 2086 | JVDM\_TOPE.raw | 9.4349E5 | 1 | 1 | 196 | 203 |  |  | PEAKS DB |
| total 7 peptides |  |  |  |  |  |  |  |  |  |  |  |  |  |  |  |  |  |  |  |
| --- | --- | --- | --- | --- | --- | --- | --- | --- | --- | --- | --- | --- | --- | --- | --- | --- | --- | --- | --- |

tr|A0A2K3E0A7|A0A2K3E0A7\_CHLRE

back to list

  

| Protein Coverage
| Supporting Peptides
|

Protein Coverage:

Supporting Peptides:

| Peptide | Uniq | -10lgP | Mass | Length | ppm | m/z | z | RT | Fraction | Scan | Source File | Area Sample 2 | #Feature | #Feature Sample 2 | Start | End | PTM | AScore | Found By |
| --- | --- | --- | --- | --- | --- | --- | --- | --- | --- | --- | --- | --- | --- | --- | --- | --- | --- | --- | --- |
| R.LTGTDAYVIPAQHGSAFYSSAEDMSAVSK.F | Y | 79.55 | 3002.3967 | 29 | 2.5 | 1001.8087 | 3 | 27.75 | 2 | 6285 | JVDM\_TOPE.raw | 7.6234E4 | 1 | 1 | 58 | 86 |  |  | PEAKS DB |
| K.SSQGVPLLAEQASSFFSSK.A | N | 76.47 | 1968.9792 | 19 | 0.6 | 657.3341 | 3 | 36.98 | 2 | 8537 | JVDM\_TOPE.raw | 1.1703E6 | 2 | 2 | 130 | 148 |  |  | PEAKS DB |
| K.FVEN(+.98)GGLVVLLGGK.A | Y | 51.27 | 1401.7867 | 14 | 1.9 | 701.9020 | 2 | 35.53 | 2 | 8156 | JVDM\_TOPE.raw | 1.1712E5 | 1 | 1 | 87 | 100 | Deamidation (NQ) | N4:Deamidation (NQ):1000.00 | PEAKS PTM |
| K.AQDAWPASLEDAK.T | N | 50.77 | 1400.6571 | 13 | 1.5 | 701.3369 | 2 | 23.58 | 2 | 5316 | JVDM\_TOPE.raw | 8.6981E5 | 1 | 1 | 149 | 161 |  |  | PEAKS DB |
| K.TISFNTWC(+57.02)R.H | N | 33.40 | 1183.5444 | 9 | 2.4 | 592.7809 | 2 | 25.61 | 2 | 5813 | JVDM\_TOPE.raw | 2.8165E5 | 1 | 1 | 162 | 170 | Carbamidomethylation | C8:Carbamidomethylation:1000.00 | PEAKS DB |
| R.LTGTDAYVIPAQ(+.98)HGSAFYSSAEDMSAVSK.F | Y | 32.18 | 3003.3806 | 29 | 6.6 | 1002.1407 | 3 | 27.78 | 2 | 6313 | JVDM\_TOPE.raw | 1.3806E5 | 1 | 1 | 58 | 86 | Deamidation (NQ) | Q12:Deamidation (NQ):1000.00 | PEAKS PTM |
| R.HEDVEAFTIPLYTAAGDEMK.V | N | 30.63 | 2236.0356 | 20 | 3.5 | 746.3551 | 3 | 33.87 | 2 | 7785 | JVDM\_TOPE.raw | 1.0031E5 | 1 | 1 | 171 | 190 |  |  | PEAKS DB |
| R.WGELLEK.L | N | 26.48 | 873.4596 | 7 | 0.8 | 437.7374 | 2 | 23.13 | 2 | 5157 | JVDM\_TOPE.raw | 5.748E5 | 1 | 1 | 220 | 226 |  |  | PEAKS DB |
| total 8 peptides |  |  |  |  |  |  |  |  |  |  |  |  |  |  |  |  |  |  |  |
| --- | --- | --- | --- | --- | --- | --- | --- | --- | --- | --- | --- | --- | --- | --- | --- | --- | --- | --- | --- |

tr|A0A2K3CQ54|A0A2K3CQ54\_CHLRE

back to list

  

| Protein Coverage
| Supporting Peptides
|

Protein Coverage:

Supporting Peptides:

| Peptide | Uniq | -10lgP | Mass | Length | ppm | m/z | z | RT | Fraction | Scan | Source File | Area Sample 2 | #Feature | #Feature Sample 2 | Start | End | PTM | AScore | Found By |
| --- | --- | --- | --- | --- | --- | --- | --- | --- | --- | --- | --- | --- | --- | --- | --- | --- | --- | --- | --- |
| K.VEYTELQILC(+57.02)PQTIDSVTGYPMDDPR.C | Y | 82.55 | 3039.4204 | 26 | 2.3 | 1520.7209 | 2 | 37.51 | 2 | 8687 | JVDM\_TOPE.raw | 1.2385E7 | 4 | 4 | 110 | 135 | Carbamidomethylation | C10:Carbamidomethylation:1000.00 | PEAKS DB |
| R.ATVAAGEEALTIR.N | Y | 75.53 | 1300.6986 | 13 | 1.8 | 651.3577 | 2 | 21.61 | 2 | 4773 | JVDM\_TOPE.raw | 2.6005E7 | 1 | 1 | 141 | 153 |  |  | PEAKS DB |
| R.ATVAAGEEALTIRN.E | Y | 41.65 | 1414.7416 | 14 | 1.9 | 708.3794 | 2 | 19.90 | 2 | 4202 | JVDM\_TOPE.raw | 6.8017E4 | 1 | 1 | 141 | 154 |  |  | PEAKS DB |
| L.QILC(+57.02)PQTIDSVTGYPMDDPR.C | Y | 38.02 | 2305.0718 | 20 | 1.9 | 769.3660 | 3 | 30.14 | 2 | 6859 | JVDM\_TOPE.raw | 2.2885E5 | 1 | 1 | 116 | 135 | Carbamidomethylation | C4:Carbamidomethylation:1000.00 | PEAKS DB |
| Q.TIDSVTGYPMDDPR.C | Y | 35.43 | 1565.7031 | 14 | 2.4 | 783.8607 | 2 | 21.42 | 2 | 4705 | JVDM\_TOPE.raw | 3.8917E5 | 1 | 1 | 122 | 135 |  |  | PEAKS DB |
| K.VEYTELQILCPQT(+79.97)IDSVTGYPMDDPR.C | Y | 33.45 | 3062.3655 | 26 | 5.8 | 766.6031 | 4 | 37.44 | 2 | 8686 | JVDM\_TOPE.raw | 0 | 0 | 0 | 110 | 135 | Phosphorylation (STY) | T13:Phosphorylation (STY):5.77 | PEAKS PTM |
| K.VEYTELQILC(+57.02)PQTIDSVTGYPMDD.P | Y | 21.84 | 2786.2666 | 24 | 2.0 | 929.7647 | 3 | 39.70 | 2 | 9259 | JVDM\_TOPE.raw | 6.4405E4 | 1 | 1 | 110 | 133 | Carbamidomethylation | C10:Carbamidomethylation:1000.00 | PEAKS DB |
| K.VEYTELQILC(+57.02)PQ.T | Y | 15.18 | 1491.7279 | 12 | 1.1 | 746.8721 | 2 | 32.68 | 2 | 7400 | JVDM\_TOPE.raw | 2.4394E5 | 1 | 1 | 110 | 121 | Carbamidomethylation | C10:Carbamidomethylation:1000.00 | PEAKS DB |
| total 8 peptides |  |  |  |  |  |  |  |  |  |  |  |  |  |  |  |  |  |  |  |
| --- | --- | --- | --- | --- | --- | --- | --- | --- | --- | --- | --- | --- | --- | --- | --- | --- | --- | --- | --- |

tr|P93106|P93106\_CHLRE

back to list

  

| Protein Coverage
| Supporting Peptides
|

Protein Coverage:

Supporting Peptides:

| Peptide | Uniq | -10lgP | Mass | Length | ppm | m/z | z | RT | Fraction | Scan | Source File | Area Sample 2 | #Feature | #Feature Sample 2 | Start | End | PTM | AScore | Found By |
| --- | --- | --- | --- | --- | --- | --- | --- | --- | --- | --- | --- | --- | --- | --- | --- | --- | --- | --- | --- |
| K.KVIGVTSLDIVR.A | Y | 64.27 | 1298.7921 | 12 | 1.5 | 433.9386 | 3 | 26.02 | 2 | 5915 | JVDM\_TOPE.raw | 3.1815E5 | 1 | 1 | 177 | 188 |  |  | PEAKS DB |
| K.VIGVTSLDIVR.A | Y | 59.77 | 1170.6971 | 11 | 1.4 | 586.3567 | 2 | 29.76 | 2 | 6771 | JVDM\_TOPE.raw | 3.3453E5 | 1 | 1 | 178 | 188 |  |  | PEAKS DB |
| R.DDLFNTNAGIVK.A | Y | 50.57 | 1305.6565 | 12 | 2.5 | 653.8372 | 2 | 25.91 | 2 | 5876 | JVDM\_TOPE.raw | 3.6104E5 | 1 | 1 | 123 | 134 |  |  | PEAKS DB |
| K.GADLIVIPAGVPR.K | Y | 49.77 | 1276.7502 | 13 | 3.3 | 639.3845 | 2 | 31.38 | 2 | 7137 | JVDM\_TOPE.raw | 5.9717E5 | 1 | 1 | 104 | 116 |  |  | PEAKS DB |
| K.HAPNAVLEIITNPVN(+.98)STVPIAVETLK.L | Y | 45.71 | 2740.5010 | 26 | 0.2 | 914.5078 | 3 | 38.74 | 2 | 9023 | JVDM\_TOPE.raw | 0 | 0 | 0 | 143 | 168 | Deamidation (NQ) | N15:Deamidation (NQ):16.03 | PEAKS PTM |
| K.HAPNAVLEIITNPVNSTVPIAVETLK.L | Y | 44.37 | 2739.5171 | 26 | 2.2 | 914.1816 | 3 | 40.38 | 2 | 9416 | JVDM\_TOPE.raw | 4.828E4 | 1 | 1 | 143 | 168 |  |  | PEAKS DB |
| K.VTGYTGPEELGAC(+57.02)LK.G | Y | 38.28 | 1593.7709 | 15 | 2.7 | 797.8948 | 2 | 24.27 | 2 | 5445 | JVDM\_TOPE.raw | 4.4207E5 | 1 | 1 | 89 | 103 | Carbamidomethylation | C13:Carbamidomethylation:1000.00 | PEAKS DB |
| K.LAGVYDPK.K | Y | 23.16 | 861.4596 | 8 | 1.2 | 431.7376 | 2 | 14.37 | 2 | 2395 | JVDM\_TOPE.raw | 4.0843E5 | 1 | 1 | 169 | 176 |  |  | PEAKS DB |
| K.VLLGPN(+.98)GVAK.V | Y | 22.48 | 967.5702 | 10 | 2.2 | 484.7935 | 2 | 19.70 | 2 | 4216 | JVDM\_TOPE.raw | 8.4824E5 | 1 | 1 | 305 | 314 | Deamidation (NQ) | N6:Deamidation (NQ):1000.00 | PEAKS PTM |
| K.AGK(+14.02)GSATLSMAYAAAR.M | Y | 16.27 | 1538.7875 | 16 | -6.3 | 513.9332 | 3 | 16.59 | 2 | 3033 | JVDM\_TOPE.raw | 4.0015E6 | 1 | 1 | 254 | 269 | Methylation(KR) | K3:Methylation(KR):16.42 | PEAKS PTM |
| total 10 peptides |  |  |  |  |  |  |  |  |  |  |  |  |  |  |  |  |  |  |  |
| --- | --- | --- | --- | --- | --- | --- | --- | --- | --- | --- | --- | --- | --- | --- | --- | --- | --- | --- | --- |

tr|Q9FE86|Q9FE86\_CHLRE

back to list

  

| Protein Coverage
| Supporting Peptides
|

Protein Coverage:

Supporting Peptides:

| Peptide | Uniq | -10lgP | Mass | Length | ppm | m/z | z | RT | Fraction | Scan | Source File | Area Sample 2 | #Feature | #Feature Sample 2 | Start | End | PTM | AScore | Found By |
| --- | --- | --- | --- | --- | --- | --- | --- | --- | --- | --- | --- | --- | --- | --- | --- | --- | --- | --- | --- |
| K.AYGVLTEDGISLR.G | Y | 68.37 | 1392.7249 | 13 | 3.0 | 697.3718 | 2 | 28.21 | 2 | 6390 | JVDM\_TOPE.raw | 4.1242E5 | 1 | 1 | 152 | 164 |  |  | PEAKS DB |
| K.AQAVFDQEFQEITLSK.Y | Y | 65.92 | 1852.9207 | 16 | 1.7 | 927.4692 | 2 | 31.90 | 2 | 7243 | JVDM\_TOPE.raw | 2.9274E5 | 1 | 1 | 55 | 70 |  |  | PEAKS DB |
| A.SHAEKPLVGSVAPDFK.A | Y | 58.89 | 1680.8834 | 16 | 2.4 | 561.3031 | 3 | 20.09 | 2 | 4241 | JVDM\_TOPE.raw | 2.5658E5 | 1 | 1 | 39 | 54 |  |  | PEAKS DB |
| K.EGGLGDLAYPLVADLKK.E | Y | 58.75 | 1757.9563 | 17 | 2.1 | 586.9940 | 3 | 34.51 | 2 | 7919 | JVDM\_TOPE.raw | 2.7066E5 | 1 | 1 | 131 | 147 |  |  | PEAKS DB |
| total 4 peptides |  |  |  |  |  |  |  |  |  |  |  |  |  |  |  |  |  |  |  |
| --- | --- | --- | --- | --- | --- | --- | --- | --- | --- | --- | --- | --- | --- | --- | --- | --- | --- | --- | --- |

tr|A8JGV6|A8JGV6\_CHLRE

back to list

  

| Protein Coverage
| Supporting Peptides
|

Protein Coverage:

Supporting Peptides:

| Peptide | Uniq | -10lgP | Mass | Length | ppm | m/z | z | RT | Fraction | Scan | Source File | Area Sample 2 | #Feature | #Feature Sample 2 | Start | End | PTM | AScore | Found By |
| --- | --- | --- | --- | --- | --- | --- | --- | --- | --- | --- | --- | --- | --- | --- | --- | --- | --- | --- | --- |
| K.LVHDQELSVEER.N | Y | 70.52 | 1452.7208 | 12 | 0.5 | 485.2478 | 3 | 12.86 | 2 | 1892 | JVDM\_TOPE.raw | 1.9723E6 | 2 | 2 | 34 | 45 |  |  | PEAKS DB |
| K.EAAEHTLLAYK.A | Y | 68.49 | 1244.6400 | 11 | 1.6 | 623.3283 | 2 | 17.45 | 2 | 3370 | JVDM\_TOPE.raw | 3.9859E5 | 1 | 1 | 146 | 156 |  |  | PEAKS DB |
| K.AAQDIALVDLPPTHPIR.L | Y | 60.56 | 1826.0050 | 17 | 1.9 | 609.6768 | 3 | 28.26 | 2 | 6412 | JVDM\_TOPE.raw | 2.1308E6 | 2 | 2 | 157 | 173 |  |  | PEAKS DB |
| K.LAEQAERYDEMVEEMKK.V | Y | 37.42 | 2097.9709 | 17 | 1.4 | 700.3319 | 3 | 21.95 | 2 | 4799 | JVDM\_TOPE.raw | 8.8522E5 | 2 | 2 | 14 | 30 |  |  | PEAKS DB |
| total 4 peptides |  |  |  |  |  |  |  |  |  |  |  |  |  |  |  |  |  |  |  |
| --- | --- | --- | --- | --- | --- | --- | --- | --- | --- | --- | --- | --- | --- | --- | --- | --- | --- | --- | --- |

P06541|ATPB\_CHLRE

back to list

  

| Protein Coverage
| Supporting Peptides
|

Protein Coverage:

Supporting Peptides:

| Peptide | Uniq | -10lgP | Mass | Length | ppm | m/z | z | RT | Fraction | Scan | Source File | Area Sample 2 | #Feature | #Feature Sample 2 | Start | End | PTM | AScore | Found By |
| --- | --- | --- | --- | --- | --- | --- | --- | --- | --- | --- | --- | --- | --- | --- | --- | --- | --- | --- | --- |
| R.FVQAGAEVSALLGR.M | Y | 73.17 | 1416.7725 | 14 | 2.0 | 709.3949 | 2 | 31.52 | 2 | 7158 | JVDM\_TOPE.raw | 1.0473E5 | 1 | 1 | 268 | 281 |  |  | PEAKS DB |
| R.VALTALTMAEYFR.D | Y | 52.19 | 1484.7697 | 13 | 3.7 | 743.3948 | 2 | 37.84 | 2 | 8787 | JVDM\_TOPE.raw | 4.4348E4 | 1 | 1 | 239 | 251 |  |  | PEAKS DB |
| K.VALVYGQMNEPPGAR.M | Y | 47.79 | 1600.8031 | 15 | 7.0 | 801.4144 | 2 | 22.97 | 2 | 5073 | JVDM\_TOPE.raw | 6.9289E4 | 1 | 1 | 222 | 236 |  |  | PEAKS DB |
| R.MPSAVGYQPTLATEMGGLQER.I | Y | 47.69 | 2235.0664 | 21 | 2.1 | 746.0310 | 3 | 29.53 | 2 | 6716 | JVDM\_TOPE.raw | 0 | 0 | 0 | 282 | 302 |  |  | PEAKS DB |
| K.GQVPNIYNALTIR.A | Y | 30.58 | 1457.7991 | 13 | 1.6 | 729.9080 | 2 | 31.77 | 2 | 7232 | JVDM\_TOPE.raw | 3.9054E4 | 1 | 1 | 28 | 40 |  |  | PEAKS DB |
| R.TAPAFVDLDTR.L | Y | 30.31 | 1204.6088 | 11 | 2.2 | 603.3130 | 2 | 24.85 | 2 | 5627 | JVDM\_TOPE.raw | 1.6021E5 | 1 | 1 | 125 | 135 |  |  | PEAKS DB |
| R.GMEVVDTGKPLSVPVGK.V | Y | 19.17 | 1711.9178 | 17 | 2.4 | 571.6479 | 3 | 23.60 | 2 | 5311 | JVDM\_TOPE.raw | 3.5529E4 | 1 | 1 | 78 | 94 |  |  | PEAKS DB |
| total 7 peptides |  |  |  |  |  |  |  |  |  |  |  |  |  |  |  |  |  |  |  |
| --- | --- | --- | --- | --- | --- | --- | --- | --- | --- | --- | --- | --- | --- | --- | --- | --- | --- | --- | --- |

tr|A8J9H8|A8J9H8\_CHLRE

back to list

  

| Protein Coverage
| Supporting Peptides
|

Protein Coverage:

Supporting Peptides:

| Peptide | Uniq | -10lgP | Mass | Length | ppm | m/z | z | RT | Fraction | Scan | Source File | Area Sample 2 | #Feature | #Feature Sample 2 | Start | End | PTM | AScore | Found By |
| --- | --- | --- | --- | --- | --- | --- | --- | --- | --- | --- | --- | --- | --- | --- | --- | --- | --- | --- | --- |
| K.MIGATNPLASEPGTIR.G | Y | 76.51 | 1626.8400 | 16 | 2.4 | 814.4292 | 2 | 25.34 | 2 | 5746 | JVDM\_TOPE.raw | 3.0907E5 | 1 | 1 | 89 | 104 |  |  | PEAKS DB |
| R.KMIGATNPLASEPGTIR.G | Y | 62.15 | 1754.9348 | 17 | 2.6 | 585.9871 | 3 | 22.37 | 2 | 4932 | JVDM\_TOPE.raw | 3.1472E5 | 1 | 1 | 88 | 104 |  |  | PEAKS DB |
| R.GDFAIEVGR.N | Y | 49.99 | 962.4821 | 9 | 0.2 | 482.2484 | 2 | 22.17 | 2 | 4857 | JVDM\_TOPE.raw | 1.1754E6 | 1 | 1 | 105 | 113 |  |  | PEAKS DB |
| total 3 peptides |  |  |  |  |  |  |  |  |  |  |  |  |  |  |  |  |  |  |  |
| --- | --- | --- | --- | --- | --- | --- | --- | --- | --- | --- | --- | --- | --- | --- | --- | --- | --- | --- | --- |

tr|A0A2K3DJF1|A0A2K3DJF1\_CHLRE

back to list

  

| Protein Coverage
| Supporting Peptides
|

Protein Coverage:

Supporting Peptides:

| Peptide | Uniq | -10lgP | Mass | Length | ppm | m/z | z | RT | Fraction | Scan | Source File | Area Sample 2 | #Feature | #Feature Sample 2 | Start | End | PTM | AScore | Found By |
| --- | --- | --- | --- | --- | --- | --- | --- | --- | --- | --- | --- | --- | --- | --- | --- | --- | --- | --- | --- |
| K.MIGATNPLASEPGTIR.G | Y | 76.51 | 1626.8400 | 16 | 2.4 | 814.4292 | 2 | 25.34 | 2 | 5746 | JVDM\_TOPE.raw | 3.0907E5 | 1 | 1 | 101 | 116 |  |  | PEAKS DB |
| R.KMIGATNPLASEPGTIR.G | Y | 62.15 | 1754.9348 | 17 | 2.6 | 585.9871 | 3 | 22.37 | 2 | 4932 | JVDM\_TOPE.raw | 3.1472E5 | 1 | 1 | 100 | 116 |  |  | PEAKS DB |
| R.GDFAIEVGR.N | Y | 49.99 | 962.4821 | 9 | 0.2 | 482.2484 | 2 | 22.17 | 2 | 4857 | JVDM\_TOPE.raw | 1.1754E6 | 1 | 1 | 117 | 125 |  |  | PEAKS DB |
| total 3 peptides |  |  |  |  |  |  |  |  |  |  |  |  |  |  |  |  |  |  |  |
| --- | --- | --- | --- | --- | --- | --- | --- | --- | --- | --- | --- | --- | --- | --- | --- | --- | --- | --- | --- |

tr|A8JC04|A8JC04\_CHLRE

back to list

  

| Protein Coverage
| Supporting Peptides
|

Protein Coverage:

Supporting Peptides:

| Peptide | Uniq | -10lgP | Mass | Length | ppm | m/z | z | RT | Fraction | Scan | Source File | Area Sample 2 | #Feature | #Feature Sample 2 | Start | End | PTM | AScore | Found By |
| --- | --- | --- | --- | --- | --- | --- | --- | --- | --- | --- | --- | --- | --- | --- | --- | --- | --- | --- | --- |
| K.ELDYLDGAVSNPK.R | Y | 68.47 | 1419.6881 | 13 | 1.0 | 710.8521 | 2 | 23.11 | 2 | 5159 | JVDM\_TOPE.raw | 8.4813E4 | 1 | 1 | 240 | 252 |  |  | PEAKS DB |
| K.FAN(+.98)GTVSIANTLAGLTPK.G | Y | 50.37 | 1774.9465 | 18 | 2.5 | 888.4828 | 2 | 36.72 | 2 | 8488 | JVDM\_TOPE.raw | 4.1581E4 | 1 | 1 | 393 | 410 | Deamidation (NQ) | N3:Deamidation (NQ):61.82 | PEAKS PTM |
| K.VLPGVAALDEK | Y | 46.73 | 1110.6284 | 11 | 0.7 | 556.3219 | 2 | 24.52 | 2 | 5547 | JVDM\_TOPE.raw | 2.0309E5 | 1 | 1 | 452 | 462 |  |  | PEAKS DB |
| K.RPFVAIVGGSK.V | Y | 41.27 | 1129.6608 | 11 | 1.7 | 565.8386 | 2 | 19.92 | 2 | 4206 | JVDM\_TOPE.raw | 5.5773E4 | 1 | 1 | 253 | 263 |  |  | PEAKS DB |
| R.LTPVVAR.L | Y | 28.09 | 754.4701 | 7 | 1.6 | 378.2429 | 2 | 13.58 | 2 | 2184 | JVDM\_TOPE.raw | 1.3424E5 | 1 | 1 | 138 | 144 |  |  | PEAKS DB |
| K.FLKPSVAGFLLQK.E | Y | 22.52 | 1446.8599 | 13 | 2.2 | 483.2950 | 3 | 31.85 | 2 | 7252 | JVDM\_TOPE.raw | 4.5437E4 | 1 | 1 | 227 | 239 |  |  | PEAKS DB |
| K.ATLAITDDTR.I | Y | 17.01 | 1075.5509 | 10 | 0.7 | 538.7831 | 2 | 16.92 | 2 | 3208 | JVDM\_TOPE.raw | 5.2825E4 | 1 | 1 | 92 | 101 |  |  | PEAKS DB |
| total 7 peptides |  |  |  |  |  |  |  |  |  |  |  |  |  |  |  |  |  |  |  |
| --- | --- | --- | --- | --- | --- | --- | --- | --- | --- | --- | --- | --- | --- | --- | --- | --- | --- | --- | --- |

tr|B7U1J0|B7U1J0\_CHLRE

back to list

  

| Protein Coverage
| Supporting Peptides
|

Protein Coverage:

Supporting Peptides:

| Peptide | Uniq | -10lgP | Mass | Length | ppm | m/z | z | RT | Fraction | Scan | Source File | Area Sample 2 | #Feature | #Feature Sample 2 | Start | End | PTM | AScore | Found By |
| --- | --- | --- | --- | --- | --- | --- | --- | --- | --- | --- | --- | --- | --- | --- | --- | --- | --- | --- | --- |
| K.ASSVAQVLNTLK.E | Y | 73.85 | 1229.6979 | 12 | 2.1 | 615.8575 | 2 | 29.22 | 2 | 6641 | JVDM\_TOPE.raw | 3.0606E5 | 1 | 1 | 203 | 214 |  |  | PEAKS DB |
| R.STLTFTPEAEGLVK.Q | Y | 53.33 | 1491.7820 | 14 | 2.6 | 746.9002 | 2 | 28.36 | 2 | 6435 | JVDM\_TOPE.raw | 1.1766E5 | 1 | 1 | 478 | 491 |  |  | PEAKS DB |
| K.TAIAVDTILNQK.G | Y | 49.20 | 1285.7241 | 12 | 2.2 | 643.8707 | 2 | 26.91 | 2 | 6101 | JVDM\_TOPE.raw | 3.2313E5 | 1 | 1 | 177 | 188 |  |  | PEAKS DB |
| R.ELIIGDR.Q | Y | 26.68 | 814.4548 | 7 | 1.9 | 408.2355 | 2 | 18.74 | 2 | 3800 | JVDM\_TOPE.raw | 3.0707E5 | 1 | 1 | 166 | 172 |  |  | PEAKS DB |
| L.ATGLVAVDAMIPVGR.G | Y | 23.16 | 1468.8071 | 15 | -0.3 | 735.4106 | 2 | 33.56 | 2 | 7680 | JVDM\_TOPE.raw | 0 | 0 | 0 | 148 | 162 |  |  | PEAKS DB |
| total 5 peptides |  |  |  |  |  |  |  |  |  |  |  |  |  |  |  |  |  |  |  |
| --- | --- | --- | --- | --- | --- | --- | --- | --- | --- | --- | --- | --- | --- | --- | --- | --- | --- | --- | --- |

P26526|ATPA\_CHLRE

back to list

  

| Protein Coverage
| Supporting Peptides
|

Protein Coverage:

Supporting Peptides:

| Peptide | Uniq | -10lgP | Mass | Length | ppm | m/z | z | RT | Fraction | Scan | Source File | Area Sample 2 | #Feature | #Feature Sample 2 | Start | End | PTM | AScore | Found By |
| --- | --- | --- | --- | --- | --- | --- | --- | --- | --- | --- | --- | --- | --- | --- | --- | --- | --- | --- | --- |
| K.ASSVAQVLNTLK.E | Y | 73.85 | 1229.6979 | 12 | 2.1 | 615.8575 | 2 | 29.22 | 2 | 6641 | JVDM\_TOPE.raw | 3.0606E5 | 1 | 1 | 203 | 214 |  |  | PEAKS DB |
| R.STLTFTPEAEGLVK.Q | Y | 53.33 | 1491.7820 | 14 | 2.6 | 746.9002 | 2 | 28.36 | 2 | 6435 | JVDM\_TOPE.raw | 1.1766E5 | 1 | 1 | 478 | 491 |  |  | PEAKS DB |
| K.TAIAVDTILNQK.G | Y | 49.20 | 1285.7241 | 12 | 2.2 | 643.8707 | 2 | 26.91 | 2 | 6101 | JVDM\_TOPE.raw | 3.2313E5 | 1 | 1 | 177 | 188 |  |  | PEAKS DB |
| R.ELIIGDR.Q | Y | 26.68 | 814.4548 | 7 | 1.9 | 408.2355 | 2 | 18.74 | 2 | 3800 | JVDM\_TOPE.raw | 3.0707E5 | 1 | 1 | 166 | 172 |  |  | PEAKS DB |
| L.ATGLVAVDAMIPVGR.G | Y | 23.16 | 1468.8071 | 15 | -0.3 | 735.4106 | 2 | 33.56 | 2 | 7680 | JVDM\_TOPE.raw | 0 | 0 | 0 | 148 | 162 |  |  | PEAKS DB |
| total 5 peptides |  |  |  |  |  |  |  |  |  |  |  |  |  |  |  |  |  |  |  |
| --- | --- | --- | --- | --- | --- | --- | --- | --- | --- | --- | --- | --- | --- | --- | --- | --- | --- | --- | --- |

tr|A0A2K3DJ22|A0A2K3DJ22\_CHLRE

back to list

  

| Protein Coverage
| Supporting Peptides
|

Protein Coverage:

Supporting Peptides:

| Peptide | Uniq | -10lgP | Mass | Length | ppm | m/z | z | RT | Fraction | Scan | Source File | Area Sample 2 | #Feature | #Feature Sample 2 | Start | End | PTM | AScore | Found By |
| --- | --- | --- | --- | --- | --- | --- | --- | --- | --- | --- | --- | --- | --- | --- | --- | --- | --- | --- | --- |
| K.LPVDFPAAVASGC(+57.02)AK.T | Y | 88.47 | 1501.7599 | 15 | 2.7 | 751.8892 | 2 | 28.27 | 2 | 6408 | JVDM\_TOPE.raw | 8.9856E4 | 1 | 1 | 1703 | 1717 | Carbamidomethylation | C13:Carbamidomethylation:1000.00 | PEAKS DB |
| K.SESVIAGGADPC(+57.02)TGAVSESANLLSALAR.D | Y | 53.48 | 2702.3181 | 28 | 3.2 | 901.7828 | 3 | 37.51 | 2 | 8698 | JVDM\_TOPE.raw | 4.6999E4 | 1 | 1 | 1746 | 1773 | Carbamidomethylation | C12:Carbamidomethylation:1000.00 | PEAKS DB |
| total 2 peptides |  |  |  |  |  |  |  |  |  |  |  |  |  |  |  |  |  |  |  |
| --- | --- | --- | --- | --- | --- | --- | --- | --- | --- | --- | --- | --- | --- | --- | --- | --- | --- | --- | --- |

tr|A8HXL8|A8HXL8\_CHLRE

back to list

  

| Protein Coverage
| Supporting Peptides
|

Protein Coverage:

Supporting Peptides:

| Peptide | Uniq | -10lgP | Mass | Length | ppm | m/z | z | RT | Fraction | Scan | Source File | Area Sample 2 | #Feature | #Feature Sample 2 | Start | End | PTM | AScore | Found By |
| --- | --- | --- | --- | --- | --- | --- | --- | --- | --- | --- | --- | --- | --- | --- | --- | --- | --- | --- | --- |
| R.SLQEALASELAAR.M | Y | 70.42 | 1357.7201 | 13 | 2.1 | 679.8688 | 2 | 29.71 | 2 | 6756 | JVDM\_TOPE.raw | 2.0197E5 | 1 | 1 | 300 | 312 |  |  | PEAKS DB |
| K.SFSLGAAPSTK.E | Y | 53.49 | 1064.5502 | 11 | 1.1 | 533.2830 | 2 | 19.13 | 2 | 3962 | JVDM\_TOPE.raw | 2.2551E5 | 1 | 1 | 172 | 182 |  |  | PEAKS DB |
| K.SVLLVVLTGDR.G | Y | 51.19 | 1170.6971 | 11 | 1.9 | 586.3569 | 2 | 34.63 | 2 | 7958 | JVDM\_TOPE.raw | 2.6572E5 | 1 | 1 | 110 | 120 |  |  | PEAKS DB |
| K.VNLVC(+57.02)VGR.K | Y | 19.00 | 915.4960 | 8 | -0.6 | 458.7550 | 2 | 18.77 | 2 | 3813 | JVDM\_TOPE.raw | 1.6723E5 | 1 | 1 | 148 | 155 | Carbamidomethylation | C5:Carbamidomethylation:1000.00 | PEAKS DB |
| total 4 peptides |  |  |  |  |  |  |  |  |  |  |  |  |  |  |  |  |  |  |  |
| --- | --- | --- | --- | --- | --- | --- | --- | --- | --- | --- | --- | --- | --- | --- | --- | --- | --- | --- | --- |

tr|A8IGH1|A8IGH1\_CHLRE

back to list

  

| Protein Coverage
| Supporting Peptides
|

Protein Coverage:

Supporting Peptides:

| Peptide | Uniq | -10lgP | Mass | Length | ppm | m/z | z | RT | Fraction | Scan | Source File | Area Sample 2 | #Feature | #Feature Sample 2 | Start | End | PTM | AScore | Found By |
| --- | --- | --- | --- | --- | --- | --- | --- | --- | --- | --- | --- | --- | --- | --- | --- | --- | --- | --- | --- |
| K.SPNAVNPVVEGK.T | Y | 63.87 | 1209.6353 | 12 | 2.1 | 605.8262 | 2 | 13.19 | 2 | 2057 | JVDM\_TOPE.raw | 3.9422E5 | 1 | 1 | 176 | 187 |  |  | PEAKS DB |
| K.SPPYALDALEPHMSK.Q | Y | 58.86 | 1654.8025 | 15 | 2.5 | 552.6095 | 3 | 26.96 | 2 | 6116 | JVDM\_TOPE.raw | 2.211E5 | 1 | 1 | 39 | 53 |  |  | PEAKS DB |
| K.LINWDAVAQR.Y | Y | 51.20 | 1184.6301 | 10 | 2.3 | 593.3237 | 2 | 26.21 | 2 | 5959 | JVDM\_TOPE.raw | 3.6451E5 | 1 | 1 | 219 | 228 |  |  | PEAKS DB |
| total 3 peptides |  |  |  |  |  |  |  |  |  |  |  |  |  |  |  |  |  |  |  |
| --- | --- | --- | --- | --- | --- | --- | --- | --- | --- | --- | --- | --- | --- | --- | --- | --- | --- | --- | --- |

tr|A8J0E4|A8J0E4\_CHLRE

back to list

  

| Protein Coverage
| Supporting Peptides
|

Protein Coverage:

Supporting Peptides:

| Peptide | Uniq | -10lgP | Mass | Length | ppm | m/z | z | RT | Fraction | Scan | Source File | Area Sample 2 | #Feature | #Feature Sample 2 | Start | End | PTM | AScore | Found By |
| --- | --- | --- | --- | --- | --- | --- | --- | --- | --- | --- | --- | --- | --- | --- | --- | --- | --- | --- | --- |
| K.GSGIANTC(+57.02)PVLESGTTNLK.E | Y | 76.35 | 1917.9465 | 19 | 4.5 | 959.9849 | 2 | 23.38 | 2 | 5237 | JVDM\_TOPE.raw | 3.3615E4 | 1 | 1 | 67 | 85 | Carbamidomethylation | C8:Carbamidomethylation:1000.00 | PEAKS DB |
| K.VGSDGSAELKEDDGIDYAATTVQLPGGER.V | Y | 63.94 | 2949.3838 | 29 | 0.9 | 984.1361 | 3 | 26.62 | 2 | 6042 | JVDM\_TOPE.raw | 0 | 0 | 0 | 141 | 169 |  |  | PEAKS DB |
| A.LTFDEIQGLTYLQVK.G | Y | 31.35 | 1766.9454 | 15 | -3.8 | 884.4766 | 2 | 36.84 | 2 | 8507 | JVDM\_TOPE.raw | 0 | 0 | 0 | 52 | 66 |  |  | PEAKS DB |
| R.LTYTLDAMSGSFK.V | Y | 30.79 | 1432.6908 | 13 | 3.4 | 717.3551 | 2 | 28.93 | 2 | 6572 | JVDM\_TOPE.raw | 1.5469E4 | 1 | 1 | 128 | 140 |  |  | PEAKS DB |
| total 4 peptides |  |  |  |  |  |  |  |  |  |  |  |  |  |  |  |  |  |  |  |
| --- | --- | --- | --- | --- | --- | --- | --- | --- | --- | --- | --- | --- | --- | --- | --- | --- | --- | --- | --- |

tr|A0A2K3CWZ6|A0A2K3CWZ6\_CHLRE

back to list

  

| Protein Coverage
| Supporting Peptides
|

Protein Coverage:

Supporting Peptides:

| Peptide | Uniq | -10lgP | Mass | Length | ppm | m/z | z | RT | Fraction | Scan | Source File | Area Sample 2 | #Feature | #Feature Sample 2 | Start | End | PTM | AScore | Found By |
| --- | --- | --- | --- | --- | --- | --- | --- | --- | --- | --- | --- | --- | --- | --- | --- | --- | --- | --- | --- |
| R.IAEEAGAVAVMALER.V | Y | 66.22 | 1528.7919 | 15 | 1.7 | 765.4045 | 2 | 29.76 | 2 | 6770 | JVDM\_TOPE.raw | 8.0948E4 | 1 | 1 | 36 | 50 |  |  | PEAKS DB |
| K.GGVIMDVTTAEEAR.I | Y | 55.65 | 1447.6976 | 14 | 2.4 | 724.8578 | 2 | 23.51 | 2 | 5296 | JVDM\_TOPE.raw | 2.0724E5 | 1 | 1 | 22 | 35 |  |  | PEAKS DB |
| K.AAVTIPVMAK.A | Y | 36.17 | 999.5787 | 10 | 1.9 | 500.7976 | 2 | 24.23 | 2 | 5467 | JVDM\_TOPE.raw | 1.0159E5 | 1 | 1 | 75 | 84 |  |  | PEAKS DB |
| R.IAEGAAMIR.T | Y | 27.92 | 930.4957 | 9 | -0.2 | 466.2550 | 2 | 16.17 | 2 | 2962 | JVDM\_TOPE.raw | 1.2289E5 | 1 | 1 | 142 | 150 |  |  | PEAKS DB |
| K.SSGTFAVK.V | Y | 19.90 | 795.4127 | 8 | 1.3 | 398.7141 | 2 | 12.91 | 2 | 1946 | JVDM\_TOPE.raw | 9.2055E4 | 1 | 1 | 6 | 13 |  |  | PEAKS DB |
| R.TKGEAGTGNVVEAVR(+14.02).H | Y | 16.24 | 1500.7896 | 15 | 5.2 | 751.4059 | 2 | 30.48 | 2 | 6951 | JVDM\_TOPE.raw | 1.6497E5 | 1 | 1 | 151 | 165 | Methylation(KR) | R15:Methylation(KR):57.37 | PEAKS PTM |
| total 6 peptides |  |  |  |  |  |  |  |  |  |  |  |  |  |  |  |  |  |  |  |
| --- | --- | --- | --- | --- | --- | --- | --- | --- | --- | --- | --- | --- | --- | --- | --- | --- | --- | --- | --- |

tr|A8IKW6|A8IKW6\_CHLRE

back to list

  

| Protein Coverage
| Supporting Peptides
|

Protein Coverage:

Supporting Peptides:

| Peptide | Uniq | -10lgP | Mass | Length | ppm | m/z | z | RT | Fraction | Scan | Source File | Area Sample 2 | #Feature | #Feature Sample 2 | Start | End | PTM | AScore | Found By |
| --- | --- | --- | --- | --- | --- | --- | --- | --- | --- | --- | --- | --- | --- | --- | --- | --- | --- | --- | --- |
| K.VIDAGANALVAGSAVFK.A | Y | 86.20 | 1601.8777 | 17 | 1.9 | 801.9476 | 2 | 30.57 | 2 | 6984 | JVDM\_TOPE.raw | 2.9659E4 | 1 | 1 | 227 | 243 |  |  | PEAKS DB |
| K.SDIIVSPSILSADFSR.L | Y | 32.02 | 1705.8887 | 16 | 2.8 | 853.9540 | 2 | 35.53 | 2 | 8153 | JVDM\_TOPE.raw | 0 | 0 | 0 | 40 | 55 |  |  | PEAKS DB |
| total 2 peptides |  |  |  |  |  |  |  |  |  |  |  |  |  |  |  |  |  |  |  |
| --- | --- | --- | --- | --- | --- | --- | --- | --- | --- | --- | --- | --- | --- | --- | --- | --- | --- | --- | --- |

tr|A8ILJ9|A8ILJ9\_CHLRE

back to list

  

| Protein Coverage
| Supporting Peptides
|

Protein Coverage:

Supporting Peptides:

| Peptide | Uniq | -10lgP | Mass | Length | ppm | m/z | z | RT | Fraction | Scan | Source File | Area Sample 2 | #Feature | #Feature Sample 2 | Start | End | PTM | AScore | Found By |
| --- | --- | --- | --- | --- | --- | --- | --- | --- | --- | --- | --- | --- | --- | --- | --- | --- | --- | --- | --- |
| R.AAGLSSITVTSDINAVK.G | Y | 70.42 | 1645.8887 | 17 | 3.0 | 823.9540 | 2 | 27.14 | 2 | 6155 | JVDM\_TOPE.raw | 1.0669E5 | 1 | 1 | 254 | 270 |  |  | PEAKS DB |
| R.VLSGYNDVIMAR.L | Y | 41.74 | 1336.6809 | 12 | 1.7 | 669.3489 | 2 | 24.49 | 2 | 5542 | JVDM\_TOPE.raw | 7.1279E4 | 1 | 1 | 149 | 160 |  |  | PEAKS DB |
| R.GLETTDGVMESPQ(+.98)SVVFQEAENR.M | Y | 31.65 | 2523.1436 | 23 | 7.2 | 842.0612 | 3 | 29.85 | 2 | 6801 | JVDM\_TOPE.raw | 4.5335E4 | 1 | 1 | 324 | 346 | Deamidation (NQ) | Q13:Deamidation (NQ):61.30 | PEAKS PTM |
| R.GLETTDGVMESPQSVVFQEAENR.M | Y | 26.61 | 2522.1594 | 23 | 1.6 | 841.7284 | 3 | 29.97 | 2 | 6821 | JVDM\_TOPE.raw | 0 | 0 | 0 | 324 | 346 |  |  | PEAKS DB |
| total 4 peptides |  |  |  |  |  |  |  |  |  |  |  |  |  |  |  |  |  |  |  |
| --- | --- | --- | --- | --- | --- | --- | --- | --- | --- | --- | --- | --- | --- | --- | --- | --- | --- | --- | --- |

tr|A0A2K3DZ46|A0A2K3DZ46\_CHLRE

back to list

  

| Protein Coverage
| Supporting Peptides
|

Protein Coverage:

Supporting Peptides:

| Peptide | Uniq | -10lgP | Mass | Length | ppm | m/z | z | RT | Fraction | Scan | Source File | Area Sample 2 | #Feature | #Feature Sample 2 | Start | End | PTM | AScore | Found By |
| --- | --- | --- | --- | --- | --- | --- | --- | --- | --- | --- | --- | --- | --- | --- | --- | --- | --- | --- | --- |
| K.LVTDFADGAYTAHAAGR.F | Y | 61.52 | 1734.8325 | 17 | 1.4 | 579.2856 | 3 | 21.86 | 2 | 4774 | JVDM\_TOPE.raw | 2.5121E5 | 1 | 1 | 253 | 269 |  |  | PEAKS DB |
| K.ELEDAAIVQAFTR.C | Y | 57.92 | 1461.7463 | 13 | 0.6 | 731.8809 | 2 | 32.81 | 2 | 7441 | JVDM\_TOPE.raw | 4.1742E4 | 1 | 1 | 182 | 194 |  |  | PEAKS DB |
| R.DGPQPGWGDLLAK.L | Y | 32.46 | 1352.6724 | 13 | 0.1 | 677.3435 | 2 | 29.89 | 2 | 6810 | JVDM\_TOPE.raw | 7.162E4 | 1 | 1 | 240 | 252 |  |  | PEAKS DB |
| total 3 peptides |  |  |  |  |  |  |  |  |  |  |  |  |  |  |  |  |  |  |  |
| --- | --- | --- | --- | --- | --- | --- | --- | --- | --- | --- | --- | --- | --- | --- | --- | --- | --- | --- | --- |

tr|A8JEV1|A8JEV1\_CHLRE

back to list

  

| Protein Coverage
| Supporting Peptides
|

Protein Coverage:

Supporting Peptides:

| Peptide | Uniq | -10lgP | Mass | Length | ppm | m/z | z | RT | Fraction | Scan | Source File | Area Sample 2 | #Feature | #Feature Sample 2 | Start | End | PTM | AScore | Found By |
| --- | --- | --- | --- | --- | --- | --- | --- | --- | --- | --- | --- | --- | --- | --- | --- | --- | --- | --- | --- |
| R.IDADLDVFIQK.S | Y | 76.70 | 1275.6710 | 11 | 2.7 | 638.8445 | 2 | 30.90 | 2 | 7052 | JVDM\_TOPE.raw | 3.2681E5 | 1 | 1 | 108 | 118 |  |  | PEAKS DB |
| R.FDLNTLASTK.E | Y | 48.55 | 1108.5764 | 10 | 2.1 | 555.2966 | 2 | 25.53 | 2 | 5786 | JVDM\_TOPE.raw | 4.9507E5 | 1 | 1 | 137 | 146 |  |  | PEAKS DB |
| K.AKLDSVLAAVL | Y | 30.17 | 1098.6648 | 11 | 1.2 | 550.3403 | 2 | 37.15 | 2 | 8608 | JVDM\_TOPE.raw | 3.5223E5 | 1 | 1 | 189 | 199 |  |  | PEAKS DB |
| total 3 peptides |  |  |  |  |  |  |  |  |  |  |  |  |  |  |  |  |  |  |  |
| --- | --- | --- | --- | --- | --- | --- | --- | --- | --- | --- | --- | --- | --- | --- | --- | --- | --- | --- | --- |

tr|A8IKZ2|A8IKZ2\_CHLRE

back to list

  

| Protein Coverage
| Supporting Peptides
|

Protein Coverage:

Supporting Peptides:

| Peptide | Uniq | -10lgP | Mass | Length | ppm | m/z | z | RT | Fraction | Scan | Source File | Area Sample 2 | #Feature | #Feature Sample 2 | Start | End | PTM | AScore | Found By |
| --- | --- | --- | --- | --- | --- | --- | --- | --- | --- | --- | --- | --- | --- | --- | --- | --- | --- | --- | --- |
| K.AGGEC(+57.02)LTFDQLALQRPTGK.D | Y | 67.65 | 2061.0312 | 19 | 1.8 | 688.0189 | 3 | 26.93 | 2 | 6111 | JVDM\_TOPE.raw | 1.6893E5 | 1 | 1 | 115 | 133 | Carbamidomethylation | C5:Carbamidomethylation:1000.00 | PEAKS DB |
| K.VAVIVATVTDDVR.L | Y | 66.22 | 1356.7612 | 13 | 2.1 | 679.3893 | 2 | 27.26 | 2 | 6185 | JVDM\_TOPE.raw | 4.0074E5 | 1 | 1 | 78 | 90 |  |  | PEAKS DB |
| total 2 peptides |  |  |  |  |  |  |  |  |  |  |  |  |  |  |  |  |  |  |  |
| --- | --- | --- | --- | --- | --- | --- | --- | --- | --- | --- | --- | --- | --- | --- | --- | --- | --- | --- | --- |

tr|A0A2K3DP12|A0A2K3DP12\_CHLRE

back to list

  

| Protein Coverage
| Supporting Peptides
|

Protein Coverage:

Supporting Peptides:

| Peptide | Uniq | -10lgP | Mass | Length | ppm | m/z | z | RT | Fraction | Scan | Source File | Area Sample 2 | #Feature | #Feature Sample 2 | Start | End | PTM | AScore | Found By |
| --- | --- | --- | --- | --- | --- | --- | --- | --- | --- | --- | --- | --- | --- | --- | --- | --- | --- | --- | --- |
| R.SVGAAGGQEDIVNLVPTAAAATSTGK.S | Y | 99.14 | 2384.2183 | 26 | 2.2 | 795.7484 | 3 | 29.98 | 2 | 6827 | JVDM\_TOPE.raw | 2.6567E5 | 1 | 1 | 200 | 225 |  |  | PEAKS DB |
| K.VYGPIEIPC(+57.02)TGSQPTK.G | Y | 29.56 | 1745.8658 | 16 | 2.2 | 873.9421 | 2 | 25.09 | 2 | 5691 | JVDM\_TOPE.raw | 9.6984E4 | 1 | 1 | 305 | 320 | Carbamidomethylation | C9:Carbamidomethylation:1000.00 | PEAKS DB |
| R.VINKVYGPIEIPC(+57.02)TGSQPTK.G | Y | 24.99 | 2200.1562 | 20 | 3.2 | 734.3950 | 3 | 25.04 | 2 | 5688 | JVDM\_TOPE.raw | 3.0425E4 | 1 | 1 | 301 | 320 | Carbamidomethylation | C13:Carbamidomethylation:1000.00 | PEAKS DB |
| total 3 peptides |  |  |  |  |  |  |  |  |  |  |  |  |  |  |  |  |  |  |  |
| --- | --- | --- | --- | --- | --- | --- | --- | --- | --- | --- | --- | --- | --- | --- | --- | --- | --- | --- | --- |

tr|A8J7F8|A8J7F8\_CHLRE

back to list

  

| Protein Coverage
| Supporting Peptides
|

Protein Coverage:

Supporting Peptides:

| Peptide | Uniq | -10lgP | Mass | Length | ppm | m/z | z | RT | Fraction | Scan | Source File | Area Sample 2 | #Feature | #Feature Sample 2 | Start | End | PTM | AScore | Found By |
| --- | --- | --- | --- | --- | --- | --- | --- | --- | --- | --- | --- | --- | --- | --- | --- | --- | --- | --- | --- |
| R.GVALIAPDTSPR.G | Y | 59.73 | 1195.6560 | 12 | 0.8 | 598.8358 | 2 | 21.70 | 2 | 4725 | JVDM\_TOPE.raw | 2.0243E5 | 1 | 1 | 78 | 89 |  |  | PEAKS DB |
| R.TISAFSPISNPINAPWGVK.A | Y | 53.14 | 1998.0574 | 19 | 3.2 | 1000.0392 | 2 | 36.17 | 2 | 8336 | JVDM\_TOPE.raw | 1.373E4 | 1 | 1 | 173 | 191 |  |  | PEAKS DB |
| R.MYEYVTEELPALLR.A | Y | 20.73 | 1725.8647 | 14 | 6.9 | 863.9456 | 2 | 37.74 | 2 | 8753 | JVDM\_TOPE.raw | 0 | 0 | 0 | 122 | 135 |  |  | PEAKS DB |
| total 3 peptides |  |  |  |  |  |  |  |  |  |  |  |  |  |  |  |  |  |  |  |
| --- | --- | --- | --- | --- | --- | --- | --- | --- | --- | --- | --- | --- | --- | --- | --- | --- | --- | --- | --- |

tr|A0A2K3DW85|A0A2K3DW85\_CHLRE

back to list

  

| Protein Coverage
| Supporting Peptides
|

Protein Coverage:

Supporting Peptides:

| Peptide | Uniq | -10lgP | Mass | Length | ppm | m/z | z | RT | Fraction | Scan | Source File | Area Sample 2 | #Feature | #Feature Sample 2 | Start | End | PTM | AScore | Found By |
| --- | --- | --- | --- | --- | --- | --- | --- | --- | --- | --- | --- | --- | --- | --- | --- | --- | --- | --- | --- |
| R.GVALIAPDTSPR.G | Y | 59.73 | 1195.6560 | 12 | 0.8 | 598.8358 | 2 | 21.70 | 2 | 4725 | JVDM\_TOPE.raw | 2.0243E5 | 1 | 1 | 79 | 90 |  |  | PEAKS DB |
| R.TISAFSPISNPINAPWGVK.A | Y | 53.14 | 1998.0574 | 19 | 3.2 | 1000.0392 | 2 | 36.17 | 2 | 8336 | JVDM\_TOPE.raw | 1.373E4 | 1 | 1 | 174 | 192 |  |  | PEAKS DB |
| R.MYEYVTEELPALLR.A | Y | 20.73 | 1725.8647 | 14 | 6.9 | 863.9456 | 2 | 37.74 | 2 | 8753 | JVDM\_TOPE.raw | 0 | 0 | 0 | 123 | 136 |  |  | PEAKS DB |
| total 3 peptides |  |  |  |  |  |  |  |  |  |  |  |  |  |  |  |  |  |  |  |
| --- | --- | --- | --- | --- | --- | --- | --- | --- | --- | --- | --- | --- | --- | --- | --- | --- | --- | --- | --- |

tr|A0A2K3DW83|A0A2K3DW83\_CHLRE

back to list

  

| Protein Coverage
| Supporting Peptides
|

Protein Coverage:

Supporting Peptides:

| Peptide | Uniq | -10lgP | Mass | Length | ppm | m/z | z | RT | Fraction | Scan | Source File | Area Sample 2 | #Feature | #Feature Sample 2 | Start | End | PTM | AScore | Found By |
| --- | --- | --- | --- | --- | --- | --- | --- | --- | --- | --- | --- | --- | --- | --- | --- | --- | --- | --- | --- |
| R.GVALIAPDTSPR.G | Y | 59.73 | 1195.6560 | 12 | 0.8 | 598.8358 | 2 | 21.70 | 2 | 4725 | JVDM\_TOPE.raw | 2.0243E5 | 1 | 1 | 84 | 95 |  |  | PEAKS DB |
| R.TISAFSPISNPINAPWGVK.A | Y | 53.14 | 1998.0574 | 19 | 3.2 | 1000.0392 | 2 | 36.17 | 2 | 8336 | JVDM\_TOPE.raw | 1.373E4 | 1 | 1 | 179 | 197 |  |  | PEAKS DB |
| R.MYEYVTEELPALLR.A | Y | 20.73 | 1725.8647 | 14 | 6.9 | 863.9456 | 2 | 37.74 | 2 | 8753 | JVDM\_TOPE.raw | 0 | 0 | 0 | 128 | 141 |  |  | PEAKS DB |
| total 3 peptides |  |  |  |  |  |  |  |  |  |  |  |  |  |  |  |  |  |  |  |
| --- | --- | --- | --- | --- | --- | --- | --- | --- | --- | --- | --- | --- | --- | --- | --- | --- | --- | --- | --- |

tr|A8IMR9|A8IMR9\_CHLRE

back to list

  

| Protein Coverage
| Supporting Peptides
|

Protein Coverage:

Supporting Peptides:

| Peptide | Uniq | -10lgP | Mass | Length | ppm | m/z | z | RT | Fraction | Scan | Source File | Area Sample 2 | #Feature | #Feature Sample 2 | Start | End | PTM | AScore | Found By |
| --- | --- | --- | --- | --- | --- | --- | --- | --- | --- | --- | --- | --- | --- | --- | --- | --- | --- | --- | --- |
| R.TLVHNPENMAEAQLLPR.K | Y | 85.29 | 1931.9886 | 17 | 2.0 | 645.0048 | 3 | 25.76 | 2 | 5846 | JVDM\_TOPE.raw | 8.6679E4 | 1 | 1 | 252 | 268 |  |  | PEAKS DB |
| total 1 peptides |  |  |  |  |  |  |  |  |  |  |  |  |  |  |  |  |  |  |  |
| --- | --- | --- | --- | --- | --- | --- | --- | --- | --- | --- | --- | --- | --- | --- | --- | --- | --- | --- | --- |

tr|A0A2K3D9K7|A0A2K3D9K7\_CHLRE

back to list

  

| Protein Coverage
| Supporting Peptides
|

Protein Coverage:

Supporting Peptides:

| Peptide | Uniq | -10lgP | Mass | Length | ppm | m/z | z | RT | Fraction | Scan | Source File | Area Sample 2 | #Feature | #Feature Sample 2 | Start | End | PTM | AScore | Found By |
| --- | --- | --- | --- | --- | --- | --- | --- | --- | --- | --- | --- | --- | --- | --- | --- | --- | --- | --- | --- |
| R.LAAEAEAAAAAEAEAAAR.A | Y | 82.51 | 1655.8114 | 18 | 1.6 | 552.9453 | 3 | 24.27 | 2 | 5458 | JVDM\_TOPE.raw | 3.0578E5 | 2 | 2 | 627 | 644 |  |  | PEAKS DB |
| total 1 peptides |  |  |  |  |  |  |  |  |  |  |  |  |  |  |  |  |  |  |  |
| --- | --- | --- | --- | --- | --- | --- | --- | --- | --- | --- | --- | --- | --- | --- | --- | --- | --- | --- | --- |

Q08365|RR3\_CHLRE

back to list

  

| Protein Coverage
| Supporting Peptides
|

Protein Coverage:

Supporting Peptides:

| Peptide | Uniq | -10lgP | Mass | Length | ppm | m/z | z | RT | Fraction | Scan | Source File | Area Sample 2 | #Feature | #Feature Sample 2 | Start | End | PTM | AScore | Found By |
| --- | --- | --- | --- | --- | --- | --- | --- | --- | --- | --- | --- | --- | --- | --- | --- | --- | --- | --- | --- |
| K.ASTVADSIVDALEK.R | Y | 45.96 | 1417.7300 | 14 | 3.3 | 709.8746 | 2 | 33.28 | 2 | 7574 | JVDM\_TOPE.raw | 4.1721E4 | 1 | 1 | 612 | 625 |  |  | PEAKS DB |
| R.TTLVNLFSNLEK.E | Y | 42.92 | 1377.7504 | 12 | 1.3 | 689.8834 | 2 | 37.21 | 2 | 8627 | JVDM\_TOPE.raw | 3.2444E4 | 1 | 1 | 43 | 54 |  |  | PEAKS DB |
| K.NPLVNNDFENAEGLTK.L | Y | 42.35 | 1773.8533 | 16 | 0.9 | 887.9348 | 2 | 25.22 | 2 | 5733 | JVDM\_TOPE.raw | 2.0028E4 | 1 | 1 | 555 | 570 |  |  | PEAKS DB |
| R.YLNIISK.G | Y | 17.03 | 849.4960 | 7 | 4.7 | 425.7573 | 2 | 21.40 | 2 | 4622 | JVDM\_TOPE.raw | 2.8893E5 | 1 | 1 | 221 | 227 |  |  | PEAKS DB |
| total 4 peptides |  |  |  |  |  |  |  |  |  |  |  |  |  |  |  |  |  |  |  |
| --- | --- | --- | --- | --- | --- | --- | --- | --- | --- | --- | --- | --- | --- | --- | --- | --- | --- | --- | --- |

tr|A0A2K3D661|A0A2K3D661\_CHLRE

back to list

  

| Protein Coverage
| Supporting Peptides
|

Protein Coverage:

Supporting Peptides:

| Peptide | Uniq | -10lgP | Mass | Length | ppm | m/z | z | RT | Fraction | Scan | Source File | Area Sample 2 | #Feature | #Feature Sample 2 | Start | End | PTM | AScore | Found By |
| --- | --- | --- | --- | --- | --- | --- | --- | --- | --- | --- | --- | --- | --- | --- | --- | --- | --- | --- | --- |
| R.YEDNFDAVNNLVVIAQDTDKK.A | Y | 61.17 | 2410.1653 | 21 | 2.2 | 804.3975 | 3 | 32.79 | 2 | 7436 | JVDM\_TOPE.raw | 2.4859E5 | 1 | 1 | 106 | 126 |  |  | PEAKS DB |
| R.VSAASLLDVSTTTDKK.G | Y | 35.66 | 1634.8727 | 16 | 1.6 | 545.9657 | 3 | 24.14 | 2 | 5434 | JVDM\_TOPE.raw | 2.2817E5 | 1 | 1 | 165 | 180 |  |  | PEAKS DB |
| total 2 peptides |  |  |  |  |  |  |  |  |  |  |  |  |  |  |  |  |  |  |  |
| --- | --- | --- | --- | --- | --- | --- | --- | --- | --- | --- | --- | --- | --- | --- | --- | --- | --- | --- | --- |

tr|A8J363|A8J363\_CHLRE

back to list

  

| Protein Coverage
| Supporting Peptides
|

Protein Coverage:

Supporting Peptides:

| Peptide | Uniq | -10lgP | Mass | Length | ppm | m/z | z | RT | Fraction | Scan | Source File | Area Sample 2 | #Feature | #Feature Sample 2 | Start | End | PTM | AScore | Found By |
| --- | --- | --- | --- | --- | --- | --- | --- | --- | --- | --- | --- | --- | --- | --- | --- | --- | --- | --- | --- |
| R.VAIDQTGIATTDR.L | Y | 57.10 | 1359.6993 | 13 | 1.7 | 680.8581 | 2 | 17.14 | 2 | 3285 | JVDM\_TOPE.raw | 9.5905E4 | 1 | 1 | 525 | 537 |  |  | PEAKS DB |
| R.VAQPLGGYDSGLPK.E | Y | 41.15 | 1400.7300 | 14 | -1.3 | 701.3713 | 2 | 22.16 | 2 | 4865 | JVDM\_TOPE.raw | 8.6358E4 | 1 | 1 | 550 | 563 |  |  | PEAKS DB |
| total 2 peptides |  |  |  |  |  |  |  |  |  |  |  |  |  |  |  |  |  |  |  |
| --- | --- | --- | --- | --- | --- | --- | --- | --- | --- | --- | --- | --- | --- | --- | --- | --- | --- | --- | --- |

tr|A8JAV1|A8JAV1\_CHLRE

back to list

  

| Protein Coverage
| Supporting Peptides
|

Protein Coverage:

Supporting Peptides:

| Peptide | Uniq | -10lgP | Mass | Length | ppm | m/z | z | RT | Fraction | Scan | Source File | Area Sample 2 | #Feature | #Feature Sample 2 | Start | End | PTM | AScore | Found By |
| --- | --- | --- | --- | --- | --- | --- | --- | --- | --- | --- | --- | --- | --- | --- | --- | --- | --- | --- | --- |
| R.VAPEEHPVLLTEAPLNPK.A | Y | 70.14 | 1953.0570 | 18 | 4.1 | 977.5398 | 2 | 26.13 | 2 | 5946 | JVDM\_TOPE.raw | 1.3114E6 | 1 | 1 | 98 | 115 |  |  | PEAKS DB |
| R.GYSFTTTAER.E | Y | 22.09 | 1131.5197 | 10 | 1.1 | 566.7677 | 2 | 16.01 | 2 | 2898 | JVDM\_TOPE.raw | 2.405E5 | 1 | 1 | 199 | 208 |  |  | PEAKS DB |
| total 2 peptides |  |  |  |  |  |  |  |  |  |  |  |  |  |  |  |  |  |  |  |
| --- | --- | --- | --- | --- | --- | --- | --- | --- | --- | --- | --- | --- | --- | --- | --- | --- | --- | --- | --- |

tr|A8I2V3|A8I2V3\_CHLRE

back to list

  

| Protein Coverage
| Supporting Peptides
|

Protein Coverage:

Supporting Peptides:

| Peptide | Uniq | -10lgP | Mass | Length | ppm | m/z | z | RT | Fraction | Scan | Source File | Area Sample 2 | #Feature | #Feature Sample 2 | Start | End | PTM | AScore | Found By |
| --- | --- | --- | --- | --- | --- | --- | --- | --- | --- | --- | --- | --- | --- | --- | --- | --- | --- | --- | --- |
| K.DYGVLIEDGPDAGVTLR.G | Y | 51.51 | 1788.8893 | 17 | 1.1 | 895.4529 | 2 | 30.59 | 2 | 6987 | JVDM\_TOPE.raw | 2.5548E4 | 1 | 1 | 111 | 127 |  |  | PEAKS DB |
| R.GLFIISPTGVLR.Q | Y | 31.65 | 1271.7601 | 12 | 2.1 | 636.8887 | 2 | 37.34 | 2 | 8653 | JVDM\_TOPE.raw | 1.0668E5 | 1 | 1 | 128 | 139 |  |  | PEAKS DB |
| total 2 peptides |  |  |  |  |  |  |  |  |  |  |  |  |  |  |  |  |  |  |  |
| --- | --- | --- | --- | --- | --- | --- | --- | --- | --- | --- | --- | --- | --- | --- | --- | --- | --- | --- | --- |

tr|A8J244|A8J244\_CHLRE

back to list

  

| Protein Coverage
| Supporting Peptides
|

Protein Coverage:

Supporting Peptides:

| Peptide | Uniq | -10lgP | Mass | Length | ppm | m/z | z | RT | Fraction | Scan | Source File | Area Sample 2 | #Feature | #Feature Sample 2 | Start | End | PTM | AScore | Found By |
| --- | --- | --- | --- | --- | --- | --- | --- | --- | --- | --- | --- | --- | --- | --- | --- | --- | --- | --- | --- |
| R.LAADVMDVPTLIIAR.T | Y | 66.34 | 1596.8909 | 15 | 3.3 | 799.4553 | 2 | 37.77 | 2 | 8762 | JVDM\_TOPE.raw | 5.2588E4 | 1 | 1 | 201 | 215 |  |  | PEAKS DB |
| K.FAAAIHAQYPGK.L | Y | 26.30 | 1272.6615 | 12 | 1.9 | 425.2286 | 3 | 16.51 | 2 | 3050 | JVDM\_TOPE.raw | 1.6491E5 | 1 | 1 | 284 | 295 |  |  | PEAKS DB |
| total 2 peptides |  |  |  |  |  |  |  |  |  |  |  |  |  |  |  |  |  |  |  |
| --- | --- | --- | --- | --- | --- | --- | --- | --- | --- | --- | --- | --- | --- | --- | --- | --- | --- | --- | --- |

tr|A0A2K3CWF7|A0A2K3CWF7\_CHLRE

back to list

  

| Protein Coverage
| Supporting Peptides
|

Protein Coverage:

Supporting Peptides:

| Peptide | Uniq | -10lgP | Mass | Length | ppm | m/z | z | RT | Fraction | Scan | Source File | Area Sample 2 | #Feature | #Feature Sample 2 | Start | End | PTM | AScore | Found By |
| --- | --- | --- | --- | --- | --- | --- | --- | --- | --- | --- | --- | --- | --- | --- | --- | --- | --- | --- | --- |
| K.IEDLSAQTQAAAAEQFK.M | Y | 66.12 | 1819.8951 | 17 | 3.3 | 910.9578 | 2 | 22.90 | 2 | 5102 | JVDM\_TOPE.raw | 3.2087E4 | 1 | 1 | 105 | 121 |  |  | PEAKS DB |
| total 1 peptides |  |  |  |  |  |  |  |  |  |  |  |  |  |  |  |  |  |  |  |
| --- | --- | --- | --- | --- | --- | --- | --- | --- | --- | --- | --- | --- | --- | --- | --- | --- | --- | --- | --- |

tr|Q7X7A7|Q7X7A7\_CHLRE

back to list

  

| Protein Coverage
| Supporting Peptides
|

Protein Coverage:

Supporting Peptides:

| Peptide | Uniq | -10lgP | Mass | Length | ppm | m/z | z | RT | Fraction | Scan | Source File | Area Sample 2 | #Feature | #Feature Sample 2 | Start | End | PTM | AScore | Found By |
| --- | --- | --- | --- | --- | --- | --- | --- | --- | --- | --- | --- | --- | --- | --- | --- | --- | --- | --- | --- |
| K.AAAAELTLAAYK.V | Y | 45.17 | 1191.6499 | 12 | 1.5 | 596.8331 | 2 | 23.81 | 2 | 5355 | JVDM\_TOPE.raw | 2.1097E5 | 1 | 1 | 147 | 158 |  |  | PEAKS DB |
| R.ILQSIEQSEQAK.G | Y | 39.89 | 1372.7197 | 12 | 1.3 | 687.3680 | 2 | 15.30 | 2 | 2698 | JVDM\_TOPE.raw | 6.568E4 | 1 | 1 | 67 | 78 |  |  | PEAKS DB |
| K.VAVLANEQELSVEER.N | Y | 28.91 | 1684.8632 | 15 | -0.3 | 843.4386 | 2 | 23.18 | 2 | 5194 | JVDM\_TOPE.raw | 6.7261E4 | 1 | 1 | 33 | 47 |  |  | PEAKS DB |
| K.ADELPTTNPIR.L | Y | 22.92 | 1225.6302 | 11 | 3.3 | 613.8244 | 2 | 19.14 | 2 | 3951 | JVDM\_TOPE.raw | 0 | 0 | 0 | 164 | 174 |  |  | PEAKS DB |
| total 4 peptides |  |  |  |  |  |  |  |  |  |  |  |  |  |  |  |  |  |  |  |
| --- | --- | --- | --- | --- | --- | --- | --- | --- | --- | --- | --- | --- | --- | --- | --- | --- | --- | --- | --- |

tr|A0A2K3DP15|A0A2K3DP15\_CHLRE

back to list

  

| Protein Coverage
| Supporting Peptides
|

Protein Coverage:

Supporting Peptides:

| Peptide | Uniq | -10lgP | Mass | Length | ppm | m/z | z | RT | Fraction | Scan | Source File | Area Sample 2 | #Feature | #Feature Sample 2 | Start | End | PTM | AScore | Found By |
| --- | --- | --- | --- | --- | --- | --- | --- | --- | --- | --- | --- | --- | --- | --- | --- | --- | --- | --- | --- |
| R.LALALGDEVGQPLPLAAASNAQYIAAR.R | Y | 62.51 | 2692.4548 | 27 | 3.0 | 898.4949 | 3 | 37.69 | 2 | 8745 | JVDM\_TOPE.raw | 5.061E4 | 1 | 1 | 327 | 353 |  |  | PEAKS DB |
| total 1 peptides |  |  |  |  |  |  |  |  |  |  |  |  |  |  |  |  |  |  |  |
| --- | --- | --- | --- | --- | --- | --- | --- | --- | --- | --- | --- | --- | --- | --- | --- | --- | --- | --- | --- |

tr|A8HRZ0|A8HRZ0\_CHLRE

back to list

  

| Protein Coverage
| Supporting Peptides
|

Protein Coverage:

Supporting Peptides:

| Peptide | Uniq | -10lgP | Mass | Length | ppm | m/z | z | RT | Fraction | Scan | Source File | Area Sample 2 | #Feature | #Feature Sample 2 | Start | End | PTM | AScore | Found By |
| --- | --- | --- | --- | --- | --- | --- | --- | --- | --- | --- | --- | --- | --- | --- | --- | --- | --- | --- | --- |
| M.ADETVAVEEVAAAEPASGK.K | Y | 59.09 | 1842.8846 | 19 | 2.1 | 922.4515 | 2 | 21.98 | 2 | 4833 | JVDM\_TOPE.raw | 2.0515E4 | 1 | 1 | 2 | 20 |  |  | PEAKS DB |
| R.TGSSVPAIK.K | Y | 31.17 | 858.4811 | 9 | 3.4 | 430.2493 | 2 | 12.91 | 2 | 1936 | JVDM\_TOPE.raw | 1.1617E5 | 1 | 1 | 65 | 73 |  |  | PEAKS DB |
| M.ADETVAVEEVAAAEPASGKK.A | Y | 22.75 | 1970.9796 | 20 | 2.1 | 658.0018 | 3 | 19.97 | 2 | 4172 | JVDM\_TOPE.raw | 7.7876E5 | 1 | 1 | 2 | 21 |  |  | PEAKS DB |
| total 3 peptides |  |  |  |  |  |  |  |  |  |  |  |  |  |  |  |  |  |  |  |
| --- | --- | --- | --- | --- | --- | --- | --- | --- | --- | --- | --- | --- | --- | --- | --- | --- | --- | --- | --- |

tr|A0A2K3CQL0|A0A2K3CQL0\_CHLRE

back to list

  

| Protein Coverage
| Supporting Peptides
|

Protein Coverage:

Supporting Peptides:

| Peptide | Uniq | -10lgP | Mass | Length | ppm | m/z | z | RT | Fraction | Scan | Source File | Area Sample 2 | #Feature | #Feature Sample 2 | Start | End | PTM | AScore | Found By |
| --- | --- | --- | --- | --- | --- | --- | --- | --- | --- | --- | --- | --- | --- | --- | --- | --- | --- | --- | --- |
| R.LLAAGQGEAVVK.E | Y | 38.56 | 1154.6659 | 12 | 2.3 | 578.3416 | 2 | 17.28 | 2 | 3314 | JVDM\_TOPE.raw | 2.0075E5 | 1 | 1 | 64 | 75 |  |  | PEAKS DB |
| R.AALAAVPAVC(+57.02)R.T | Y | 38.37 | 1097.6016 | 11 | 3.0 | 549.8097 | 2 | 20.00 | 2 | 4223 | JVDM\_TOPE.raw | 1.0497E5 | 1 | 1 | 110 | 120 | Carbamidomethylation | C10:Carbamidomethylation:1000.00 | PEAKS DB |
| total 2 peptides |  |  |  |  |  |  |  |  |  |  |  |  |  |  |  |  |  |  |  |
| --- | --- | --- | --- | --- | --- | --- | --- | --- | --- | --- | --- | --- | --- | --- | --- | --- | --- | --- | --- |

tr|A0A2K3CQP7|A0A2K3CQP7\_CHLRE

back to list

  

| Protein Coverage
| Supporting Peptides
|

Protein Coverage:

Supporting Peptides:

| Peptide | Uniq | -10lgP | Mass | Length | ppm | m/z | z | RT | Fraction | Scan | Source File | Area Sample 2 | #Feature | #Feature Sample 2 | Start | End | PTM | AScore | Found By |
| --- | --- | --- | --- | --- | --- | --- | --- | --- | --- | --- | --- | --- | --- | --- | --- | --- | --- | --- | --- |
| R.LLAAGQGEAVVK.E | Y | 38.56 | 1154.6659 | 12 | 2.3 | 578.3416 | 2 | 17.28 | 2 | 3314 | JVDM\_TOPE.raw | 2.0075E5 | 1 | 1 | 83 | 94 |  |  | PEAKS DB |
| R.AALAAVPAVC(+57.02)R.T | Y | 38.37 | 1097.6016 | 11 | 3.0 | 549.8097 | 2 | 20.00 | 2 | 4223 | JVDM\_TOPE.raw | 1.0497E5 | 1 | 1 | 129 | 139 | Carbamidomethylation | C10:Carbamidomethylation:1000.00 | PEAKS DB |
| total 2 peptides |  |  |  |  |  |  |  |  |  |  |  |  |  |  |  |  |  |  |  |
| --- | --- | --- | --- | --- | --- | --- | --- | --- | --- | --- | --- | --- | --- | --- | --- | --- | --- | --- | --- |

tr|A0A2K3E350|A0A2K3E350\_CHLRE

back to list

  

| Protein Coverage
| Supporting Peptides
|

Protein Coverage:

Supporting Peptides:

| Peptide | Uniq | -10lgP | Mass | Length | ppm | m/z | z | RT | Fraction | Scan | Source File | Area Sample 2 | #Feature | #Feature Sample 2 | Start | End | PTM | AScore | Found By |
| --- | --- | --- | --- | --- | --- | --- | --- | --- | --- | --- | --- | --- | --- | --- | --- | --- | --- | --- | --- |
| K.IGSLLDQSITR.H | Y | 57.14 | 1201.6666 | 11 | 0.2 | 601.8407 | 2 | 24.69 | 2 | 5594 | JVDM\_TOPE.raw | 4.4689E4 | 1 | 1 | 136 | 146 |  |  | PEAKS DB |
| total 1 peptides |  |  |  |  |  |  |  |  |  |  |  |  |  |  |  |  |  |  |  |
| --- | --- | --- | --- | --- | --- | --- | --- | --- | --- | --- | --- | --- | --- | --- | --- | --- | --- | --- | --- |

tr|Q6UKY5|Q6UKY5\_CHLRE

back to list

  

| Protein Coverage
| Supporting Peptides
|

Protein Coverage:

Supporting Peptides:

| Peptide | Uniq | -10lgP | Mass | Length | ppm | m/z | z | RT | Fraction | Scan | Source File | Area Sample 2 | #Feature | #Feature Sample 2 | Start | End | PTM | AScore | Found By |
| --- | --- | --- | --- | --- | --- | --- | --- | --- | --- | --- | --- | --- | --- | --- | --- | --- | --- | --- | --- |
| R.SIISTQLGTELEK.V | Y | 56.52 | 1417.7664 | 13 | 2.1 | 709.8920 | 2 | 26.83 | 2 | 6082 | JVDM\_TOPE.raw | 1.8672E5 | 1 | 1 | 46 | 58 |  |  | PEAKS DB |
| total 1 peptides |  |  |  |  |  |  |  |  |  |  |  |  |  |  |  |  |  |  |  |
| --- | --- | --- | --- | --- | --- | --- | --- | --- | --- | --- | --- | --- | --- | --- | --- | --- | --- | --- | --- |

tr|A8J1U1|A8J1U1\_CHLRE

back to list

  

| Protein Coverage
| Supporting Peptides
|

Protein Coverage:

Supporting Peptides:

| Peptide | Uniq | -10lgP | Mass | Length | ppm | m/z | z | RT | Fraction | Scan | Source File | Area Sample 2 | #Feature | #Feature Sample 2 | Start | End | PTM | AScore | Found By |
| --- | --- | --- | --- | --- | --- | --- | --- | --- | --- | --- | --- | --- | --- | --- | --- | --- | --- | --- | --- |
| K.GIVDSEDLPLNISR.E | Y | 53.87 | 1526.7939 | 14 | 2.5 | 764.4061 | 2 | 29.52 | 2 | 6715 | JVDM\_TOPE.raw | 0 | 0 | 0 | 360 | 373 |  |  | PEAKS DB |
| total 1 peptides |  |  |  |  |  |  |  |  |  |  |  |  |  |  |  |  |  |  |  |
| --- | --- | --- | --- | --- | --- | --- | --- | --- | --- | --- | --- | --- | --- | --- | --- | --- | --- | --- | --- |

tr|Q9LD42|Q9LD42\_CHLRE

back to list

  

| Protein Coverage
| Supporting Peptides
|

Protein Coverage:

Supporting Peptides:

| Peptide | Uniq | -10lgP | Mass | Length | ppm | m/z | z | RT | Fraction | Scan | Source File | Area Sample 2 | #Feature | #Feature Sample 2 | Start | End | PTM | AScore | Found By |
| --- | --- | --- | --- | --- | --- | --- | --- | --- | --- | --- | --- | --- | --- | --- | --- | --- | --- | --- | --- |
| R.LSTLNIQAYDAAR.T | Y | 53.09 | 1434.7467 | 13 | 2.2 | 718.3822 | 2 | 24.56 | 2 | 5549 | JVDM\_TOPE.raw | 6.5093E4 | 1 | 1 | 229 | 241 |  |  | PEAKS DB |
| total 1 peptides |  |  |  |  |  |  |  |  |  |  |  |  |  |  |  |  |  |  |  |
| --- | --- | --- | --- | --- | --- | --- | --- | --- | --- | --- | --- | --- | --- | --- | --- | --- | --- | --- | --- |

tr|A8HVQ1|A8HVQ1\_CHLRE

back to list

  

| Protein Coverage
| Supporting Peptides
|

Protein Coverage:

Supporting Peptides:

| Peptide | Uniq | -10lgP | Mass | Length | ppm | m/z | z | RT | Fraction | Scan | Source File | Area Sample 2 | #Feature | #Feature Sample 2 | Start | End | PTM | AScore | Found By |
| --- | --- | --- | --- | --- | --- | --- | --- | --- | --- | --- | --- | --- | --- | --- | --- | --- | --- | --- | --- |
| K.NAIVQVDATPFK.Q | Y | 53.07 | 1301.6979 | 12 | 2.4 | 651.8578 | 2 | 25.75 | 2 | 5842 | JVDM\_TOPE.raw | 1.0858E5 | 1 | 1 | 100 | 111 |  |  | PEAKS DB |
| total 1 peptides |  |  |  |  |  |  |  |  |  |  |  |  |  |  |  |  |  |  |  |
| --- | --- | --- | --- | --- | --- | --- | --- | --- | --- | --- | --- | --- | --- | --- | --- | --- | --- | --- | --- |

tr|A0A2K3E7M7|A0A2K3E7M7\_CHLRE

back to list

  

| Protein Coverage
| Supporting Peptides
|

Protein Coverage:

Supporting Peptides:

| Peptide | Uniq | -10lgP | Mass | Length | ppm | m/z | z | RT | Fraction | Scan | Source File | Area Sample 2 | #Feature | #Feature Sample 2 | Start | End | PTM | AScore | Found By |
| --- | --- | --- | --- | --- | --- | --- | --- | --- | --- | --- | --- | --- | --- | --- | --- | --- | --- | --- | --- |
| K.AGAAASAVQSTAASAVEK.A | Y | 51.60 | 1588.8057 | 18 | 1.2 | 795.4111 | 2 | 16.39 | 2 | 3021 | JVDM\_TOPE.raw | 2.7426E3 | 1 | 1 | 166 | 183 |  |  | PEAKS DB |
| K.AGAAASAAQSSAAAAVEKASG.A | Y | 16.34 | 1745.8544 | 21 | 9.5 | 873.9427 | 2 | 26.96 | 2 | 6120 | JVDM\_TOPE.raw | 0 | 0 | 0 | 238 | 258 |  |  | PEAKS DB |
| K.AGAAAS(+79.97)AAQSTAASAVEKAGSLTQR.A | Y | 15.85 | 2354.1226 | 25 | 3.6 | 785.7176 | 3 | 34.77 | 2 | 7989 | JVDM\_TOPE.raw | 0 | 0 | 0 | 382 | 406 | Phosphorylation (STY) | S6:Phosphorylation (STY):0.00 | PEAKS PTM |
| K.AGAAASAAQ(+.98)STAASAVEKASGAATAAQ(+.98)AAAGSAAEK.A | Y | 15.82 | 3047.4641 | 36 | 8.0 | 1016.8368 | 3 | 33.66 | 2 | 7789 | JVDM\_TOPE.raw | 3.8661E5 | 1 | 1 | 58 | 93 | Deamidation (NQ) | Q9:Deamidation (NQ):1000.00;Q27:Deamidation (NQ):1000.00 | PEAKS PTM |
| total 4 peptides |  |  |  |  |  |  |  |  |  |  |  |  |  |  |  |  |  |  |  |
| --- | --- | --- | --- | --- | --- | --- | --- | --- | --- | --- | --- | --- | --- | --- | --- | --- | --- | --- | --- |

tr|A0A2K3CRJ3|A0A2K3CRJ3\_CHLRE

back to list

  

| Protein Coverage
| Supporting Peptides
|

Protein Coverage:

Supporting Peptides:

| Peptide | Uniq | -10lgP | Mass | Length | ppm | m/z | z | RT | Fraction | Scan | Source File | Area Sample 2 | #Feature | #Feature Sample 2 | Start | End | PTM | AScore | Found By |
| --- | --- | --- | --- | --- | --- | --- | --- | --- | --- | --- | --- | --- | --- | --- | --- | --- | --- | --- | --- |
| R.GLGPGGVELNK.Q | Y | 50.86 | 1039.5662 | 11 | 0.8 | 520.7908 | 2 | 18.12 | 2 | 3574 | JVDM\_TOPE.raw | 7.1637E5 | 1 | 1 | 119 | 129 |  |  | PEAKS DB |
| total 1 peptides |  |  |  |  |  |  |  |  |  |  |  |  |  |  |  |  |  |  |  |
| --- | --- | --- | --- | --- | --- | --- | --- | --- | --- | --- | --- | --- | --- | --- | --- | --- | --- | --- | --- |

tr|Q1ALD7|Q1ALD7\_CHLRE

back to list

  

| Protein Coverage
| Supporting Peptides
|

Protein Coverage:

Supporting Peptides:

| Peptide | Uniq | -10lgP | Mass | Length | ppm | m/z | z | RT | Fraction | Scan | Source File | Area Sample 2 | #Feature | #Feature Sample 2 | Start | End | PTM | AScore | Found By |
| --- | --- | --- | --- | --- | --- | --- | --- | --- | --- | --- | --- | --- | --- | --- | --- | --- | --- | --- | --- |
| R.GLGPGGVELNK.Q | Y | 50.86 | 1039.5662 | 11 | 0.8 | 520.7908 | 2 | 18.12 | 2 | 3574 | JVDM\_TOPE.raw | 7.1637E5 | 1 | 1 | 119 | 129 |  |  | PEAKS DB |
| total 1 peptides |  |  |  |  |  |  |  |  |  |  |  |  |  |  |  |  |  |  |  |
| --- | --- | --- | --- | --- | --- | --- | --- | --- | --- | --- | --- | --- | --- | --- | --- | --- | --- | --- | --- |

tr|A0A2K3CRI8|A0A2K3CRI8\_CHLRE

back to list

  

| Protein Coverage
| Supporting Peptides
|

Protein Coverage:

Supporting Peptides:

| Peptide | Uniq | -10lgP | Mass | Length | ppm | m/z | z | RT | Fraction | Scan | Source File | Area Sample 2 | #Feature | #Feature Sample 2 | Start | End | PTM | AScore | Found By |
| --- | --- | --- | --- | --- | --- | --- | --- | --- | --- | --- | --- | --- | --- | --- | --- | --- | --- | --- | --- |
| R.GLGPGGVELNK.Q | Y | 50.86 | 1039.5662 | 11 | 0.8 | 520.7908 | 2 | 18.12 | 2 | 3574 | JVDM\_TOPE.raw | 7.1637E5 | 1 | 1 | 137 | 147 |  |  | PEAKS DB |
| total 1 peptides |  |  |  |  |  |  |  |  |  |  |  |  |  |  |  |  |  |  |  |
| --- | --- | --- | --- | --- | --- | --- | --- | --- | --- | --- | --- | --- | --- | --- | --- | --- | --- | --- | --- |

tr|A0A2K3DMT2|A0A2K3DMT2\_CHLRE

back to list

  

| Protein Coverage
| Supporting Peptides
|

Protein Coverage:

Supporting Peptides:

| Peptide | Uniq | -10lgP | Mass | Length | ppm | m/z | z | RT | Fraction | Scan | Source File | Area Sample 2 | #Feature | #Feature Sample 2 | Start | End | PTM | AScore | Found By |
| --- | --- | --- | --- | --- | --- | --- | --- | --- | --- | --- | --- | --- | --- | --- | --- | --- | --- | --- | --- |
| K.AAATGAAAAAGSQLASAQEAGK.A | Y | 50.67 | 1871.9336 | 22 | 0.9 | 624.9857 | 3 | 16.37 | 2 | 3018 | JVDM\_TOPE.raw | 0 | 0 | 0 | 156 | 177 |  |  | PEAKS DB |
| total 1 peptides |  |  |  |  |  |  |  |  |  |  |  |  |  |  |  |  |  |  |  |
| --- | --- | --- | --- | --- | --- | --- | --- | --- | --- | --- | --- | --- | --- | --- | --- | --- | --- | --- | --- |

tr|A8J1A3|A8J1A3\_CHLRE

back to list

  

| Protein Coverage
| Supporting Peptides
|

Protein Coverage:

Supporting Peptides:

| Peptide | Uniq | -10lgP | Mass | Length | ppm | m/z | z | RT | Fraction | Scan | Source File | Area Sample 2 | #Feature | #Feature Sample 2 | Start | End | PTM | AScore | Found By |
| --- | --- | --- | --- | --- | --- | --- | --- | --- | --- | --- | --- | --- | --- | --- | --- | --- | --- | --- | --- |
| R.SIVGTSLEVIQK.K | Y | 49.93 | 1272.7289 | 12 | 2.5 | 637.3733 | 2 | 26.87 | 2 | 6099 | JVDM\_TOPE.raw | 2.2313E5 | 1 | 1 | 84 | 95 |  |  | PEAKS DB |
| total 1 peptides |  |  |  |  |  |  |  |  |  |  |  |  |  |  |  |  |  |  |  |
| --- | --- | --- | --- | --- | --- | --- | --- | --- | --- | --- | --- | --- | --- | --- | --- | --- | --- | --- | --- |

tr|A0A2K3DE61|A0A2K3DE61\_CHLRE

back to list

  

| Protein Coverage
| Supporting Peptides
|

Protein Coverage:

Supporting Peptides:

| Peptide | Uniq | -10lgP | Mass | Length | ppm | m/z | z | RT | Fraction | Scan | Source File | Area Sample 2 | #Feature | #Feature Sample 2 | Start | End | PTM | AScore | Found By |
| --- | --- | --- | --- | --- | --- | --- | --- | --- | --- | --- | --- | --- | --- | --- | --- | --- | --- | --- | --- |
| R.SIVGTSLEVIQK.K | Y | 49.93 | 1272.7289 | 12 | 2.5 | 637.3733 | 2 | 26.87 | 2 | 6099 | JVDM\_TOPE.raw | 2.2313E5 | 1 | 1 | 108 | 119 |  |  | PEAKS DB |
| total 1 peptides |  |  |  |  |  |  |  |  |  |  |  |  |  |  |  |  |  |  |  |
| --- | --- | --- | --- | --- | --- | --- | --- | --- | --- | --- | --- | --- | --- | --- | --- | --- | --- | --- | --- |

tr|A0A2K3DZB3|A0A2K3DZB3\_CHLRE

back to list

  

| Protein Coverage
| Supporting Peptides
|

Protein Coverage:

Supporting Peptides:

| Peptide | Uniq | -10lgP | Mass | Length | ppm | m/z | z | RT | Fraction | Scan | Source File | Area Sample 2 | #Feature | #Feature Sample 2 | Start | End | PTM | AScore | Found By |
| --- | --- | --- | --- | --- | --- | --- | --- | --- | --- | --- | --- | --- | --- | --- | --- | --- | --- | --- | --- |
| K.LFASLGNPVSLEK.L | Y | 43.07 | 1373.7554 | 13 | 1.7 | 687.8861 | 2 | 29.92 | 2 | 6812 | JVDM\_TOPE.raw | 0 | 0 | 0 | 159 | 171 |  |  | PEAKS DB |
| total 1 peptides |  |  |  |  |  |  |  |  |  |  |  |  |  |  |  |  |  |  |  |
| --- | --- | --- | --- | --- | --- | --- | --- | --- | --- | --- | --- | --- | --- | --- | --- | --- | --- | --- | --- |

tr|A8IV40|A8IV40\_CHLRE

back to list

  

| Protein Coverage
| Supporting Peptides
|

Protein Coverage:

Supporting Peptides:

| Peptide | Uniq | -10lgP | Mass | Length | ppm | m/z | z | RT | Fraction | Scan | Source File | Area Sample 2 | #Feature | #Feature Sample 2 | Start | End | PTM | AScore | Found By |
| --- | --- | --- | --- | --- | --- | --- | --- | --- | --- | --- | --- | --- | --- | --- | --- | --- | --- | --- | --- |
| D.AAEEAGLDLPYSC(+57.02)R.A | Y | 42.06 | 1550.7035 | 14 | 2.6 | 776.3610 | 2 | 23.53 | 2 | 5283 | JVDM\_TOPE.raw | 0 | 0 | 0 | 57 | 70 | Carbamidomethylation | C13:Carbamidomethylation:1000.00 | PEAKS DB |
| total 1 peptides |  |  |  |  |  |  |  |  |  |  |  |  |  |  |  |  |  |  |  |
| --- | --- | --- | --- | --- | --- | --- | --- | --- | --- | --- | --- | --- | --- | --- | --- | --- | --- | --- | --- |

tr|A0A2K3D7P8|A0A2K3D7P8\_CHLRE

back to list

  

| Protein Coverage
| Supporting Peptides
|

Protein Coverage:

Supporting Peptides:

| Peptide | Uniq | -10lgP | Mass | Length | ppm | m/z | z | RT | Fraction | Scan | Source File | Area Sample 2 | #Feature | #Feature Sample 2 | Start | End | PTM | AScore | Found By |
| --- | --- | --- | --- | --- | --- | --- | --- | --- | --- | --- | --- | --- | --- | --- | --- | --- | --- | --- | --- |
| R.LLAPSADVVAAGEAAQPHC(+57.02)SR.L | Y | 41.48 | 2119.0481 | 21 | 3.5 | 707.3591 | 3 | 22.84 | 2 | 5090 | JVDM\_TOPE.raw | 1.4618E4 | 1 | 1 | 1484 | 1504 | Carbamidomethylation | C19:Carbamidomethylation:1000.00 | PEAKS DB |
| total 1 peptides |  |  |  |  |  |  |  |  |  |  |  |  |  |  |  |  |  |  |  |
| --- | --- | --- | --- | --- | --- | --- | --- | --- | --- | --- | --- | --- | --- | --- | --- | --- | --- | --- | --- |

tr|A0A2K3CPA4|A0A2K3CPA4\_CHLRE

back to list

  

| Protein Coverage
| Supporting Peptides
|

Protein Coverage:

Supporting Peptides:

| Peptide | Uniq | -10lgP | Mass | Length | ppm | m/z | z | RT | Fraction | Scan | Source File | Area Sample 2 | #Feature | #Feature Sample 2 | Start | End | PTM | AScore | Found By |
| --- | --- | --- | --- | --- | --- | --- | --- | --- | --- | --- | --- | --- | --- | --- | --- | --- | --- | --- | --- |
| R.VLILDADNAGSAVR.T | Y | 39.81 | 1412.7623 | 14 | 3.3 | 707.3907 | 2 | 24.90 | 2 | 5659 | JVDM\_TOPE.raw | 8.3093E3 | 1 | 1 | 18 | 31 |  |  | PEAKS DB |
| total 1 peptides |  |  |  |  |  |  |  |  |  |  |  |  |  |  |  |  |  |  |  |
| --- | --- | --- | --- | --- | --- | --- | --- | --- | --- | --- | --- | --- | --- | --- | --- | --- | --- | --- | --- |

tr|O48949|O48949\_CHLRE

back to list

  

| Protein Coverage
| Supporting Peptides
|

Protein Coverage:

Supporting Peptides:

| Peptide | Uniq | -10lgP | Mass | Length | ppm | m/z | z | RT | Fraction | Scan | Source File | Area Sample 2 | #Feature | #Feature Sample 2 | Start | End | PTM | AScore | Found By |
| --- | --- | --- | --- | --- | --- | --- | --- | --- | --- | --- | --- | --- | --- | --- | --- | --- | --- | --- | --- |
| K.AAGLDAVDTVSVVK.N | Y | 38.48 | 1343.7296 | 14 | 4.5 | 672.8751 | 2 | 24.68 | 2 | 5596 | JVDM\_TOPE.raw | 9.6926E4 | 1 | 1 | 218 | 231 |  |  | PEAKS DB |
| K.TGPPAVTVEDADKLK.S | Y | 27.07 | 1539.8143 | 15 | 1.2 | 514.2794 | 3 | 18.99 | 2 | 3898 | JVDM\_TOPE.raw | 9.9186E4 | 1 | 1 | 156 | 170 |  |  | PEAKS DB |
| K.TEDVVFVQTTSADVAK.A | Y | 19.58 | 1708.8519 | 16 | 0.1 | 855.4333 | 2 | 23.91 | 2 | 5371 | JVDM\_TOPE.raw | 0 | 0 | 0 | 202 | 217 |  |  | PEAKS DB |
| R.ATAVLATDIDTDSLTAFVK.S | Y | 17.12 | 1951.0149 | 19 | 2.1 | 976.5167 | 2 | 35.63 | 2 | 8183 | JVDM\_TOPE.raw | 1.5207E4 | 1 | 1 | 239 | 257 |  |  | PEAKS DB |
| K.ALEGEIYDTFK.S | Y | 17.03 | 1284.6238 | 11 | 2.4 | 643.3207 | 2 | 27.26 | 2 | 6186 | JVDM\_TOPE.raw | 5.6701E4 | 1 | 1 | 186 | 196 |  |  | PEAKS DB |
| total 5 peptides |  |  |  |  |  |  |  |  |  |  |  |  |  |  |  |  |  |  |  |
| --- | --- | --- | --- | --- | --- | --- | --- | --- | --- | --- | --- | --- | --- | --- | --- | --- | --- | --- | --- |

tr|A8HMY6|A8HMY6\_CHLRE

back to list

  

| Protein Coverage
| Supporting Peptides
|

Protein Coverage:

Supporting Peptides:

| Peptide | Uniq | -10lgP | Mass | Length | ppm | m/z | z | RT | Fraction | Scan | Source File | Area Sample 2 | #Feature | #Feature Sample 2 | Start | End | PTM | AScore | Found By |
| --- | --- | --- | --- | --- | --- | --- | --- | --- | --- | --- | --- | --- | --- | --- | --- | --- | --- | --- | --- |
| R.TNIVLEATR.W | Y | 36.79 | 1015.5662 | 9 | 1.1 | 508.7909 | 2 | 19.58 | 2 | 4062 | JVDM\_TOPE.raw | 2.2058E5 | 1 | 1 | 154 | 162 |  |  | PEAKS DB |
| total 1 peptides |  |  |  |  |  |  |  |  |  |  |  |  |  |  |  |  |  |  |  |
| --- | --- | --- | --- | --- | --- | --- | --- | --- | --- | --- | --- | --- | --- | --- | --- | --- | --- | --- | --- |

tr|A0A2K3E3X9|A0A2K3E3X9\_CHLRE

back to list

  

| Protein Coverage
| Supporting Peptides
|

Protein Coverage:

Supporting Peptides:

| Peptide | Uniq | -10lgP | Mass | Length | ppm | m/z | z | RT | Fraction | Scan | Source File | Area Sample 2 | #Feature | #Feature Sample 2 | Start | End | PTM | AScore | Found By |
| --- | --- | --- | --- | --- | --- | --- | --- | --- | --- | --- | --- | --- | --- | --- | --- | --- | --- | --- | --- |
| R.ALTSGVDAATAGR.Q | Y | 36.18 | 1188.6099 | 13 | 1.1 | 595.3129 | 2 | 13.37 | 2 | 2116 | JVDM\_TOPE.raw | 8.6069E4 | 1 | 1 | 150 | 162 |  |  | PEAKS DB |
| total 1 peptides |  |  |  |  |  |  |  |  |  |  |  |  |  |  |  |  |  |  |  |
| --- | --- | --- | --- | --- | --- | --- | --- | --- | --- | --- | --- | --- | --- | --- | --- | --- | --- | --- | --- |

tr|A8IXZ0|A8IXZ0\_CHLRE

back to list

  

| Protein Coverage
| Supporting Peptides
|

Protein Coverage:

Supporting Peptides:

| Peptide | Uniq | -10lgP | Mass | Length | ppm | m/z | z | RT | Fraction | Scan | Source File | Area Sample 2 | #Feature | #Feature Sample 2 | Start | End | PTM | AScore | Found By |
| --- | --- | --- | --- | --- | --- | --- | --- | --- | --- | --- | --- | --- | --- | --- | --- | --- | --- | --- | --- |
| K.LAVNLIPFPR.L | Y | 35.27 | 1138.6863 | 10 | 2.1 | 570.3516 | 2 | 35.74 | 2 | 8215 | JVDM\_TOPE.raw | 9.359E4 | 1 | 1 | 253 | 262 |  |  | PEAKS DB |
| total 1 peptides |  |  |  |  |  |  |  |  |  |  |  |  |  |  |  |  |  |  |  |
| --- | --- | --- | --- | --- | --- | --- | --- | --- | --- | --- | --- | --- | --- | --- | --- | --- | --- | --- | --- |

tr|A0A2K3DFK4|A0A2K3DFK4\_CHLRE

back to list

  

| Protein Coverage
| Supporting Peptides
|

Protein Coverage:

Supporting Peptides:

| Peptide | Uniq | -10lgP | Mass | Length | ppm | m/z | z | RT | Fraction | Scan | Source File | Area Sample 2 | #Feature | #Feature Sample 2 | Start | End | PTM | AScore | Found By |
| --- | --- | --- | --- | --- | --- | --- | --- | --- | --- | --- | --- | --- | --- | --- | --- | --- | --- | --- | --- |
| K.AAPAANGGGASGAAAAVK.A | Y | 32.97 | 1410.7214 | 18 | 2.0 | 706.3694 | 2 | 20.31 | 2 | 4318 | JVDM\_TOPE.raw | 0 | 0 | 0 | 536 | 553 |  |  | PEAKS DB |
| R.GAPAVAAAAAAAAAGGGAGGGAGGAP.A | Y | 20.06 | 1818.8972 | 26 | -0.1 | 607.3063 | 3 | 23.31 | 2 | 5187 | JVDM\_TOPE.raw | 5.7435E6 | 1 | 1 | 1626 | 1651 |  |  | PEAKS DB |
| total 2 peptides |  |  |  |  |  |  |  |  |  |  |  |  |  |  |  |  |  |  |  |
| --- | --- | --- | --- | --- | --- | --- | --- | --- | --- | --- | --- | --- | --- | --- | --- | --- | --- | --- | --- |

tr|A0A2K3D7S0|A0A2K3D7S0\_CHLRE

back to list

  

| Protein Coverage
| Supporting Peptides
|

Protein Coverage:

Supporting Peptides:

| Peptide | Uniq | -10lgP | Mass | Length | ppm | m/z | z | RT | Fraction | Scan | Source File | Area Sample 2 | #Feature | #Feature Sample 2 | Start | End | PTM | AScore | Found By |
| --- | --- | --- | --- | --- | --- | --- | --- | --- | --- | --- | --- | --- | --- | --- | --- | --- | --- | --- | --- |
| K.RGSAAGGGAAAPPSLPGATAAATSSSGTGGGGGGGGR.R | Y | 32.62 | 2910.3928 | 37 | -4.0 | 728.6025 | 4 | 20.60 | 2 | 4404 | JVDM\_TOPE.raw | 8.5138E4 | 1 | 1 | 531 | 567 |  |  | PEAKS DB |
| total 1 peptides |  |  |  |  |  |  |  |  |  |  |  |  |  |  |  |  |  |  |  |
| --- | --- | --- | --- | --- | --- | --- | --- | --- | --- | --- | --- | --- | --- | --- | --- | --- | --- | --- | --- |

tr|A0A2K3E270|A0A2K3E270\_CHLRE

back to list

  

| Protein Coverage
| Supporting Peptides
|

Protein Coverage:

Supporting Peptides:

| Peptide | Uniq | -10lgP | Mass | Length | ppm | m/z | z | RT | Fraction | Scan | Source File | Area Sample 2 | #Feature | #Feature Sample 2 | Start | End | PTM | AScore | Found By |
| --- | --- | --- | --- | --- | --- | --- | --- | --- | --- | --- | --- | --- | --- | --- | --- | --- | --- | --- | --- |
| A.HDVGIGGGGNGGGGALSSGGPR.S | Y | 32.58 | 1834.8669 | 22 | 5.4 | 918.4457 | 2 | 20.83 | 2 | 4469 | JVDM\_TOPE.raw | 0 | 0 | 0 | 760 | 781 |  |  | PEAKS DB |
| total 1 peptides |  |  |  |  |  |  |  |  |  |  |  |  |  |  |  |  |  |  |  |
| --- | --- | --- | --- | --- | --- | --- | --- | --- | --- | --- | --- | --- | --- | --- | --- | --- | --- | --- | --- |

tr|A0A2K3E262|A0A2K3E262\_CHLRE

back to list

  

| Protein Coverage
| Supporting Peptides
|

Protein Coverage:

Supporting Peptides:

| Peptide | Uniq | -10lgP | Mass | Length | ppm | m/z | z | RT | Fraction | Scan | Source File | Area Sample 2 | #Feature | #Feature Sample 2 | Start | End | PTM | AScore | Found By |
| --- | --- | --- | --- | --- | --- | --- | --- | --- | --- | --- | --- | --- | --- | --- | --- | --- | --- | --- | --- |
| A.HDVGIGGGGNGGGGALSSGGPR.S | Y | 32.58 | 1834.8669 | 22 | 5.4 | 918.4457 | 2 | 20.83 | 2 | 4469 | JVDM\_TOPE.raw | 0 | 0 | 0 | 760 | 781 |  |  | PEAKS DB |
| total 1 peptides |  |  |  |  |  |  |  |  |  |  |  |  |  |  |  |  |  |  |  |
| --- | --- | --- | --- | --- | --- | --- | --- | --- | --- | --- | --- | --- | --- | --- | --- | --- | --- | --- | --- |

tr|A0A2K3CWQ4|A0A2K3CWQ4\_CHLRE

back to list

  

| Protein Coverage
| Supporting Peptides
|

Protein Coverage:

Supporting Peptides:

| Peptide | Uniq | -10lgP | Mass | Length | ppm | m/z | z | RT | Fraction | Scan | Source File | Area Sample 2 | #Feature | #Feature Sample 2 | Start | End | PTM | AScore | Found By |
| --- | --- | --- | --- | --- | --- | --- | --- | --- | --- | --- | --- | --- | --- | --- | --- | --- | --- | --- | --- |
| K.LESARGGGPAATAAAR.A | Y | 31.86 | 1454.7589 | 16 | 8.0 | 728.3926 | 2 | 17.13 | 2 | 3277 | JVDM\_TOPE.raw | 0 | 0 | 0 | 243 | 258 |  |  | PEAKS DB |
| total 1 peptides |  |  |  |  |  |  |  |  |  |  |  |  |  |  |  |  |  |  |  |
| --- | --- | --- | --- | --- | --- | --- | --- | --- | --- | --- | --- | --- | --- | --- | --- | --- | --- | --- | --- |

Peptide List

  
  

---

Prepared with PEAKS ™ (bioinfor.com)
