## Supplementary figures and images for "Efficient secretion of a plastic degrading enzyme from the green algae *Chlamydomonas reinhardtii*"

### cov_7.png

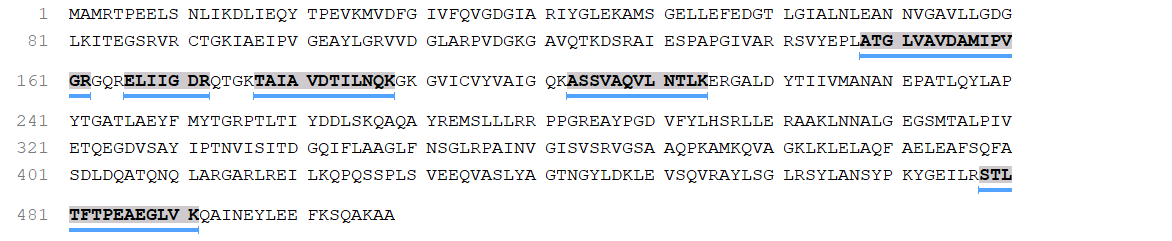

### cov_8.png

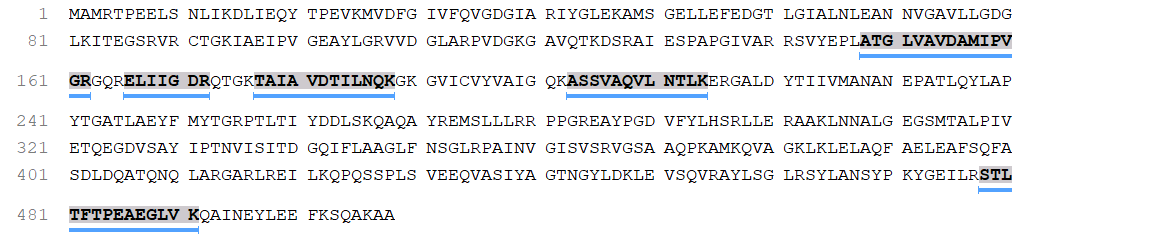

### cov_9.png

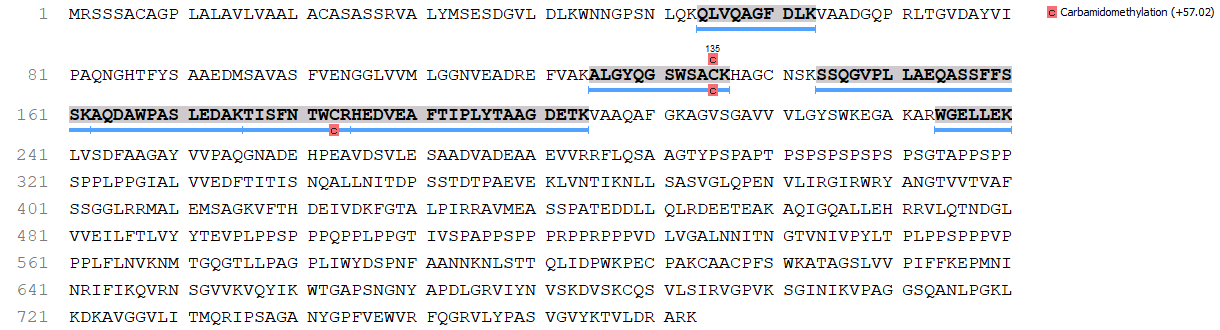

### cov_693.png

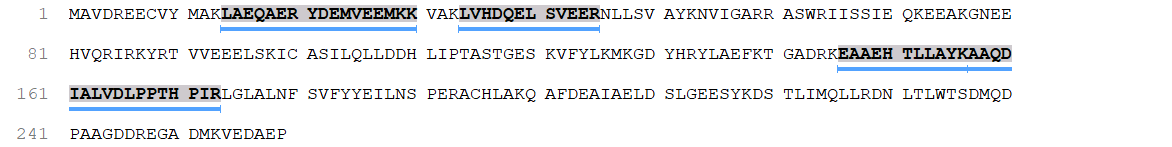

### cov_735.png

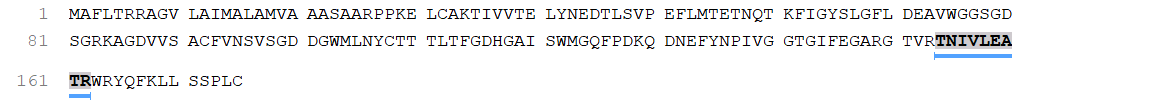

### cov_989.png

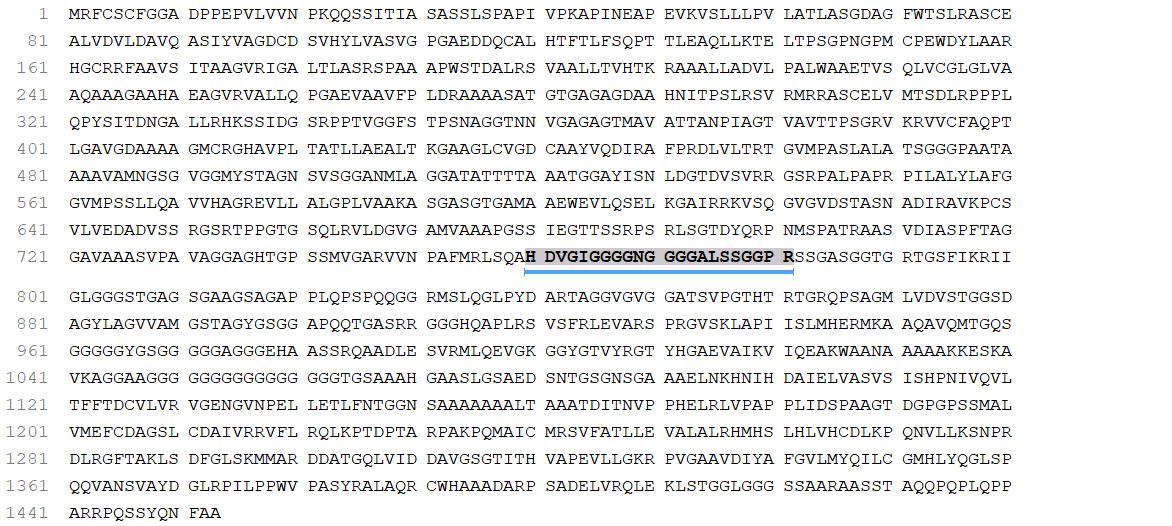

### cov_990.png

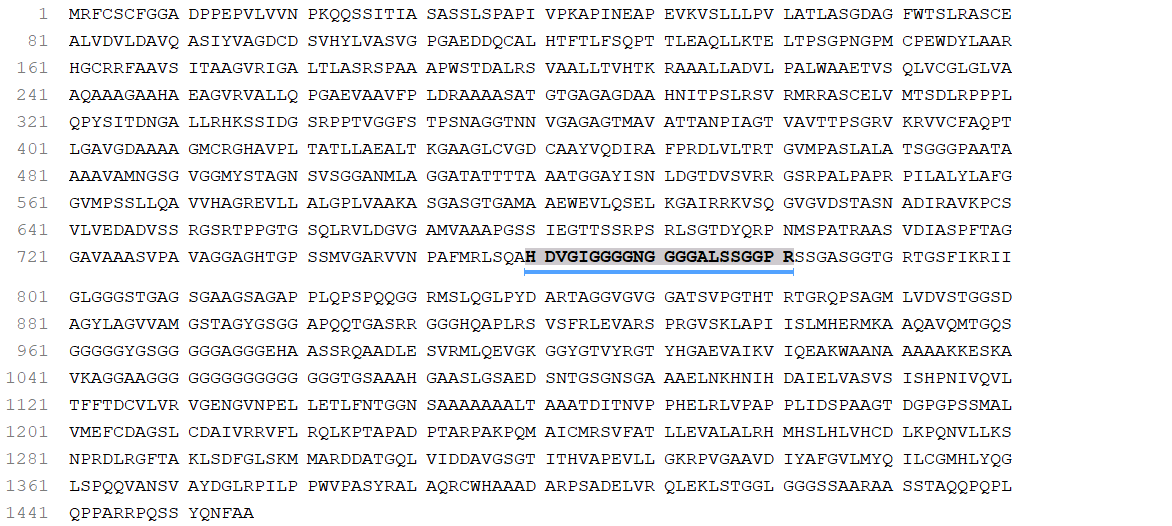

### cov_5748.png

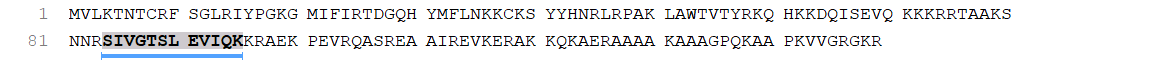

### cov_5749.png

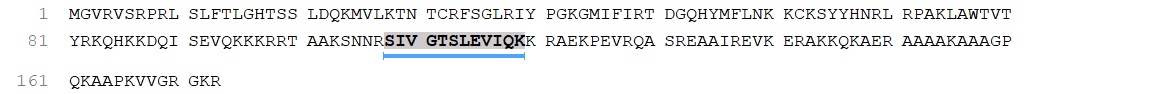

### cov_5750.png

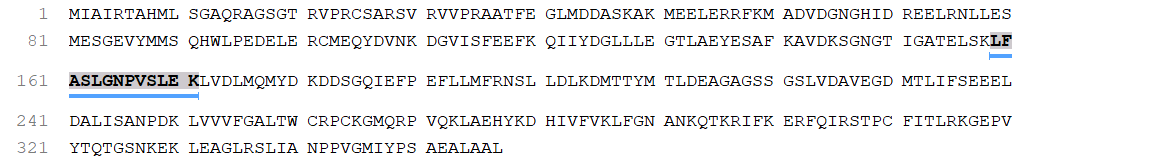

### cov_5751.png

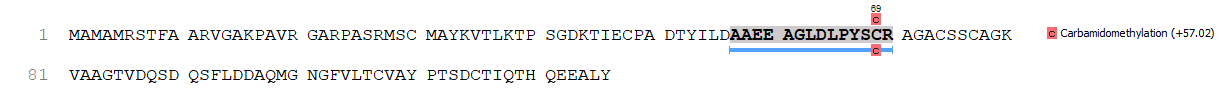

### cov_5753.png

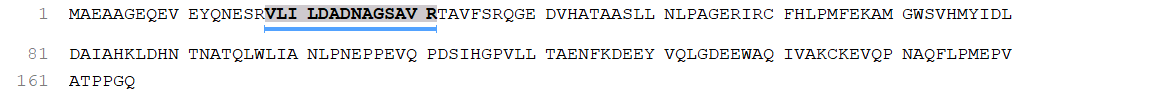

### cov_5754.png

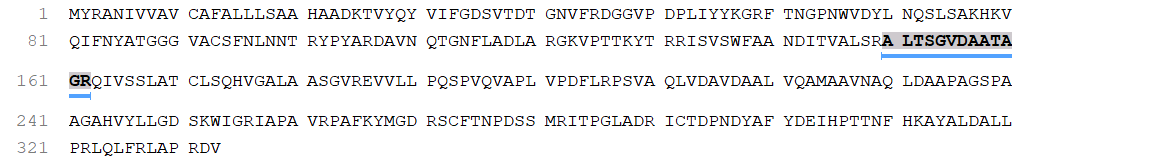

### cov_5757.png

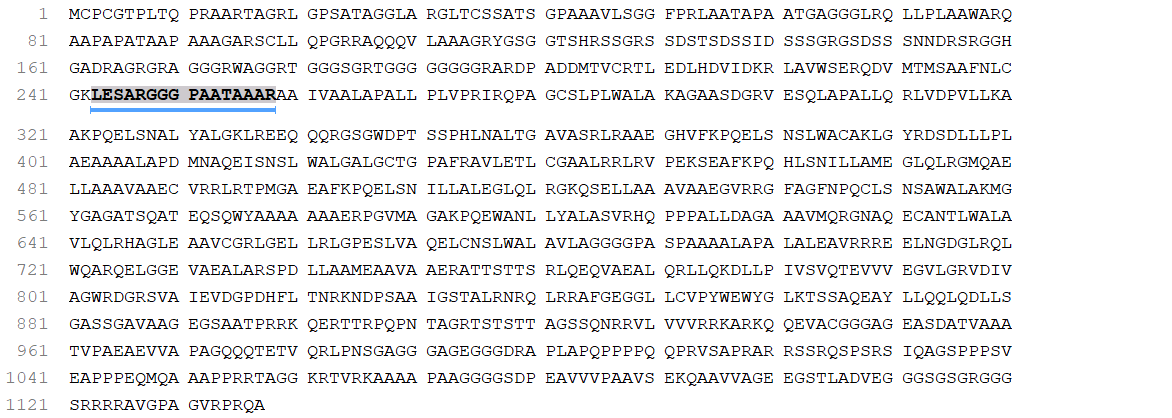

### DenovoFDRCurveFigure725967651367319998.png

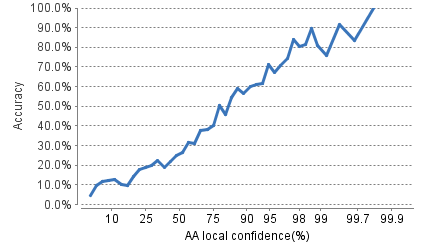

### ErrorCalibratedHistogram4295230005781691970.png

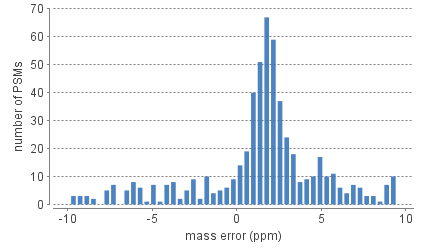

### ErrorPlotFigure3812743119609832443.png

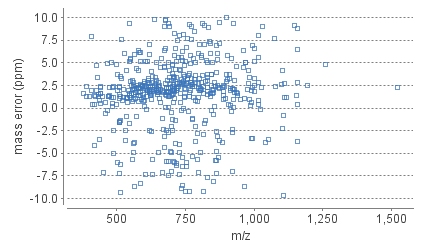

### FDRFigure4854596623094368.png

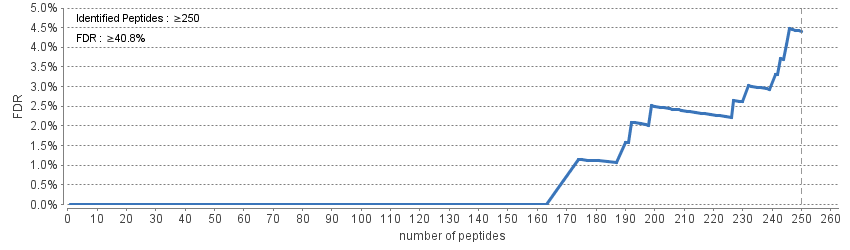

### FeatureIntensityDistributionHistogram2354080603349882113.png

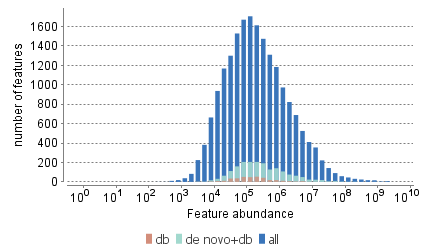

### FeatureMzHistogram1974660684123570834.png

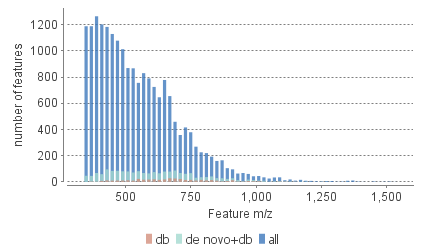

### FeatureRtHistogram9138977435263175233.png

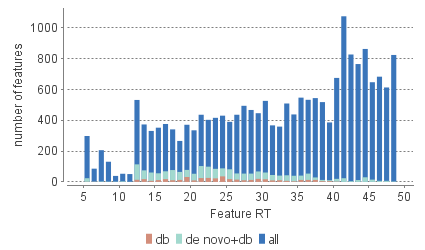

### IMG_0002.JPG

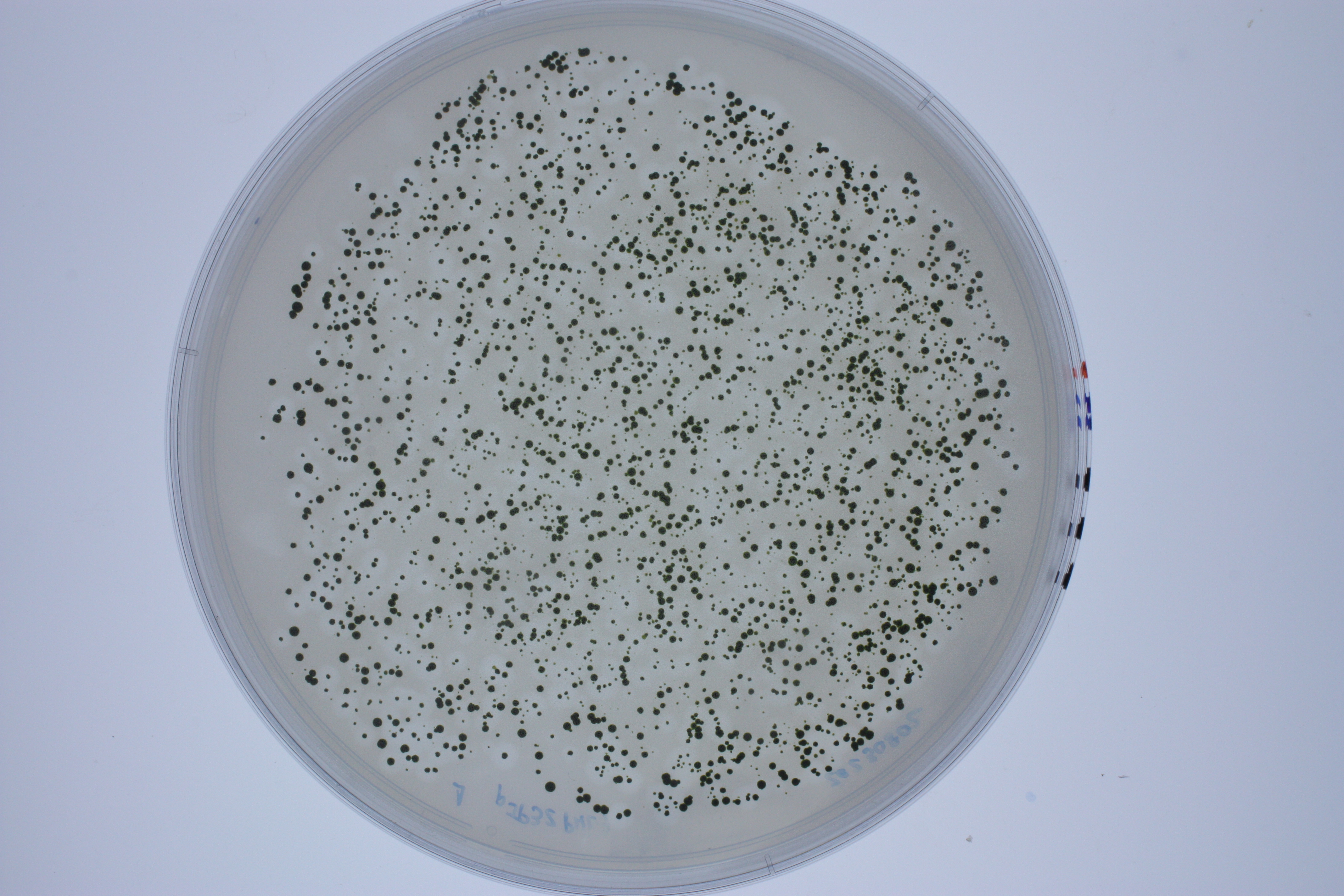

### IMG_0002_edit.png

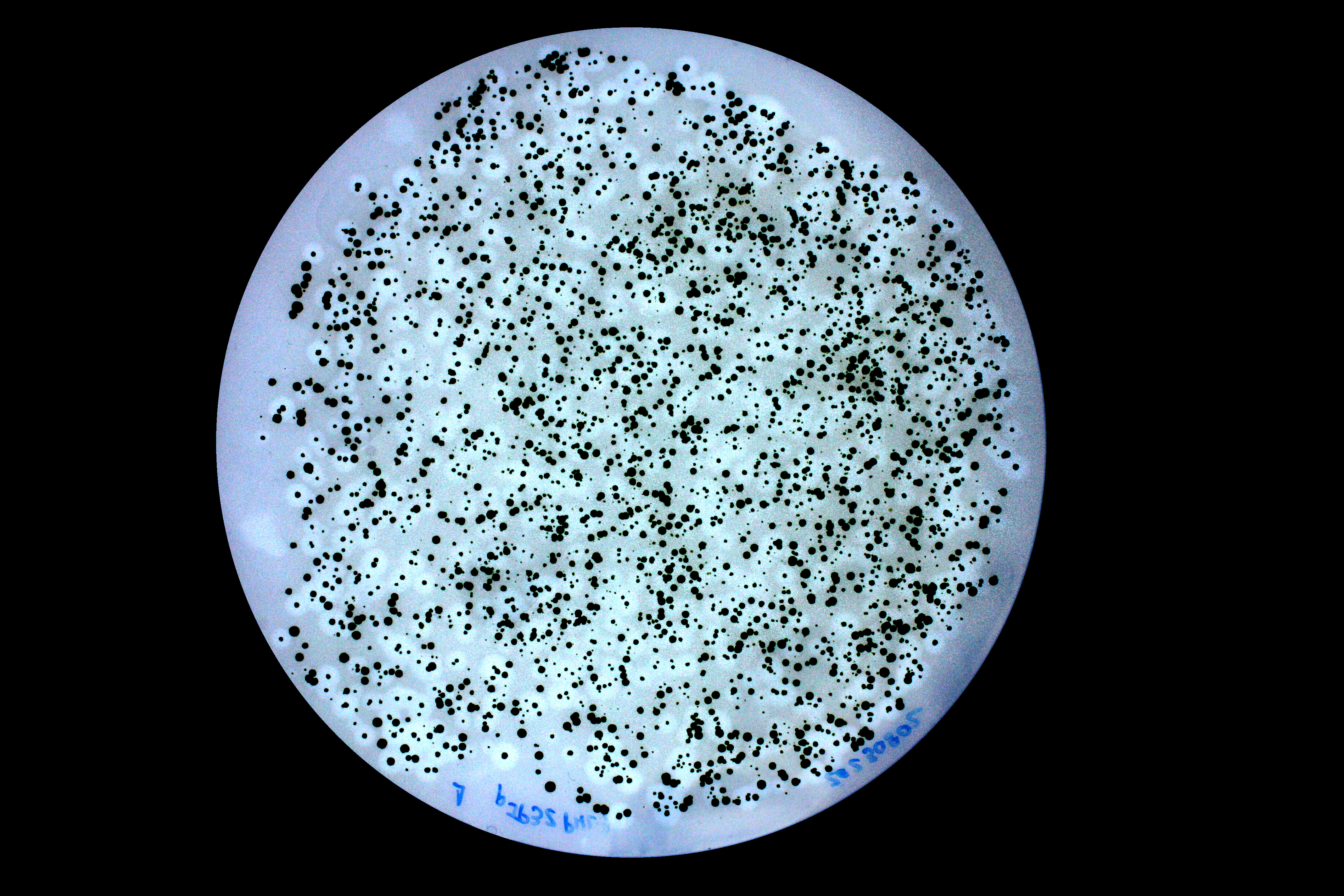

### IMG_0004.JPG

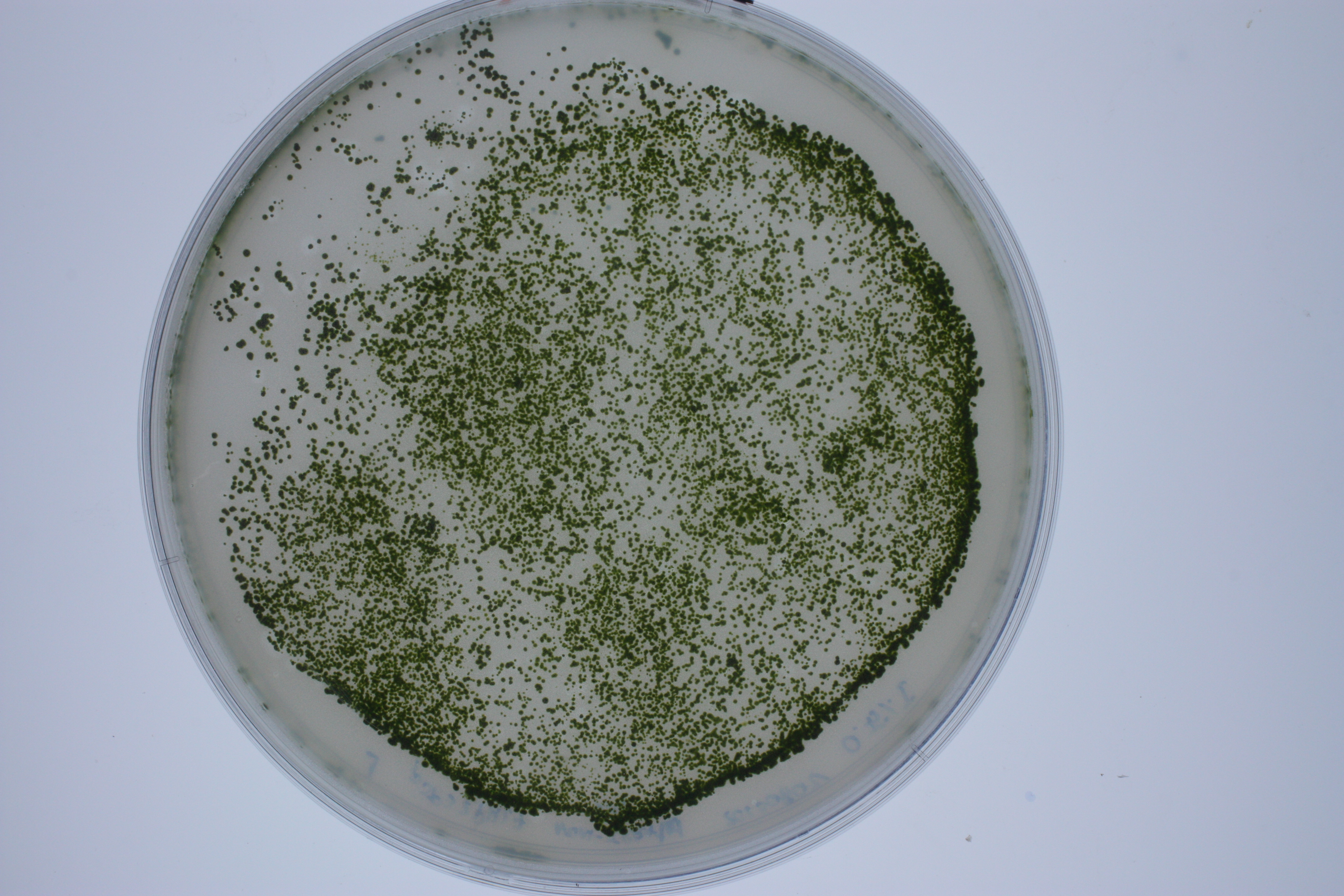

### IMG_0006.JPG

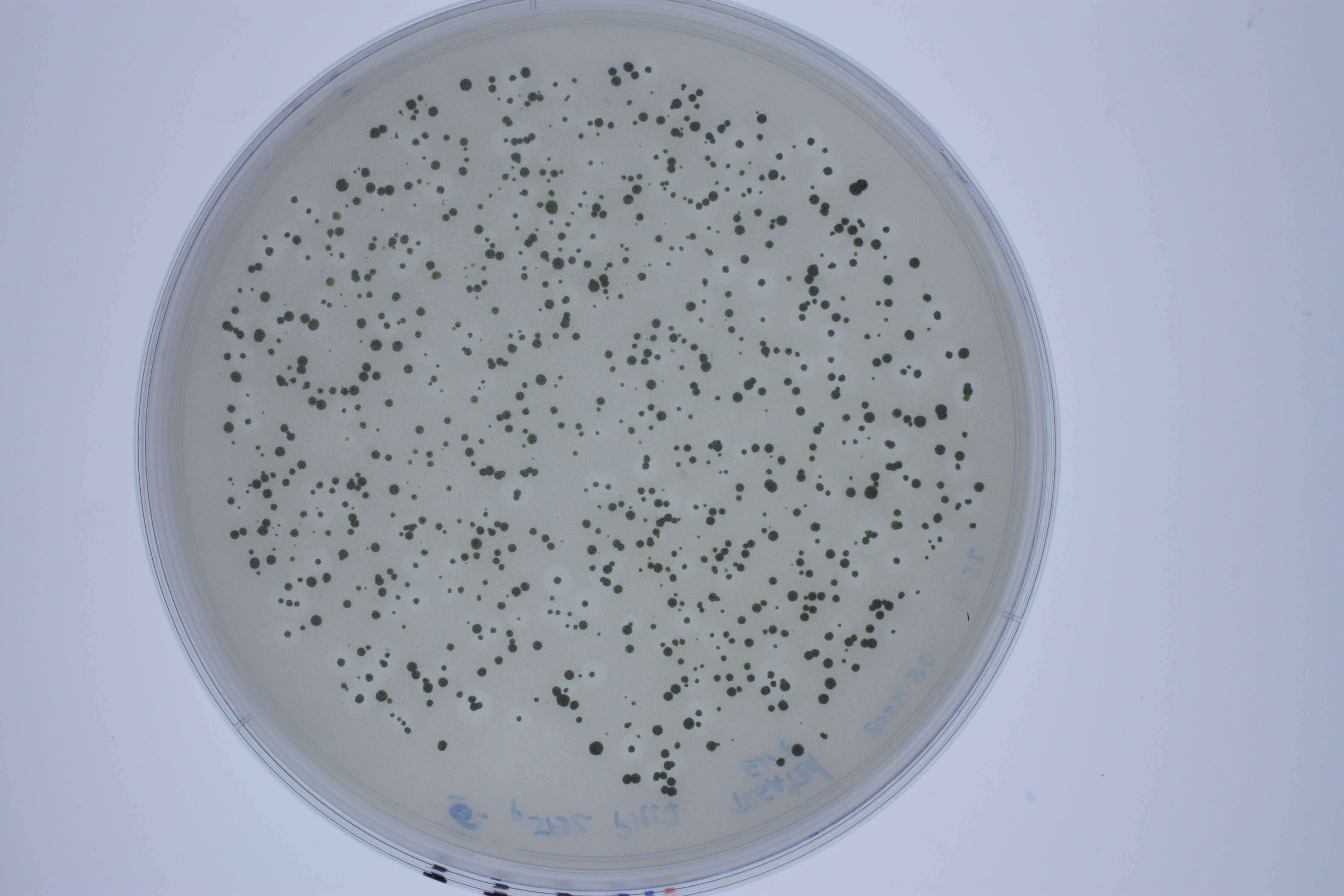

### IMG_0006_edit.png

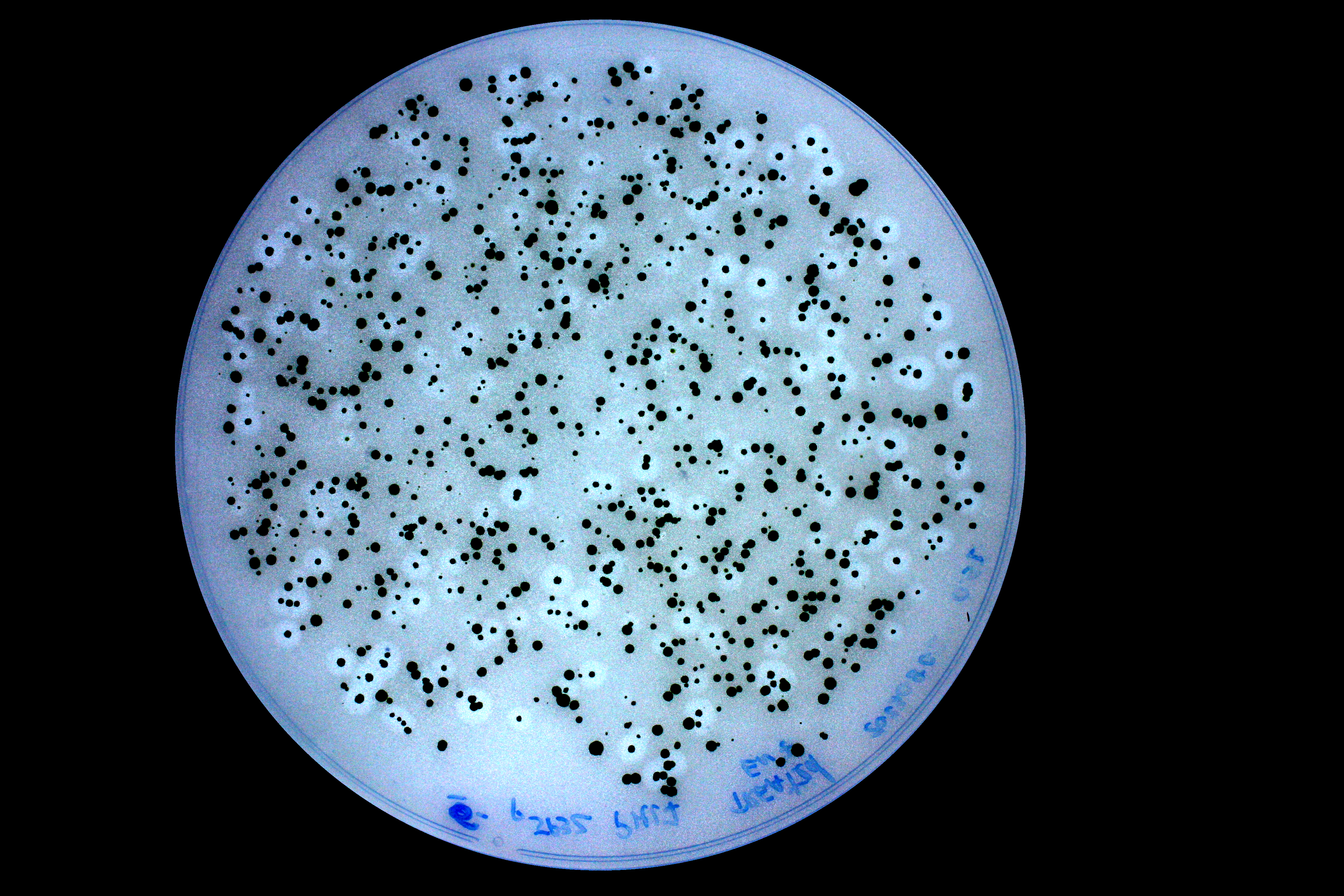

### IMG_0008.JPG

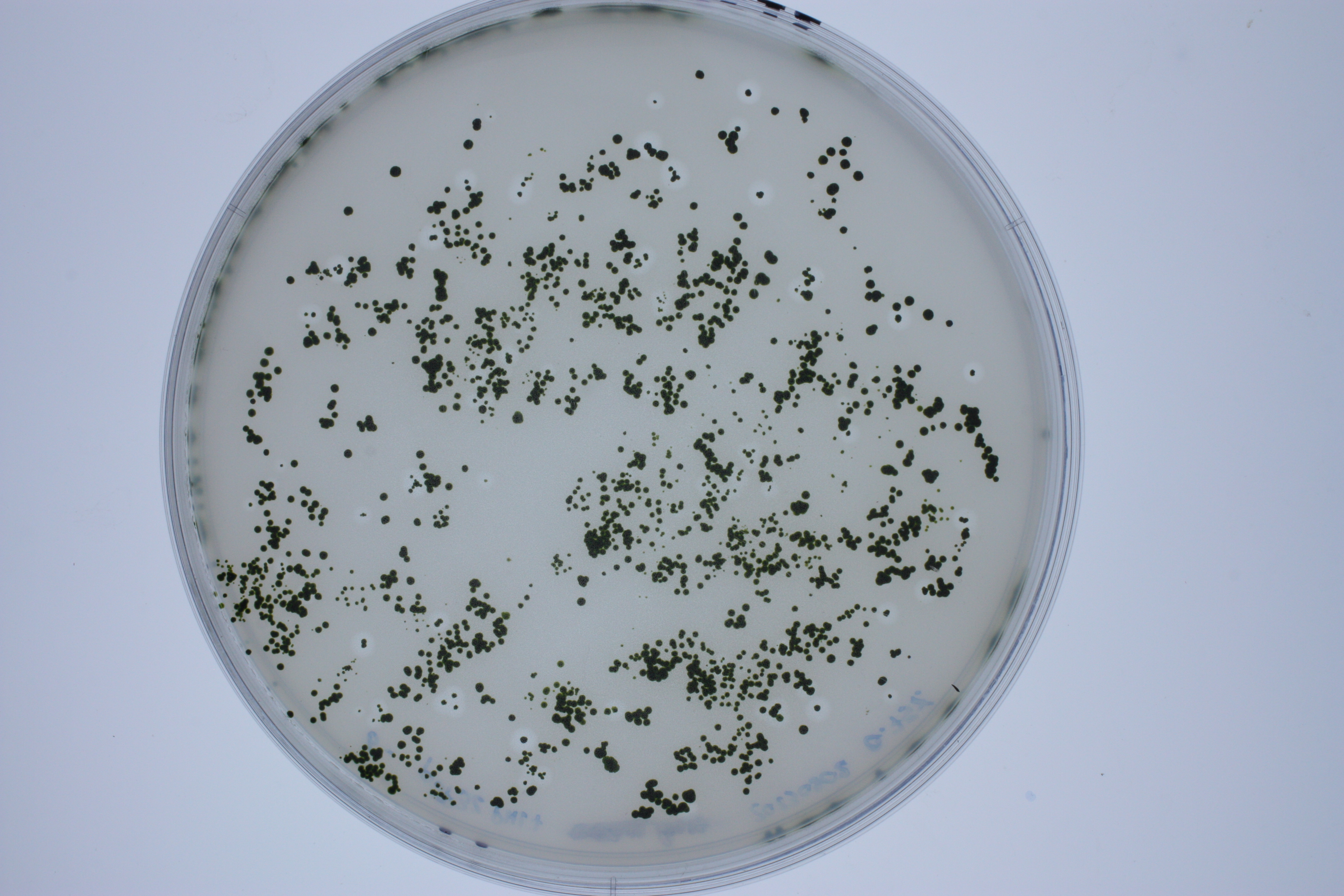

### IMG_0010.JPG

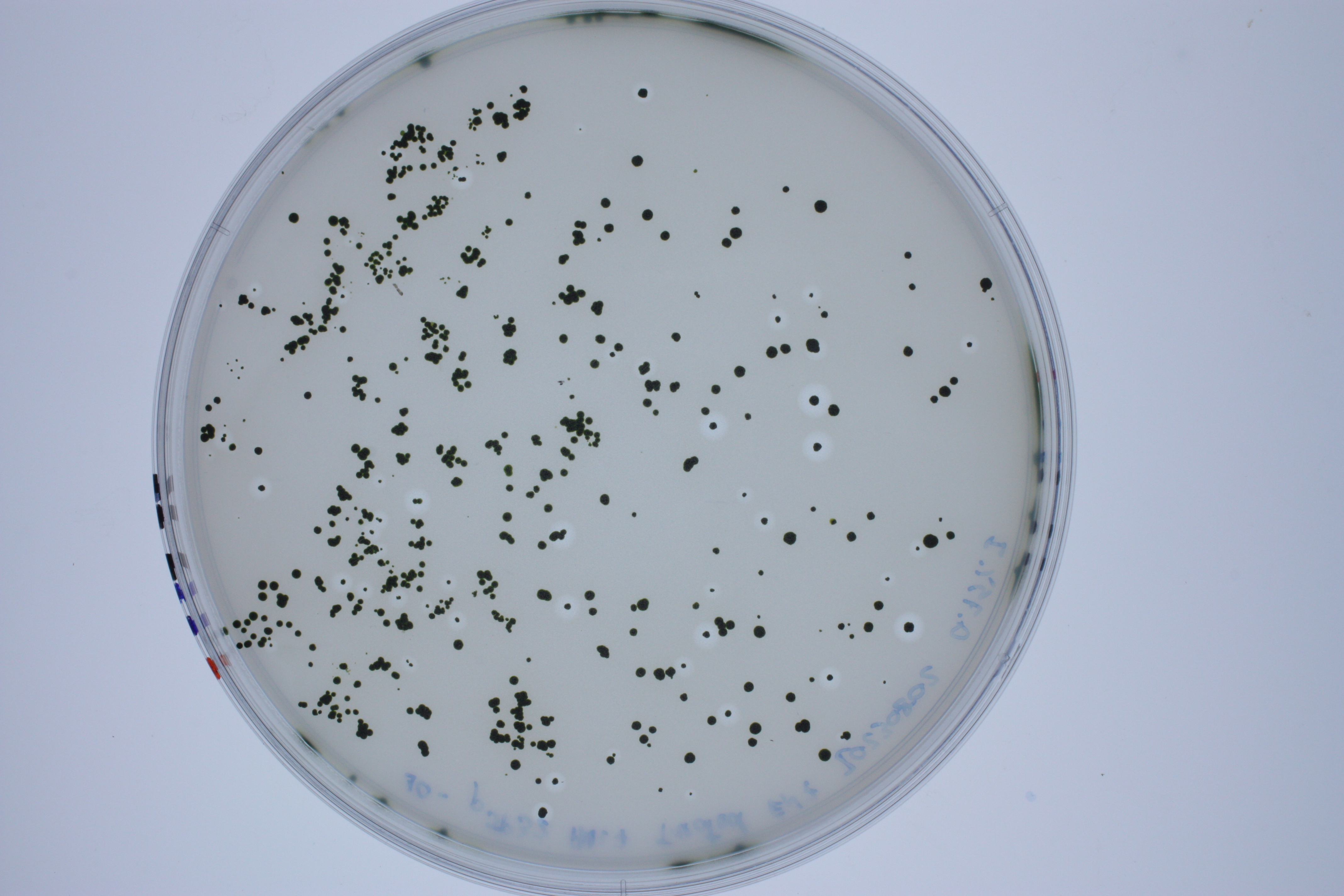

### IMG_0012.JPG

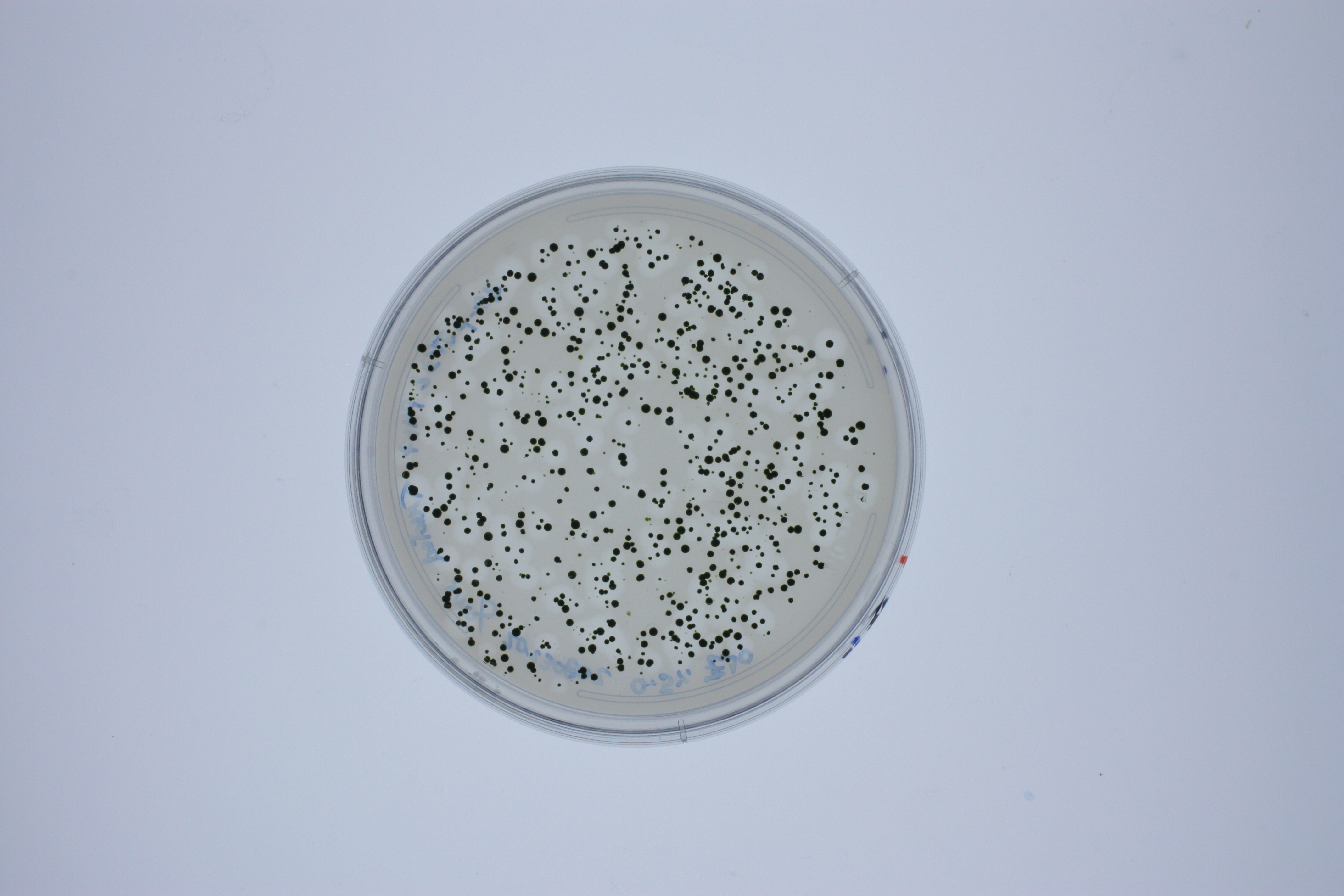

### IMG_0012_edit.png

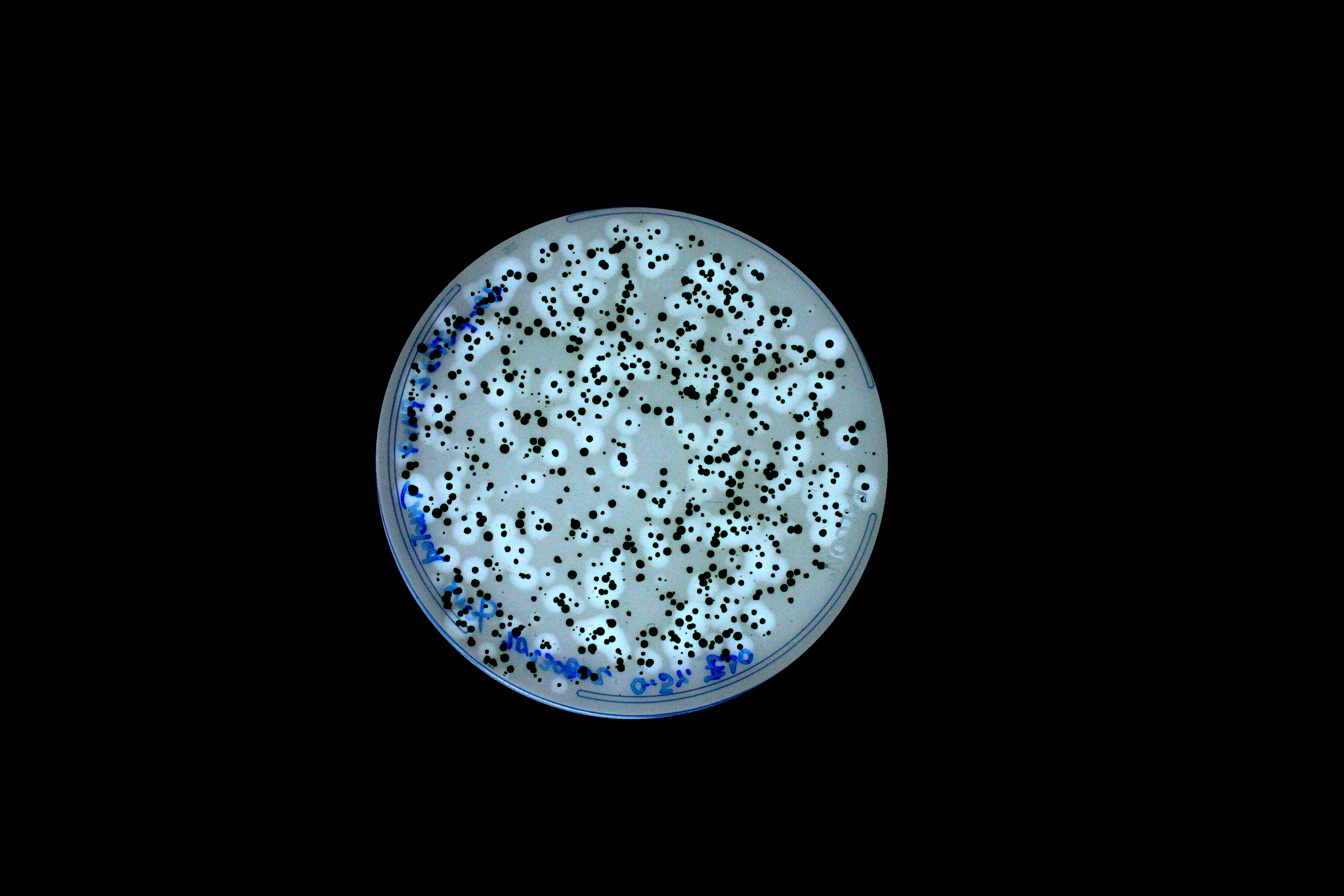

### IMG_0014.JPG

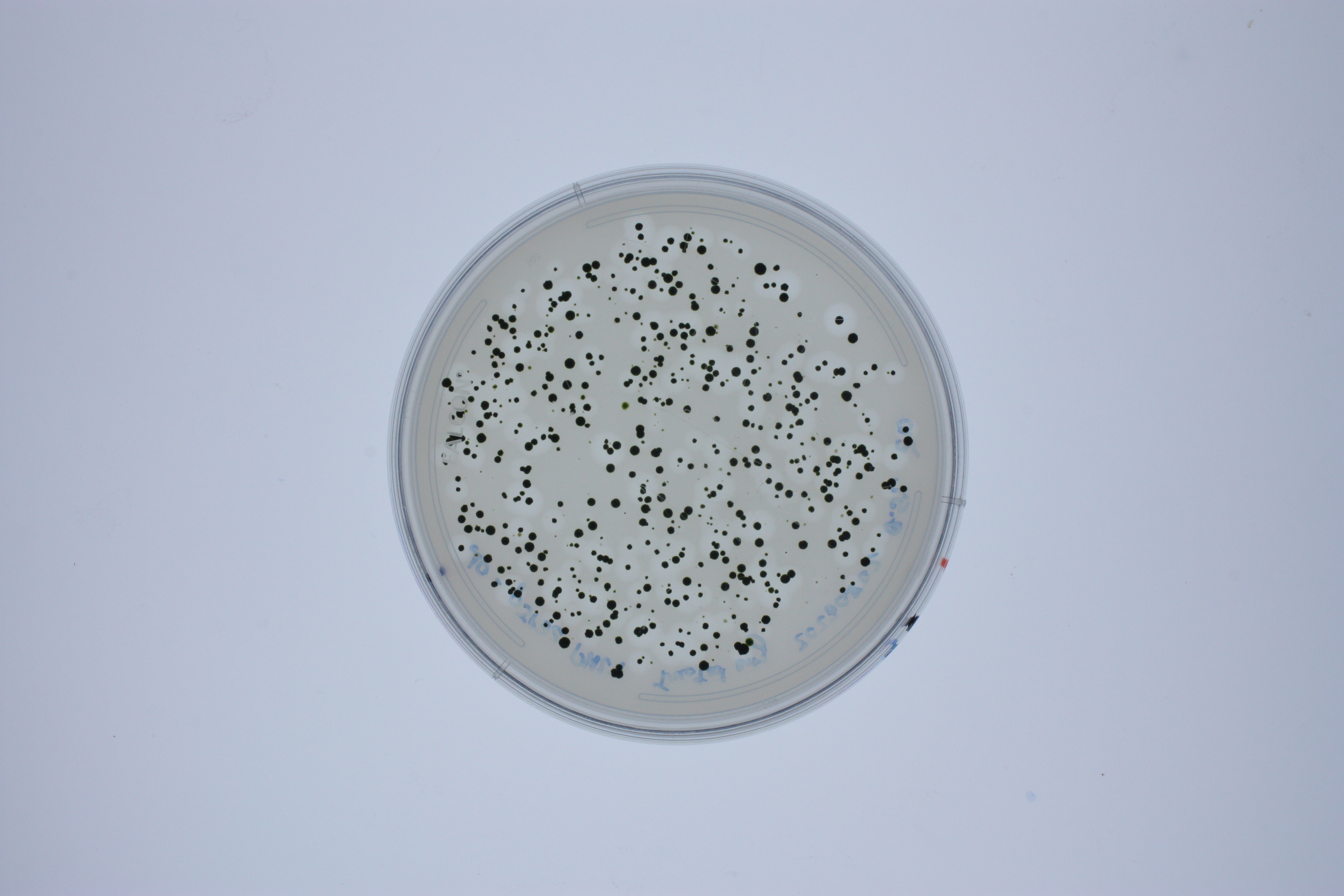

### IMG_0014_edit.png

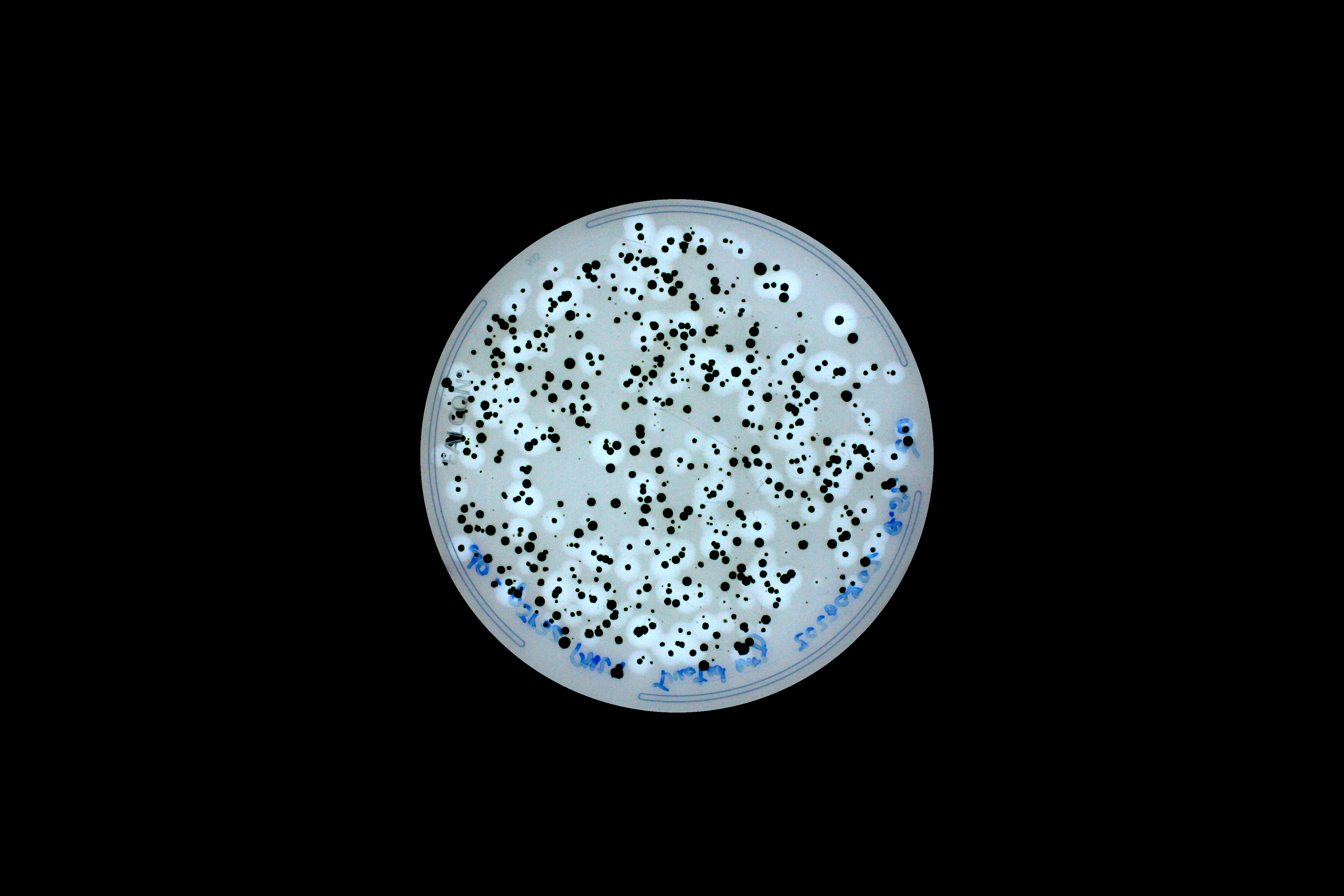

### ScoreHistogram7891267701088096522.png

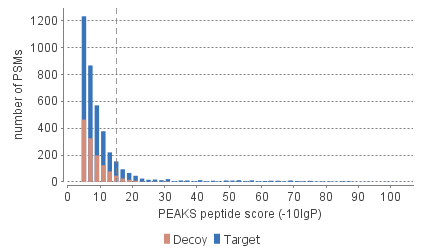

### ScorePlotFigure7242268655030745994.png

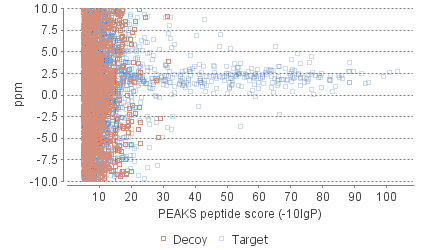
