## Supplementary figures and images for "Efficient secretion of a plastic degrading enzyme from the green algae *Chlamydomonas reinhardtii*"

### cov_1.png

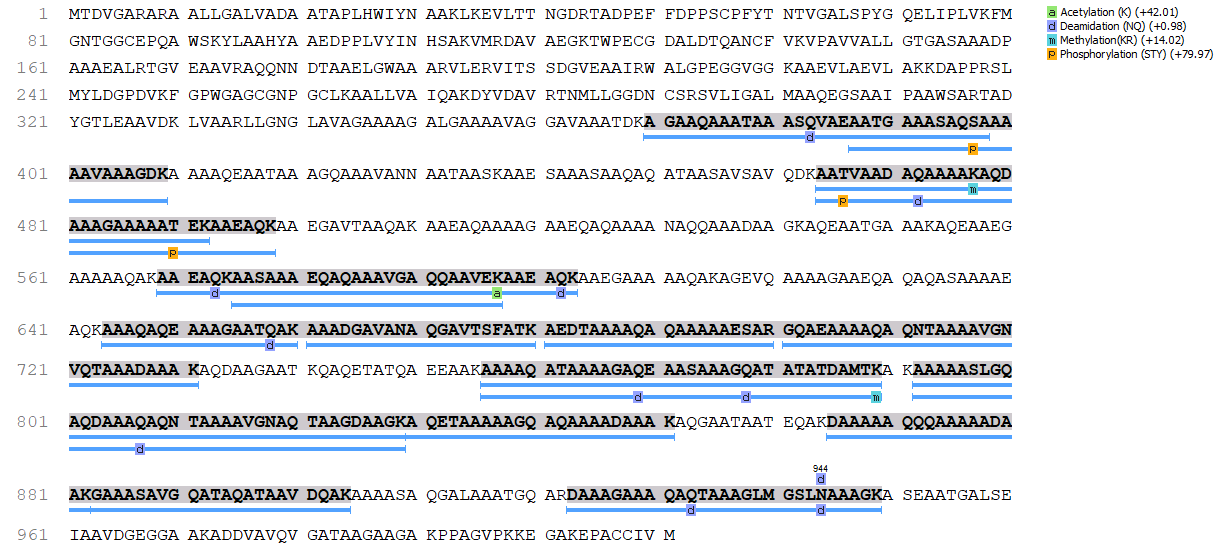

### cov_1.png

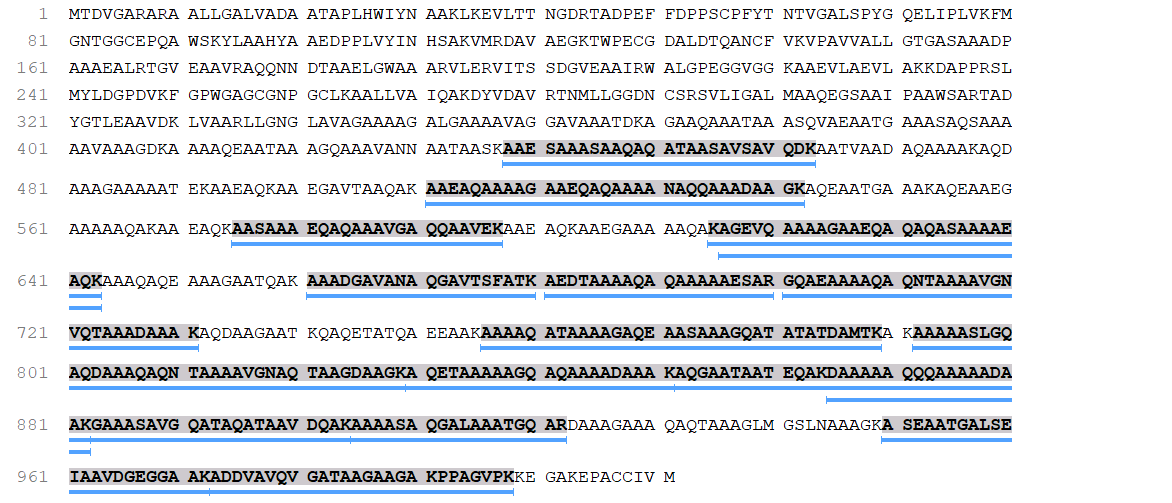

### cov_2.png

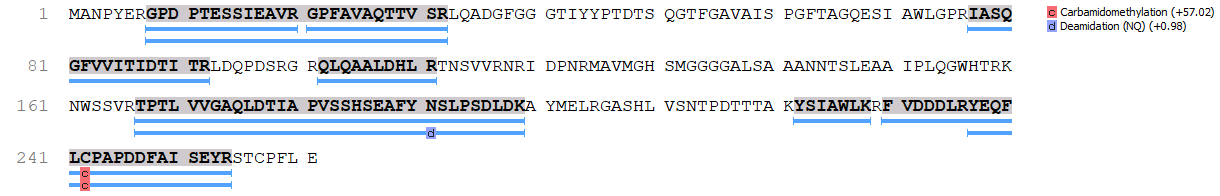

### cov_2.png

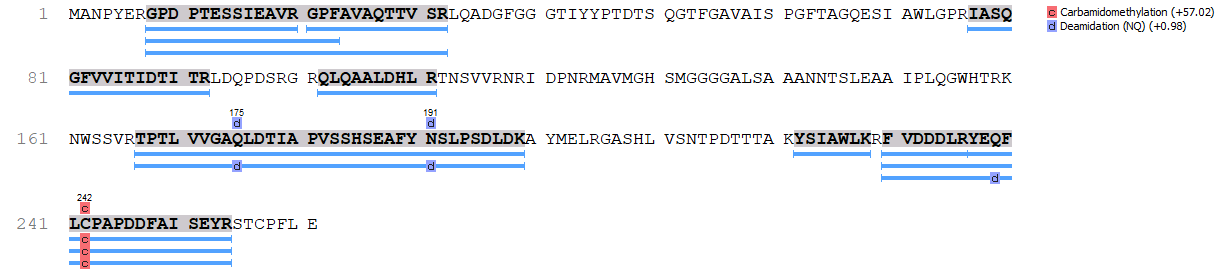

### cov_3.png

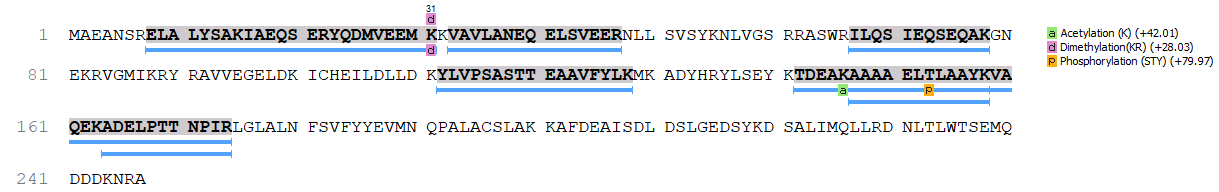

### cov_3.png

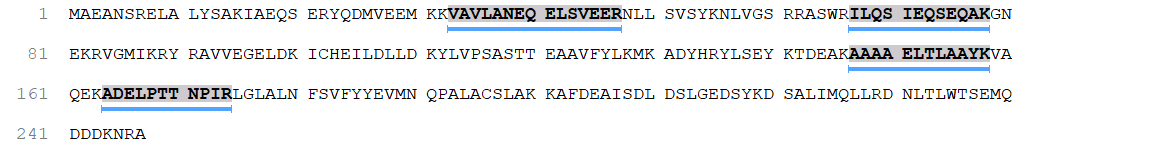

### cov_4.png

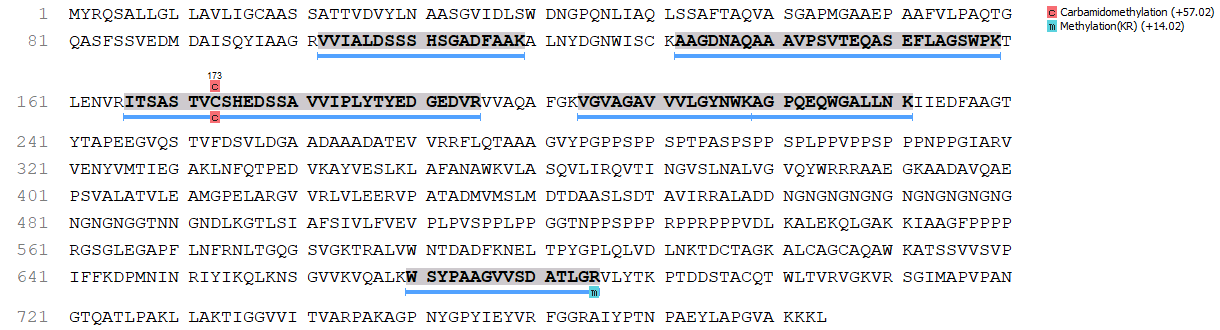

### cov_4.png

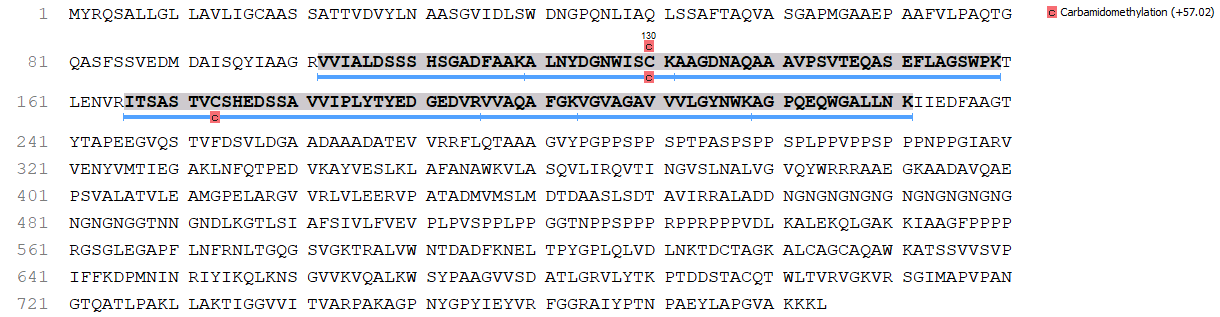

### cov_5.png

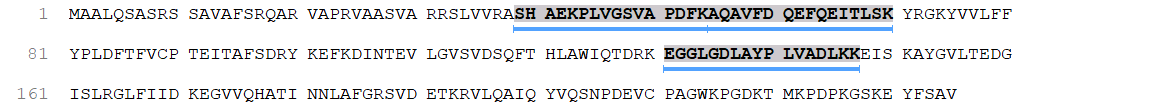

### cov_5.png

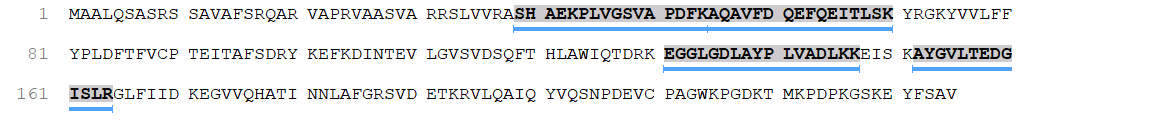

### cov_6.png

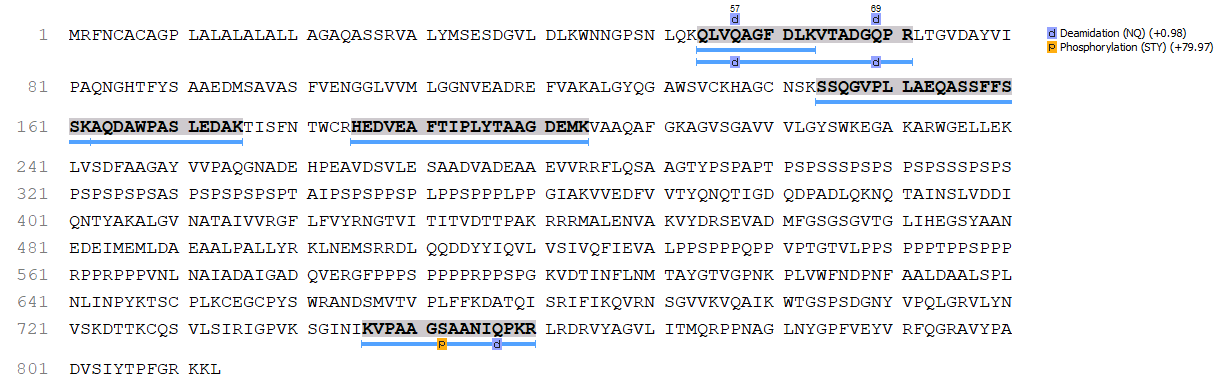

### cov_7.png

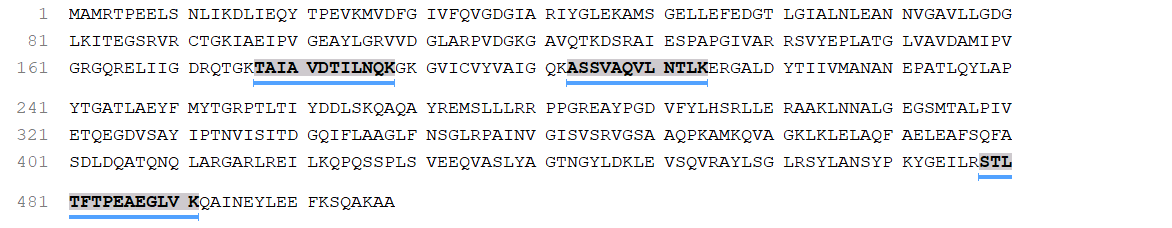

### cov_8.png

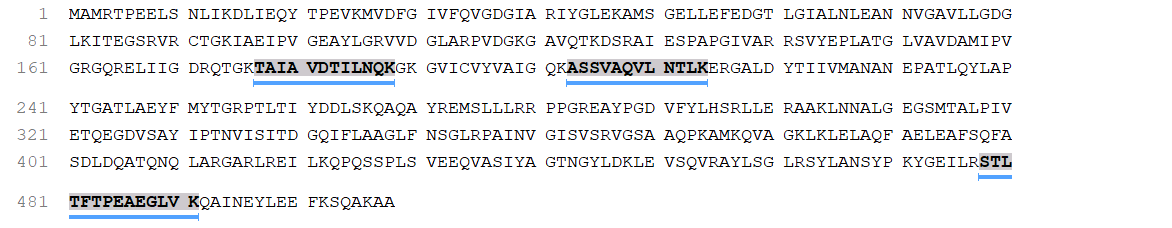

### cov_9.png

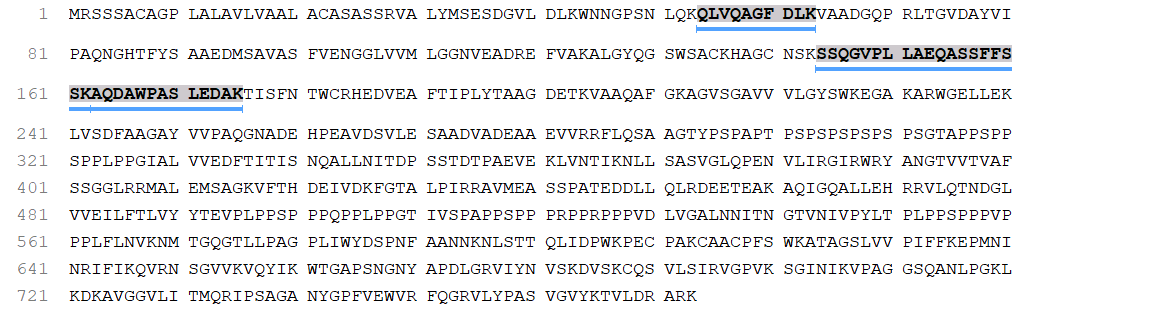

### cov_10.png

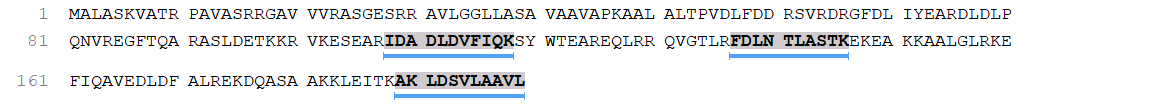

### cov_11.png

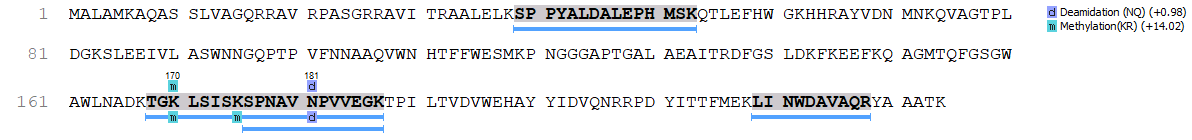

### cov_11.png

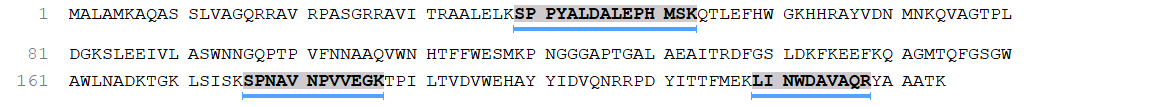

### cov_12.png

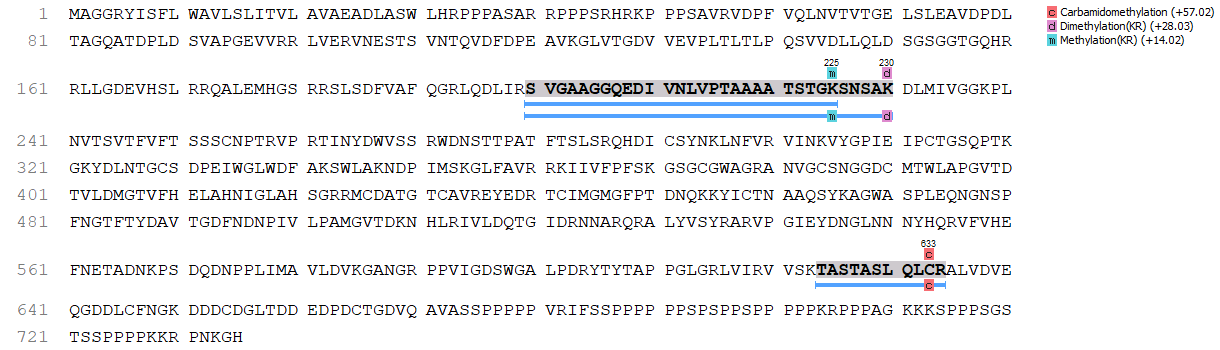

### cov_12.png

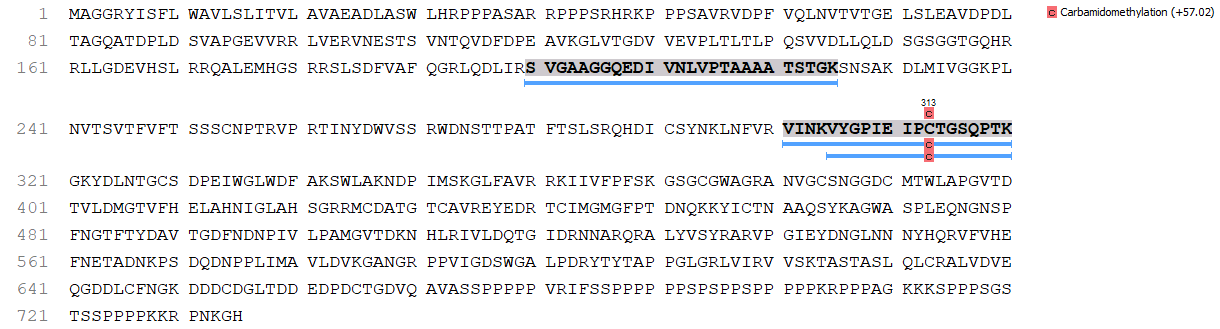

### cov_13.png

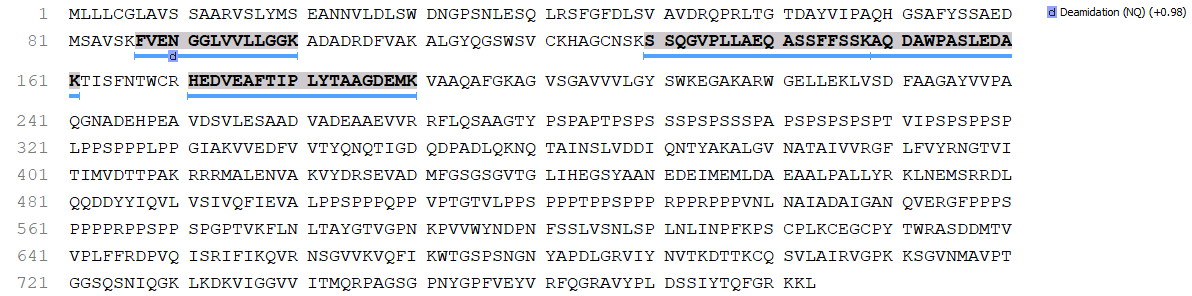

### cov_13.png

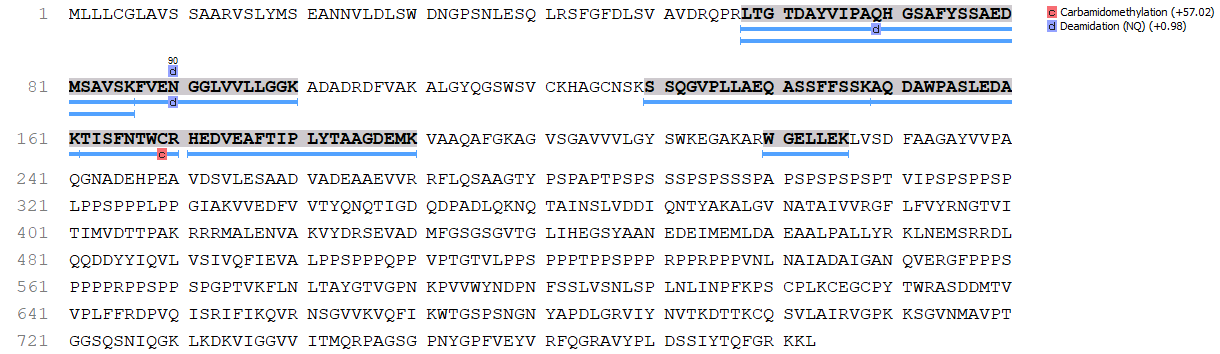

### cov_14.png

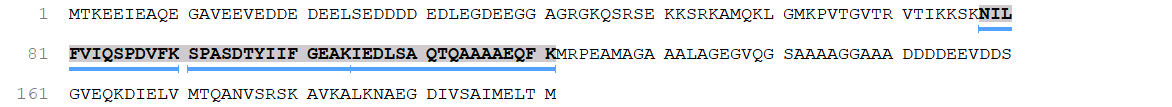

### cov_14.png

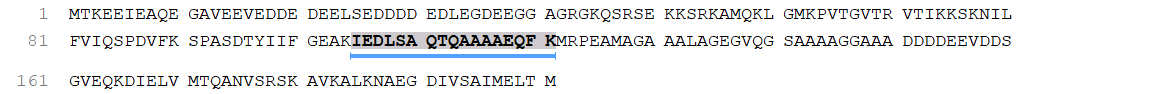

### cov_15.png

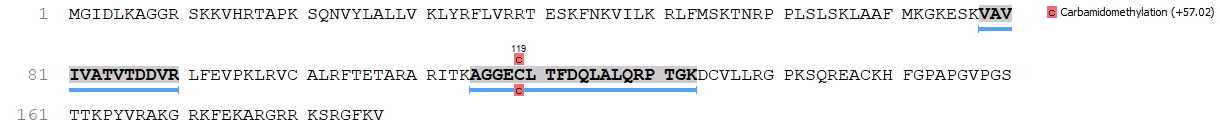

### cov_15.png

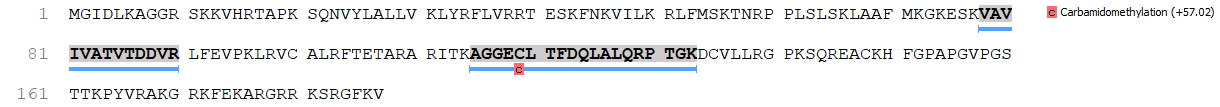

### cov_16.png

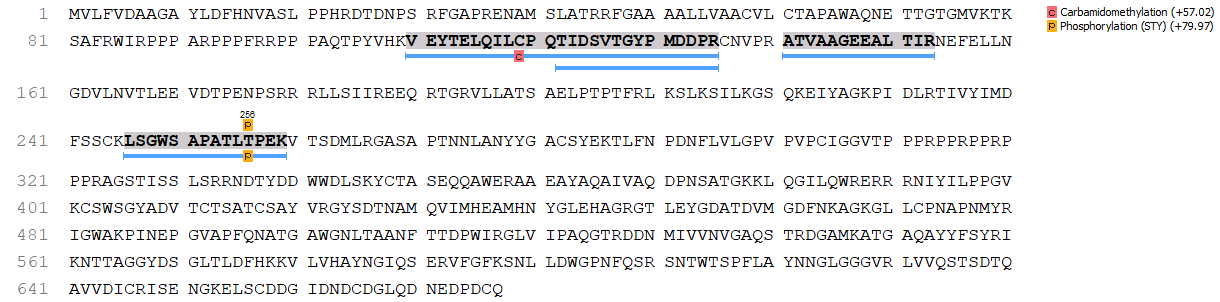

### cov_16.png

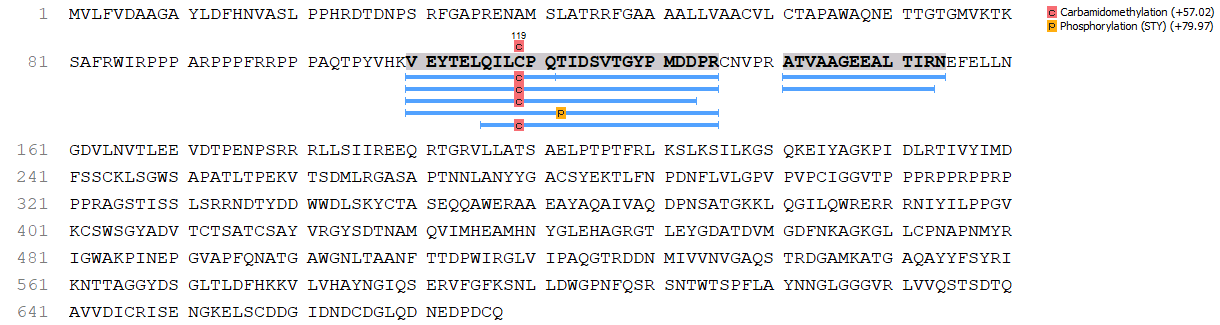

### cov_17.png

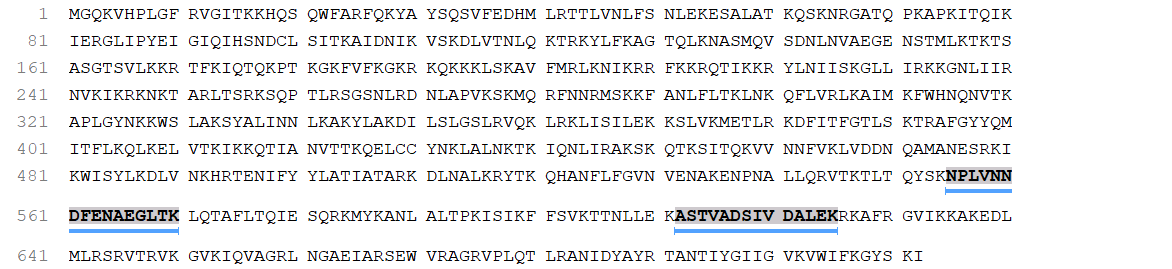

### cov_17.png

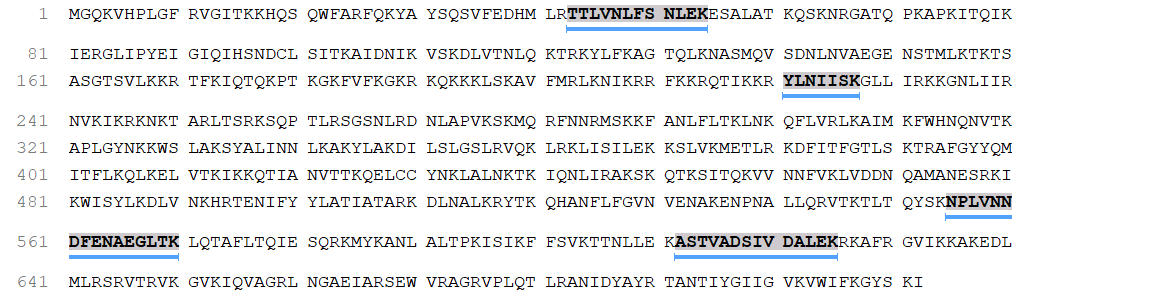

### cov_18.png

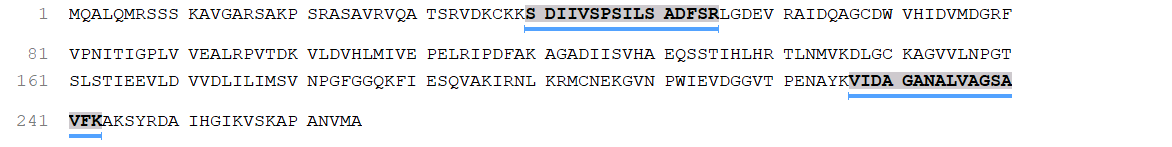

### cov_19.png

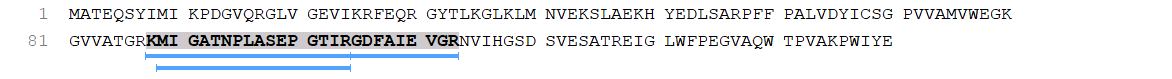

### cov_20.png

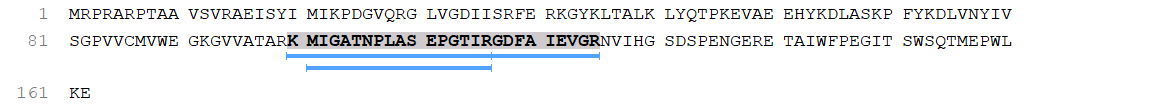

### cov_21.png

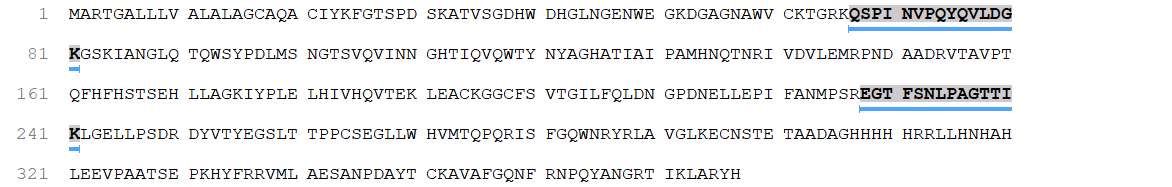

### cov_22.png

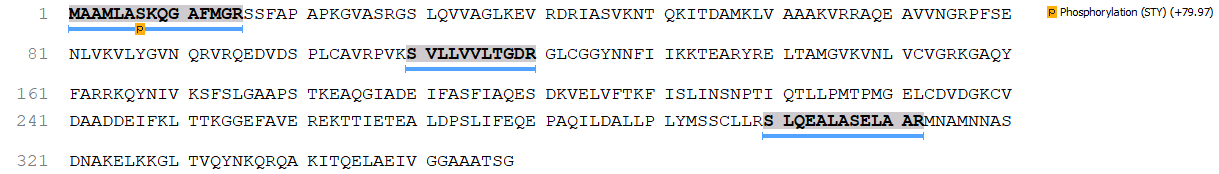

### cov_22.png

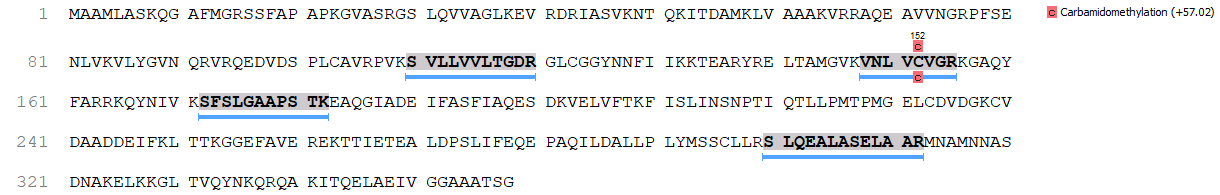

### cov_23.png

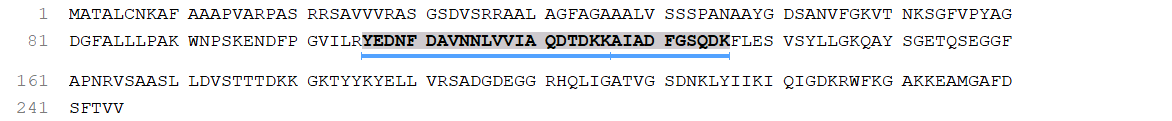

### cov_23.png

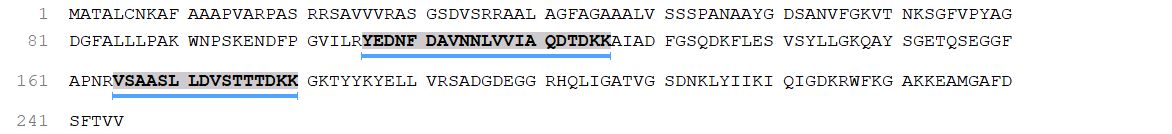

### cov_24.png

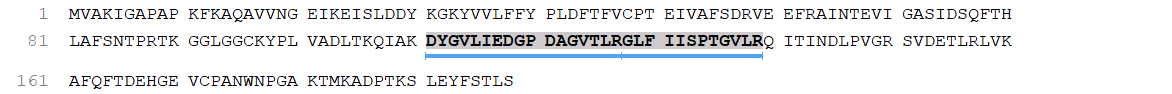

### cov_26.png

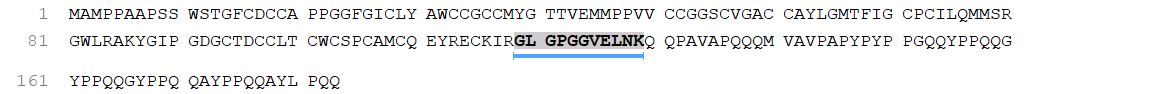

### cov_27.png

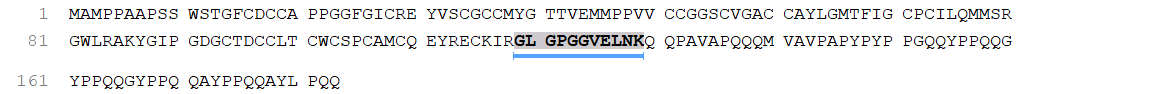

### cov_28.png

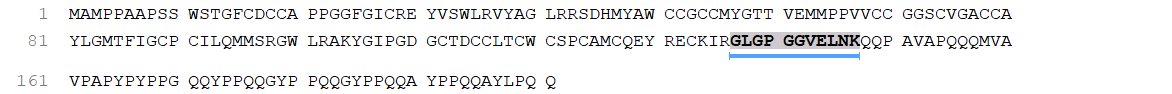

### cov_29.png

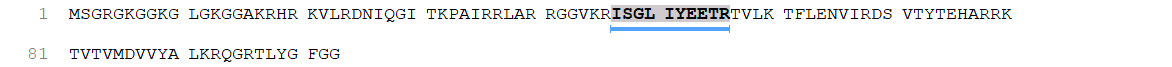

### cov_30.png

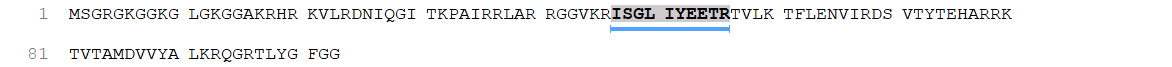

### cov_33.png

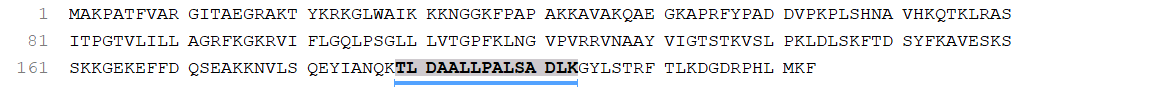

### cov_34.png

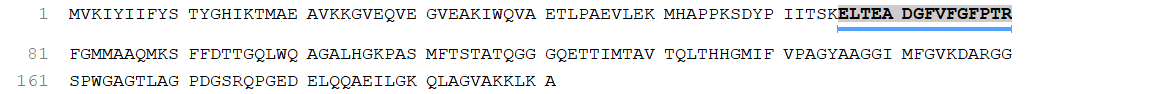

### cov_35.png

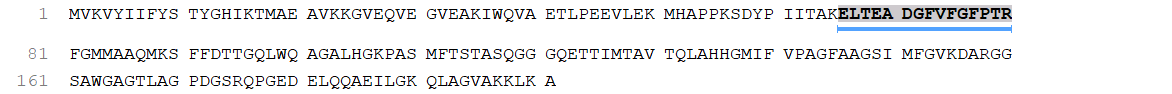

### cov_36.png

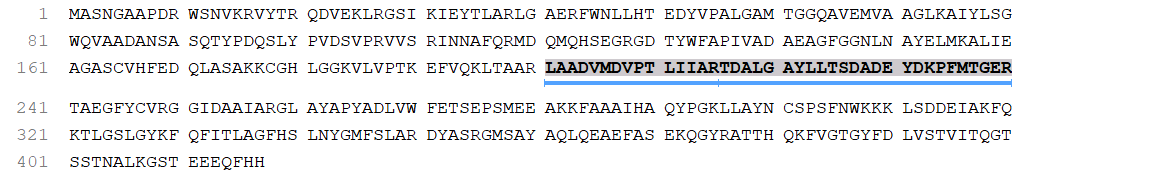

### cov_36.png

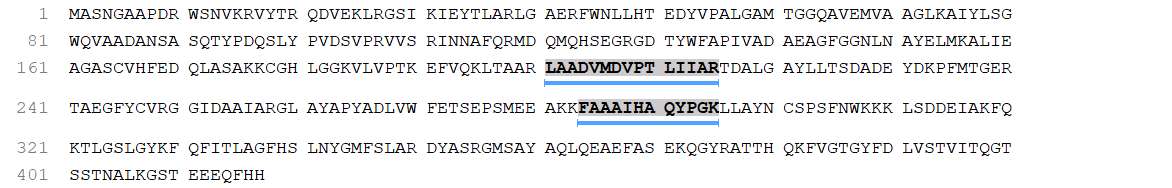

### cov_37.png

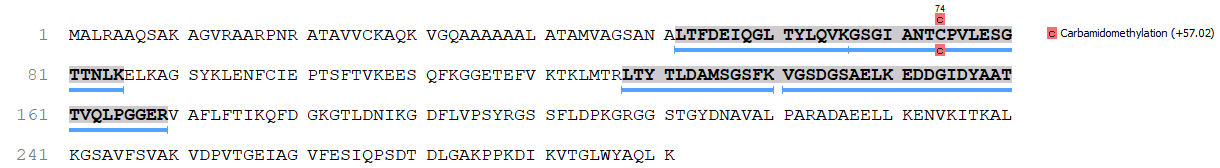

### cov_42.png

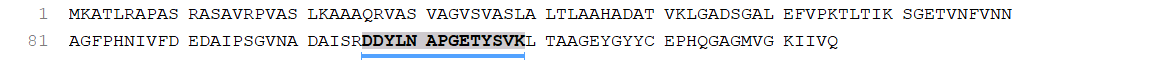

### cov_43.png

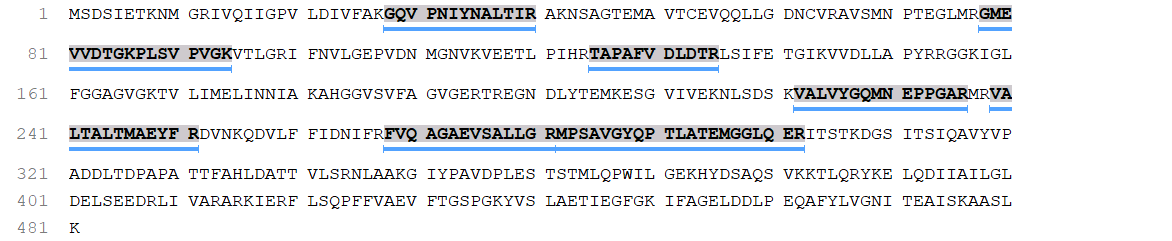

### cov_44.png

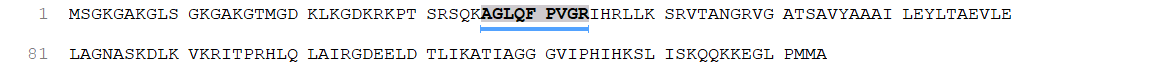

### cov_47.png

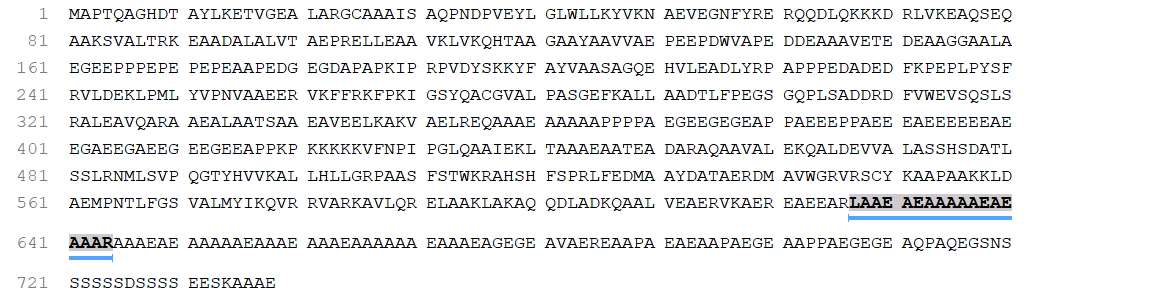

### cov_50.png

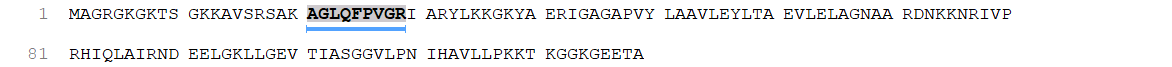

### cov_51.png

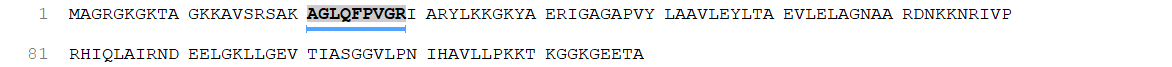

### cov_52.png

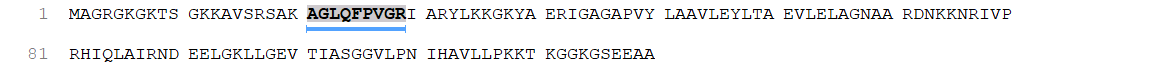

### cov_53.png

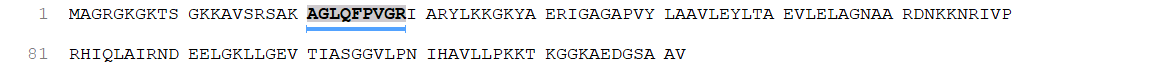

### cov_54.png

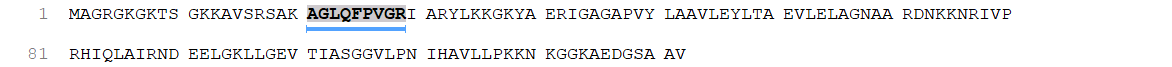

### cov_55.png

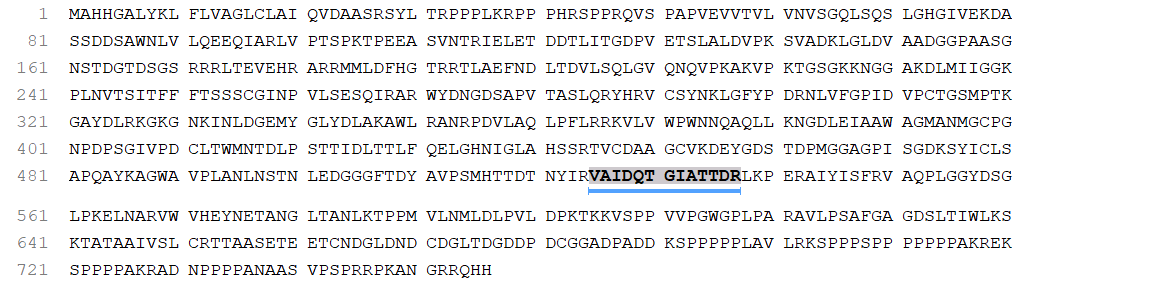

### cov_55.png

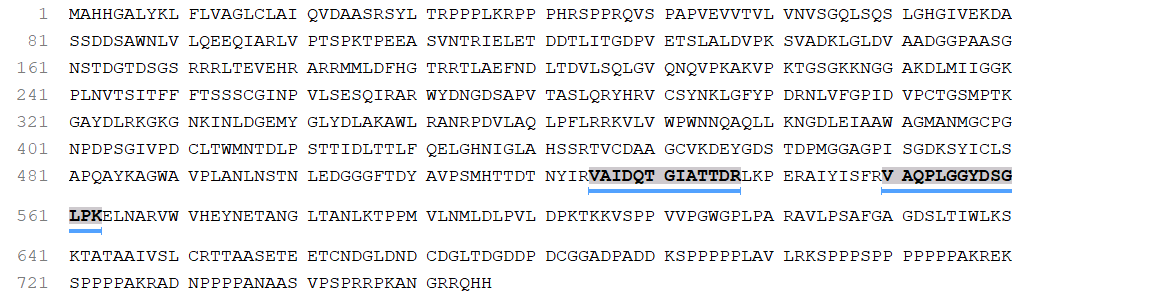

### cov_212.png

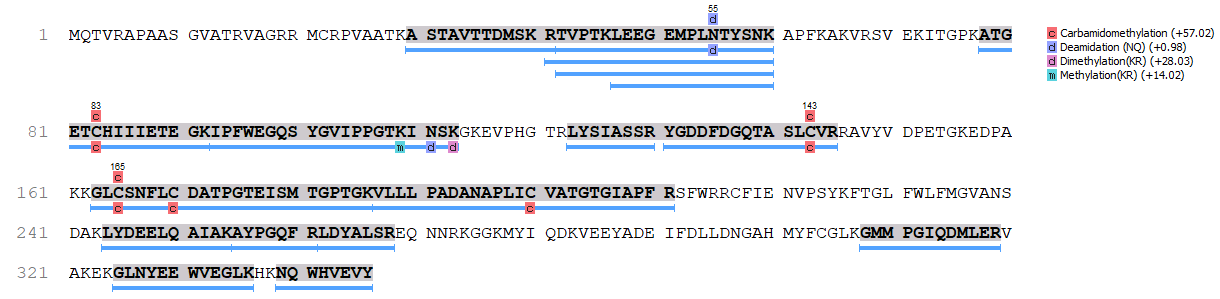

### cov_223.png

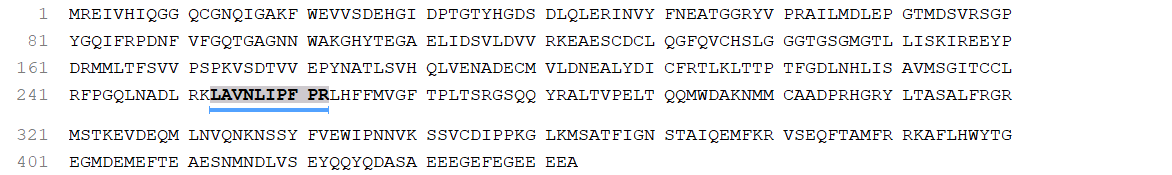

### cov_294.png

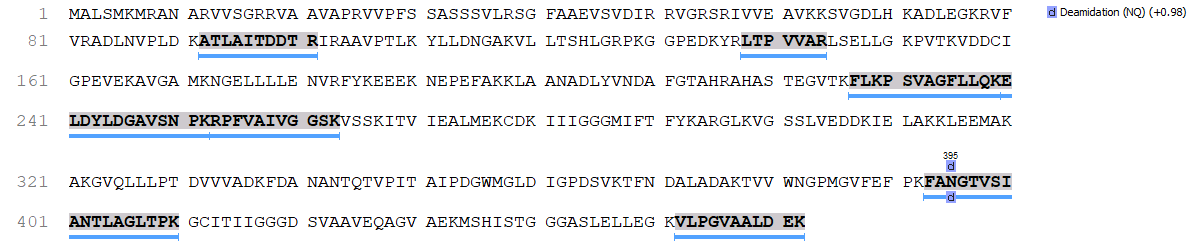

### cov_511.png

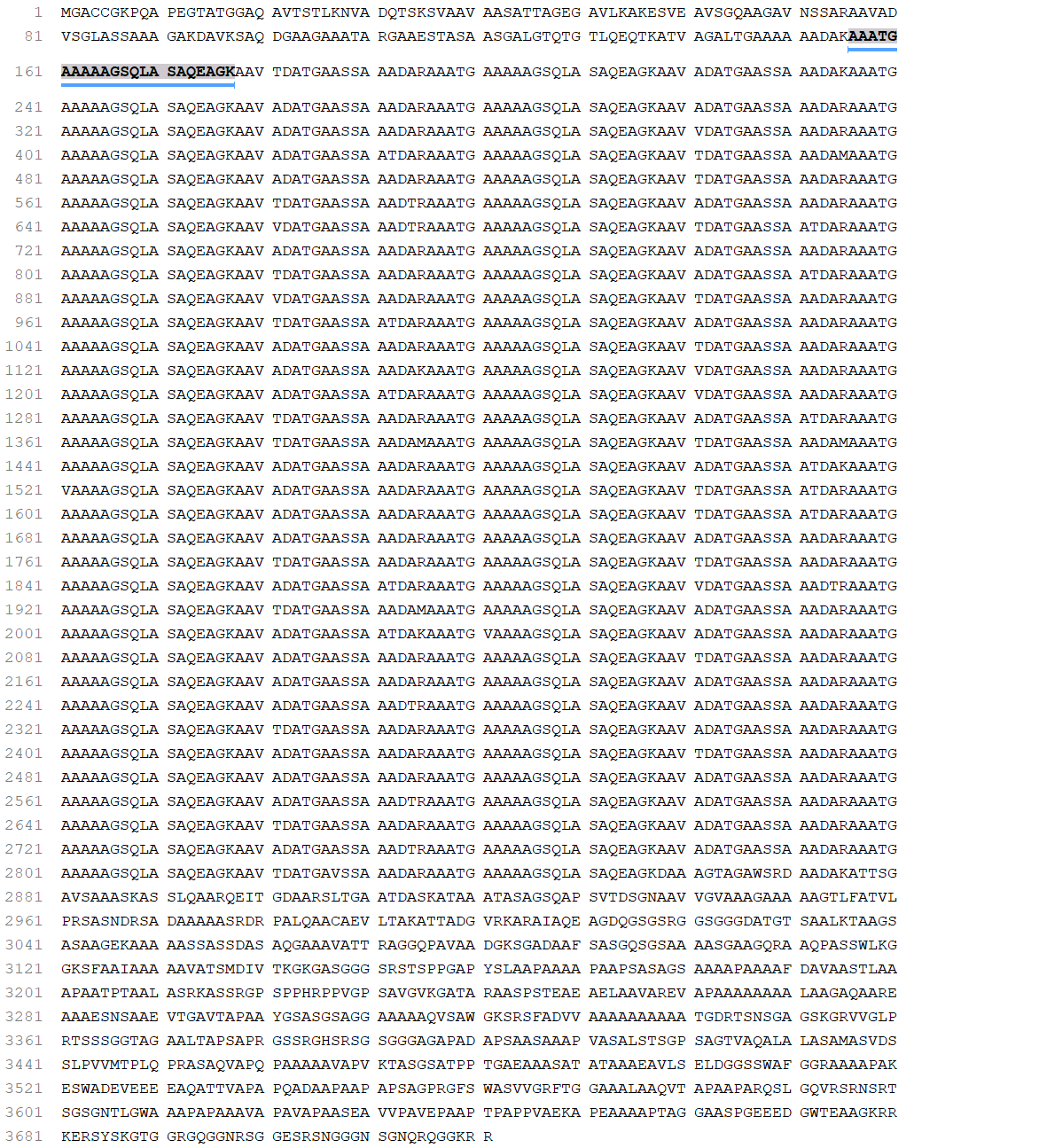

### cov_564.png

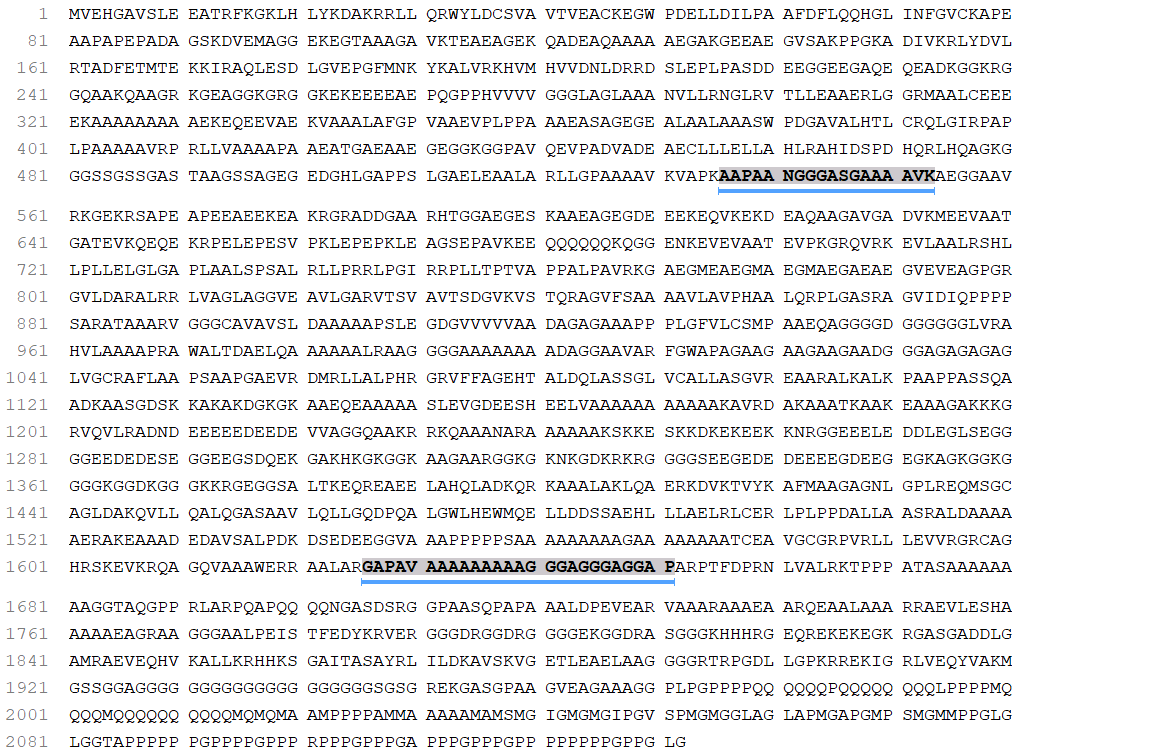

### cov_1031.png

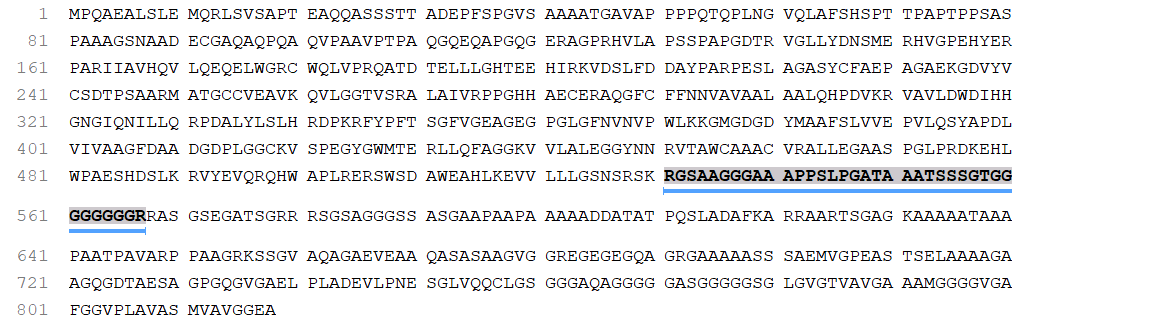

### cov_5722.png

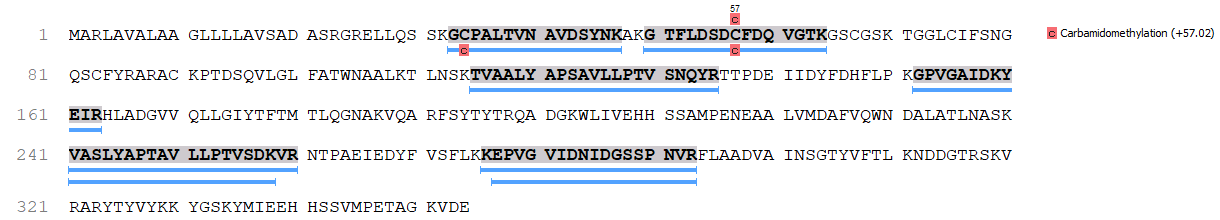

### cov_5723.png

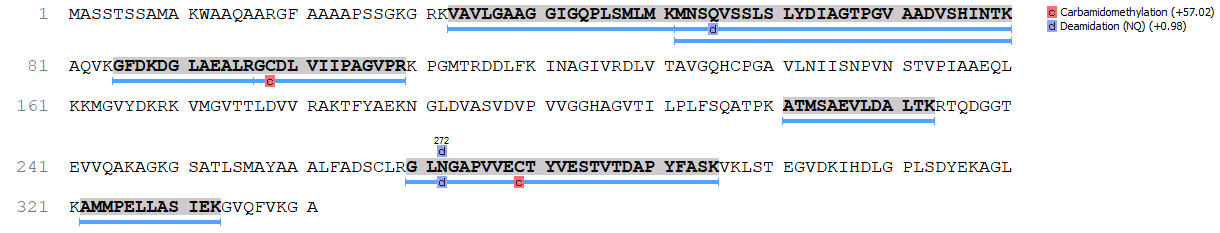

### cov_5725.png

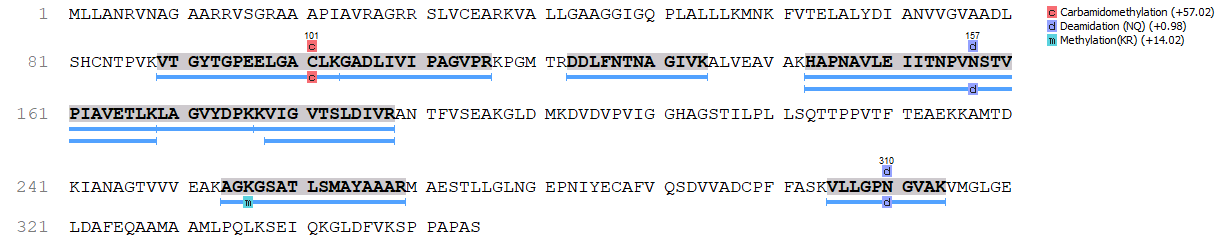

### cov_5726.png

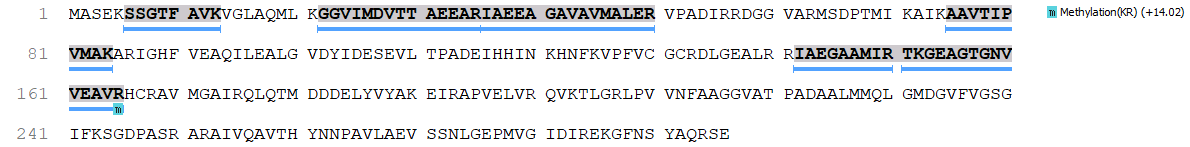

### cov_5727.png

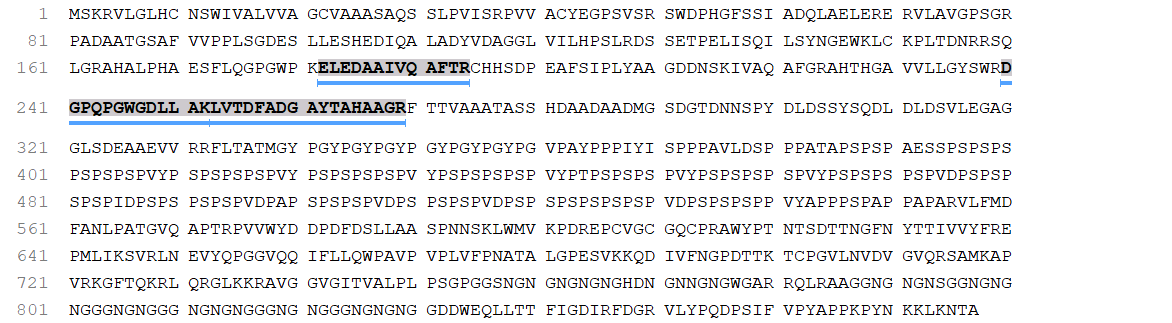

### cov_5728.png

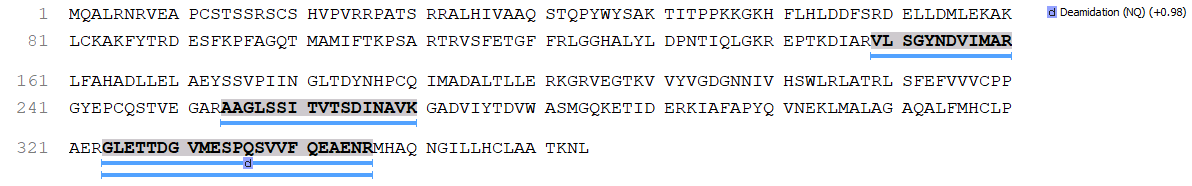

### cov_5729.png

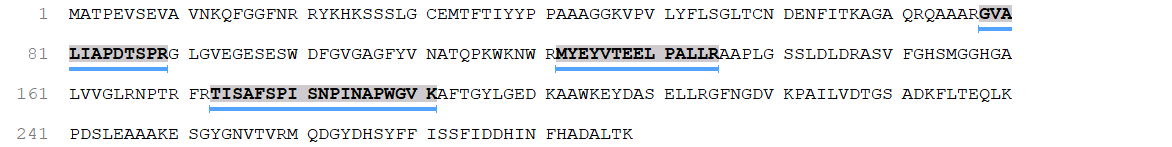

### cov_5730.png

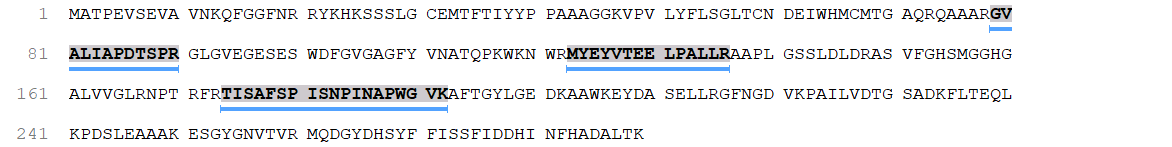

### cov_5731.png

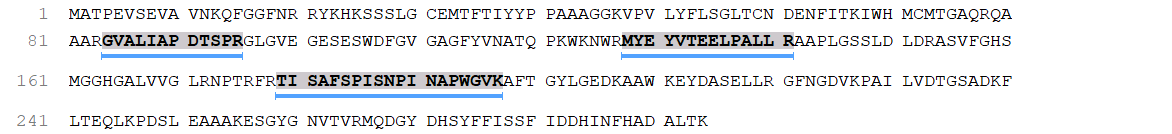

### cov_5732.png

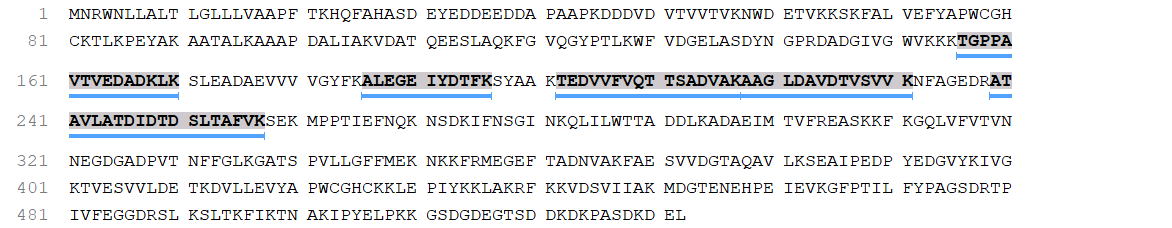

### cov_5733.png

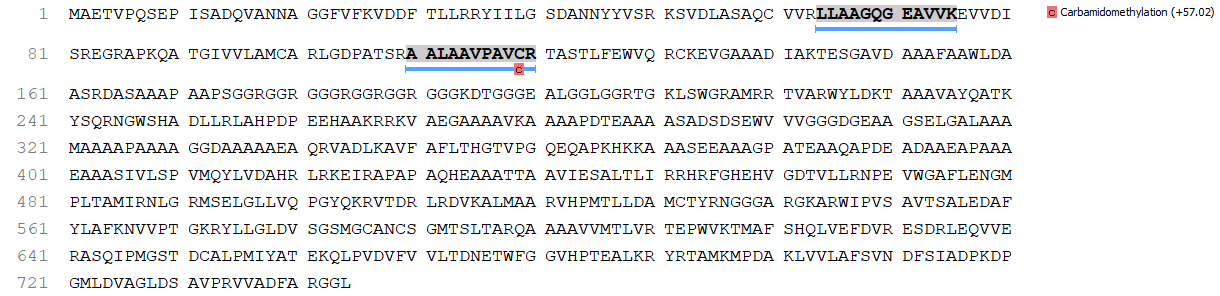

### cov_5734.png

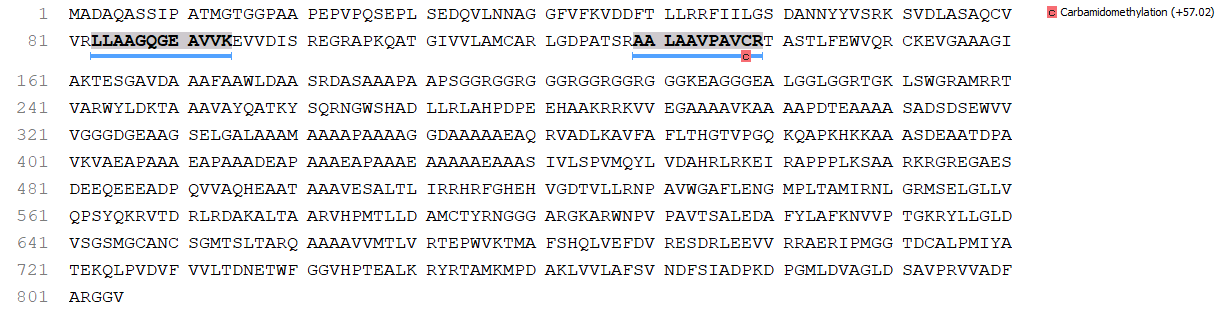

### cov_5735.png

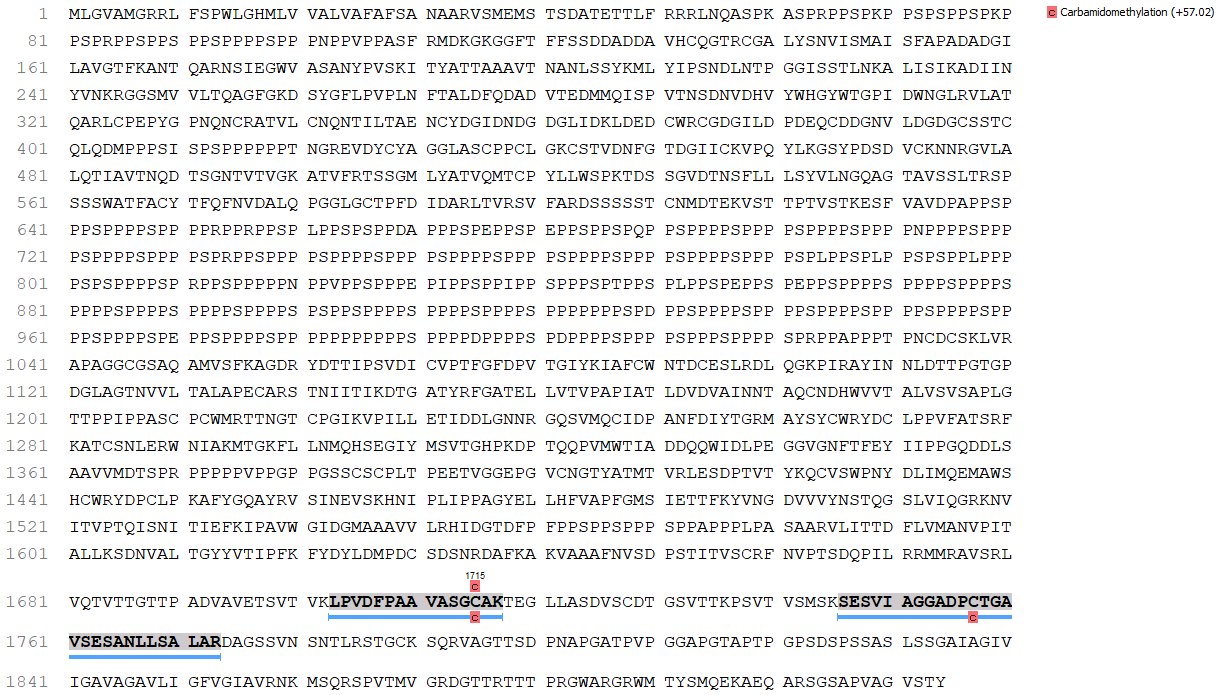

### cov_5736.png

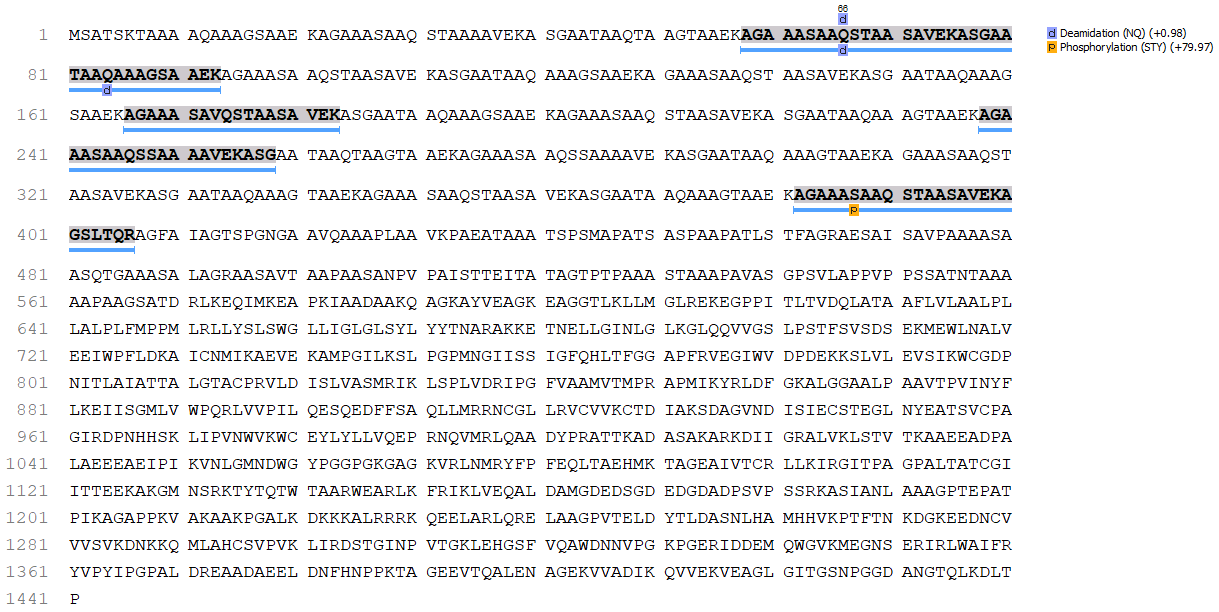

### cov_5737.png

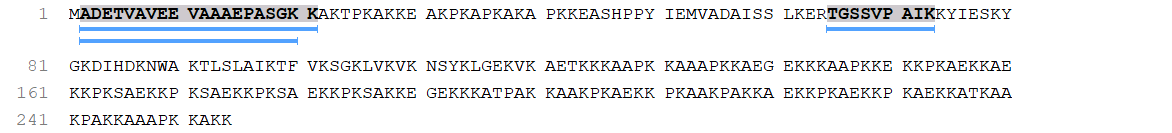

### cov_5738.png

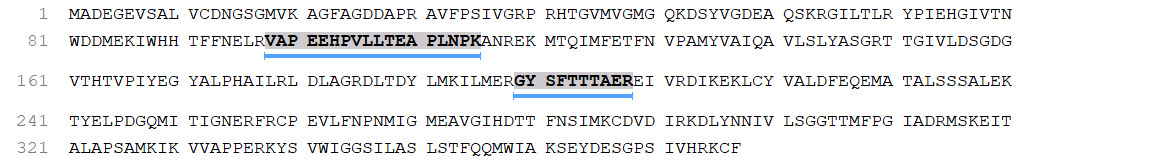

### cov_5739.png

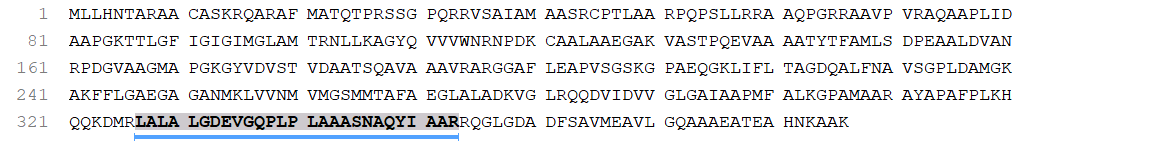

### cov_5740.png

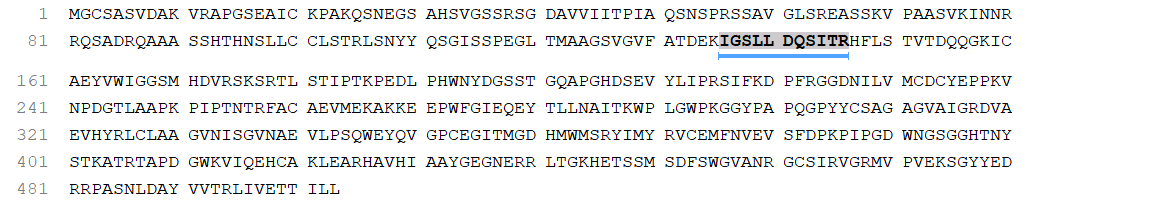

### cov_5741.png

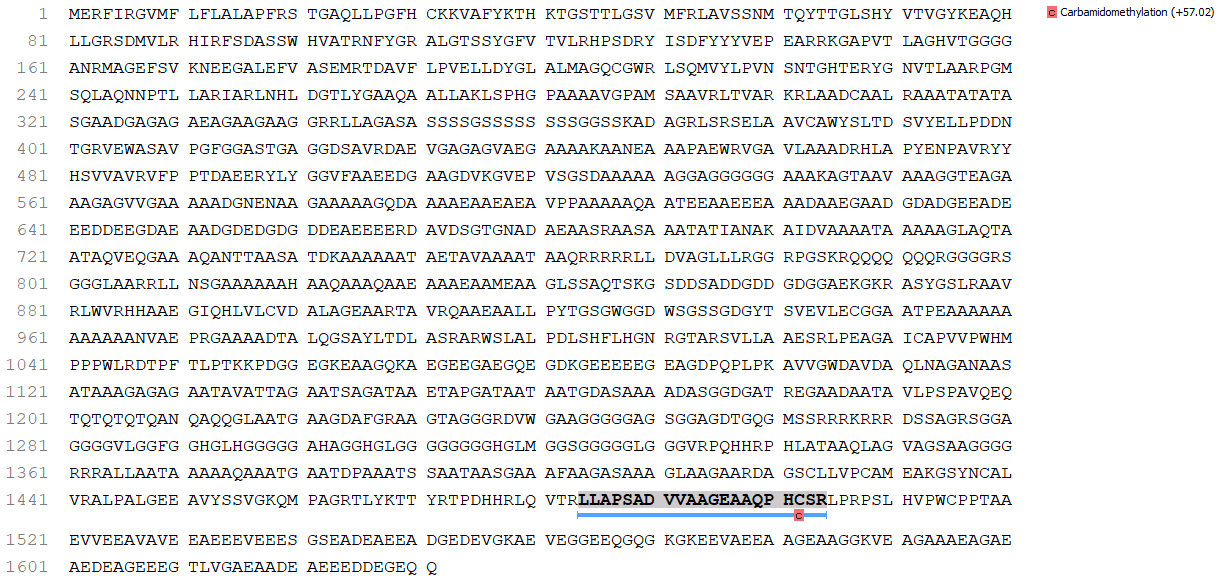

### cov_5743.png

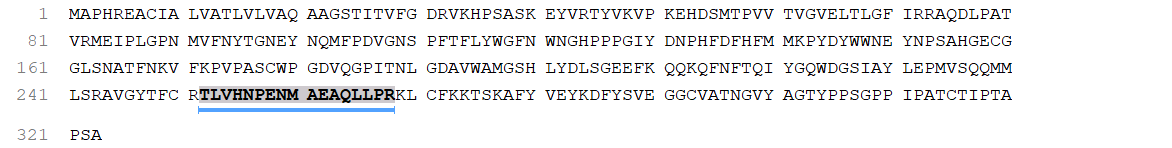

### cov_5744.png

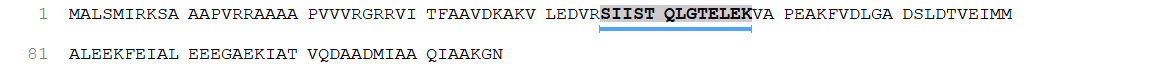

### cov_5745.png

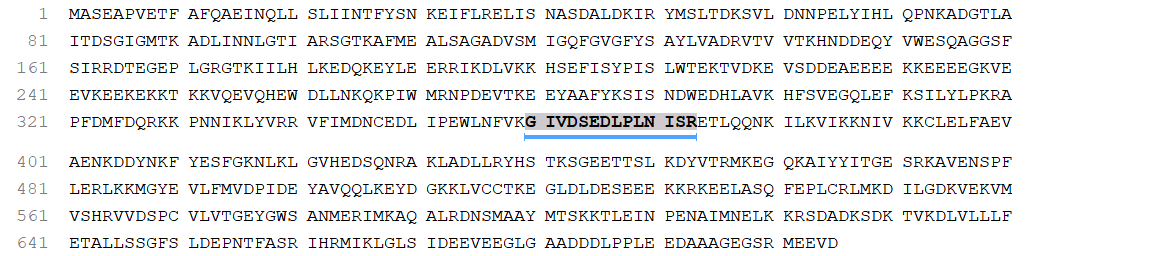

### cov_5746.png

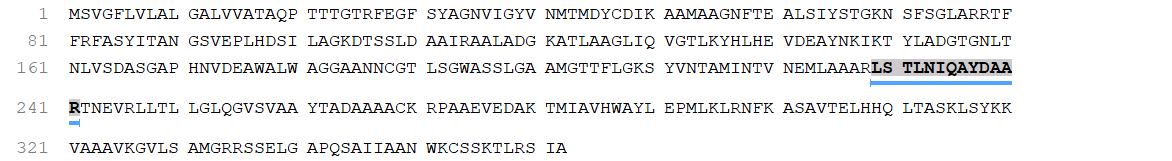

### cov_5747.png

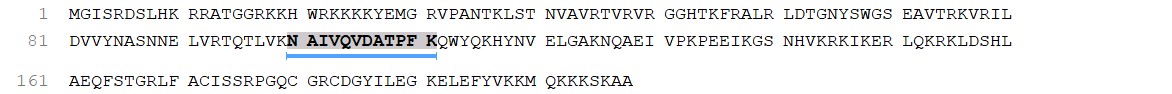

### DenovoFDRCurveFigure1960048827367275892.png

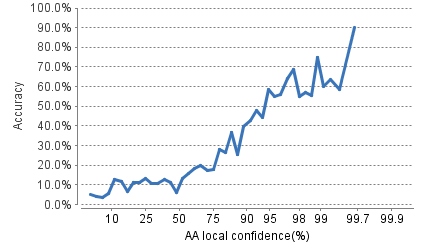

### ErrorCalibratedHistogram400687697950143094.png

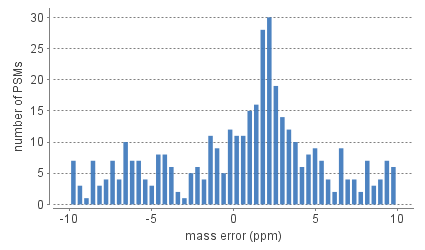

### ErrorPlotFigure3654833654735647081.png

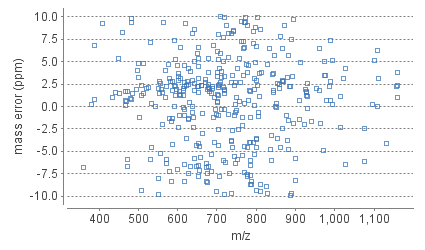

### FDRFigure1065976972405576925.png

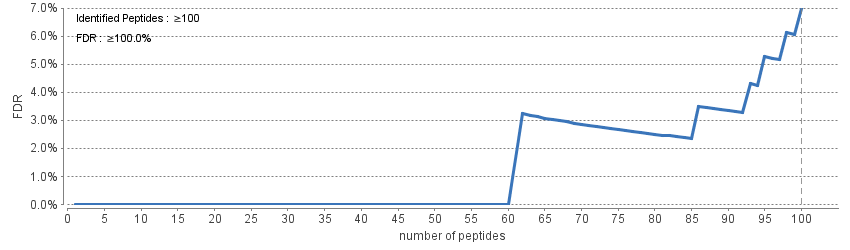

### FeatureIntensityDistributionHistogram7717308136477350083.png

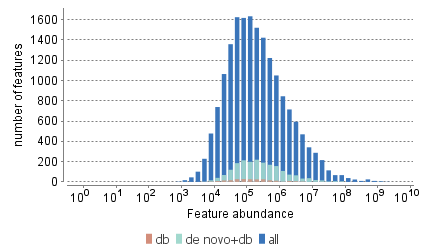

### FeatureMzHistogram5280862637344542542.png

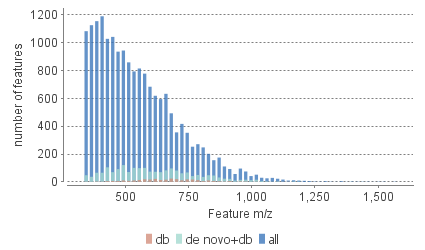

### FeatureRtHistogram7902428588763694180.png

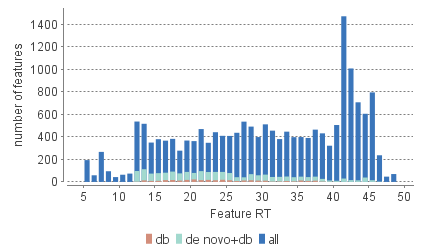

### q.png

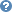

### ScoreHistogram8459785599774466543.png

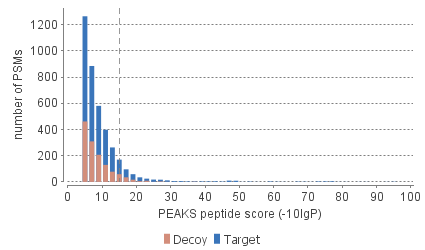

### ScorePlotFigure8859431838719737712.png

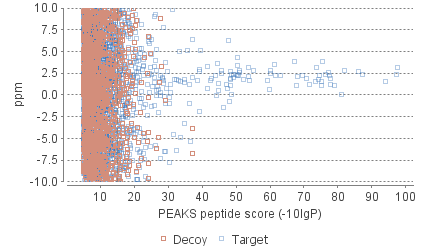
