## Supplementary Figures for "Efficient secretion of a plastic degrading enzyme from the green algae *Chlamydomonas reinhardtii*"

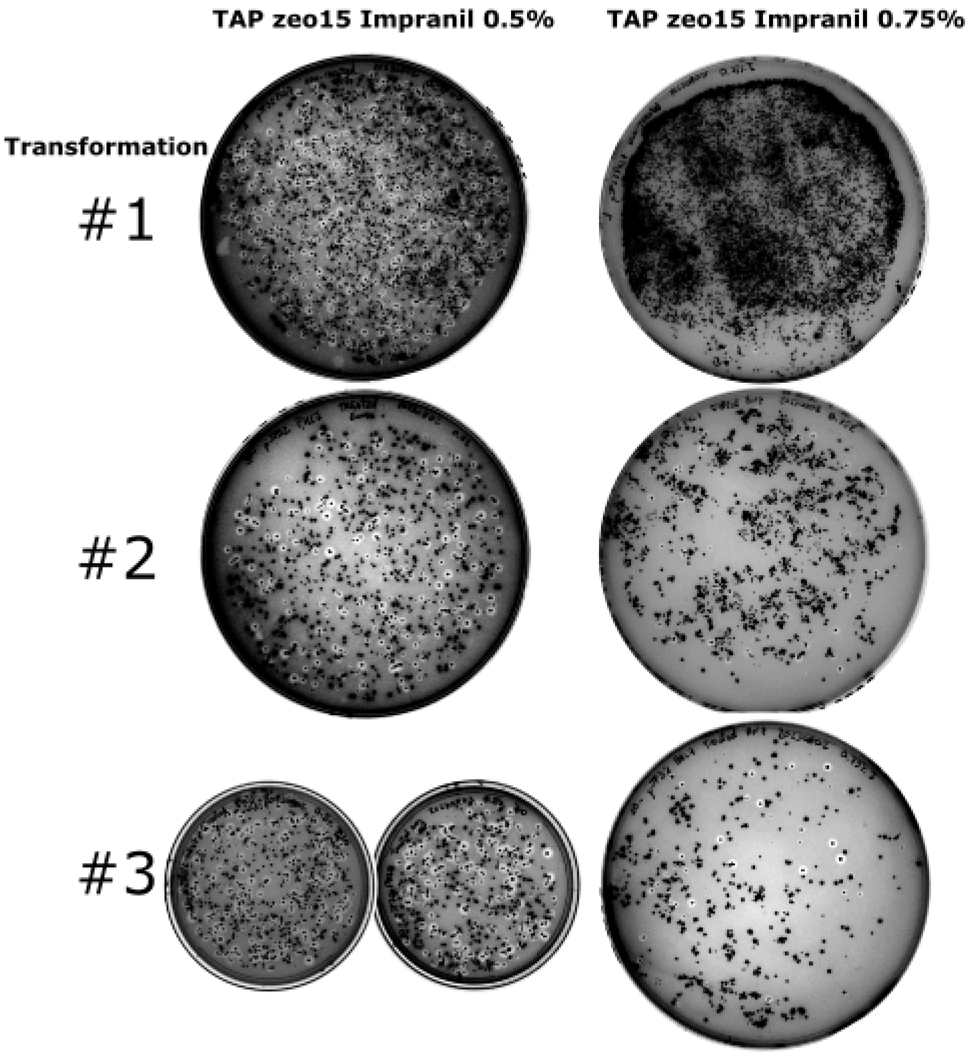


**Supplementary Figure. 1:** Selection/screening plates. Composition: TAP, agar 1.5% (w/v), zeocin 15 ug/mL, and 0.5 or 0.75% Impranil DLN (v/v). Transformants from three independent events with pJP32PHL7 vector harboring the secreting vector for PHL7. Each transformation was split equally among 0.5% and 0.75% plates.

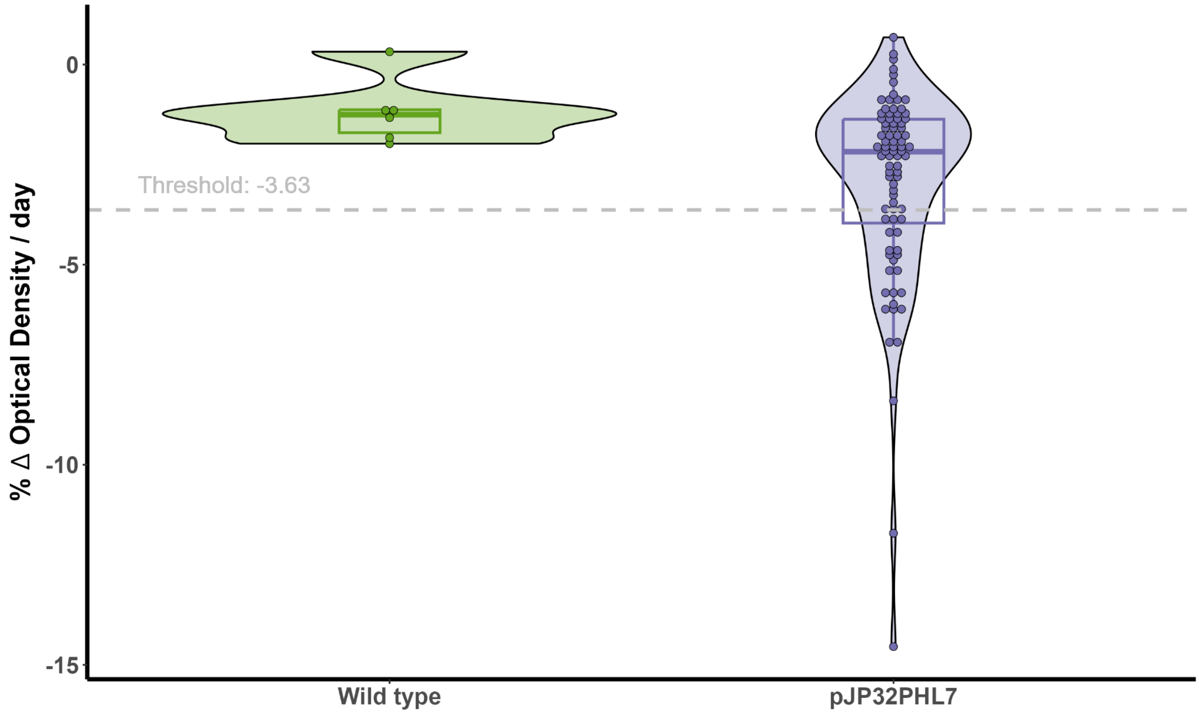


**Supplementary Figure 2: Screening of PHL7 secreting strains.** Comparison of activity based on Delta OD/d for Wild Type and pJP32PHL7 Strain. Violin plots display the entire distribution of values with overlaid boxplots. Individual data points are shown as dots. The dashed horizontal line represents the calculated threshold, determined as the wild-type data's mean minus three standard deviations. Values below this threshold suggest significantly higher activity compared to the wild type.

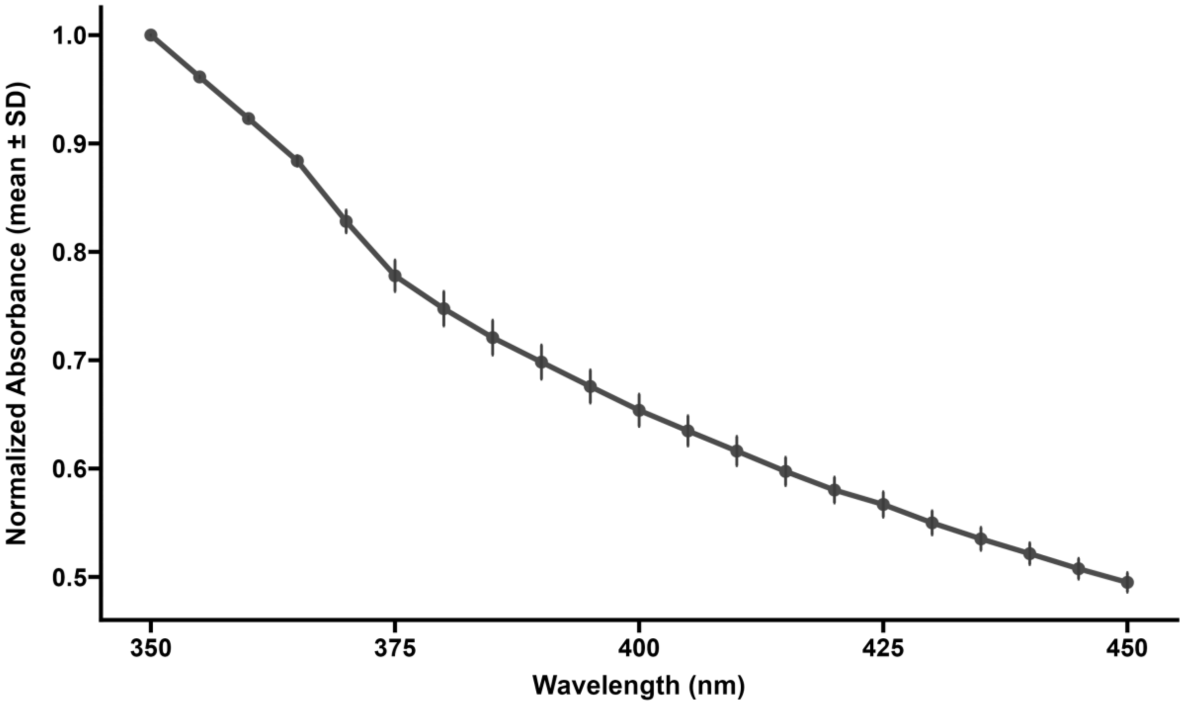


**Supp. Figure 3:** Normalized absorption spectra of 0.2% (w/v) agarose and 0.25% (v/v) Impranil® DLN in 100 mM phosphate buffer at pH 8, measured across wavelengths from 350 to 450 nm. The solid line represents the mean absorbance calculated from three independent measurements, each normalized to the absorbance at 350 nm. Error bars indicate the standard deviation (mean ± SD), reflecting the variability between the replicates.

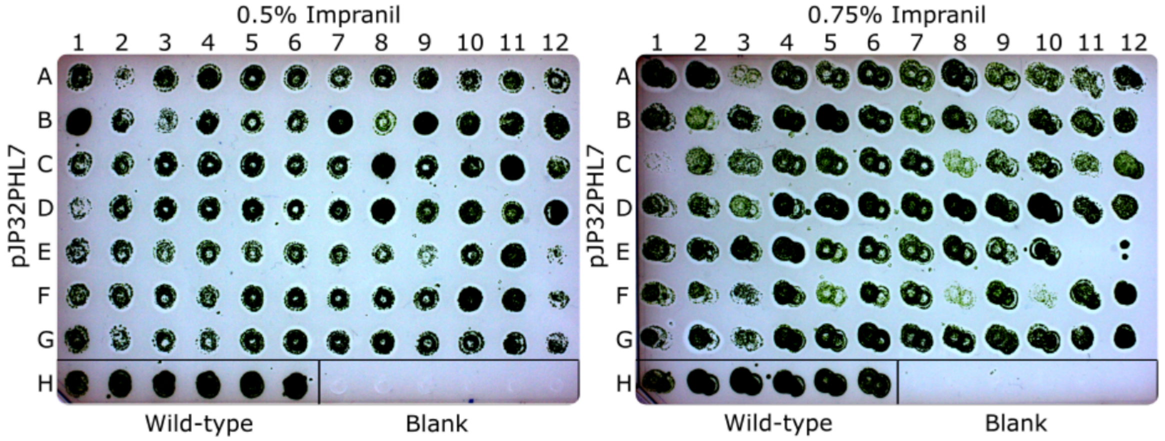


**Supplementary Figure 4**: Stamped cells from 0.5% Impranil plate (left) and 0.75% Impranil plate (right) onto agar plates containing zeocin 15µg/mL and 0.5% Impranil DLN. The 0.5% Impranil plate contains 81 colonies with halos and three without halos (positions B1, E12, and G1) due to the selection of colonies in a densely populated region, resulting in false positives. The 0.75% Impranil plate contains 83 colonies with halos and one missing colony in position E11 due to uneven stamping during the transfer that caused the blotter to miss capturing the specific colony in the designated well. Each plate contains six wild-type colonies (parental cc1690 strain) and six blanks at the most bottom row.

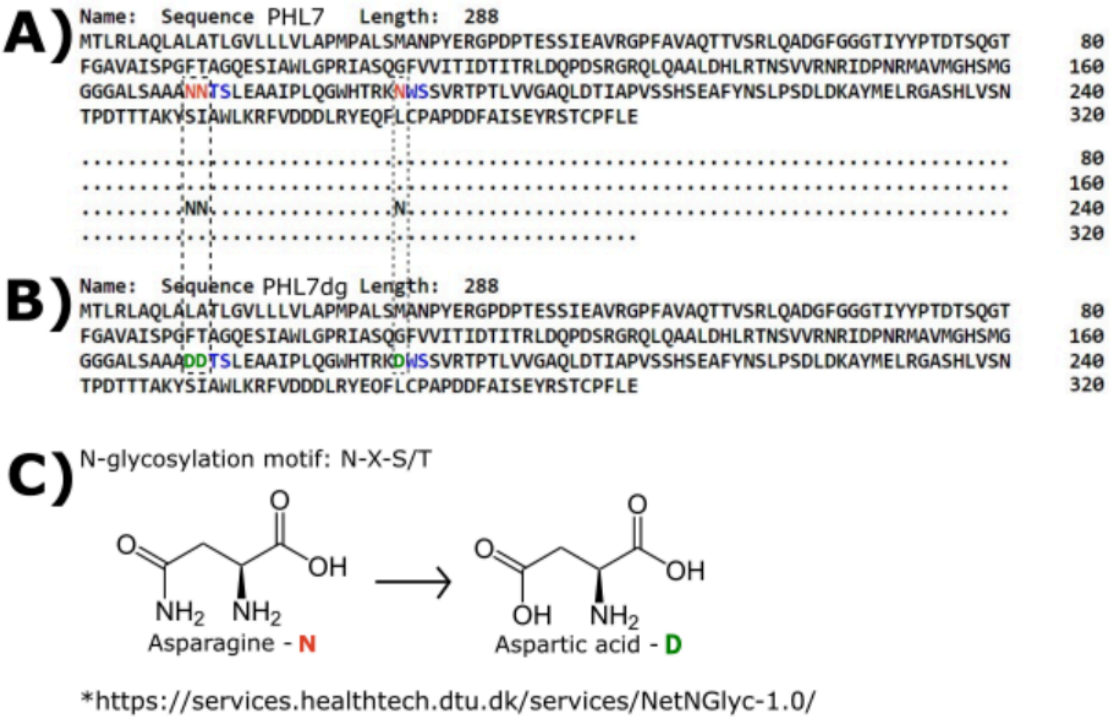


**Supplementary Figure 5**: (A) *In silico* analysis of the PHL7 protein sequence using NetN-Glyc-1.0 displayed three glycosylation sites (asparagine residues denoted as N in red text). From NetN-Glyc-1.0, N (for asparagine) represents glycosylation sites, and dots (•)

represent any other amino acid besides asparagine. (B) The protein sequence of PHL7dg displays the removal of glycosylation sites by replacing asparagine with aspartic acid (aspartic acid residues are denoted as D in green text). (C) Scheme displaying replacement of asparagine with aspartic acid to generate pJP32PHL7dg, demonstrating the difference between both amino acids, a hydroxyl group substituting an amine.

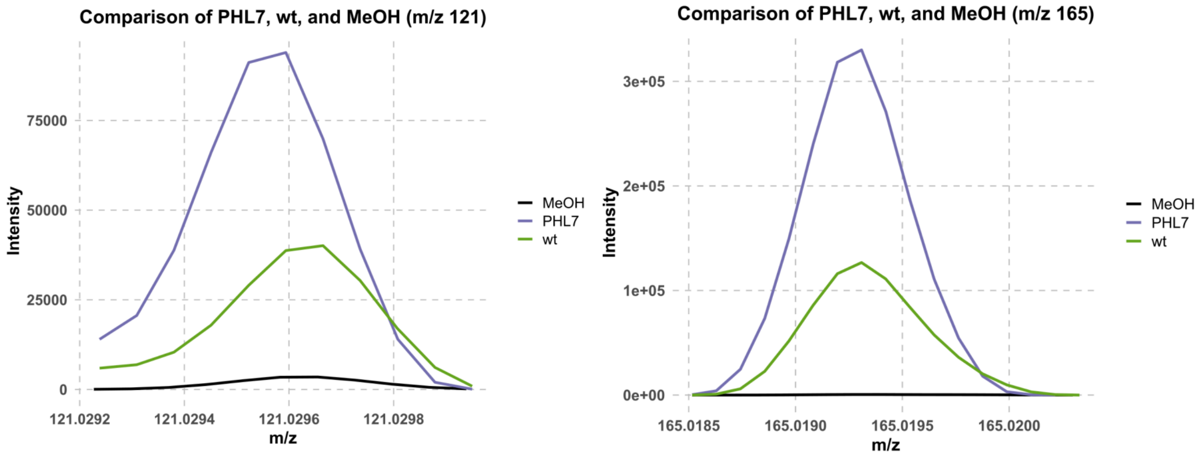


**Supplementary Figure 6:** Comparison of mass spectrometry signals for PHL7, wt, and MeOH at m/z 121 and 165. The graphs depict the intensity profiles for PHL7 (purple), wt (green), and MeOH (black) across two mass-to-charge (m/z) ranges: (A) m/z 121 and (B) m/z 165. The intensities represent the relative abundance of the detected ions corresponding to each sample. Dashed gray lines indicate the major grid, and the bold axis labels enhance clarity. The spectra highlight the variation in ion intensities between the biological strains (PHL7 and wt) and the blank (MeOH).

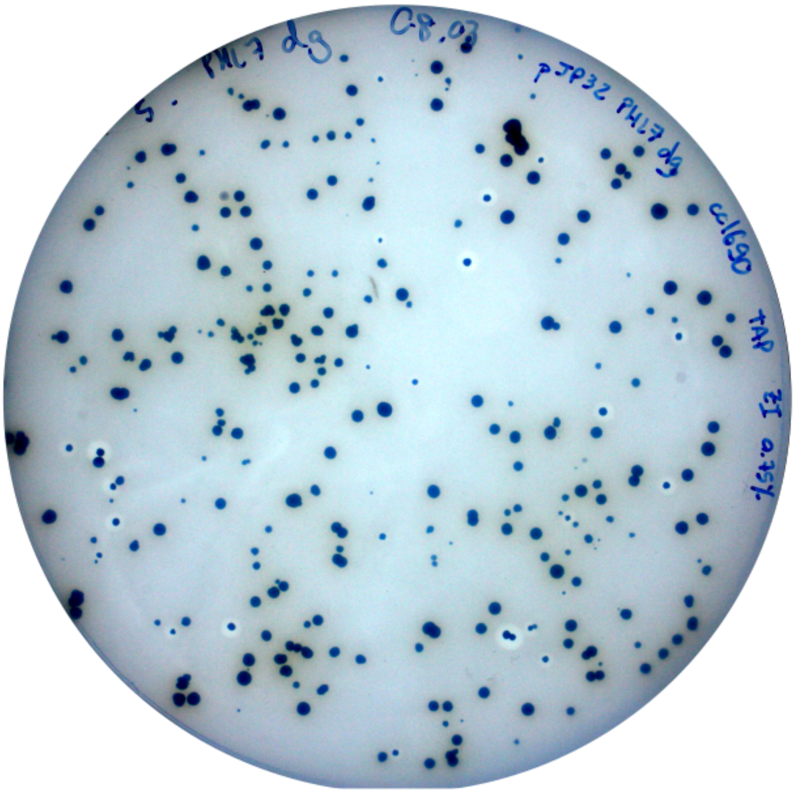


**Supplementary Figure 7**: TAP, agar 1.5% (w/v), zeocin 15µg/mL, and 0.75% Impranil DLN (v/v) plate containing transformants from a single transformation event with pJP32PHL7dg vector harboring the secretion of the deglycosylated version of PHL7dg.

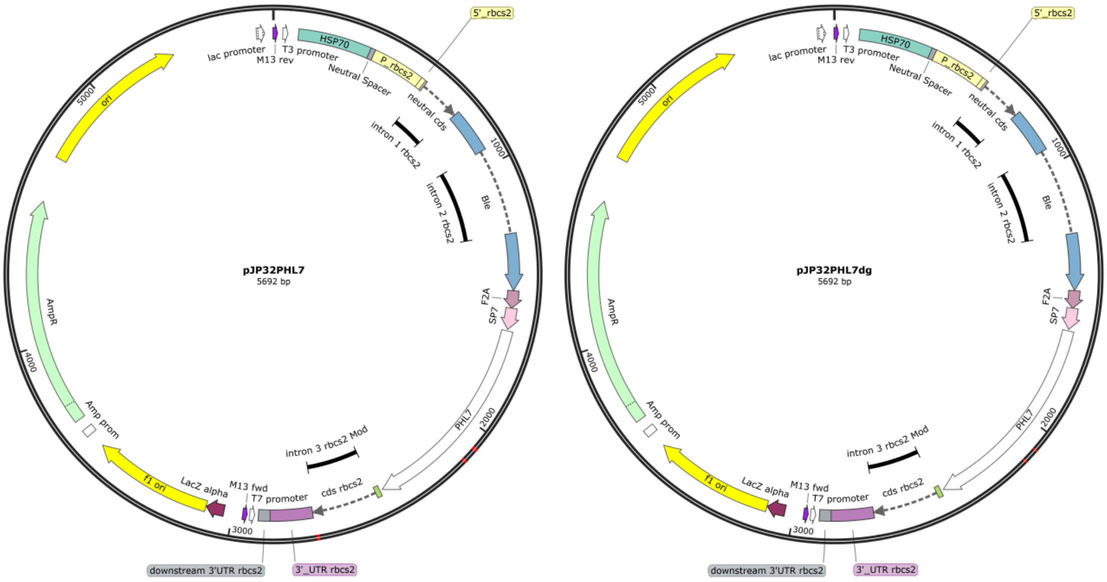


**Supplementary Figure 8:** Plasmid Maps of Vectors pJP32PHL7 and pJP32PHL7dg. PHL7 denotes the non-modified sequence of the plastic degrading enzyme. PHL7dg denotes the modified sequence replacing the asparagine (N) residues present in potential glycosylation sites by aspartic acid (D).
